# The genome of the oomycete *Peronosclerospora sorghi*, a cosmopolitan pathogen of maize and sorghum, is inflated with dispersed pseudogenes

**DOI:** 10.1101/2022.07.13.499355

**Authors:** Kyle Fletcher, Frank Martin, Thomas Isakeit, Keri Cavanaugh, Clint Magill, Richard Michelmore

## Abstract

Several species in the oomycete genus *Peronosclerospora* cause downy mildew on maize and can result in significant yield losses in Asia. Biosurveillance of these pathogens is a high priority to prevent epidemics on maize in the US and consequent damage to the US economy. The unresolved taxonomy and dearth of molecular resources for *Peronosclerospora* spp. hinder these efforts. *P. sorghi* is a pathogen of sorghum and maize with a global distribution, for which limited diversity has been detected in the southern USA. We characterized the genome, transcriptome, and mitogenome of an isolate, representing the US pathotype 6. The highly homozygous genome was assembled using 10x Genomics linked reads and scaffolded using Hi-C into 13 chromosomes. The total assembled length was 319.6 Mb—larger than any other oomycete previously assembled. The mitogenome was 38 kb, similar in size to other oomycetes, although it had a unique gene order. Nearly 20,000 genes were annotated in the nuclear genome, more than described for other downy mildew causing oomycetes. The 13 chromosomes of *P. sorghi* were highly syntenic with the 17 chromosomes of *Peronospora effusa* with conserved centromeric regions and distinct chromosomal fusions. The increased assembly size and gene count of *P. sorghi* is due to extensive retrotransposition, resulting in putative pseudogenization. Ancestral genes had higher transcript abundance and were enriched for differential expression. This study provides foundational resources for analysis of *Peronosclerospora* and comparisons to other oomycete genera. Further genomic studies of global *Peronosclerospora* spp. will determine the suitability of the mitogenome, ancestral genes, and putative pseudogenes for marker development and taxonomic relationships.

## Introduction

Oomycetes are destructive pathogens of plants and animals that have caused several historical and contemporary epidemics [1]. The genus *Peronosclerospora* contains multiple, poorly resolved species of oomycetes, which cause downy mildews on graminaceous plants including grain crops and sugarcane [2–4]. Maize is a host to many of these species and downy mildew outbreaks can result in significant yield losses [4–7]. Maize production plays a major role in the economy of the US and was valued at over $61 billion dollars by the USDA National Agricultural Statistics Service for 2020. Consequently, measures for surveillance and strategies for response to species that cause major downy mildew epidemics on maize elsewhere, but are not established in the US, are desirable. This threat to US agriculture resulted in *P. philippinensis* being placed on the select agent list [3]; however, the taxonomic relationships of *Peronosclerospora* spp., including *P. philippinensis*, is unresolved [3, 4, 8, 9]. Therefore, it may be prudent to evaluate whether the introduction of any species that causes downy mildew on maize could threaten US production.

Currently, the only *Peronosclerospora* species reported in the US is *P. sorghi,* which causes downy mildew on maize and sorghum [4]. *P. sorghi* can complete its asexual cycle on both hosts. The species is homothallic; however, oospores generated during the sexual cycle have only been reported in sorghum in the US [3, 10]. Previously, *P. sorghi* has been managed through crop rotation, resistant hybrids, and seed treatment with metalaxyl (methyl N- (2, 6-dimethylphenyl)-N-(methoxyacetyl)-DL-alaninate). Outbreaks of metalaxyl resistant downy mildew on sorghum occurred in Texas in 2001 and 2002 [3, 11], and in subsequent years, associated with sorghum monoculture. After growers in Texas switched to monoculturing with resistant hybrids in response to metalaxyl resistance, a new pathotype, 6, was detected [12], which was also resistant to metalaxyl. Consequently, constant monitoring of this pathogen is required. At present, six pathotypes have been described from the Americas, based on virulence phenotypes on 10 differential inbred lines of sorghum [13]. Pathotypes 1 to 3 and 6 were identified from Texas, pathotype 4 from Brazil, and pathotype 5 from Honduras [13]. The line SC155 was initially thought to differentiate pathotype 6 from 3 [12], but this was not subsequently confirmed [13]. Pathotype 6 overcomes resistance of proprietary Pioneer brand germplasm resistant to pathoype 3 [13]. American cultivars of maize are highly resistant to *P. sorghi* but are reported to be susceptible to other *Peronosclerospora* spp. found in Asia [3, 14]. The variability of *P. sorghi* in the USA is believed to be low [13]; however, this has not been characterized at the genomic level.

Genomic resources are foundational for accurately defining the variation present in populations of a pathogen, conducting bio-surveillance, and identifying vulnerabilities and opportunities that can be utilized for disease management. Recently, two 17-chromosome-scale genome assemblies have been reported for the oomycetes *Bremia lactucae* and *Peronospora effusa,* which cause lettuce and spinach downy mildew, respectively [15, 16]. Despite being taxonomically distant, these oomycetes share a high degree of synteny. Because downy mildew oomycetes are polyphyletic [16–20], this high level of synteny suggests that similar or derived chromosome architectures are expected for all downy mildew causing oomycetes, and several clades of *Phytophthora* spp. [16].

The present study describes the chromosome-scale genome assembly of *P. sorghi*. This is the first genome assembly for any species in the genus *Peronosclerospora* and will be a foundational resource for surveillance of this pathogen globally. *P. sorghi* has the largest genome of any oomycete sequenced to date. Comparative genomics revealed a unique genomic architecture with 13 chromosomes derived from the ancestral architecture. In addition, the transcriptome of *P. sorghi* was characterized to generate reliable gene models and investigate differential gene expression of infected leaves pre- and post-sporulation of the pathogen.

## Results

Our initial genome assembly of *P. sorghi* contained 342 Mb of nucleotide sequence that was scaffolded using 10x Genomics linked and Hi-C reads into 12 large superscaffolds. This genome assembly was manually corrected by inspecting the Hi-C contact matrix (Fig. S1). Centromeres were identified on each superscaffold by short distance cis contacts, which resulted in crosses on the Hi-C plot (Fig. 1A). The initial Hi-C plot indicated that there were three mis-assembled superscaffolds (Supplementary Fig 1). The arms of Chromosome (Chr.) 11 were joined at the telomeric regions not the centromere; the chromosomes arms were therefore reoriented and scaffolded over the centromere. The two arms of Chr. 8 were split in the initial assembly and joined to Chr. 1 and Chr. 2; the false joins were evident in the Hi-C plot because one end of each scaffold was enriched for short distance cis contacts in addition to those at the centromere. The two arms of Chr. 8 were separated and then scaffolded together. Hi-C reads were remapped to validate the revised assembly (Fig. 1A). These manual corrections were subsequently confirmed by synteny analysis (see below). Full assembly statistics are available in Supplementary Table S1.

**Fig. 1.**
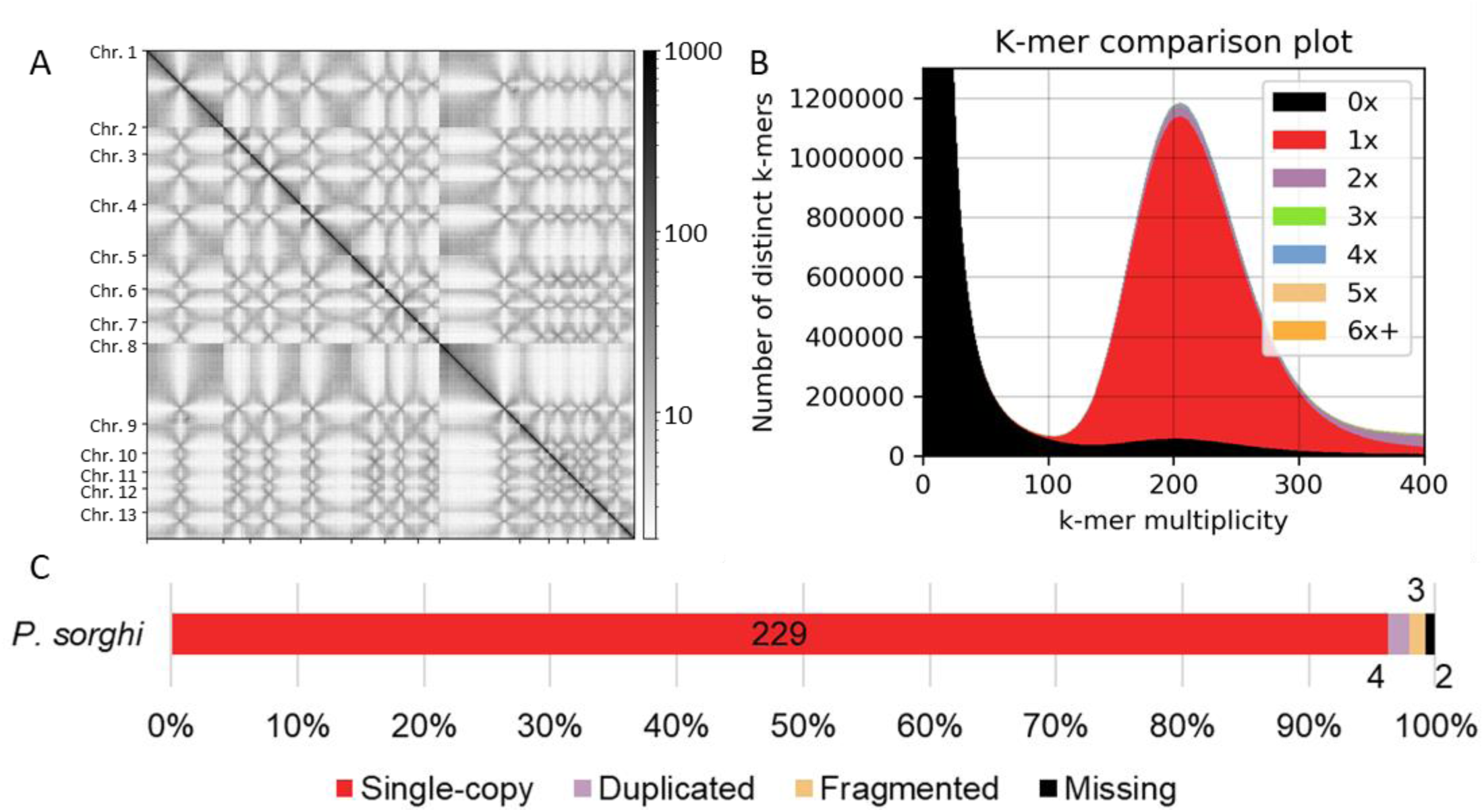
The genome assembly of *Peronosclerospora sorghi*. A) Hi-C contact matrix of 13 chromosome-scale scaffolds. The strong diagonal reflects the high contact frequency between physically close sequences indicating their correct linear order along each chromosome-scale scaffold. Cross patterns along the x- and y-planes are indicative of high frequencies of trans contacts between centromeres and are likely due to the Rabl-like chromosome configurations. B) Visualization of assembly completeness using *k*-mer inclusion demonstrating that the majority of single-copy *k*-mers are present in the assembly only once (red). The Poisson distribution is consistent with high homozygosity in the genome of *P. sorghi*. C) Stacked bar-chart showing the completeness of the genome assembly as calculated by BUSCO. The color scheme is similar to B so that single-copy BUSCOs are depicted as red. The annotated numbers indicate gene count. The total number of BUSCOs in the protist database was 234.

Our final assembly contained thirteen chromosomes with a total scaffolded length of 283.5 Mb; there were 8,767 unplaced small scaffolds (Chr. 0) with an average length of 6.7 kb and a total length of 58.3 Mb (Table 1). Analysis of *k*-mers determined that the genome of *P. sorghi* was highly homozygous because only a single Poisson distribution was present (Fig. 1B). In addition, the majority of single-copy *k*-mers were present in the genome assembly. Benchmarking with BUSCO demonstrated that the assembly was 97.9% complete with little duplication (Fig. 1C), supporting a high level of completeness. All but one BUSCO was predicted from the assembled chromosomes of *P. sorghi*. The single BUSCO gene located on Chr. 0 was scored as duplicate and occurred three times on three separate scaffolds.

**Table 1.** Assembly statistics for the genome of *Peronosclerospora sorghi*.

| *P. sorghi* sequence | Length | Primogenitors<sup>1</sup> | Total Gaps | Percent Gaps | AT content | Genes | Genes/Mb | Repeat-masked bases | Percent Repeats |
| --- | --- | --- | --- | --- | --- | --- | --- | --- | --- |
| Chr. 1 | 44403666 | Chr. 1 | 3569330 | 8.04% | 51.0% | 2755 | 62.04 | 34200987 | 77.02% |
| Chr. 2 | 15566821 | Chr. 2 | 1090110 | 7.00% | 51.6% | 999 | 64.17 | 11673129 | 74.99% |
| Chr. 3 | 29316722 | Chr. 3, Chr. 5 | 2441500 | 8.33% | 51.2% | 1864 | 63.58 | 22246130 | 75.88% |
| Chr. 4 | 29556142 | Chr. 4, Chr. 9 | 2216080 | 7.50% | 51.4% | 1960 | 66.31 | 22262399 | 75.32% |
| Chr. 5 | 19437638 | Chr. 6 | 1668350 | 8.58% | 51.6% | 1252 | 64.41 | 14321970 | 73.68% |
| Chr. 6 | 19380668 | Chr. 7, Chr. 13 | 1459800 | 7.53% | 51.3% | 1319 | 68.06 | 14626688 | 75.47% |
| Chr. 7 | 12369872 | Chr. 8 | 1079370 | 8.73% | 51.9% | 905 | 73.16 | 9186674 | 74.27% |
| Chr. 8 | 46690312 | Chr. 10, Chr. 12 | 3572980 | 7.65% | 51.0% | 2681 | 57.42 | 35816060 | 76.71% |
| Chr. 9 | 16992261 | Chr. 11 | 1138780 | 6.70% | 51.5% | 1044 | 61.44 | 13069339 | 76.91% |
| Chr. 10 | 10915484 | Chr. 14 | 752700 | 6.90% | 51.8% | 750 | 68.71 | 8382571 | 76.80% |
| Chr. 11 | 9655334 | Chr. 15 | 706550 | 7.32% | 51.8% | 569 | 58.93 | 7357013 | 76.20% |
| Chr. 12 | 13731139 | Chr. 16 | 909030 | 6.62% | 51.5% | 1055 | 76.83 | 10492191 | 76.41% |
| Chr. 13 | 15489776 | Chr. 17 | 1128150 | 7.28% | 51.7% | 1109 | 71.60 | 11615788 | 74.99% |
| Chr. 0 | 58304996 | n/a | 655880 | 1.12% | 50.0% | 1386 | 23.77 | 53971640 | 92.57% |

<sup>1</sup>Primogenitors retain numbering from the chromosomes of *P. effusa* GCA\_021491665.1.

A subset of single-copy orthologs identified in *P. sorghi* by BUSCO were added to a previously reported phylogenetic analysis using 18 BUSCO genes found to be single-copy across 31 genome assemblies [16]. This determined that the two graminicolous downy mildew assemblies, those of *P. sorghi* and *Sclerospora graminicola*, formed a monophyletic clade within downy mildew clade 2 (Fig. 2). This phylogeny supports that these two graminicolous downy mildews share a common biotrophic ancestor with other genera of downy mildews in this clade, including *Peronospora*, *Pseudoperonospora*, and *Hyaloperonospora* [16–20].

**Fig. 2.**
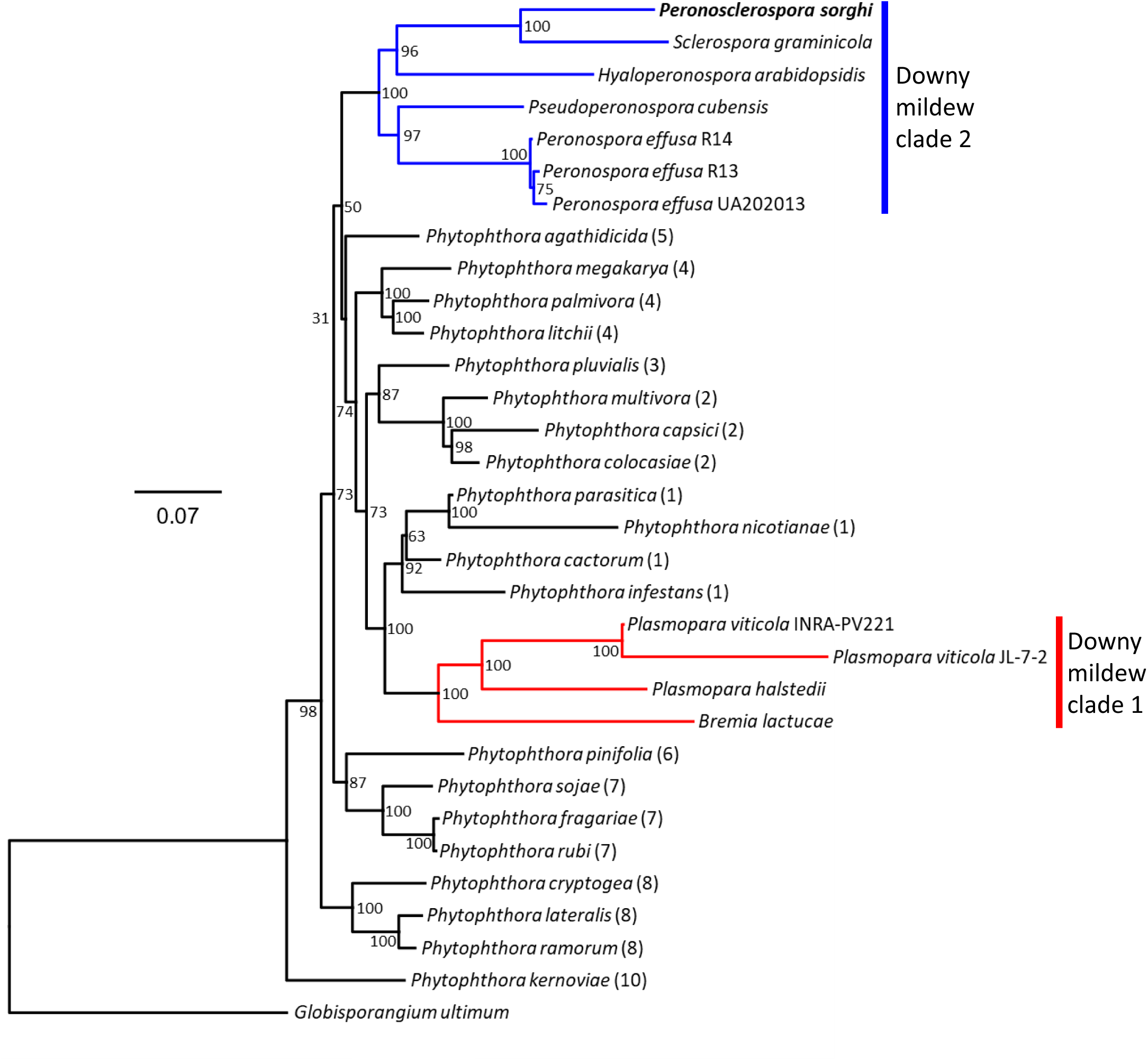
Phylogenetic analysis of *Peronosclerospora sorghi*. Maximum likelihood phylogenetic tree calculated from the concatenated alignment of 18 BUSCO proteins surveyed in 32 assemblies of 28 oomycete species. All species analyzed belong to the Peronosporaceae, except *Globisporangium ultimum* (Pythiaceae), which was used as an outgroup. Colored branches indicate the two downy mildew clades, consistent with polyphyly of the downy mildews. *P. sorghi* was assigned to downy mildew clade 2 (blue). The scale indicates the mean number of amino acid substitutions per site. Branch support was calculated from 1,000 bootstraps. Most of the data for this figure is the same as used in Fletcher *et al*. 2022, Figure 5.

Over 84% (269 Mb) of the assembled sequence was repetitive. The most common repeat elements were long terminal-repeat retrotransposons, which comprised 235 Mb of the assembly. The density of repeats was higher in Chr. 0 scaffolds compared to the chromosome-scale scaffolds (Table 1). Repeat density in 100 kb windows along the chromosomal scaffolds ranged from 0.29 to 1.00, and the average repeat density of windows was 0.82 (Fig. 3).

**Fig. 3.**
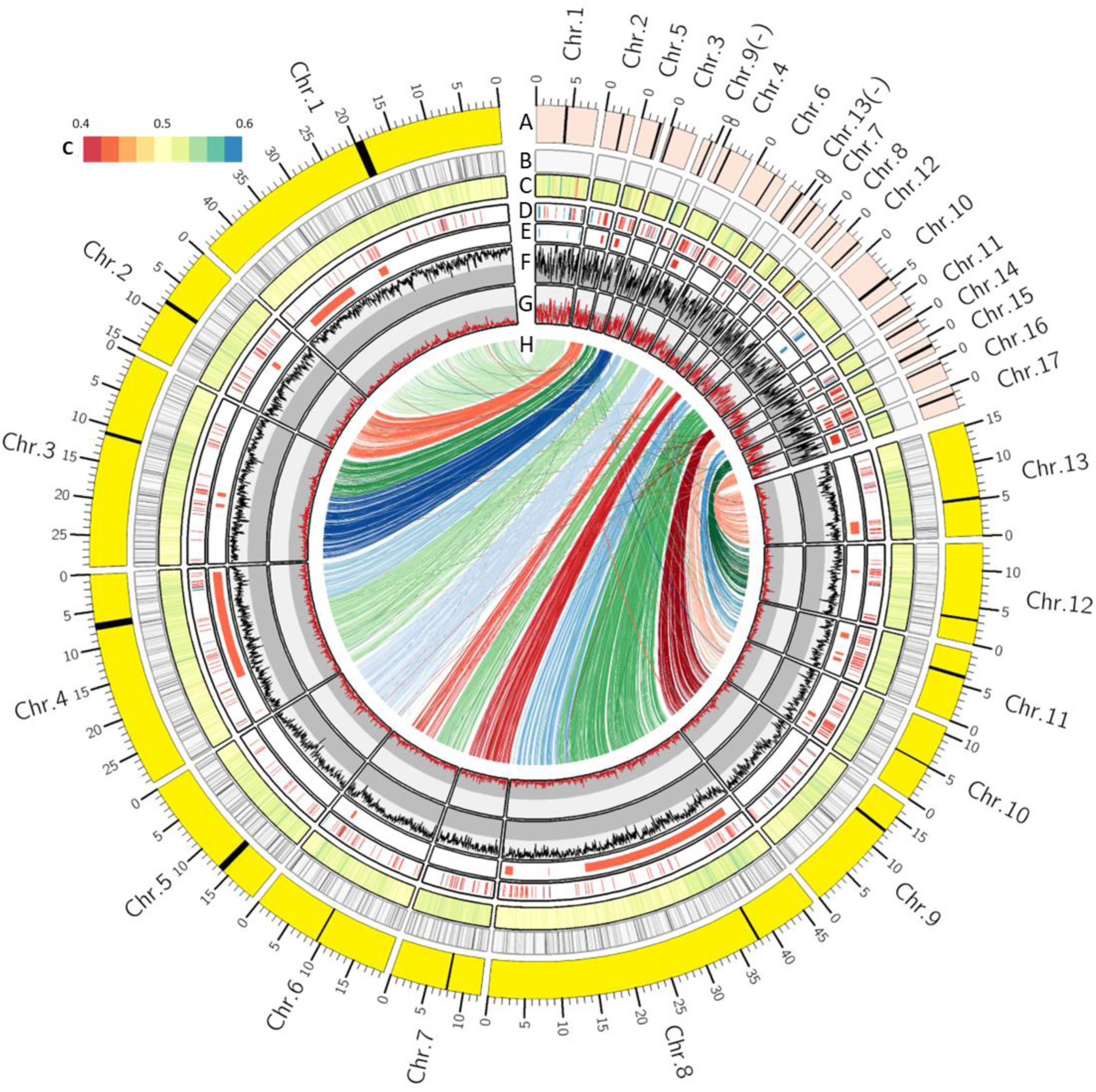
Comparison of the genome architectures of *Perononsclerospora sorghi* and *Peronospora effusa*. A) Scaled chromosomes showing lengths for *P. sorghi* (yellow) and *P. effusa* (red). Chromosome numbers of *P. sorghi* were ordered based on their designation in *P. effusa*. The scale shows the sizes in Mb. Black bars indicate the putative positions of centromeres (see Fig. 6). B) Distribution of gaps between contigs in the genome assemblies of *P. effusa* (1) and *P. sorghi* (9,804). C) Heatmap of AT content, ranging from 0.4 to 0.6. D) Distribution of annotated effectors. Blue bars represent genes encoding crinklers (98 in *P. effusa* and only 32 in *P. sorghi*). Red bars indicate genes encoding RXLR-like effectors (209 in *P. effusa* and 428 in *P. sorghi*). E) Effector clusters called in the genomes of *P. effusa* and *P. sorghi*. Clusters were called if three or more effectors on the same chromosome had high peptide identity (also see Supplementary Fig. S3). F) Repeat density in 100 kb windows with a 25 kb step. Split in the grey background indicates 0.5. G) Gene density in 100 kb windows with a 25 kb step. Split in the grey background indicates 0.5. H) High levels of synteny depicted by 3,476 links that indicate the genomic positions of single copy orthologs in the genomes of *P. effusa* and *P. sorghi*. The color of each link reflects the chromosome of *P. effusa* that the ortholog is encoded on.

The mitochondrial genome had a read depth of 2,359x and was assembled into a 38,497 bp circular molecule with a GC content of 22.4% (Fig. S2; GenBank accessions #####). Coding regions constituted 90.3% of the genome with 6.0% of this total representing hypothetical genes. A total of 35 known genes (encoding 18 respiratory chain proteins, 16 ribosomal proteins, and an import protein, *ymf16* of the *sec*Y-independent pathway), the *rnl* and *rns*, and 25 tRNA genes encoding for 19 amino acids were present. In addition, there were five hypothetical proteins (*ymf98*, *ymf99, ymf100, ymf101* and *orf32*) in common with other oomycete mitochondrial genomes [19, 21] and one putative ORF (*orf161*) that was present in *P. sorghi* downstream from the *cox1* gene (Fig. 4). BLAST queries to GenBank identified no significant sequence similarities for *orf161.* The mitochondrial gene order in *P. sorghi* was similar to *Peronospora* species with some exceptions (Fig. 4). There was a large inversion in the mitogenome of *P. sorghi*, relative to all other oomycete taxa. This inversion split the mitogenome into two parts representing approximately 71% and 29% of the sequence respectively. The larger portion encoded 31 genes plus four putative orfs (*ymf100* to *cob*) and spanned a region with a predominantly conserved gene order across several downy mildew taxa and *Phytophthora* species. Another difference in gene order for *P. sorghi* was the placement of the *nad5*-*nad6*-*trnR* genes, which were in the same location but inverted and encoded on the opposite strand rather than downstream from the *atp1* gene (Fig. 4). The inversion of genes *cob* to *atp1* (highlighted blue in Fig. 4) common to *B. lactucae*, *Plasmopara viticola,* and *P. infestans*, relative to the other taxa sampled is consistent with polyphyly of oomycetes which cause downy mildew diseases [17, 19].

**Fig. 4.**
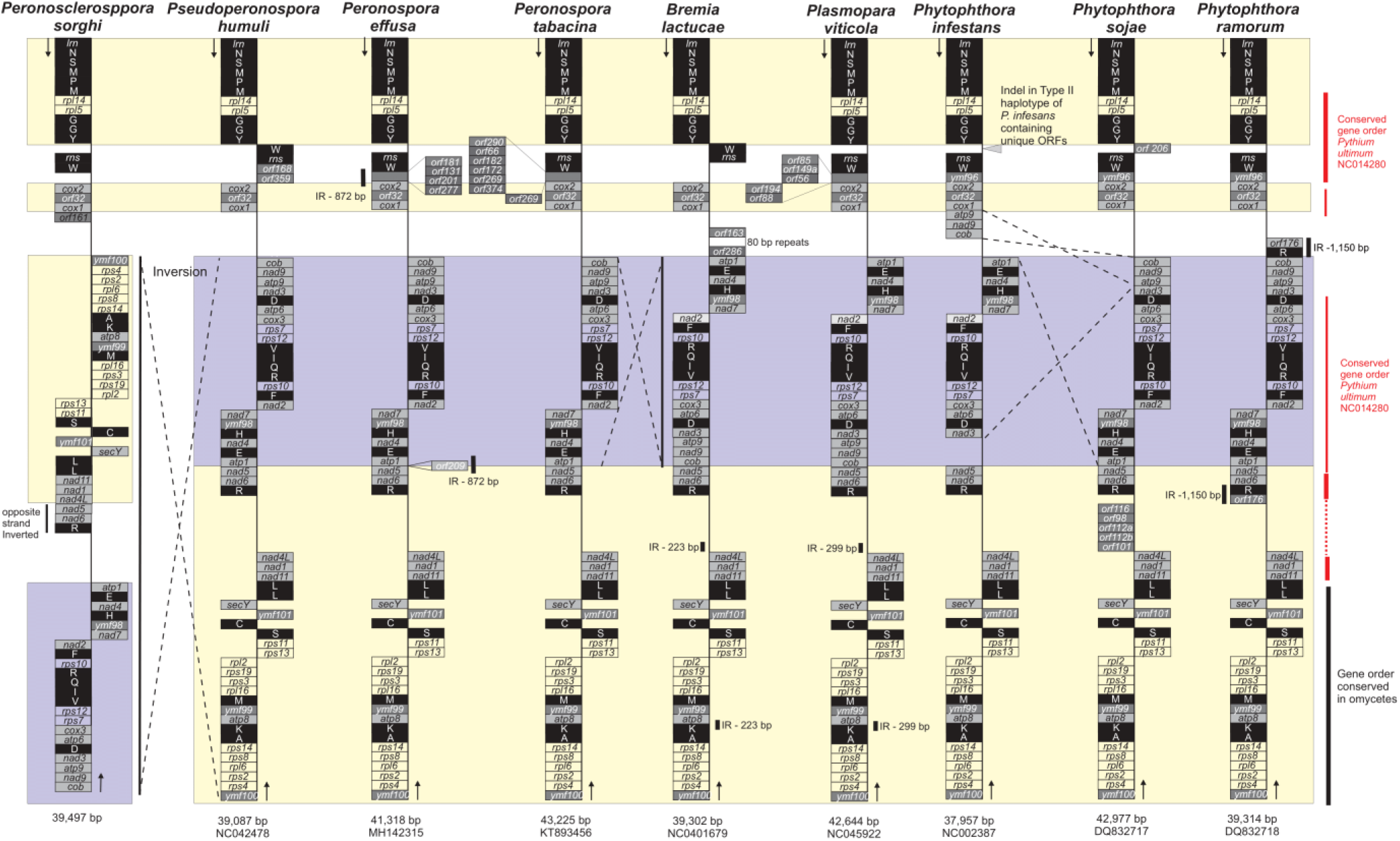
Comparison of mitochondrial gene order between six genera of the Peronosporaceae. Assemblies of nine species from six genera were analysed. The species are ordered to be consistent with phylogenetics (Fig. 2). Length and NCBI accession of each sequence is at the bottom. All genomes were oriented from the gene encoding the large subunit of the mitochondrial ribosome (*lrn*). Approximate locations of genes are indicated in boxes (not scaled to gene size). Boxes on the left indicate the gene was predicted on the top strand. Boxes on the right indicate the gene was predicted on the bottom (reverse-complemented) strand. Major inversions between assemblies of different species are highlighted by dashed lines. Repeats are annotated by black vertical bars. Colored backgrounds indicate regions of conserved gene order across taxa.

The transcriptome of *P. sorghi* isolate P6 was characterized using RNA extracted from lesions on leaves of *Sorghum bicolor* harvested both prior to and post sporulation. The transcriptome assembly contained 365,281 sequences totaling 482 Mb. Aligning transcripts to the genomes of *P. sorghi* and *S. bicolor* assigned 73,563 (20%) of the transcripts to *P. sorghi* (Fig. 5). The total length of transcripts assigned to *P. sorghi* was 98 Mb; the largest transcript was 16.8 kb. The N_50_ of the *P. sorghi* transcriptome was 2.2 kb, the mean length was 1.3 kb. Over 95% of BUSCO genes were detected in the transcriptome with 49.1% duplicated. Over 10% of the *P. sorghi* transcriptome was derived from repetitive genomic sequences, 184 transcripts of which encoded reverse transcriptase (PF07727), ranking it the 12^th^ most frequently identified domain encoded in the transcriptome. Most of the assembled transcriptome (62%) aligned to the host assembly of *S. bicolor* (225,170 transcripts totaling 356.8 Mb). The largest *S. bicolor* transcript was 18.4 kb, the transcript N_50_ was 2.6 kb, and the mean transcript length was 1.6 kb. Over 93% of BUSCO genes were detected in the transcriptome of the host with 58.4% duplicated. An additional 66,548 transcripts totaling 27.3 Mb (18%) aligned to neither pathogen nor host. Of these, 40% were assigned to true fungi, 18% to metazoa, 13% to bacteria, 8% to viridiplantae, 0.3% to Oomycota, 0.2% to archaea, and 0.1% to viruses; Kraken2 was unable to assign a taxonomic identity to 15% of the unaligned transcripts (Supplementary File S1).

**Fig. 5.**
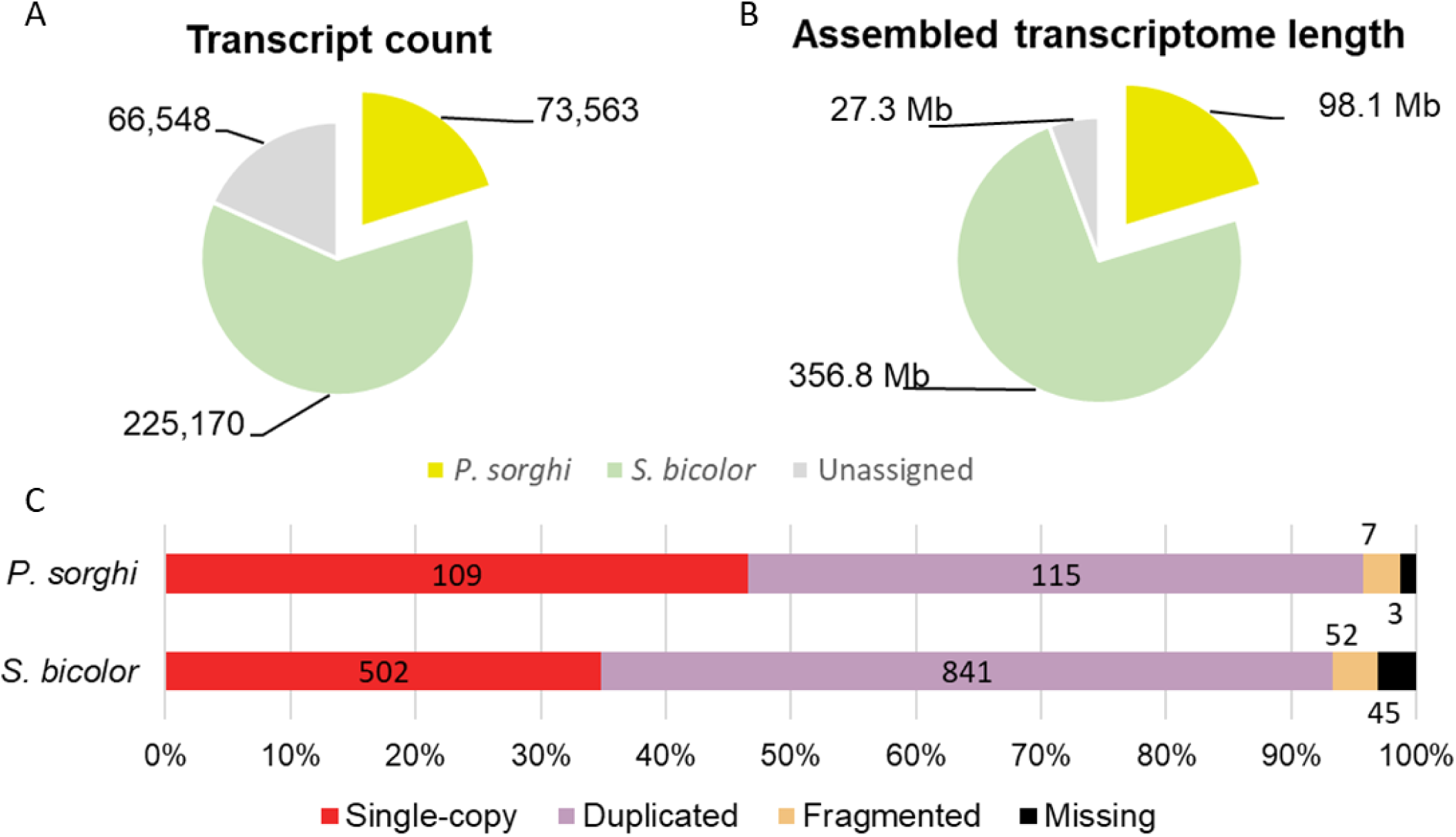
Transcriptome assembly of the pathogen and host. A) Pie charts showing the proportion and length of sequence assigned to *P. sorghi*, *S. bicolor*, and unassigned to either pathogen or host. B) Stacked bar-chart showing the BUSCO results on the transcriptome assemblies of the pathogen and host.

A total of 19,648 genes were annotated in the assembly of *P. sorghi,* which covered 17.4% of the genome assembly. The mean length of gene models was 3.0 kb. The density of genes was higher on chromosome-scale scaffolds than unplaced scaffolds, with 18,623 assigned to chromosomes (Table 1). Signal peptides, indicative of extra-cellular secretion, were identified in 1,518 gene models. These included 360 genes encoding secreted RXLR-like effectors (defined in methods) and eight secreted Crinkler (CRN) effectors. Evidence for transcription was found for 282 of these effector-encoding genes (Supplementary Table S2) and 233 were sufficiently expressed to be included in the subsequent analysis of differential gene expression (DGE; see below). In addition, 68 gene models encode peptides containing an LWY domain, but lacking a secretion signal and 24 gene models encoded peptides containing a CRN motif but no secretion signal (Table 2). Evidence for transcription was found for 79 of these genes and 63 were included in the subsequent DGE analysis. Clustering peptide sequences of putative effectors identified nineteen clusters of genes each encoding three or more high-identity RXLR effectors on ten of the thirteen chromosomes; these clusters spanned 54 kb (Chr. 4) to 28 Mb (Chr. 8). No clusters of genes encoding CRN effectors were localized to a single chromosome. The physical location of clusters of effector genes were not conserved between *P. sorghi* and *P. effusa* (Supplementary Fig. S3). There were 196 genes encoding RXLR effectors, which clustered with no other RXLR and so were classified as singletons. There were 24 singleton genes encoding CRN effectors. The majority of high identity pairs of RXLR effectors were not found on the same chromosome (Fig. 6).

**Fig. 6.**
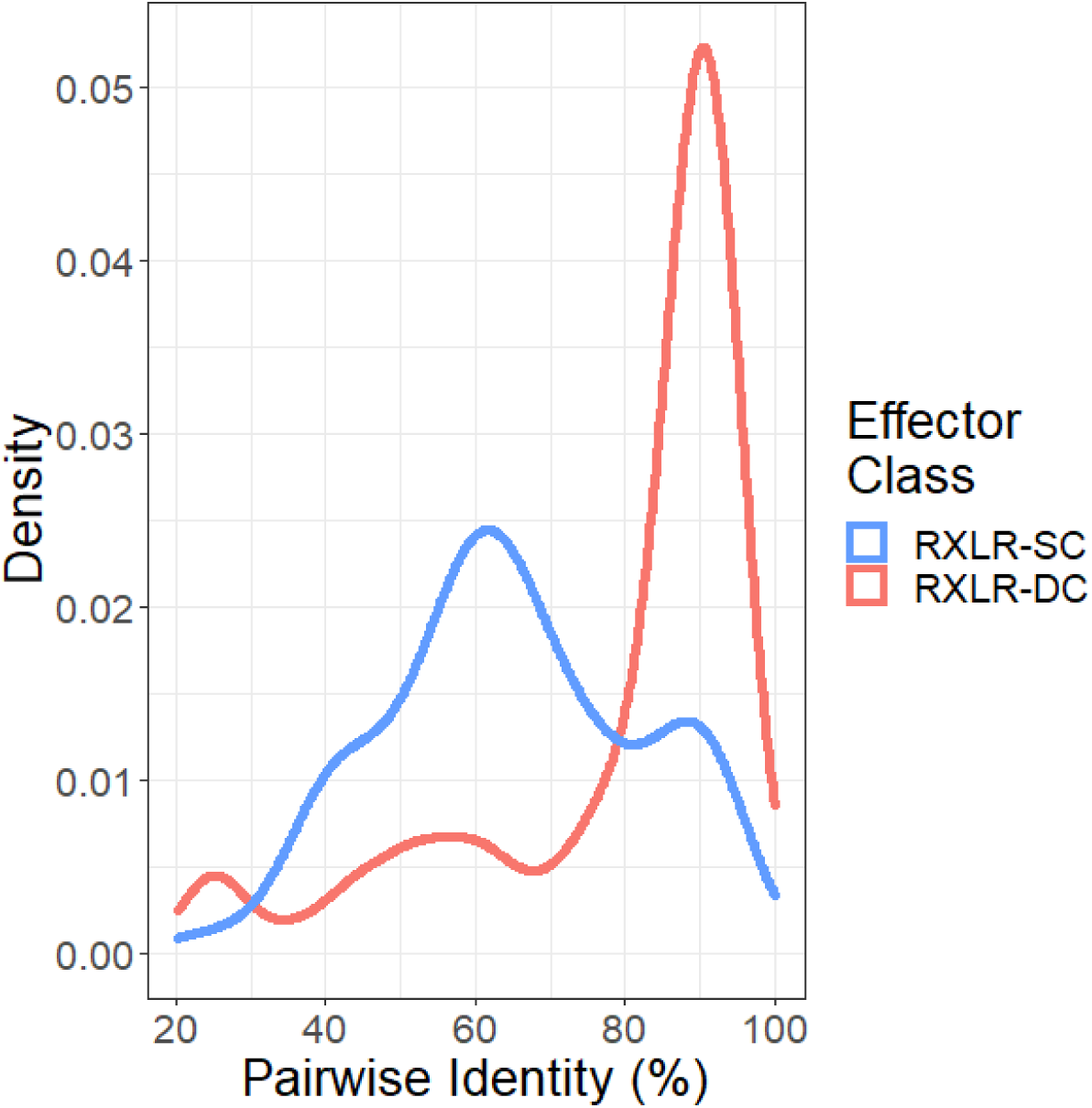
Effector distribution in the genome assembly of *Peronosclerospora sorghi*. Density of 493 pairwise amino acid identities of 232 of 428 putative RXLR effector proteins assigned to 71 clusters by CD-hit. Pairwise identities of genes were more similar when the genes were located on different chromosomes (RXLR-DC) than when the genes were located on the same chromosome (RXLR-SC). Only eight of the 32 genes encoding CRNs were identified in the three clusters, so pairwise similarities are not plotted.

**Table 2.** Annotated effectors in the genome of *Peronosclerospora sorghi*.

| Signal Peptide | RXLR | EER | (L)WY | CRN | Count | Annotations by MAKER | Annotations from ORFs | Number assigned to orthogroups |
| --- | --- | --- | --- | --- | --- | --- | --- | --- |
| + | + | - | - | - | 141 | 141 | 0 | 93 |
| + | + | + | - | - | 130 | 53 | 77 | 43 |
| + | + | + | + | - | 11 | 4 | 7 | 6 |
| + | + | - | + | - | 24 | 13 | 11 | 18 |
| + | - | + | + | - | 38 | 23 | 15 | 33 |
| + | - | - | + | - | 16 | 8 | 8 | 10 |
| - | - | - | + | - | 68 | 31 | 37 | 58 |
| + | - | - | - | + | 8 | 3 | 5 | 7 |
| - | - | - | - | + | 24 | 17 | 7 | 23 |
| + | - | - | - | - | 1150 | 1150 | 0 | 865 |

Of the 19,648 gene models, 15,236 (77.5%) were assigned to 6,231 multi-species orthogroups (Fig. 7A-C). This orthology analysis utilized 39 genome assemblies of 34 species spanning 14 genera across five families of the Oomycota: Peronosporaceae, Pythiaceae, Albuginaceae, Leptolegniaceae, and Saprolegniaceae (Supplementary Table S2). This analysis defined 4,399 core orthogroups common to at least one assembly from each classification of oomycete (Fig. 7E). Core orthogroups accounted for 53.8% of all *P. sorghi* annotations (Fig 7D). In total, 14,264 *P. sorghi* gene models were assigned to 5,799 orthogroups, which also contained gene models from at least one clade 2 downy mildew species. In contrast, 5,325 of these 5,799 orthogroups contained 7,247 *P. effusa* annotations; the 474 orthogroups, which did not have any *P. effusa* gene models assigned, contained 1,375 gene models from *P. sorghi*. For the majority of the 5,325 orthogroups shared between *P. sorghi* and *P. effusa*, the ratio of genes for each species was 1:1 (3,660 orthogroups; 3,901 genes), for 405 orthogroups (625 genes) it was under 1:1, and for 1,260 orthogroups (8,363 genes) it was greater than 1:1 (Fig. 7F). This indicates that the majority of the orthogroups were not duplicated between the two species and is therefore not consistent with a whole-genome duplication event. An additional 973 *P. sorghi* proteins were assigned to 259 interspecies orthogroups that excluded clade 2 downy mildew species. The 4,412 genes annotated in *P. sorghi* but not assigned to multi-species orthogroups were enriched for putative effectors (169 of 460, *X^2^* = 54.365, *p* = 1.665×10^-13^) and proteins predicted to be secreted (443/1518, *X^2^* = 42.363 p = 7.582×10^-11^, Table 2). Relative to *Phytophthora* spp., *P. sorghi* was missing similar orthogroups to other downy mildew clade 2 species (Fig. 7), indicating that the lineage has, most likely, undergone a similar gene-loss event during the transition to obligate biotrophy. In addition, 703 *P. sorghi* gene models were assigned to 332 orthogroups with *Phytophthora* spp., which were not detected in other downy mildew clade 2 species. These results suggest that the increased gene count in *P. sorghi*, relative to other downy mildew species, is not due to gene retention since divergence from a shared ancestor with *Phytophthora* spp.

**Fig. 7.**
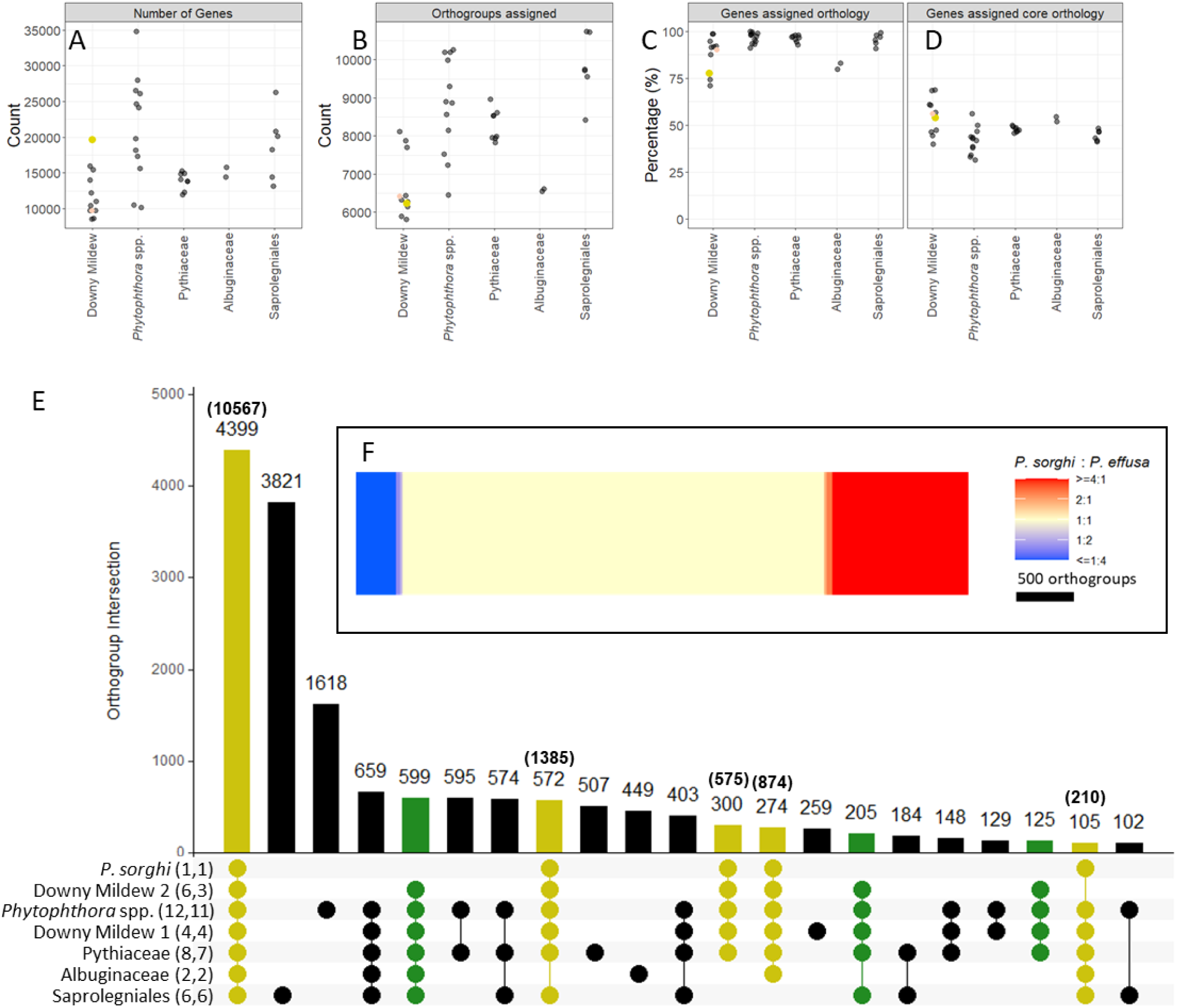
Orthology analysis of *Peronosclerospora sorghi* with 38 other genome assemblies, representing 31 other species of oomycetes. A) Illustration of the number of genes annotated in the genomes used for orthology analysis. B) Scatter plot showing the number of orthogroups protein sequences from each species were assigned to. C) Scatter plot showing the percentage of genes for each species assigned to multi-species orthogroups. D) Same as C, but only considering the 4,399 core orthogroups. In B to D, both downy mildew clades 1 and 2 are condensed to a single category on the x-axis. The large yellow dot indicates *P. sorghi*, the red dot indicates *P. effusa* UA202013. E) An UpSet plot showing multi-species intersections of orthogroups between oomycete phylogenetic classifications. Numbers in parentheses next to the classification indicate the number of assemblies and number of species surveyed under that classification. Only intersections totaling more than 100 orthogroups were plotted; therefore 16,027 of the 17,588 orthogroups calculated are illustrated. The number of orthogroups in each intersection is annotated above the bar. Numbers in parentheses above the bar indicate the number of genes annotated in *P. sorghi* assigned to the orthogroups in the intersection. In total, 13,612 of the 15,237 genes annotated in *P. sorghi* and assigned to multi-species orthogroups are illustrated. Blue bars highlight intersections containing *P. sorghi*. Red bars indicate intersections containing downy mildew clade 2 species, but not *P. sorghi*. Therefore, red bars may indicate unique gene losses in the lineage leading to *P. sorghi* but retained in other related downy mildew clade 2 species. F) Heatmap demonstrating the ratio of sequences annotated in the genomes of *P. sorghi* and *P. effusa* assigned to 5,326 orthogroups that were assigned proteins from both species. The majority of orthogroups are balanced, with the same number of proteins assigned from each species.

The 13 chromosome-scale scaffolds of *P. sorghi* were compared to the seventeen-chromosome assembly of *P. effusa* based on positions of 3,476 single-copy orthologs. This revealed a high degree of synteny between the two genomes. The gene order was significantly correlated between the two assemblies (*r* = 0.937, *p* < 2.2×10^-16^); this clearly validated the assembly of *P. sorghi*. The lineage leading to *P. sorghi* had undergone four chromosome fusions since diverging from the last common ancestor with *P. effusa*. Fusions have occurred between ancestral Chr. 3 and Chr.5, Chr. 4 and Chr.9, Chr. 7 and Chr. 13, and Chr. 10 and Chr.12 of *P. effusa* to form contemporary Chr. 3, Chr. 4, Chr. 6, and Chr. 8, respectively, in *P. sorghi*. The other chromosomes were numbered to retain chromosome order with *P. effusa* (i.e., *P. sorghi* Chr. 1 **≡** *P. effusa* Chr. 1, … *P. sorghi* Chr. 13 **≡** *P. effusa* Chr. 17; Fig. 3H).

Hi-C data allowed the identification of putative coordinates for centromeres. Aligning Hi-C reads back to the genome assembly and calculating the mean cis-interaction distance in 100 kb windows identified chromosomal regions enriched for short distance interactions (Fig. 8). This demonstrated that the chromosomes were likely organized in Rabl-like configurations ([22]; Fig. 1A) that resulted in enriched short-range interactions. Because centromeres have been annotated in *P. effusa*, synteny of flanking single copy orthologs could be used to validate the coordinates in *P. sorghi*. This analysis showed that orthologs assigned to either chromosome arm in *P. effusa* were located on either side of the putative centromere in *P. sorghi*, indicating that centromere positions were similar between these species. Similar centromere positions could be identified for one of the primogenitors in three of the four chromosome fusions identified in *P. sorghi* relative to *P. effusa*. Chromosome 4 of *P. sorghi* retained the centromere syntenic to *P. effusa* Chr. 9; Chr. 6 of *P. sorghi* retained the centromere syntenic to *P. effusa* Chr. 13, and Chr. 8 retained the centromere syntenic to Chr. 10. For *P. sorghi* Chr. 3, it was not apparent if a centromere was retained from either primogenitor. While the mean cis interaction distance for bins in the regions syntenic to the centromere of *P. effusa* Chr. 5 was low, the lowest mean cis interaction distance was at the point of fusion. This could indicate a neocentromere or a false signal because the chromosome arms of the primogenitor are short; the length of regions in *P. sorghi* that contained single copy orthologs conferring to the chromosome arms was ∼2.0 Mb and ∼1.3 Mb. Additional evidence for centromere location was sought through identification of Copia Like Transposons (CoLT). These elements were identified on every chromosome, but they were not enriched in the vicinity of putative centromeres. However, the centromeres of *P. sorghi* were not fully assembled and CoLTs were identified in Chr. 0. Therefore, an improved assembly is required to fully characterize the centromeres of *P. sorghi* and to determine if CoLTs are present in each centromere.

**Fig. 8.**
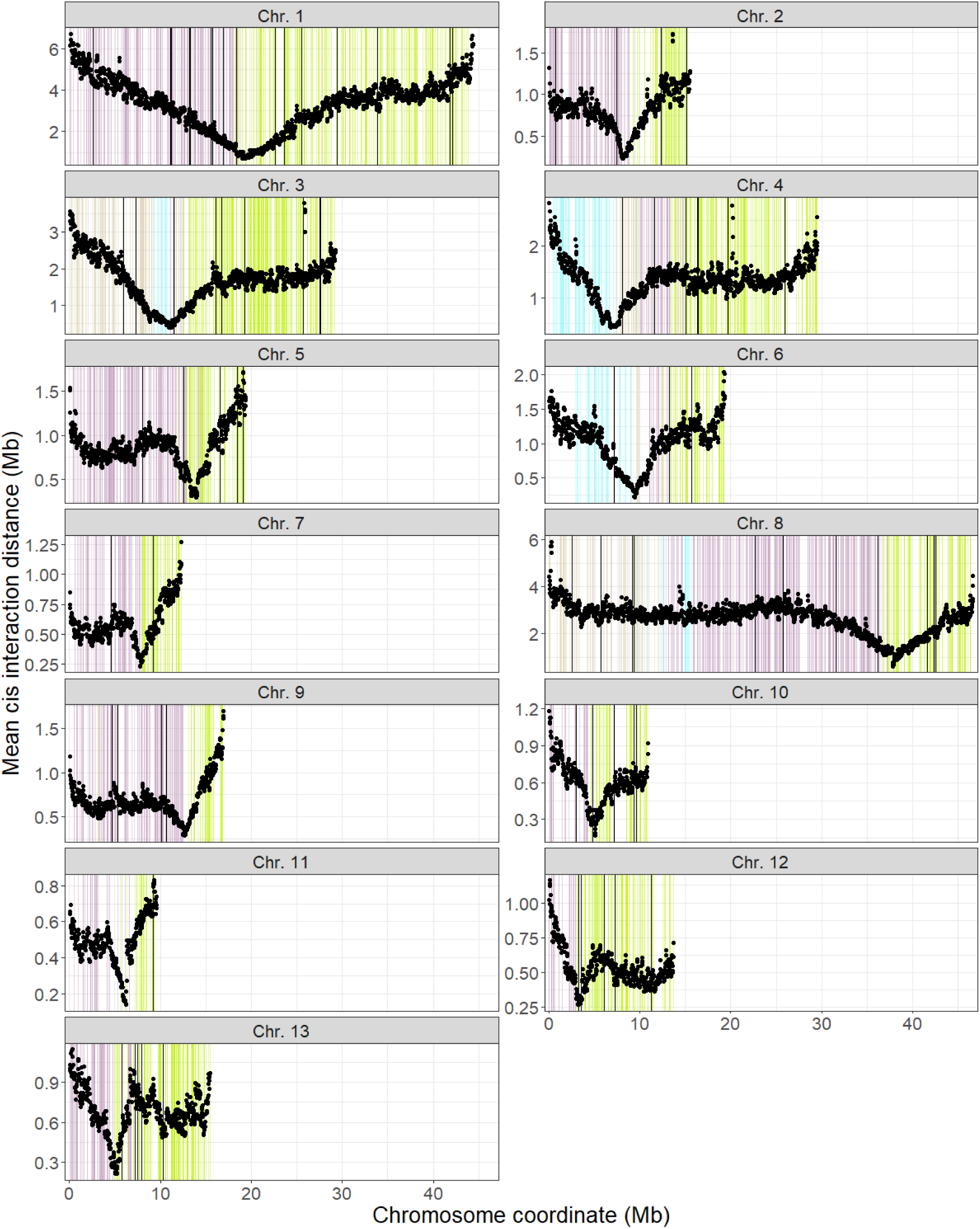
Conservation of centromeric regions between the genomes of *Peronosclerospora sorghi* and *Peronospora effusa*. The mean cis interaction distance between Hi-C reads was calculated in 100 kb windows, with a 25 kb step, across the genome of *P. sorghi*. There was a single region enriched for short distance *cis* interactions per chromosome, corresponding to the centromeric region. Coordinates of single copy orthologs in the assembly of *P. sorghi*, used to establish synteny with *P. effusa* (Fig. 5), were plotted in the background. The colors indicate which chromosome arm of *P. effusa* the ortholog is located on; proximal chromosome arms are purple, and distal chromosome arms are green for each chromosome. Most chromosomes have only two colors because they share the same ancestral conformation. Chromosomes 3, 4, 6, and 8 have four colors because these chromosomes have undergone fusions relative to *P. effusa*. Brown depicts orthologs derived from a proximal region, and blue depicts orthologs derived from a distal region. Except for Chr. 3, the region enriched for short distance *cis* interactions co-locates with the change in color, indicating that the centromere positions are similar between the two species. In Chr. 3 the putative centromere appears to be at located at the fusion point between primogenitors. Black bars indicate the location of Copia-like transposons in the genome of *P. sorghi*.

Synteny between *P. sorghi* and *P. effusa* was established using 3,476 genes that were single-copy in both genomes (Fig. 3H, Fig. 9A). We then investigated whether the expansion in gene number in *P. sorghi* was the result of local duplication or genome-wide dispersal. A gene was considered expanded in *P. sorghi* if multiple genes annotated in *P. sorghi* were assigned to an orthogroup containing only one *P. effusa* gene (>1:1); in total, there were 5,675 such multicopy genes in *P. sorghi*. Comparison of the chromosomal positions of these expanded genes in *P. sorghi* established the presence of two broad categories of orthologous genes; 1,506 were located on syntenic chromosomes and 3,510 genes were scattered through the genome on non-syntenic chromosomes (Fig. 9B); in addition, 659 were on Chr. 0. Only a few genes (798) were expanded in *P. effusa* relative to 320 *P. sorghi* genes (Table 3). In total, 666 of these were located on the syntenic chromosome (Fig. 9C). The remaining 10,177 genes in *P. sorghi* were classified into other categories: 3,417 genes were orthologous to multiple *P. sorghi* and multiple *P. effusa* annotations (>1:>1; Table 3), which were located on both syntenic and non-chromosomes (Supplementary Fig. S4); 608 genes annotated in *P. sorghi,* which were assigned as the lone representative of *P. sorghi* to inter-species orthogroups lacking *P. effusa* (1:0); 1,740 genes assigned to inter-species orthogroups containing multiple *P. sorghi* annotations but no *P. effusa* annotations (>1:0); and 4,412 *P. sorghi* annotations lacking orthology with any other oomycete. The pattern of genome-wide dispersal of expanded genes is consistent with duplication by retrotransposition rather than local duplication.

**Fig. 9.**
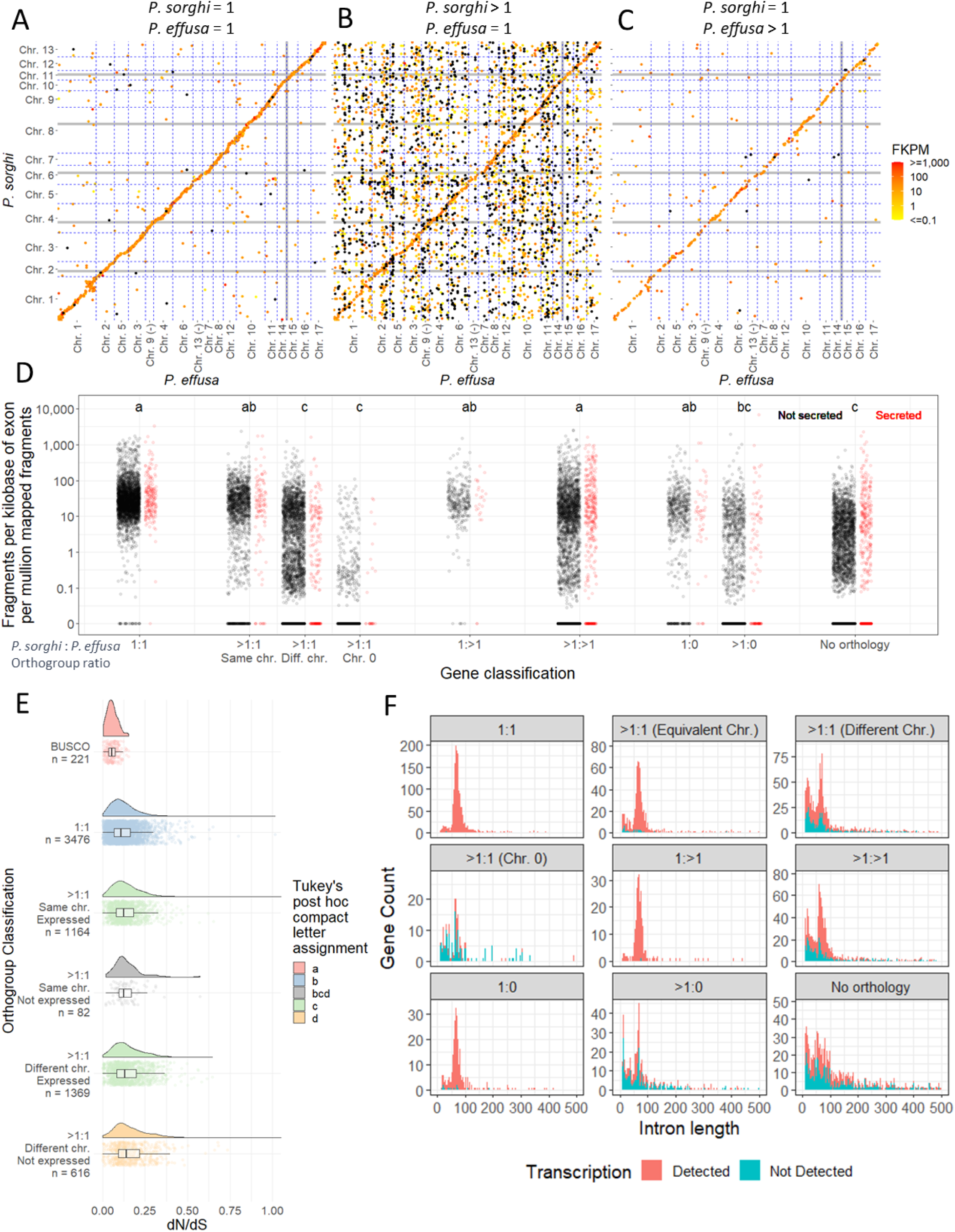
Multiplication and dispersal of genes in the genome of *Peronosclerospora sorghi*. A) Scatter plot showing the coordinates of single-copy genes in the genomes of *P. effusa* and *P. sorghi*. The x-axis is ordered to demonstrate collinearity, like Fig. 3H. Axes are not scaled to size. Grey gridlines indicate 50 Mb boundaries. Blue dotted lines indicate chromosome boundaries. Points are colored by their transcript abundance (FPKM). Black dots indicate 0 FPKM. B) Like A, except genes were expanded in *P. sorghi*, single copy in *P. effusa*. C) Like A, genes were single copy in *P. sorghi*, expanded in *P. effusa*. D) Scatter plots showing the transcript abundance of genes annotated in *P. sorghi*. Genes were assigned to different categories (x-axis) based on their orthology to *P. effusa*. Genes expanded in *P. sorghi* (>1:1) were sub-categorized based on their synteny with *P. effusa*. Genes encoding peptides predicted to be secreted were plotted in red for each category. Lowercase letters indicate different groups of significance as calculated by Tukey’s HSD test. E) Raincloud plots depicting signatures of selection on orthologs single copy in both *P. sorghi* and *P. effusa* and expanded in *P. sorghi* but single copy in *P. effusa*. Genes expanded in *P. sorghi* were split on the basis of synteny with *P. effusa* and detection of transcripts (FPKM > 0). BUSCO genes are shown as representative of highly conserved genes. Colors indicate different groups of significance as calculated by Tukey’s test. F) Distribution of intron lengths for genes annotated in *P. sorghi*. Genes were split based on their synteny with *P. effusa*. The color of the stacked histogram indicates the number of introns originating from genes lacking or with detected transcripts.

**Table 3.** Overview of orthologous gene assignment for *Peronosclerospora sorghi* annotations relative to *Peronospora effusa*.

| *P. sorghi* : *P. effusa*<br>orthogroup<br>ratio | Assigned genes | Expressed genes | Differential<br>gene<br>expression<br>(up : down). | Mean FPKM | Mean encoded<br>peptide length* | Mean exon<br>count* | Genes with<br>secreted<br>product | Expressed<br>genes with<br>secreted<br>product | Mean<br>FPKM |
| --- | --- | --- | --- | --- | --- | --- | --- | --- | --- |
| 1:1 | 3,476 | 3,447 (99.1%) | 91 : 25 | 59.35 | 491.7 (a) | 1.66 (a) | 202 | 201 (99.5%) | 101.0 |
| >1:1 – syntenic<br>chr. | 1,506 | 1,346 (89.3%) | 30 : 10 | 46.42 | 472.9 (a) | 1.67 (a) | 113 | 104 (92.0%) | 89.55 |
| >1:1 – non-syntenic chr. | 3,510 | 2,181 (62.1%) | 17 : 15 | 19.16 | 201.6 (b) | 1.29 (b) | 191 | 129 (67.5%) | 34.79 |
| >1:1 Chr. 0 | 659 | 499 (75.7%) | 0 : 0 | 6.22 | 268.1 (c) | 1.30 (bc) | 25 | 10 (40%) | 8.31 |
| 1:>1 | 320 | 313 (97.8%) | 8 : 5 | 53.89 | 515.4 (a) | 2.78 (e) | 24 | 22 (91.7%) | 54.98 |
| >1:>1 | 3,417 | 2,257 (66.1%) | 126 : 29 | 50.78 | 339.5 (d) | 1.21 (bc) | 364 | 297 (81.6%) | 90.05 |
| 1:0 | 608 | 524 (86.2%) | 15 : 6 | 40.91 | 293.8 (c) | 1.68 (a) | 52 | 47 (90.4%) | 83.33 |
| >1:0 | 1,740 | 716 (41.1%) | 9 : 13 | 32.55 | 198.8 (b) | 1.14 (cd) | 104 | 59 (56.7%) | 164.66 |
| No orthology | 4,412 | 2,274 (51.5%) | 32 : 27 | 16.08 | 133.0 (e) | 1.01 (d) | 443 | 269 (60.1%) | 59.79 |
| Total | 19,648 | 13,557 (69.0%) | 328 : 130 |  |  |  | 1,518 | 1,138 (75.0%) |  |

\*Parenthesized letters indicate significant differences in pairwise analysis (Tukey’s HSD).

Transcript abundance of syntenic and non-syntenic genes was compared to determine the frequency of potential pseudogenes in each set. In total, transcripts were detected for 13,281 of the 19,648 annotated genes. Transcripts were detected for 3,447 of the 3,476 single copy orthologs identified between *P. sorghi* and *P. effusa* (1:1). Their mean level of abundance was 58.86 fragments per kilobase of exon per million mapped fragments (FPKM; Fig. 9D). No transcripts were identified for the other 29 of these genes, 20 of which were located on non-syntenic chromosomes. Transcripts were detected for 1,346 of 1,506 expanded genes located on syntenic chromosomes relative to *P. effusa*, 2,181 of 3,510 expanded genes on non-syntenic chromosomes, and 499 of 659 expanded genes on Chr. 0 (Table 3). The mean FPKM of expressed, expanded genes in syntenic positions was lower, but not significantly lower than for genes classified as 1:1 (Fig. 9D; Table 3). In addition, many expanded genes with high transcript abundance were in syntenic positions (Fig. 9B), indicating that these genes were the ancestral copies. The mean FPKM for expanded genes on non-syntenic chromosomes and Chr. 0 was significantly lower than the mean FPKM for both genes classified as 1:1 and expanded genes in syntenic positions (Table 3; Fig. 9D). The distribution of FPKM for genes on non-syntenic chromosomes was bimodal (Fig. 7D; Supplementary Fig. S5), suggesting that some duplicated genes on non-syntenic chromosomes had higher transcript abundance (1,378 genes with mean 30.08 FPKM), while others had lower transcript abundance (803 genes with mean 0.43 FPKM). Treated independently, both these subsets had significantly lower transcript levels than expanded genes located in syntenic positions (p = 0.00128 & p < 2.2×10^-16^). Transcripts were detected for 313 of 320 *P. sorghi* single copy genes that appeared expanded in the genome of *P. effusa* (Table 3). The transcript abundance of these genes was not significantly different from single copy genes in *P. sorghi* or genes expanded in *P. sorghi* located on syntenic chromosomes (Fig. 9D). Levels of transcripts for genes assigned interspecies orthology but lacking orthologs in *P. effusa* (1:0 & >1:0) did not differ significantly from genes that were syntenic with *P. effusa* (Table 3; Fig. 9D). Genes lacking interspecies orthology had significantly lower transcript abundance than most other genes (Fig. 9D). Overall, these results show that genes duplicated to non-syntenic positions in the genome of *P. sorghi* have fewer detectable transcripts than their ancestral, syntenic counterparts.

The transcript abundance of the 1,518 annotated genes encoding predicted secreted proteins was also investigated. Transcripts were detected for 75% of these genes, which was significantly higher than the portion of transcripts detected for genes encoding non-secreted peptides (*X^2^*=27.088, p=1.94×10^-7^). The mean transcript abundance of genes predicted to encode a secreted product was also higher than genes not predicted to encode secreted proteins (Table 3). Some of the genes with the most abundant transcripts encoded secreted peptides (Fig. 9D) including annotated effectors. Therefore, transcripts of genes encoding signal peptides, including those on non-syntenic chromosomes or when lacking inter-species orthologs, were detected at higher abundances than the genome-wide average.

Coding sequences were compared to determine if gene expansion in *P. sorghi* had resulted in changes in selection pressures acting on single copy and duplicated genes. The ratio of non-synonymous to synonymous polymorphisms (dN/dS) in *P. sorghi* genes since divergence from the common ancestor with *P. effusa* was calculated and summarized considering chromosomal location of duplicated genes and whether transcripts were detected for duplicated genes. The mean dN/dS of the 3,476 single copy orthologs (1:1) was 0.125, consistent with these genes undergoing purifying selection since divergence. The mean dN/dS for genes that were probably ancestral (>1:1 on syntenic chromosomes), and for which transcripts were detected, was 0.140. While this is consistent with purifying selection, it was significantly higher than the dN/dS for single copy orthologs (Fig. 9E), suggesting relaxed purifying selection. The mean dN/dS for expanded genes (>1:1) on non-syntenic chromosomes for which transcripts were detected was 0.150. This was also significantly different from single copy orthologs (1:1), but not expanded genes (>1:1) on syntenic chromosomes with transcripts (Fig. 9E). The mean dN/dS for expanded genes (>1:1) on non-syntenic chromosomes lacking transcripts was 0.168 and was significantly higher than the means for the other described subset of genes (Fig. 9E). The mean dN/dS for expanded genes on syntenic chromosomes and lacking transcripts was 0.142 and did not differ significantly from the previously described subsets (Fig. 9E). Selection may have been lost on some duplicated genes, consistent with pseudogenization. Therefore, comparative sequence analysis suggests that duplicated genes have significantly different selection pressures than conserved, single copy genes, and that duplicated genes lacking transcription have been under even weaker purifying selection.

Additional evidence for pseudogenization of expanded genes was sought via global characterization of genes. The mean encoded peptide length (MEPL) for the 3,476 single copy orthologs was 491.7 residues. This was not significantly different from the MEPL for the 1,506 expanded genes (>1:1) on syntenic chromosomes or the 320 single copy genes expanded in *P. effusa* (1:>1; Table 3). All other subsets of proteins had significantly shorter MEPLs, including >1:1 genes on non-syntenic chromosomes. Shorter MPELs may indicate nonsense mutations introducing premature stop codons in the ORF. The mean exon count (MEC) of single copy orthologs was 1.66 and did not significantly differ from the MEC of >1:1 genes on syntenic chromosomes or single copy *P. sorghi* genes not found in *P. effusa* (1:0; Table 3). The MEC of 1:>1 genes was significantly higher than 1:1 genes. The MEC of all other gene classifications was significantly lower, including >1:1 genes on non-syntenic chromosomes. Reduced exon counts could be due to nonsense mutations or intron loss due to duplication by retrotransposition. The distribution of intron lengths was summarized for each gene category in the context of transcript detection. For 1:1 genes, a major peak was detected with a mode intron length (MIL) of 67 base pair (bp; a minor peak was present with a MIL of 25 bp that may be an annotation artifact). Similar profiles were obtained for >1:1 genes on syntenic chromosomes, 1:>1 genes, and 1:0 genes (Fig. 9F). Transcripts were detected for most of the genes in these categories (Table 3; Fig. 9F). For the other gene categories, two similar peaks could be detected; however, the major and minor peaks had similar counts and genes with zero transcripts detected were found under both peaks (Fig. 9F). For all annotations, the peak of larger MILs may represent introns of optimal size for splicing in *P. sorghi*; the peak of smaller MILs might represent introns incorrectly predicted by the annotation software. It therefore seems likely that the 19,458 genes annotated for *P. sorghi* contains true protein-coding genes often retaining synteny with *P. effusa* as well as a large number of pseudogenes that lacked evidence for transcription and were predicted to encode shorter peptides in fewer exons.

The *P. sorghi* assembly was investigated to determine whether the genome was compartmentalized in relation to transcription and gene classification. Both the 5’ and 3’ intergenic distances between genes were bimodal indicative of gene-dense (intergenic distances of less than 6.5 kb either side) and gene-sparse (intergenic distances greater than 6.5 kb on either or both sides) regions in the genome (Fig. 10A). Genes for which transcription was and was not detected were located in both such gene-dense and the gene-sparse regions. When orthology was considered, all previously ascribed categories were distributed across the different genomic compartments (Fig. 10B). Therefore, conserved single-copy orthologs were annotated in both the gene-sparse and gene-dense region. In addition, annotations that may represent pseudogenes were also located in the gene-sparse and gene-dense regions, regardless of transcriptional status. The dN/dS of genes in the gene-dense region did not significantly differ from the dN/dS of genes in the gene-sparse regions (p=0.128). Therefore, pseudogenes and high-confidence protein coding genes were present in both the gene-dense and gene-sparse compartments of the *P. sorghi* genome.

**Fig. 10.**
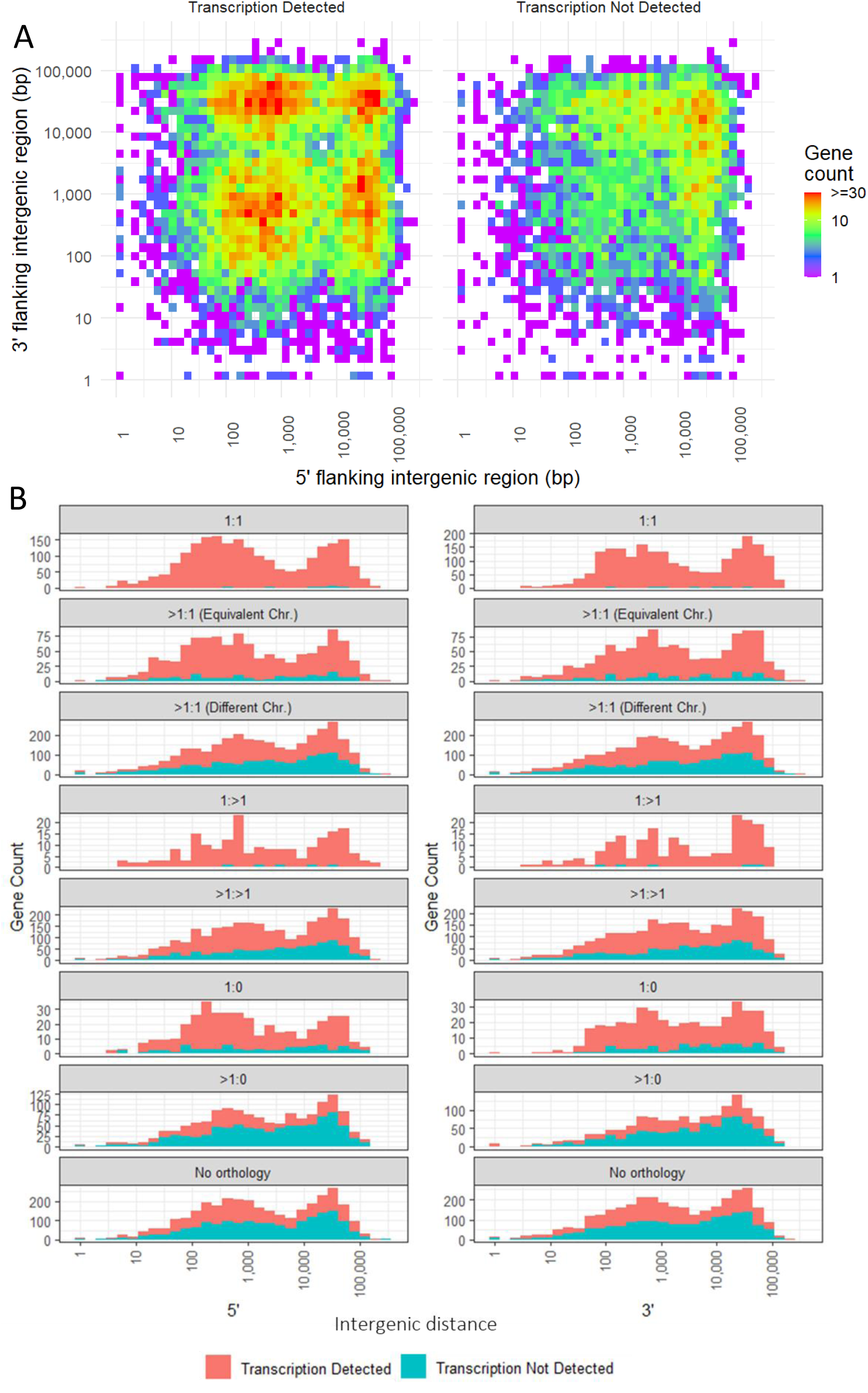
Compartmentalization of the *Peronosclerospora sorghi* genome based on intergenic distances. A) For each annotated gene, the 5’ distance (bp) to the next gene was plotted on the y-axis, and the 3’ distance (bp) was plotted on the y-axis. A heatmap was plotted for genes both with and without detectable transcripts. The heatmaps demonstrate that intergenic distances on both flanks were bimodal. Some genes had short intergenic distances (<6.5 kb) on both flanks, consistent with high gene-density. Others had long intergenic distances (>6.5 kb) on both or either flank, consistent with gene-sparsity. B) Stacked histograms on the left show the 5’ intergenic distance, while the right show the 3’ intergenic distance. Separate histograms were plotted for genes based on their synteny with *P. effusa*. All gene categories had genes consistent with high gene density and gene sparsity. The stacked histogram indicated the number of genes for which transcripts could and could not be detected. Genes with and without detectable transcripts had intergenic distances consistent with high gene density and gene sparsity.

Differential gene expression was analyzed to investigate transcriptional differences pre- and post-sporulation of *P. sorghi*. Multi-dimensional scaling indicated that normalized read counts of pathogen and host genes, for each of the two biological conditions could be distinguished from one another across the three replicates (Fig. 11A, B). A total of 10,451 genes passed the 5 counts per-million threshold. There was evidence for 328 up-regulated and 130 down-regulated genes post-sporulation, compared to pre-sporulation after correction for false discovery (Fig. 11 C). These two subsets of genes were enriched for peptides predicted to be secreted, with 75 up-regulated genes predicted to be secreted (*p* = 1.03 x10^-24^) and 27 down-regulated genes (*p* = 2.60×10^-8^). Pfam domains encoded by differentially expressed genes suggests that up-regulated proteins include transporters, cutinase, chitinase, cellulose synthase, chitin synthase, and a necrosis inducing protein (Supplementary Table S4). Fewer down-regulated genes could be due to mycelium within the plant tissue after sporulation. In contrast, more genes from the host were downregulated (1,498) than upregulated (522) post-sporulation (Fig. 11D). These included six downregulated genes encoding disease resistance proteins, one upregulated, 21 down-regulated genes encoding transporters, 13 up-regulated, and 10 down-regulated genes encoding dehydration response element binding proteins. These differentially regulated host genes could include genes in response to the pathogen or in response to the environment. There are extensive data available for further study of the host complement that is beyond the focus of this paper. The differentially expressed pathogen genes were enriched for genes syntenic with *P. effusa*. Synteny could be established for 356 of the 458 differentially expressed genes, of which: 116 were single copy in both *P. sorghi* and *P. effusa*; 72 were expanded in *P. sorghi* relative to *P. effusa*; 13 were single copy in *P. sorghi*, but expanded in *P. effusa*; 155 were multi-copy in both (Table 3). The expanded, differentially regulated genes in *P. sorghi* were enriched for genes which retained synteny (*X*^2^ = 8.68, p = 0.0032). Therefore, genes which retain synteny between these two distinct species are also differentially regulated during the lifecycle of *P. sorghi*.

**Fig. 11.**
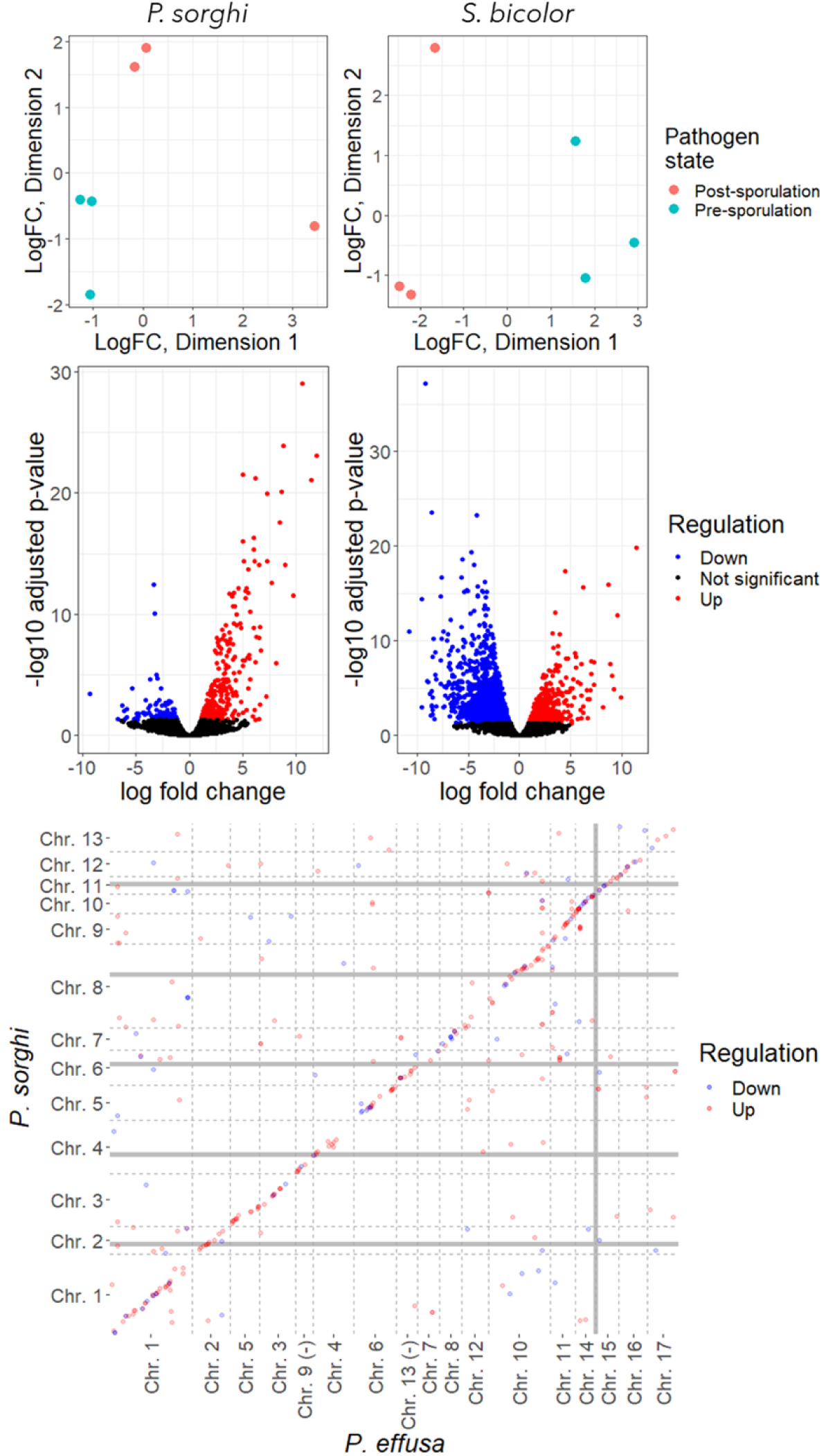
Differential gene expression of *Peronoscleropsora sorghi* and *Sorghum bicolor*. A) Multi-dimensional scaling (MDS) plots demonstrating that the gene counts of the pathogen can be distinguished based on their pre- or post-sporulation source. B) MDS plot demonstrating that the gene counts of the host could be distinguished based on their pre- or post-sporulation source. C) Volcano plot demonstrating up- and down-regulated genes of the pathogen. D) Same as C, but for the host. E) Scatter plot of differentially expressed genes showing enrichment of genes syntenic with *P. effusa*.

## Discussion

This paper describes a third chromosome-scale genome assembly for an oomycete following the genetically oriented genome assembly of *Bremia lactucae* [15] and the telomere-to-telomere (T2T) genome assembly of *Peronospora effusa* [16]. All three of these species represent different genera of oomycetes, which cause downy mildew diseases on different plants. *P. sorghi* is phylogenetically closer to *P. effusa* (downy mildew clade 2) than *B. lactucae* (downy mildew clade 1) (Fig. 2). The genome architecture of *P. sorghi* was unique compared to the highly syntenic genomes of *B. lactucae* and *P. effusa* because it was more homozygous, larger, and had fewer chromosomes. Increased homozygosity was computationally beneficial when generating the genome assembly and may explain why, in combination with Hi-C, it was possible to obtain a chromosome-scale genome assembly of *P. sorghi*. Previously, the ancestral chromosome-state of downy mildews and many *Phytophthora* spp. has been described as that of *B. lactucae* and *P. effusa*, which share high collinearity between their respective chromosomes [16]. While the genome of *P. sorghi* has fewer chromosomes, it retains a high level of synteny with the ancestral state; nine *P. sorghi* chromosomes were colinear with nine chromosomes of *P. effusa* and *B. lactucae* (Fig. 3) [16]. The four other *P. sorghi* chromosomes are unique fusions compared to *P. effusa* and *B. lactucae*, but retain a high level of synteny with their primogenitors (Fig. 3). Therefore, this genome assembly presents a contemporary state derived from the conserved ancestral configuration.

Identification of centromeres also supports the 13-chromosome architecture of *P. sorghi*. Analysis of Hi-C data identified one centromeric region on each chromosome (Fig. 1A, 8). Aligning the assembly of *P. sorghi* to the assembly of *P. effusa* showed which centromeres were lost after chromosome fusions and that the retained centromeres were in approximately syntenic positions (Fig. 8). It was not possible to compare centromere sequences between the two species because the current assembly of *P. sorghi* is based on Illumina 10x Genomics sequencing in contrast to the gapless chromosomes of *P. effusa* that is based on Pacific BioSciences HiFi reads [16]. It will be informative in the future to generate long read assemblies of *Peronosclerospora* spp. to assemble the centromeres completely and determine whether they harbor sequences conserved with *P. effusa*.

Genome-wide synteny indicates that proliferation of repeat sequences underlies the increased genome size of *P. sorghi* relative to *P. effusa* rather than a whole genome duplication event. The assembled length of *P. sorghi* was more than six times that of the T2T assembly of *P. effusa*, although their chromosomes were highly syntenic. Therefore, the inflated genome size was not caused by auto- or allopolyploidy. Instead, since diverging from their most recent common ancestor, the genome of *P. sorghi* has accumulated or retained more transposable elements than the genome of *P. effusa*. Comparing 100 kb windows across the two assemblies, *P. sorghi* had a higher density of repeats and a lower density of genes annotated than *P. effusa* (Fig. 3F, G). This is consistent with the previously reported correlation of repeat content with genome size across multiple oomycete species, including downy mildews [18]. Inflation of genome size exclusively by retention of transposable elements is different from some *Phytophthora* spp., where whole-genome duplication in addition to proliferation of transposable elements have resulted in an increased genome sizes [23, 24].

Despite its high gene-model count, *P. sorghi* probably underwent similar gene-loss events to other downy mildews due to its biotrophic and non-flagellate lifestyle. Evidence for this comes from the high number of orthogroups that are present in *Phytophthora* spp. and species in the Pythiaceae, but absent in *P. sorghi* and other downy mildew species; only 105 inter-species orthogroups were assigned a total of 210 sequences from *P. sorghi* but no other downy mildew clade 2 species (Fig. 7). This is consistent with *P. sorghi* undergoing similar gene-loss events as other clade 2 downy mildews. More contiguous assemblies of *Phytophthora* spp. from multiple *Phytophthora* clades as well as more downy mildew species, may reveal genes lost in downy mildew species, which encode proteins dispensable in the biotrophic lifecycle, but are required for the necrotrophic phase of the lifecycles of *Phytophthora* species. Since *Phytophthora* species produce flagellated zoospores, the flagellated state is likely ancestral to all downy mildews. Loss of flagella genes likely occurred multiple times in clade 2 downy mildews because *Sclerospora* and *Pseudoperonospora* spp. retain flagellated zoospores, but *Peronosclerospora*, *Hyaloperonospora*, and *Peronospora* spp. do not (Fig. 2) [4, 19, 25].

The increased number of genes in *P. sorghi* relative to other downy mildews is likely due to gene expansion through retrotransposition, which has led to the accumulation of pseudogenes (Fig. 9) and dispersed, transcribed genes, including genes encoding putative effectors (Fig. 3). For the majority of orthogroups, the ratio of genes assigned from *P. sorghi* compared to *P. effusa* was 1:1 (Fig. 7F), indicating that the majority of orthologs were conserved. Further to this, the majority of these orthogroups were single copy in *P. effusa* and *P. sorghi*, syntenic, transcribed, had signals of purifying selection, and a normal distribution of intron lengths (Fig. 9). A minority of orthogroups (1,260) were expanded in *P. sorghi* relative to *P. effusa*, accounting for the majority of orthologous genes (8,363 genes) annotated in *P. sorghi* (Fig. 7); a minority of these genes were syntenic between *P. sorghi* and *P. effusa* and retained similar characteristics with the single copy-set, suggesting that they were ancestral (Fig. 9, Table 3). The remainder of these genes were on non-syntenic chromosomes, suggesting duplication by retrotransposition. Transcripts could not be detected for a larger portion of these genes (Table 3) and when transcripts were detected, their transcript abundance was significantly lower (Fig. 9). In addition, genes on non-syntenic chromosomes encoded shorter peptides in fewer exons and had a large number of short introns predicted, characteristic of pseudogenes. Of the expanded genes, those that were transcribed had significantly weaker signals of selection than single copy genes; however, they had similar signals of selection regardless of whether they were syntenic, suggesting that both the ancestral and transposed copies were under similar relaxed selection [26]. Some of the transcribed non-syntenic genes had short introns predicted suggesting nonsense alleles or transcribed pseudogenes (Fig. 9F). Expanded genes with undetected transcription had weaker signals of purifying selection consistent with pseudogenization (Fig. 9E). While the genome of *P. sorghi* was consistent with compartmentalization (Fig. 10A), the compartments did not have distinct complements of putative pseudogenes and protein-coding genes; syntenic genes annotated as single copy in both *P. sorghi* and *P. effusa* were located in both gene-dense and gene-sparse compartments, as were the syntenic and non-syntenic subset of the expanded genes (Fig. 10B). Genes lacking transcription, expanded in *P. sorghi*, which had weaker signals of purifying selection (Fig. 9E, i.e. pseudogenes), were also present in both compartments (Fig. 10B). Although both single copy and expanded genes were differentially expressed (Table 3), the majority were syntenic (Fig. 11 E). In summary, the majority of orthologs between *P. sorghi* and *P. effusa* are single copy, but the majority of annotations in the *P. sorghi* genome originate from a minor portion of expanded orthologs. Some of the expanded genes may be functional paralogs, while others are likely pseudo-genes. These findings should be considered when analyzing other oomycete assemblies, especially assemblies with high gene counts because they may also harbor a significant portion of pseudogenes. Identification of the syntenic, ancestral gene pairs will also be important if attempting reference-based scaffolding across species.

The evolutionary advantage of increased genome sizes in currently reported assemblies of graminicolous downy mildews compared to other species remains unknown. One consequence of genome-inflation by retrotransposition in *P. sorghi* is the presence of highly similar genes encoding effectors on distinct chromosomes, which is not present in *P. effusa* [16]. In *P. effusa,* gene duplications, including of effector-encoding genes, appear to have more often resulted in clusters of genes at similar chromosome positions (Fig. 9C). Local gene duplication is likely the result of unequal crossing over or non-homologous recombination, rather than retrotransposition [27]. The coenocytic nature of oomycetes means that within their mycelia there may be rapid somatic evolution with the fittest nuclei proliferating and being preferentially represented in the next asexual or sexual generation. For *P. effusa*, this has resulted in an organism with a compact genome and tight clusters of effectors. For *P. sorghi* this has resulted in an organism with a genome bloated with repeats, putative pseudogenes, and widely dispersed effector genes (Figs. 3, 5). Comparative genomics with other oomycetes, including downy mildews and *Phytophthora* spp., will determine whether there is a relationship between genome size and effector distribution.

The mitochondrial genome of *P. sorghi* is circular in orientation and at approximately 38.5 kb is similar in size to other oomycetes (Fig. 4). The mitogenome encoded the same common suite of genes observed in other oomycetes, including 35 genes, ribosomal RNAs, tRNAs and the putative ORFs *ymf16, ymf98, ymf99, ymf100*, and *ymf101*. There was also an additional putative ORF (*orf161*) of unknown function encoded between *cox1* and *ymf100* that is unique to *P. sorghi.* The presence of species-specific putative ORFs is a common phenomenon and was observed in all nine of the taxa illustrated in Fig. 4.

The mitochondrial gene order between the nine Peronosporaceae taxa representing six genera was largely conserved with the exceptions of inversions of gene-blocks which retained internal gene order (Fig. 4). This finding parallels a previous study examining a broader array of taxa, in particular *Phytophthora* species [28]. The mechanisms driving this type of genome evolution has yet to be determined. Recently inverted repeats were reported as flanking a region of the mitogenome, which was inverted in some, but not all isolates of *P. effusa* [29]. Recombination between the small 1,150 bp inverted repeats in *P. ramorum* generated isomers of the mitochondrial genome where the region between the repeats was also present in an inverted orientation [30], but this did not lead to structural changes in the mitochondrial genome. However, some taxa, including *P. sorghi*, lack inverted repeats flanking inversions/translocations relative to other taxa (Fig. 4). Therefore, it is possible that other mechanisms may underlie these differences.

The genome of *P. sorghi* provides the molecular foundation to characterize taxonomic relationships within and between species of *Peronosclerospora* and to deploy diagnostic molecular markers. Establishing molecular diagnostics to detect and distinguish potential immigration of graminicolous downy mildews, including *P. sorghi* and other *Peronosclerospora* spp., such as *P. philippinensis,* is a priority to prevent epidemics on corn in the USA. This is desirable since US corn is potentially highly susceptible to tropical downy mildew causing pathogens [3, 14]. Highly conserved markers may provide genus-level resolution suitable for screening at ports of entry. The mitogenome is likely to be a good source for molecular marker development given its unique configuration in *P. sorghi* compared to other Peronosporaceae species (Fig. 4) [28] and its high-copy number relative to the nuclear genome. Variation of the mitogenome within and between *Peronosclerospora* species should be assayed to confirm its suitability as a diagnostic marker. Single copy protein coding genes in the nucleus may provide additional sequences for conserved markers; however, high sequence conservation could result in false species assignment. The expanded paralogs and pseudogenes may provide opportunities for lineage-specific markers that are useful for taxonomic and population studies of *Peronosclerospora* spp. [31] because sequences under weaker purifying selection may accumulate unique polymorphisms quicker than sequences under stronger purifying selection. All marker types should be validated to determine their variation within and between species; consequently, additional studies of the mitochondrial and nuclear genome of *Peronosclerospora* spp., including global isolates of *P. sorghi*, are required to efficiently monitor these important pathogens.

## Methods

### Isolate propagation and collection

The P6 isolate of *Peronosclerospora sorghi* was previously collected from Wharton County, Texas [12] and maintained on inoculated plants in a greenhouse at Texas A&M as described previously [32]. Sporangia were collected into distilled water from multiple plants and shipped frozen to UC Davis for DNA extraction and to Dovetail Genomics to generate Hi-C libraries. For RNAseq, pairs of segments from leaves of an infected plant were split longitudinally prior to sporulation. One segment of each pair was placed immediately in RNAlater (ThermoFisher Scientific, Waltham, MA) to characterize the transcriptome pre-sporulation. The second segment was placed in a 20°C incubator overnight to trigger sporulation and then placed in RNAlater. This material was sent to UC Davis for RNA extraction.

### DNA extraction, sequencing, and genome assembly

Genomic DNA was extracted from a pellet of sporangia to generate 10x Genomics linked-read libraries. Briefly, the pellet was vortexed for two minutes in a microcentrifuge tube with ∼200 μL of Rainex-treated beads and 500 µL of 2× extraction buffer (100 mM Tris-HCl pH 8.0, 1.4 M NaCl, 20 mM EDTA, 2% [wt/vol] cetyltrimethylammonium bromide, and Β-mercaptoethanol at 20 μL/mL). After vortexing, the material was transferred to a fresh 2 mL tube and subsequently treated with RNase (20 μL/mL) at 65°C for 30 min. Next, an equal volume of 1:1 phenol/chloroform was added, mixed, and centrifuged at maximum speed (5,200×g) for 15 min. The aqueous phase was retained, mixed with 24:1 chloroform/isoamyl alcohol, and again centrifuged as maximum speed for 15 min. The aqueous phase was mixed with 0.7 volumes of isopropanol and DNA precipitated at −20°C for one hour. DNA was pelleted by centrifuging at maximum speed for 30 min, washed with 70% ethanol, dried, and suspended in 10 mM Tris-HCl. Genomic libraries were constructed using the 10x Genomics Chromium [33] and sequenced on an Illumina x10 at Novogene (Sacramento, CA). All raw data, plus the subsequent assemblies and annotation, are available under NCBI BioProject ######.

Genomic sequence data generated to study the genome of *P. sorghi* included 449,879,160 linked-read pairs (10x Genomics) for isolate P6, equivalent to 294x genome coverage. Genome assemblies were constructed using the 10x Genomics SuperNova v2.0 pipeline [33] and positively filtered for oomycete scaffolds using BLASTn [34, 35]. Optimal assembly conditions using different barcode fractions were determined empirically (Supplementary Table S1). Hi-C libraries were prepared by Dovetail Genomics. The Hi-C libraries were sequenced on a HiSeq 4000 at the UC Davis DNA Technologies Core to yield 321,243,925 Hi-C paired-end reads that were used for scaffolding the Supernova assembly with the Dovetail Genomics Inc. HiRise pipeline [36]. Hi-C interactions were visualized by re-aligning Hi-C reads using bwa mem and plotting contact matrices with HiCExplorer [37]. Centromeres were identified in regions of the assembly enriched for short-distance cis-interactions and regional trans-interactions, resulting in crosses on the Hi-C plot. This is because centromeres of distinct chromosomes may be physically close to one another in Rabl-like conformations [22]. In the initial assembly, some chromosome-scale scaffolds were inconsistent with the expectation of one centromere per scaffold (Fig. S1). These potential misjoins were investigated with Hi-C reads and broken at gaps (strings on unknown bases; N). Coordinates of Hi-C reads were recalculated and enrichment of read-pairs aligned to chromosome arms was sought. Sequences were rejoined so the centromeres formed expected crosses when contact matrices were plotted. Centromere coordinates in the final assembly were defined by identifying 100 kb windows of the genome with short mean cis-interaction distance of Hi-C, first-mate reads. These windows were visualized as scatter plots using ggplot2 [38]. Assembly completeness was evaluated using KAT [39] and BUSCO [40]. Results from the BUSCO analysis were added to a previous phylogenomic analysis [16]. Briefly, peptide sequences for each BUSCO prediction found to be single copy in all 32 species surveyed were aligned independently with MAFFT v7.245 [41]. The aligned sequences were concatenated, and a Maximum Likelihood phylogeny was produced with RAxML v8.2.9 [42] using the PROTGAMMAAUTO model and 1,000 bootstraps. The phylogenetic tree was visualized using Geneious v8.0.5 [43] and labels were italicized in Microsoft PowerPoint. The Newick tree produced by RAxML is available as Supplementary File S2.

### Mitochondrial assembly

Contigs from a *de novo* genomic assembly in CLC Genomics Workbench (v9; Qiagen, Redwood City, CA) were identified as mitochondrial due to sequence similarity with *P. tabacina* mitochondrial sequences (KT893455) by BLAST analysis. These were used as templates for further assembly with SeqMan NGen (v16.0.0, DNASTAR, Madison, WI, USA). The resulting assemblies were evaluated for uniformity and depth of coverage. Contigs were broken when gaps/low coverage or inconsistencies were observed and the set of smaller contigs reassembled using the reference-guided assembly – special workflows assembly option of SeqMan NGen to extend the ends of the contigs and the close gaps. ORFs were predicted and annotated with Geneious v9.1.8 (Biomatters, New Zealand) using the universal genetic code. Encoded products of genes were identified using BLAST [34] analysis against mitochondrial genome sequences published for *Peronospora tabacina* [44]. Genes encoding tRNAs were identified using tRNAscan-SE v1.3.1 [45].

### RNA extraction, sequencing, and transcript assembly

Total RNA was extracted from pairs of infected leaf segments pre- and post-sporulation of *P. sorghi* collected at Texas A&M, using a Qiagen RNeasy kit Cat. No. / ID: 74904, followed by poly-A cDNA generation. Strand-specific libraries were generated using the KAPA mRNA HyperPrep Kit (KR1352 v5.17) per supplier instructions. Fragments were 150 bp paired-end sequenced on a HiSeq 4000 at the UC Davis DNA Technologies Core. Between 37,570,743 and 123,050,654 strand-specific cDNA read pairs were generated across three biological replicates of the two conditions for a total of 290,220,630 cDNA read pairs. The resulting reads were trimmed using bbduk.sh [46], assembled with Trinity v2.4.0 in strand-specific mode [47], and mapped to a combined reference containing the *S. bicolor* [48] and *P. sorghi* assemblies using minimap2 [49]. The pathogen’s transcriptome was defined as those that primarily aligned to *P. sorghi* scaffolds, while the host’s transcriptome was defined as those that primarily aligned to *S. bicolor* scaffolds. Transcriptome completeness for the host and pathogen was evaluated by BUSCO [40] in transcriptome mode. The sense (top) strand of the *P. sorghi* transcriptome was translated into a set of longest open reading frames (ORFs) using TransDecoder.LongOrfs v5.5.0 [50], and conserved Pfam domains were identified using InterProScan [51, 52]. Unmapped transcripts were evaluated with Kraken2 [53] using the NCBI non-redundant protein (nr) database. Results were visualized using KronaTools [54].

### Annotation

Repeats in the genome assembly of *P. sorghi* were defined with RepeatModeler v1.73 [55] and masked with RepeatMasker v4.0.9 [56]. The same library was used to identify repeats in the transcriptome assembly. Gene models were annotated in the genome assembly using MAKER [57], with additional putative effectors identified using hidden Markov models (HMM) with HMMER (ref) and regular expression string searches of ORFs (Fletcher 2018). The MAKER pipeline was provided with the RepeatModeler profile as well as assembled transcripts and translated ORFs from the transcriptome of *P. sorghi*, all described above, plus ESTs (option: altest) and protein sequences of other oomycete species available from NCBI. MAKER was initially run without a SNAP HMM, inferring genes using est2genome and protein2genome. These predictions were used to train a SNAP HMM [58] that was used for a subsequent run of MAKER with both est2genome and protein2genome set to 0. The predicted proteins were again used to train a new SNAP HMM [59]. This process was repeated twice to generate three SNAP HMMs, which were used sequentially in three independent runs of MAKER. The annotations produced were evaluated as previously described [19] to select a single optimal run. This involved contrasting the number of gene models predicted, mean protein length, BLASTp hits to other oomycete annotations, and Pfam domains annotated by InterProScan [51, 52]. Annotation of genes encoding putative effectors was performed as previously described [19]. Briefly, the entire genome was translated into ORFs. These ORFs were surveyed for secretion signals using SignalP3.1 and SignalP4.0, and CRN motifs of LWY domains using HMMs. For peptides with secretion signals, the 60 residues beyond the predicted cleavage site were surveyed for an RXLR or RXLR-like motif and subsequently for a downstream EER or EER-like motif. ORFs encoding peptides that were predicted to be secreted and contained an (L)WY domain or a CRN motif were considered high-confidence putative effectors (HCPEs). ORFs encoding peptides that were predicted to be secreted and encoded an RXLR and EER domain, but did contain an (L)WY domain, or encoding peptides not predicted to be secreted, but contained an (L)WY domain, or a CRN motif were considered low-confidence putative effectors (LCPEs). The putative effectors and MAKER annotations were reconciled so that annotations did not overlap on the same strand. This was performed so that 1) any HQE or LQE annotations that did not overlap a MAKER annotation were added to the master annotation; for *P. sorghi* this was 12 HCPEs and 122 LCPEs. 2) HCPEs that overlapped MAKER annotations with the same start coordinates but earlier stop coordinates were discarded; for *P. sorghi* this was six peptides. 3) HCPEs that overlapped MAKER annotations with the same start coordinates but later stop coordinates replaced the model proposed by MAKER if they had a higher BLASTp score to the NCBI nr database than the overlapping MAKER model; for *P. sorghi* this was six peptides. 4) HCPEs that overlapped MAKER annotations but had different start coordinates and later or identical stop coordinates were retained over proposed MAKER models; for *P. sorghi* this was 27 peptides. 5) HCPEs that overlapped MAKER annotations but had different start coordinates and earlier stop coordinates were investigated to determine if the MAKER model should have a modified start coordinate; for *P. sorghi* this was six peptides. 6) Any LCPEs that overlapped MAKER annotations were discarded; for *P. sorghi* this was 142 ORFs. The same effector prediction workflow was then applied to the reconciled annotation set to determine the reported effector counts. Tracks for repeats, transcript coverage, annotation, and effector annotations were generated in 100 kb windows along each chromosome using Bedtools v2.29.2 and plotted using Circos [60].

### Comparative Genomics

Predicted peptide sequences of *P. sorghi* were used in an orthology analysis including another 39 genome assemblies of 34 different species (Supplementary Table S2). The number of genes for each assembly surveyed was calculated by counting the number of FASTA entries in the peptide file. Orthology was calculated using OrthoFinder v2.2.1 [61]. Resulting orthogroups were filtered for interspecies orthogroups, which required assignment of proteins from at least two different oomycete species. The number of inter-specific orthogroups assigned to each assembly and the number of proteins represented in these orthogroups was calculated. Interspecies orthogroups were then summarized at different taxonomic levels for visualization. For the purposes of this study, core orthogroups had proteins from *P. sorghi* and at least one other downy mildew clade 2 species; one downy mildew clade 1 species; one *Phytophthora* spp.; one species in the Family Pythiaceae; one species in the Family Albuginaceae; one species in the Order Saprolegniales. For all orthogroups, a ratio of *P. sorghi* to *P. effusa* gene assignment was calculated, ordered, and plotted as a heatmap using ggplot2 [38]. Single copy orthologs between *P. sorghi* and *P. effusa* were identified as members of orthogroups assigned exactly one protein sequence from *P. sorghi*, one from *P. effusa*. Coordinates of the genes encoding these protein sequences were extracted from GFF files and plotted as links using Circos [60], scatter plots using ggplot2 to establish synteny, or vertical lines using ggplot2 to establish centromere conservation. Genes expanded in *P. sorghi* relative to *P. effusa* (notated throughout as >1:1) were identified as members of interspecific orthogroups assigned more than one *P. sorghi* sequence and exactly one *P. effusa* sequence. Genes expanded in *P. effusa* relative to *P. sorghi* (notated throughout as 1:>1) were identified as members of interspecific orthogroups assigned exactly one *P. sorghi* sequence and more than one *P. effusa* sequence. Genes expanded in both (>1:>1) were identified as members of interspecific orthogroups containing multiple *P. sorghi* and multiple *P. effusa* sequences. Genes absent in *P. effusa* but present as single-copy (1:0) or multi-copy in *P. sorghi* (>1:0) were identified as members of interspecific orthogroups lacking *P. effusa* annotations with either one or multiple *P. sorghi* annotations, respectively. Genes not assigned to interspecific orthogroups were classified as lacking orthology for the purpose of this study.

Genes expression was investigated using the six RNAseq data sets generated here (three replicates of two conditions; see above). A single reference containing the assembled sequence and annotation of *P. sorghi* and *S. bicolor* (GCF_000003195.3; [48]) using STAR v2.7.9 in runMode genomeGenerate. Paired-end RNAseq data was aligned to this reference using STAR with in quantMode GeneCounts setting sjdbOverhang to 99. Transcript abundance was determined by calculating the FPKM across all generated RNAseq reads. The total exon length of each gene was calculated by summing the length of annotated exons in the GFF file. The raw counts generated by STAR were summed for each gene and scaled to counts per million (CPM). This value was further scaled by the total exon length for each gene to obtain the FPKM. Similar results were obtained when the CPM was used instead of FPKM (Supplementary Fig. S6). Analysis of variance (ANOVA) was conducted in R using aov(FPKM ∼ Gene Classification) and pairwise comparisons generated using Tukey’s honestly significant difference (HSD) [62] in R using TukeyHSD(). Groups of significance were identified using multcompView [63] in R. Dot plots showing the synteny and transcript abundance of *P. sorghi* genes were plotted in R ggplot() and geom_point() [38]. Scatter plots showing the transcript abundance for secreted and non-secreted genes in each category were plotted using ggplot() and geom_jitter() [38] setting height to 0. DGE analysis was conducted using edgeR [64]. Read counts of pathogen and host genes were independently analyzed in R. Genes with fewer than five CPM were dropped from the analysis and normalization was conducted on the reduced data set using calcNormFactors(). Multidimensional scaling coordinates were obtained using plotMDS() and plotted using ggplot() [38]. A design matrix was constructed using limma [65] model.matrix(), dispersion estimated using estimateDisp() from edgeR [64], and a negative binomial generalized linear model was fit using glmFit(). The two conditions were then compared using makeContrasts() and a negative binomial generalized log-linear model was fit using glmLRT(). Differentially regulated genes were identified as those that have an adjusted p-value <0.05. For each gene, the log fold change and -log10 adjusted p-value was plotted in R using ggplot() and geom_point(). For the host, the function of differentially regulated genes was investigated by identifying the annotated product produced in the GFF file. For the pathogen, the function of differentially regulated genes was investigated by identifying conserved Pfam domains annotated by InterProsScan [51, 52]. Chi-squared tests were run in R using chisq.test().

The dN/dS for genes annotated in *P. sorghi* that had single copy orthologs in *P. effusa* was performed if the orthologs had a blastp alignment coverage greater than 20%. Pairwise codon-based alignments were conducted for each pair of orthologs using PRANK. The dN/dS for each alignment was calculated using codeml in pairwise mode [66]. This analysis was run as pairwise instead of per orthogroup because the *P. sorghi* genes assigned to the same orthogroup may cover different parts of the *P. effusa* ortholog, which resulted in codeml using a smaller portion of sequence for the calculation. An ANOVA for dN/dS of different gene classifications was calculated as above using Tukey’s HSD. Data was plotted as a raincloud plot in R using ggplot() [38, 67].

The length of each peptide sequence annotated in the genome of *P. sorghi* was obtained by building an index file with SAMtools faidx [68]. The mean exon count of each gene was obtained by counting the number of exon entries per annotation in the GFF file. An ANOVA was run to compare the mean encoded peptide length and mean exon count for each of the gene categories as described above. The length of every intron encoded in the genome of *P. sorghi* was determined from the GFF file and plotted as a histogram using ggplot() [38]. Histograms were colored depending on whether transcripts could be detected (FPKM >0) for the gene from which the intron originates.

Compartmentalization of the *P. sorghi* genome assembly was determined by calculating the 5’ and 3’ intergenic distances between genes on chromosomal scaffolds [69, 70]. This information was extracted and oriented from the GFF file. Density of intergenic distances for genes with FPKM equal to 0 and FPKM >0 were plotted on a log10-scaled heatmap using ggplot() and geom_bin2d() [38] with 40 bins on each axis. A histogram of intergenic distance for each flank was plotted for gene categories defined by orthology with *P. effusa* using ggplot() and geom_histogram() [38]. Histograms were colored by the presence or absence of transcript detection.

## Supporting information

Supplemental File 1: Compressed HTML of Krona output, classifying unaligned transcripts with KRAKEN2

Supplemental File 2: Newick format phylogenetic tree.

## Acknowledgments

We thank H. Xu (UC Davis) for raw data submissions to NCBI, and E. Georgian (UC Davis) for editorial services. The sequencing was carried out by the DNA Technologies and Expression Analysis Cores at the UC Davis Genome Center, supported by NIH Shared Instrumentation Grant 1S10OD010786-01. The bioinformatic analysis was carried out using the UC Davis LSSC0 High Performance Computing cluster maintained by the UC Davis Bioinformatics Core.

## Data availability

All raw reads and the nuclear genome assembly and annotation are available at NCBI under BioProject PRJNA845776. The mitochondrial genome and annotation is available at NCBI under accession #######.

## Supplementary Information

**Table S1. Intermediate assembly statistics.**

**Table S2. RNAseq Read count of *P. sorghi* genes used for differential gene expression analysis.**

**Table S3. List of oomycete assemblies used for orthology analysis.**

**Table S4. Pfam domains encoded by *P. sorghi* genes identified as differentially regulated**

**Fig. S1. Hi-C contact matrix of the genome assembly produced by HiRise.** The strong diagonal reflects the high contact frequency between physically close sequences, indicating their correct linear order along each chromosome-scale scaffold. Cross patterns along the x- and y-planes are indicative of high frequencies of trans contacts between centromeres and are likely due to the Rabl-like chromosome configurations. Labels highlighted red indicate misassembled sequences, evident from the cross pattern within the scaffold.

**Fig. S2. Mitochondrial genome of *Peronosclerospora sorghi* isolate P6.** The map starts at the gene encoding the large subunit of the mitochondrial ribosome to parallel Fig. 4. Colored bars indicate the genes encoded in the mitogenome to scale. Genes colored red are ribosomal, purple are tRNAs, and green are protein coding. The arrow indicates the direction of transcription.

**Fig. S3. Distribution of dispersed effectors in the genome of *Peronosclerospora sorghi*.** A–E are like Fig. 3, but only *P. sorghi* data is plotted. F) Links indicate effectors with peptide similarity as detected by CD-Hit. Red links join pairs of RXLR effectors assigned to the same cluster, which required a 40% identity to the centroid sequence. Blue links join pairs of Crinkler effectors assigned to the same cluster, which required a 70% identity to the centroid sequence.

**Fig. S4. Scatter plot showing the coordinates of genes from the genomes of *P. effusa* and *P. sorghi* assigned to orthogroups as multicopy in both.** The x-axis is ordered to demonstrate collinearity, like Fig. 3H. Axes are not scaled to size. Grey gridlines indicate 50 Mb boundaries. Blue dotted lines indicate chromosome boundaries. Points are colored by their transcript abundance (FPKM). Black dots indicate 0 FPKM.

**Fig. S5. Transcript abundance was bimodal for genes expanded in *P. sorghi* on non-syntenic chromosomes relative to *P. effusa*.**

**Fig. S6. Scatter plots showing the read counts per million for genes annotated in *P. sorghi*.** Genes were assigned to different categories (x-axis) based on their orthology to *P. effusa*. Genes expanded in *P*. *sorghi* (>1:1) were sub-categorized based on their synteny with *P. effusa*. Genes encoding peptides predicted to be secreted were plotted in red for each category. Lowercase letters indicate different groups of significance as calculated by Tukey’s HSD test. Plot is similar to Fig. 9D, which plotted fragments per kilobase of exon per million mapped fragments.

**File S1. Metagenomic classification of unassigned, assembled transcripts.** Interactive HTML produced using Krona tools. Very few of the unassigned transcripts were assigned to the Oomycota.

**File S2. Maximum likelihood tree calculated by RAxML**. Newick format file used to generate Fig. 2.

