## Supplementary Figures S1 to S6 for "The genome of the oomycete *Peronosclerospora sorghi*, a cosmopolitan pathogen of maize and sorghum, is inflated with dispersed pseudogenes"

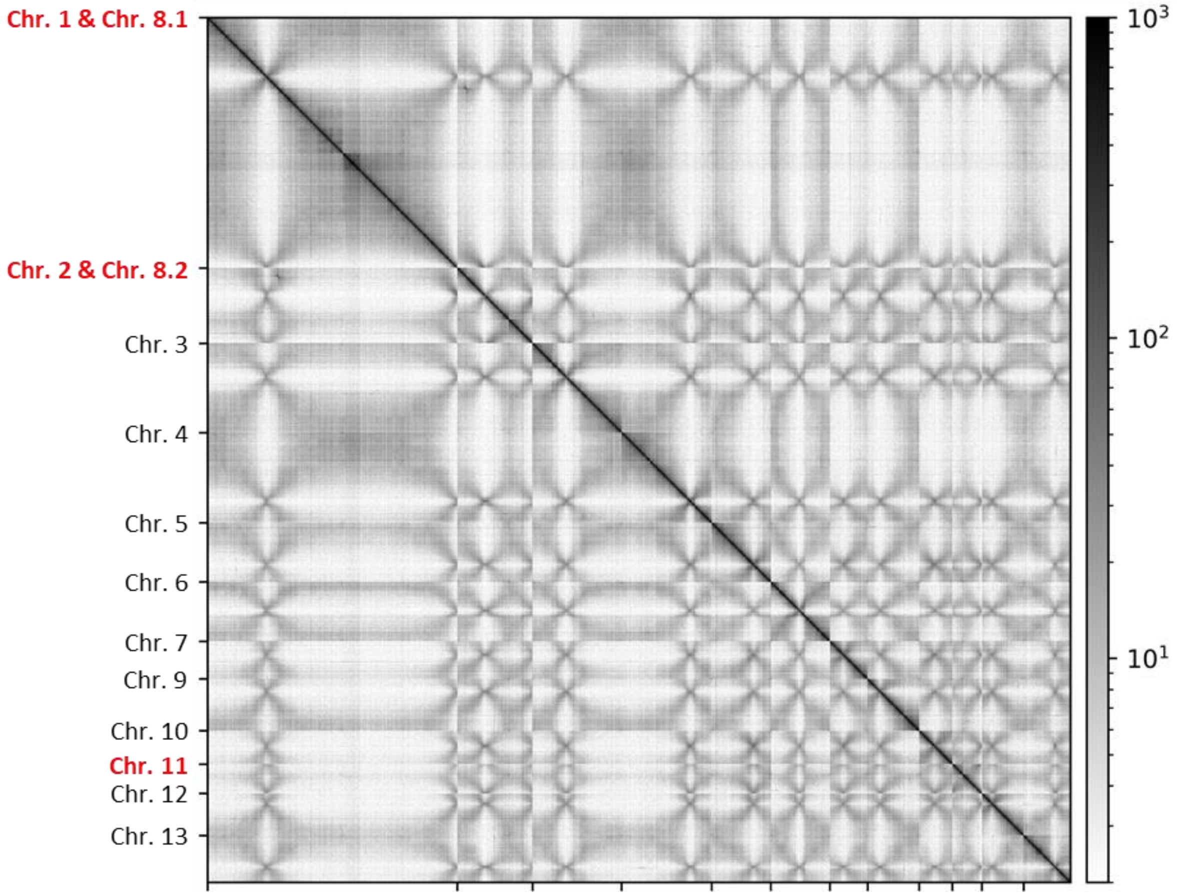


**Fig. S1. Hi-C contact matrix of the genome assembly produced by HiRise.** The strong diagonal reflects the high contact frequency between physically close sequences indicating their correct linear order along each chromosome-scale scaffold. Cross patterns along the x- and y-planes are indicative of high frequencies of trans contacts between centromeres and are likely due to the Rabl-like chromosome configurations. Labels highlighted red indicate misassembled sequences, evident from the cross pattern within the scaffold.

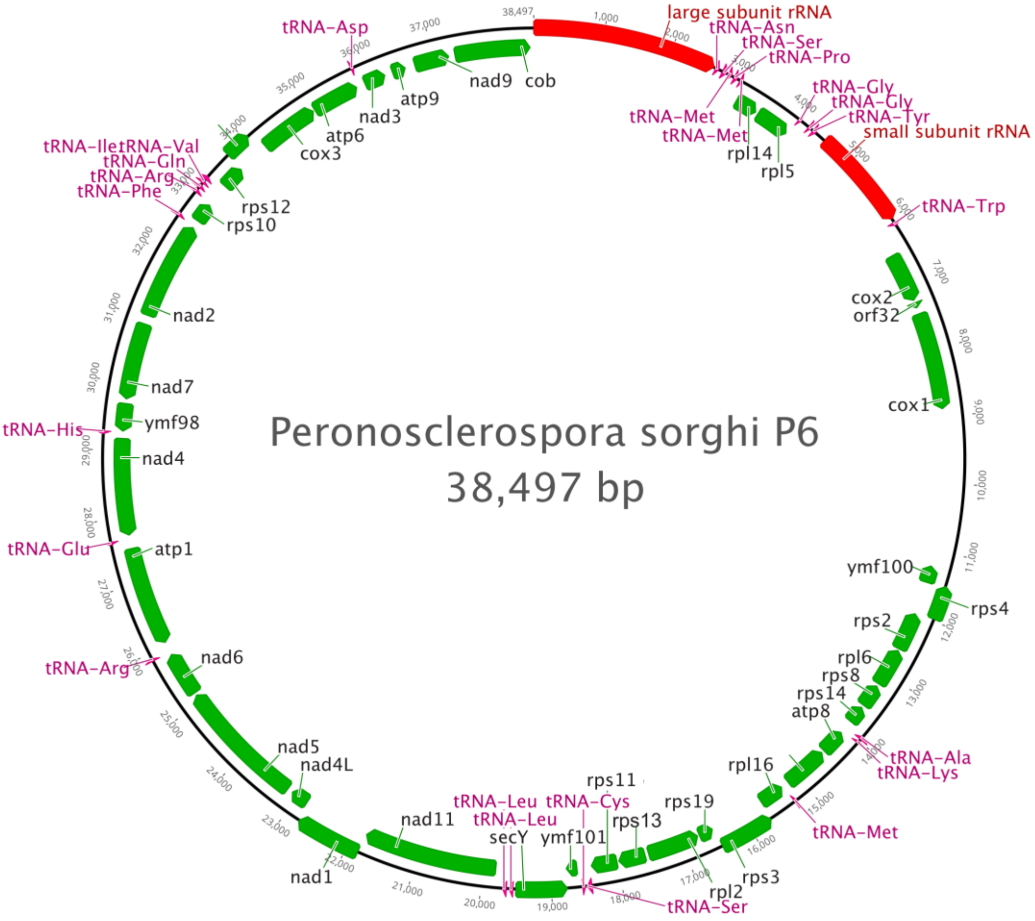


**Fig. S2. Mitochondrial genome of *Peronosclerospora sorghi* isolate P6.** The map starts at the gene encoding the large subunit of the mitochondrial ribosome to parallel Fig. 4. Colored bars indicate the genes encoded in the mitogenome to scale. Genes colored red are ribosomal, purple are tRNAs, and green are protein coding. The arrow indicates the direction of transcription.

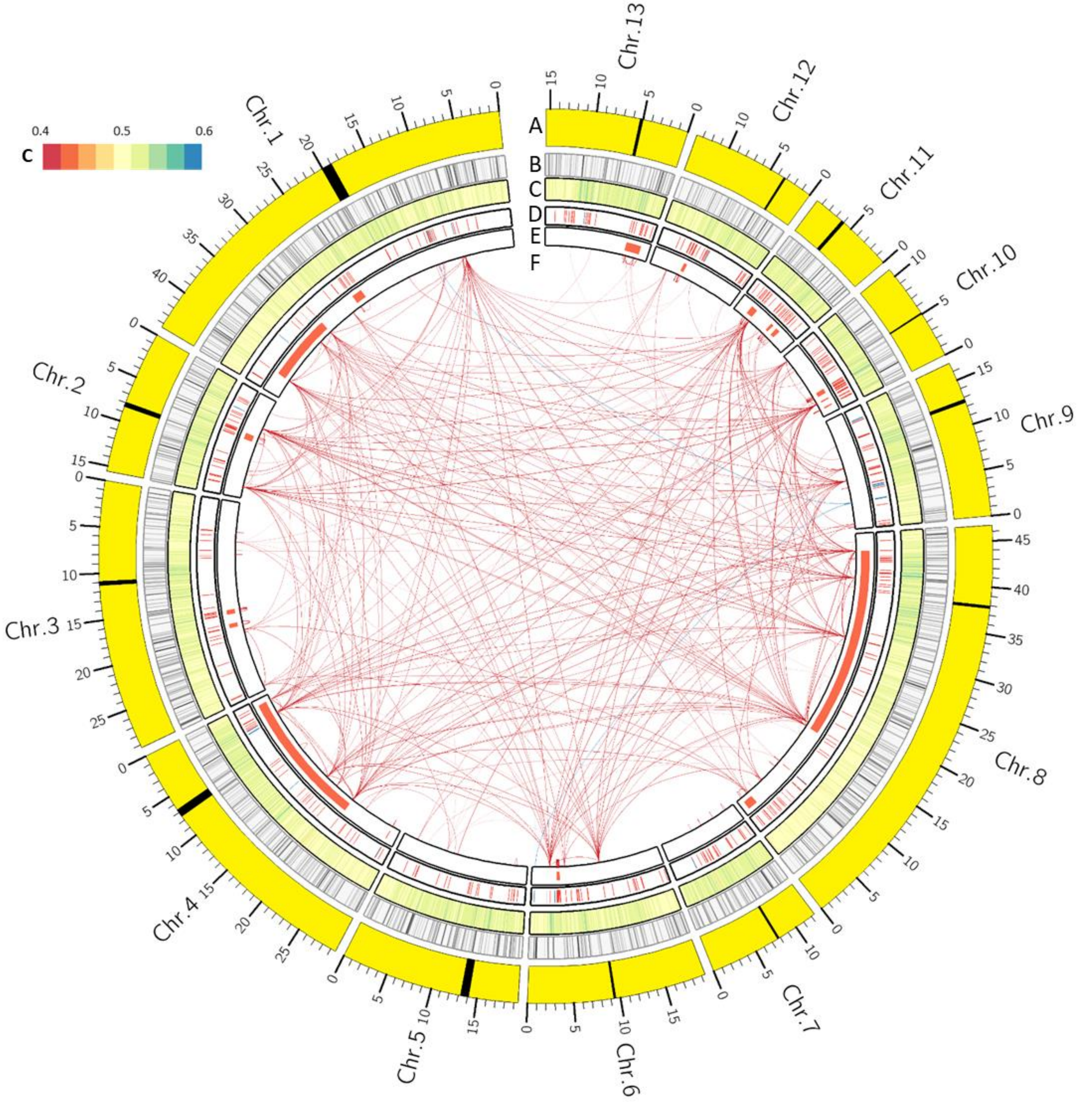


**Fig. S3. Distribution of dispersed effectors in the genome of *Peronosclerospora sorghi*.** A–E are the same as Fig. 3, but only *P. sorghi* data is plotted. F) Links indicate effectors with peptide similarity as detected by CD-Hit. Red links join pairs of RXLR effectors assigned to the same cluster, which required a 40% identity to the centroid sequence. Blue links join pairs of Crinkler effectors assigned to the same cluster, which required a 70% identity to the centroid sequence.

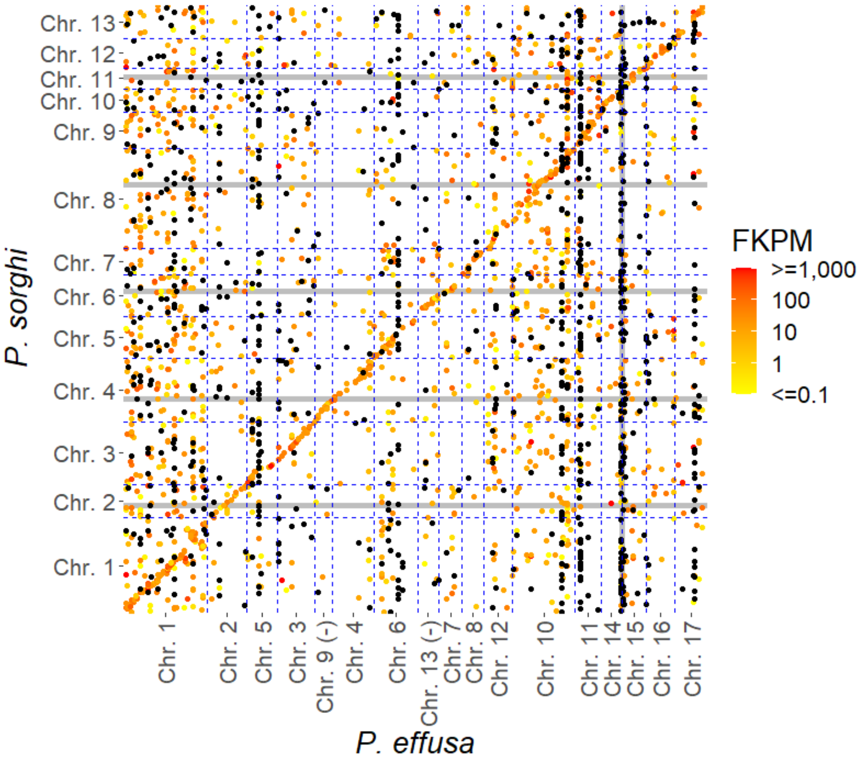


**Fig. S4. Scatter plot showing the coordinates of genes from the genomes of *P. effusa* and *P. sorghi* assigned to orthogroups as multicopy in both**. The x-axis is ordered to demonstrate collinearity, like Fig. 3H. Axes are not scaled to size. Grey gridlines indicate 50 Mb boundaries. Blue dotted lines indicate chromosome boundaries. Points are colored by their transcript abundance (FPKM). Black dots indicate 0 FPKM.

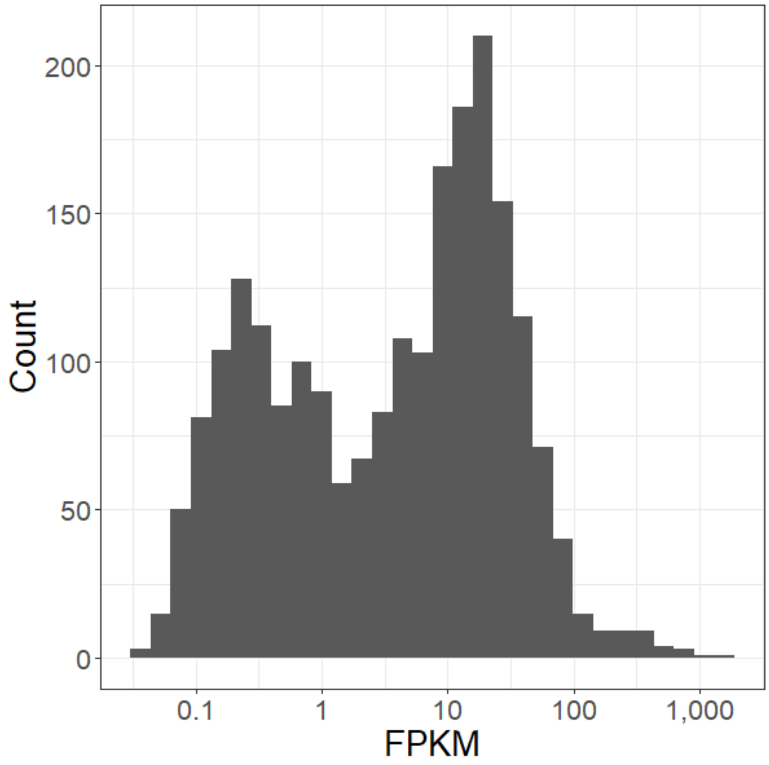


**Fig. S5. Transcript abundance was bimodal for genes expanded in *P. sorghi* on non-syntenic chromosomes relative to *P. effusa*.**

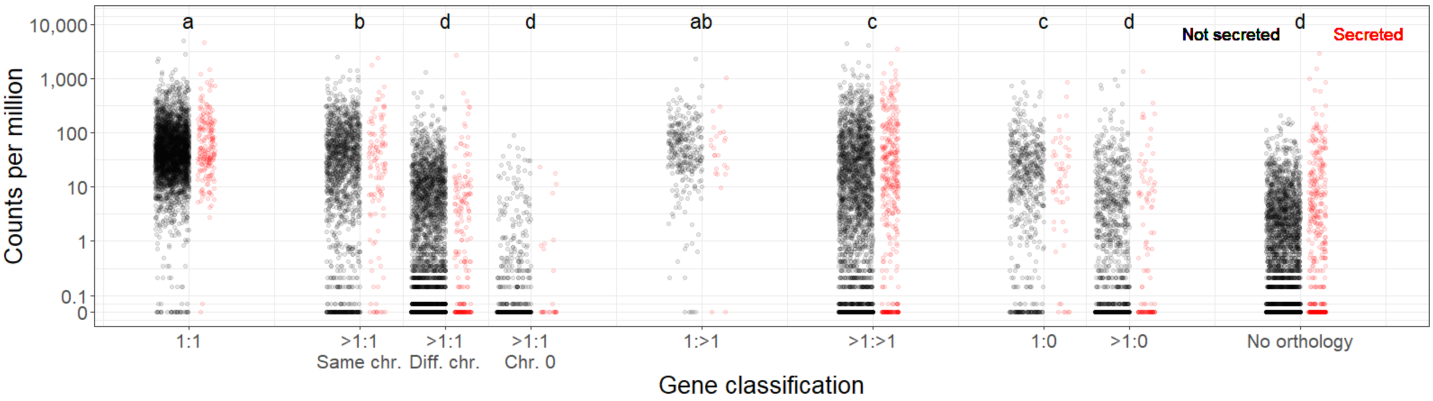


**Fig. S6. Scatter plots showing the read counts per million for genes annotated in *P. sorghi*.** Genes were assigned to different categories (x-axis), based on their orthology to *P. effusa*. Genes expanded in *P. sorghi* (>1:1) were sub-categorized based on their synteny with *P. effusa*. Genes encoding peptides predicted to be secreted were plotted in red for each category. Lower case letters indicate different groups of significance as calculated by Tukey's HSD test. Plot is similar to Fig. 8D which plotted fragments per kilobase of exon per million mapped fragments.
