## Supplemental File 1: Compressed HTML of Krona output, classifying unaligned transcripts with KRAKEN2 for "The genome of the oomycete *Peronosclerospora sorghi*, a cosmopolitan pathogen of maize and sorghum, is inflated with dispersed pseudogenes": SupplementaryFileS1.html

Javascript must be enabled to view this page.

members
magnitude
magnitudeUnassigned
count
unassigned
taxon
rank

KrakenDefault\_nt

node0.members.0.js
7769
365281

71805
node1.members.0.js

15
2787823
no rank

15
12908
no rank

151659
15
no rank

no rank
1
44770

1
44909
species
node6.members.0.js

5
81490
no rank

198431
node8.members.0.js
species
5

9
node9.members.0.js
species
155900

235
4
10239
superkingdom
node10.members.0.js

family
687329
2

no rank
363628
2

432261
species
node13.members.0.js
1

1
species
node14.members.0.js
68887

10482
2
family

10485
2
genus

no rank
2
357261

1
1836595
node18.members.0.js
species

1
452647
species

1
no rank
node20.members.0.js
452648

4
2731342
clade

1
2732091
kingdom

phylum
2732412
1

class
1
2732413

order
1
2732414

family
10841
1

1
117574
no rank

species
node28.members.0.js
2202644
1

3
2732092
kingdom

3
2732416
phylum

2732424
2
class

2
2732539
order

10811
2
family

genus
10814
2

1
node35.members.0.js
species
223287

2712833
node36.members.0.js
species
1

class
1
2732423

order
2732536
1

1
39724
family

no rank
1
742826

species
node41.members.0.js
1379717
1

12333
9
no rank

clade
2693321
1

2693720
species
node44.members.0.js
1

1
284052
node45.members.0.js
species

6
38018
node46.members.0.js
species

1
2827298
species
node47.members.0.js

12429
9
no rank

1
node49.members.0.js
species
1685769

no rank
2204151
8

no rank
1
35342

1
2058757
family

clade
1
2502018

2569970
1
no rank

node55.members.0.js
species
2761302
1

51368
6
no rank

genus
3
2060084

1349410
species
node58.members.0.js
3

2591644
node59.members.0.js
species
3

node60.members.0.js
species
2714171
1

84
2559587
clade

no rank
17
2585030

12
1922348
no rank

12
species
node64.members.0.js
1922378

439490
1
no rank

1
35278
clade

1
species
node67.members.0.js
2605289

species
node68.members.0.js
2743153
4

52
2732396
kingdom

2732407
21
phylum

class
2732498
2

2732502
2
order

family
186766
2

1077831
2
no rank

2819083
species
node75.members.0.js
2

class
2732500
17

17
2732504
order

2560063
17
family

no rank
1
2587553

1
2719868
node80.members.0.js
species

16
186783
genus

2
16
no rank
node82.members.0.js
1629669

3
species
node83.members.0.js
2741767

1
species
node84.members.0.js
2741861

1
node85.members.0.js
species
2741761

2741811
node86.members.0.js
species
1

species
node87.members.0.js
2741810
1

2741797
species
node88.members.0.js
3

2686482
node89.members.0.js
species
1

1
node90.members.0.js
species
2741780

1
2741850
node91.members.0.js
species

node92.members.0.js
species
2741812
1

2732499
2
class

order
2732503
2

2
2732892
family

genus
2
186768

343865
2
no rank

2
2587224
species
node98.members.0.js

5
2732405
phylum

4
2732459
class

2732541
4
order

10880
4
family

subfamily
3
689832

10912
3
genus

species
2
28875

2
no rank
node106.members.0.js
10967

species
1
28876

1
335102
no rank

1
no rank
node109.members.0.js
449582

1
36446
no rank

1
node111.members.0.js
species
907191

class
2732458
1

order
2732540
1

1
249310
family

genus
2560100
1

1
2560291
species

no rank
node117.members.0.js
2066695
1

2732406
11
phylum

class
2732461
7

order
1
2732543

1
2560066
family

genus
1
11040

2846071
1
species

node124.members.0.js
no rank
11041
1

order
675063
5

1
675068
family

subfamily
1914297
1

1
12163
genus

1
12172
species

12458
no rank
node130.members.0.js
1

family
4
675064

genus
4
12176

species
node133.members.0.js
35286
1

1
112228
species
node134.members.0.js

2
node135.members.0.js
species
12177

2732544
1
order

69973
1
family

217160
1
genus

node139.members.0.js
species
47985
1

class
2732463
2

2
2732548
order

family
39738
2

2560072
1
subfamily

1
39734
genus

1
473783
no rank

species
node146.members.0.js
2908239
1

no rank
1
137555

1
2735226
node148.members.0.js
species

class
2
2732462

order
2732545
2

family
11050
2

genus
2
11102

node153.members.0.js
species
11103
2

phylum
2497569
3

2497571
2
subphylum

class
2
2497577

order
2
2499411

family
11308
2

genus
197912
1

11520
node160.members.0.js
species
1

197911
1
genus

species
11320
1

no rank
1
102797

1929075
node164.members.0.js
no rank
1

subphylum
1
2497570

class
2497574
1

11157
1
order

1
11244
family

genus
1
1868215

11250
1
species

1
208893
no rank
node171.members.0.js

2732408
12
phylum

class
2732506
12

order
5
2732553

2169577
5
family

no rank
2600328
5

2715208
node177.members.0.js
species
5

76804
3
order

suborder
3
2499399

3
11118
family

subfamily
2501931
3

693996
3
genus

1
2509505
subgenus

1
28295
species
node184.members.0.js

2509514
2
subgenus

species
2
693997

no rank
node187.members.0.js
11153
2

4
464095
order

family
1
232795

1
no rank
node190.members.0.js
336635

2
675074
no rank

2
1955153
node192.members.0.js
species

12058
1
family

12059
1
genus

species
1
138950

12089
serotype
node196.members.0.js
1

kingdom
2732397
15

phylum
15
2732409

class
2732514
15

15
2169561
order

family
15
11632

14
327045
subfamily

140052
1
genus

347957
1
no rank

species
node205.members.0.js
2594425
1

11646
13
genus

species
node207.members.0.js
11676
12

species
node208.members.0.js
11665
1

no rank
1
35276

1435008
species
node210.members.0.js
1

10
2732004
clade

kingdom
10
2732005

10
2732007
phylum

class
2732523
10

2732524
2
order

family
10501
2

genus
1
181083

1
346674
no rank
node218.members.0.js

1
181086
genus

no rank
358403
1

1
251749
node221.members.0.js
species

2732554
3
order

family
3
549779

no rank
1
1737587

species
node225.members.0.js
1737588
1

985780
2
no rank

1
985782
species
node227.members.0.js

2487768
species
node228.members.0.js
1

order
2732555
5

family
944644
5

1513461
5
no rank

2506606
species
node232.members.0.js
4

1
species
node233.members.0.js
2506605

family
1
185751

genus
1
147262

1
12893
species
node236.members.0.js

110
2731341
clade

kingdom
110
2731360

phylum
2731361
3

2731363
3
class

3
548681
order

family
3
10292

subfamily
3
10357

10358
3
genus

1535247
node245.members.0.js
species
2

1
50290
species
node246.members.0.js

phylum
2731618
107

class
2731619
107

node249.members.0.js
order
28883
3
107

102294
1
no rank

species
node251.members.0.js
2202568
1

10699
63
family

subfamily
1
2842522

2852964
1
no rank

1147042
species
node255.members.0.js
1

genus
2733186
1

species
1
2734244

no rank
node258.members.0.js
2283015
1

2133759
1
no rank

1
1407671
node260.members.0.js
species

1623306
1
genus

no rank
2562717
1

1
2024005
node263.members.0.js
species

196894
51
no rank

2686309
node265.members.0.js
species
1

1
2825654
species
node266.members.0.js

1
species
node267.members.0.js
2826389

2827838
species
node268.members.0.js
1

1
1775140
species
node269.members.0.js

2825740
species
node270.members.0.js
1

2170413
species
node271.members.0.js
44

2825325
node272.members.0.js
species
1

genus
1623304
2

2
node274.members.0.js
no rank
2231643

1
2560121
genus

species
1
2560782

2044561
node277.members.0.js
no rank
1

5
1910976
subfamily

5
1910991
node279.members.0.js
genus

family
2
2560065

no rank
2
2572872

2
species
node282.members.0.js
2831613

family
26
10662

1
2842519
subfamily

genus
1
2842820

2843637
1
species

1
no rank
node287.members.0.js
2686289

19
2
196896
node288.members.0.js
no rank

species
node289.members.0.js
2202564
15

species
node290.members.0.js
1493513
1

1
2769337
node291.members.0.js
species

subfamily
1198136
2

1913651
2
genus

no rank
2
2315206

2836116
species
node295.members.0.js
2

genus
1
2733088

node297.members.0.js
no rank
2788002
1

2
680115
genus

no rank
node299.members.0.js
1231361
2

genus
1
2843389

no rank
2852399
1

species
node302.members.0.js
2878008
1

10744
11
family

10
196895
no rank

10
node305.members.0.js
species
2202567

1984798
genus
node306.members.0.js
1

1
2842329
family

2842641
1
genus

2844071
1
species

1
node310.members.0.js
no rank
1563661

2
node311.members.0.js
superkingdom
9103
804

phylum
23
32066

class
23
203490

order
203491
23

family
203492
16

848
genus
node316.members.0.js
16
2

1755100
species
node317.members.0.js
1

no rank
2648384
2

species
node319.members.0.js
2764326
2

7
2
851
species
node320.members.0.js

node321.members.0.js
subspecies
76859
4
5

strain
node322.members.0.js
457405
1

1
861
species

1
469617
node324.members.0.js
strain

2
2663009
node325.members.0.js
species

species
1
850

469616
strain
node327.members.0.js
1

1
7
node328.members.0.js
family
1129771

6
32067
genus

node330.members.0.js
species
157687
2

2633022
node331.members.0.js
no rank
3
2

712357
species
node332.members.0.js
1

1
554406
node333.members.0.js
species

1
200940
phylum

class
1
67799

order
1
188710

family
188711
1

1
1740
genus

species
1295609
1

1
node340.members.0.js
strain
795359

203691
21
phylum

1
1130380
no rank

2202144
node343.members.0.js
species
1

203692
20
class

order
1643686
1

family
1
143786

29521
genus
node347.members.0.js
1

order
4
1643688

family
170
4

4
1
171
genus
node350.members.0.js

1
2564040
node351.members.0.js
species

2633828
1
no rank

1
2838237
node353.members.0.js
species

node354.members.0.js
species
408139
1

order
136
15

family
1643685
2

genus
2
64895

29518
2
species

2
390236
strain
node359.members.0.js

2845253
8
family

8
1
157
node361.members.0.js
genus

1
node362.members.0.js
species
744512

2638727
1
no rank

166
species
node364.members.0.js
1

species
88058
1

545694
strain
node366.members.0.js
1

3
node367.members.0.js
species
158

species
node368.members.0.js
221027
1

3
2791015
family

genus
3
399320

species
1131703
3

3
node372.members.0.js
strain
158189

family
2
137

1
146
genus

node375.members.0.js
species
46355
1

1911556
1
genus

species
55206
1

573413
node378.members.0.js
strain
1

774
1
1783270
node379.members.0.js
clade

61
142182
phylum

no rank
234665
3

3
2026742
node382.members.0.js
species

142185
1
no rank

1
203437
node384.members.0.js
species

219685
57
class

57
219686
order

57
2
219687
node387.members.0.js
family

21
1
173479
node388.members.0.js
genus

1
1379270
species
node389.members.0.js

no rank
3
221168

3
node391.members.0.js
species
221169

3
173480
species
node392.members.0.js

13
2732249
node393.members.0.js
species

genus
1706036
34

32
861299
node395.members.0.js
species

2
1706037
no rank

2
node397.members.0.js
species
1706038

711
68336
clade

phylum
1662329
2

2
species
node400.members.0.js
2447898

phylum
1134404
4

4
1852932
no rank

2026749
node403.members.0.js
species
4

976
phylum
node404.members.0.js
705
5

1100069
4
order

family
563843
4

1196022
1
no rank

1
1779382
node408.members.0.js
species

146918
2
genus

node410.members.0.js
species
146919
2

genus
1
29548

1
node412.members.0.js
species
29549

43
1853228
class

1853229
43
order

1
43
node415.members.0.js
family
563835

genus
8
79328

4
2
2619133
no rank
node417.members.0.js

1
species
node418.members.0.js
2829818

1
node419.members.0.js
species
2853437

2
2703787
node420.members.0.js
species

2
1
79329
node421.members.0.js
species

485918
node422.members.0.js
strain
1

3
569836
no rank

3
species
node424.members.0.js
1869212

1
649461
genus

2630913
1
no rank

1
2760713
node427.members.0.js
species

2
1874621
genus

2
1813871
node429.members.0.js
species

1004303
genus
node430.members.0.js
8
1

4
563172
node431.members.0.js
species

2647248
3
no rank

3
species
node433.members.0.js
2875540

genus
354354
3

node435.members.0.js
species
354356
3

genus
2
1860196

2
661488
species
node437.members.0.js

1
379899
genus

species
1
446683

929713
strain
node440.members.0.js
1

genus
2812019
2

2
2502779
species
node442.members.0.js

398041
genus
node443.members.0.js
11
2

1492898
species
node444.members.0.js
4

5
node445.members.0.js
species
661481

1
2698688
genus

1
node447.members.0.js
species
2315862

1
50742
no rank

1
358068
node449.members.0.js
species

no rank
8
171558

order
2923288
8

genus
8
2923290

8
species
node453.members.0.js
2875962

768503
214
class

node455.members.0.js
order
768507
2
214

2762286
6
family

genus
396811
4

2
species
node458.members.0.js
2810512

no rank
2685541
2

2904245
node460.members.0.js
species
1

2904246
node461.members.0.js
species
1

genus
2
1433993

2
node463.members.0.js
species
2321403

family
25
2896860

1
319458
genus

1
316068
species

649349
strain
node467.members.0.js
1

120831
genus
node468.members.0.js
17
1

2747268
species
node469.members.0.js
7

no rank
7
2625061

4
species
node471.members.0.js
2861765

species
node472.members.0.js
538966
3

species
94254
2

strain
node474.members.0.js
471854
2

2
861914
genus

no rank
2620963
2

node477.members.0.js
species
1834519
2

1949218
node478.members.0.js
genus
1

2
312278
genus

species
312279
2

2
929562
strain
node481.members.0.js

105
2
genus

species
node483.members.0.js
2259595
2

family
1937968
1

genus
1
1937972

1
999
species

1
880071
node487.members.0.js
strain

family
3
200667

59739
3
genus

species
node490.members.0.js
373891
1

node491.members.0.js
species
2494373
2

563798
11
family

68288
2
genus

species
2
104

2
880070
strain
node495.members.0.js

4
280472
genus

species
4
280473

strain
node498.members.0.js
758820
4

genus
1
232244

species
232259
1

1
866536
node501.members.0.js
strain

1
390846
genus

no rank
1
2621165

node504.members.0.js
species
2859768
1

579921
3
no rank

species
node506.members.0.js
2099675
3

family
node507.members.0.js
1853232
3
133

genus
7
323449

no rank
2648980
1

node510.members.0.js
species
2571030
1

2694930
node511.members.0.js
species
1

2
species
node512.members.0.js
2694929

3
species
node513.members.0.js
400092

genus
299566
5

2745197
node515.members.0.js
species
4

2086471
node516.members.0.js
species
1

116
22
89966
genus
node517.members.0.js

2607656
species
node518.members.0.js
3

2615202
no rank
node519.members.0.js
58
3

2596915
species
node520.members.0.js
6

5
node521.members.0.js
species
2735321

1484116
node522.members.0.js
species
9

node523.members.0.js
species
1385663
5

4
2835648
node524.members.0.js
species

6
2584940
node525.members.0.js
species

2761579
species
node526.members.0.js
2

node527.members.0.js
species
2675878
5

2
node528.members.0.js
species
1484118

2714932
species
node529.members.0.js
2

node530.members.0.js
species
1356852
3

2675877
node531.members.0.js
species
5

1385664
species
node532.members.0.js
1

node533.members.0.js
species
1446467
3
4

1
node534.members.0.js
strain
1227739

3
1385715
node535.members.0.js
species

14
species
node536.members.0.js
1411621

4
2319843
species
node537.members.0.js

no rank
170015
1

1
170016
species
node539.members.0.js

2502781
species
node540.members.0.js
1

3
node541.members.0.js
species
1850093

3
species
node542.members.0.js
2615203

genus
2
1379908

no rank
2
2639626

node545.members.0.js
species
1379909
2

family
89373
32

1
978
genus

985
1
species

1
269798
strain
node549.members.0.js

genus
455076
7

7
2704465
node551.members.0.js
species

3
23
genus
node552.members.0.js
107

108
2
species

node554.members.0.js
strain
504472
2

2621999
3
no rank

2894082
species
node556.members.0.js
1

node557.members.0.js
species
2520506
2

564064
species
node558.members.0.js
15

1
94253
no rank

species
node560.members.0.js
2026729
1

1
1501348
family

273135
1
genus

node563.members.0.js
species
247481
1

37452
3
no rank

1898104
species
node565.members.0.js
2

node566.members.0.js
species
1690483
1

class
200643
126

171549
order
node568.members.0.js
122
3

family
2005473
2

2518495
2
genus

2
2649562
node571.members.0.js
no rank

family
171551
27

genus
836
27

4
3
28123
node574.members.0.js
species

strain
node575.members.0.js
879243
1

2645799
3
no rank

3
species
node577.members.0.js
712435

node578.members.0.js
species
36874
15

species
node579.members.0.js
28124
1

4
node580.members.0.js
species
322095

family
48
171552

47
4
838
genus
node582.members.0.js

2801997
node583.members.0.js
species
1

no rank
84374
3

3
159272
node585.members.0.js
species

1
28133
node586.members.0.js
species

species
1
589437

1
node588.members.0.js
strain
1236518

2
node589.members.0.js
species
839

species
1
28131

node591.members.0.js
strain
1122984
1

2638335
2
no rank

node593.members.0.js
species
2913621
1

2913616
species
node594.members.0.js
1

282402
species
node595.members.0.js
1

28132
node596.members.0.js
species
1

node597.members.0.js
species
28127
7

2
77095
node598.members.0.js
species

28135
node599.members.0.js
species
5

node600.members.0.js
species
1177574
14

1
species
node601.members.0.js
28129

1
node602.members.0.js
species
52227

genus
1
2884814

species
node604.members.0.js
2133944
1

171550
12
family

11
239759
genus

1
626932
species
node607.members.0.js

1
species
node608.members.0.js
2585118

2
species
node609.members.0.js
1288121

214856
species
node610.members.0.js
1

species
node611.members.0.js
328813
1
4

3
2585117
node612.members.0.js
subspecies

2
node613.members.0.js
species
2364787

genus
1
1611681

1433126
node615.members.0.js
species
1

family
2005520
5

307628
1
genus

1642646
species
node618.members.0.js
1

genus
2
294702

species
node620.members.0.js
1642647
2

156973
2
genus

no rank
node622.members.0.js
2630389
2

815
13
family

2
909656
genus

821
node625.members.0.js
species
1

1
node626.members.0.js
species
357276

816
node627.members.0.js
genus
11
5

1
818
node628.members.0.js
species

291644
node629.members.0.js
species
1

1
node630.members.0.js
species
371601

node631.members.0.js
species
47678
3

family
2005525
11

2
375288
node633.members.0.js
genus

195950
9
genus

2
712710
species
node635.members.0.js

7
species
node636.members.0.js
28112

1853231
node637.members.0.js
family
1

4
1970189
order

family
4
1471398

1471399
2
genus

1
2706887
node641.members.0.js
species

node642.members.0.js
species
1168034
1

genus
2
2678352

2
2681766
node644.members.0.js
species

231
117743
class

231
5
200644
order
node646.members.0.js

family
1
246874

253244
1
no rank

1
species
node649.members.0.js
2829795

node650.members.0.js
family
49546
4
144

2
252306
genus

2
252307
species

313596
strain
node653.members.0.js
2

527198
1
genus

no rank
1
2643887

2027857
species
node656.members.0.js
1

1
2045416
genus

no rank
2631961
1

1
2529032
species
node659.members.0.js

genus
363408
4

species
930802
4

node662.members.0.js
strain
1454201
4

252356
genus
node663.members.0.js
8
2

no rank
6
2615042

node665.members.0.js
species
2496865
1

5
313603
species
node666.members.0.js

417127
2
genus

species
1
398743

655815
strain
node669.members.0.js
1

no rank
2632541
1

node671.members.0.js
species
2779359
1

237
node672.members.0.js
genus
50
6

species
node673.members.0.js
2748320
1

node674.members.0.js
species
2816357
1

node675.members.0.js
no rank
196869
4
31

2897181
node676.members.0.js
species
1

14
2830782
node677.members.0.js
species

2893884
species
node678.members.0.js
1

2893883
node679.members.0.js
species
1

node680.members.0.js
species
1179672
1

1751095
species
node681.members.0.js
2

2478552
species
node682.members.0.js
1

1
species
node683.members.0.js
2893886

1
species
node684.members.0.js
2739062

1
species
node685.members.0.js
2724135

3
935222
node686.members.0.js
species

2602769
node687.members.0.js
species
1

1784713
species
node688.members.0.js
2

species
node689.members.0.js
2175091
1

998845
node690.members.0.js
species
1

species
node691.members.0.js
2518177
4

node692.members.0.js
species
1306519
1

1
node693.members.0.js
species
2172098

290174
3
genus

2627091
3
no rank

2
1714860
node696.members.0.js
species

1
2736252
species
node697.members.0.js

389486
2
genus

2648869
2
no rank

2
1453352
species
node700.members.0.js

no rank
5
61432

species
node702.members.0.js
1871037
3

species
node703.members.0.js
1150389
1

531844
node704.members.0.js
species
1

52959
genus
node705.members.0.js
6
1

1
species
node706.members.0.js
996801

no rank
196858
4

1
313598
node708.members.0.js
species

1
node709.members.0.js
species
1336804

2
species
node710.members.0.js
2808900

111500
8
genus

254955
node712.members.0.js
species
2

1
516051
species
node713.members.0.js

1
2615049
no rank

species
node715.members.0.js
2908843
1

1
1383885
node716.members.0.js
species

species
1
111501

886377
node718.members.0.js
strain
1

2698672
species
node719.members.0.js
2

1
336276
genus

2615019
no rank
node721.members.0.js
1

1
83612
genus

no rank
196864
1

node724.members.0.js
species
2777186
1

3
104267
genus

no rank
2635139
2

2
species
node727.members.0.js
754423

node728.members.0.js
species
669041
1

2058174
node729.members.0.js
genus
3
1

2
2585771
node730.members.0.js
species

genus
2
76831

256
node732.members.0.js
species
2

genus
286104
5

2686077
node734.members.0.js
species
3

no rank
2615021
1

species
node736.members.0.js
754409
1

2686078
node737.members.0.js
species
1

762641
1
genus

2507538
species
node739.members.0.js
1

2
261827
genus

species
node741.members.0.js
1736674
1

no rank
1
2615009

1
species
node743.members.0.js
2686365

genus
292691
1

2126553
species
node745.members.0.js
1

15
3
1016
node746.members.0.js
genus

2640652
3
no rank

node748.members.0.js
species
1705617
1

1
1316596
species
node749.members.0.js

species
node750.members.0.js
2748316
1

1
1017
species
node751.members.0.js

species
node752.members.0.js
28188
1

5
species
node753.members.0.js
28189

1019
species
node754.members.0.js
2

genus
1
112040

1
node756.members.0.js
species
63186

genus
221065
3

no rank
2685202
3

3
2282170
species
node759.members.0.js

genus
143222
3

1
2633436
no rank

1
1729720
species
node762.members.0.js

2
270918
species
node763.members.0.js

genus
1
291183

no rank
1
2647285

1
2653681
species
node766.members.0.js

genus
153265
3

2615031
3
no rank

species
node769.members.0.js
2494375
3

genus
1434045
2

no rank
2636168
2

node772.members.0.js
species
2900738
2

genus
1
225842

no rank
1
2644710

1
1336794
node775.members.0.js
species

76
16
2762318
family
node776.members.0.js

genus
308865
2

1
1117645
species
node778.members.0.js

238
node779.members.0.js
species
1

1
node780.members.0.js
genus
59734

1013
2
genus

2
1014
species
node782.members.0.js

1
1778601
genus

1
node784.members.0.js
species
2500547

genus
1
28250

1
28251
node786.members.0.js
species

3
8
genus
node787.members.0.js
501783

1
237258
node788.members.0.js
species

2004710
species
node789.members.0.js
4

2782232
no rank
node790.members.0.js
45
2

genus
2782229
2

2
2487072
species
node792.members.0.js

59732
genus
node793.members.0.js
40
5

651561
node794.members.0.js
species
1

1
558152
species
node795.members.0.js

254
node796.members.0.js
species
1

1
12
node797.members.0.js
no rank
2593645

2039166
node798.members.0.js
species
6

1
2724619
node799.members.0.js
species

2
species
node800.members.0.js
2879938

species
node801.members.0.js
2487065
1

2015076
species
node802.members.0.js
1

250
species
node803.members.0.js
2

node804.members.0.js
species
253
13

1
536441
node805.members.0.js
species

1265445
species
node806.members.0.js
1

1
112234
node807.members.0.js
species

2
node808.members.0.js
species
1685010

2782231
1
genus

1
node810.members.0.js
species
1241979

family
3
1853230

genus
332102
1

species
node813.members.0.js
1798018
1

genus
246873
2

no rank
2
2619166

2
2862509
node816.members.0.js
species

family
39782
1

34098
1
genus

1
node819.members.0.js
species
1653831

1333713
1
family

1
2828338
genus

1
node822.members.0.js
species
2761580

class
70
117747

70
200666
order

node825.members.0.js
family
84566
6
70

genus
node826.members.0.js
28453
3
17

7
2609468
no rank

node828.members.0.js
species
2557994
1

1
1538644
species
node829.members.0.js

species
node830.members.0.js
2907623
4

1
2713573
species
node831.members.0.js

node832.members.0.js
species
561061
1

species
node833.members.0.js
371142
1

2886510
node834.members.0.js
species
2

species
node835.members.0.js
258
1

node836.members.0.js
species
649196
1

259
species
node837.members.0.js
1

1
376469
genus

no rank
2632301
1

node840.members.0.js
species
2592345
1

genus
1
929509

species
1
995

strain
node843.members.0.js
929556
1

genus
node844.members.0.js
423349
1
11

species
node845.members.0.js
551996
5

2
node846.members.0.js
species
398053

no rank
2617802
1

1
2728022
species
node848.members.0.js

398049
species
node849.members.0.js
1

1550579
species
node850.members.0.js
1

84567
node851.members.0.js
genus
34
4

12
1
2628915
no rank
node852.members.0.js

4
species
node853.members.0.js
2762321

1
node854.members.0.js
species
2856523

1
species
node855.members.0.js
2578106

5
2482728
species
node856.members.0.js

363852
node857.members.0.js
species
1

984
species
node858.members.0.js
5

332999
node859.members.0.js
species
1

species
node860.members.0.js
2605747
1

node861.members.0.js
species
2201271
6

node862.members.0.js
species
430522
4

phylum
1379697
1

no rank
1
2751484

1
2053527
species
node865.members.0.js

48479
no rank
node866.members.0.js
475
27

node867.members.0.js
species
698386
1

1
species
node868.members.0.js
698384

1
node869.members.0.js
species
1181499

1
164851
species
node870.members.0.js

77133
species
node871.members.0.js
405

node872.members.0.js
species
1191434
1

species
node873.members.0.js
1781347
1

node874.members.0.js
species
698387
3

698379
node875.members.0.js
species
1

1
1130999
node876.members.0.js
species

1
698391
species
node877.members.0.js

1
node878.members.0.js
species
1425034

1
1256537
species
node879.members.0.js

698395
species
node880.members.0.js
5

1425163
species
node881.members.0.js
4

2
node882.members.0.js
species
155098

1242644
species
node883.members.0.js
2

node884.members.0.js
species
542421
1

5
node885.members.0.js
species
526686

1562172
node886.members.0.js
species
1

3
node887.members.0.js
species
526666

526687
node888.members.0.js
species
1

2
node889.members.0.js
species
698394

1
1701325
species
node890.members.0.js

2
species
node891.members.0.js
698390

phylum
40117
11

class
203693
11

order
11
189778

family
11
189779

node896.members.0.js
genus
179
1
6

species group
655606
5

180
node898.members.0.js
species
5

2
1234
genus

1
node900.members.0.js
species
330214

1715989
species
node901.members.0.js
1

no rank
3
688163

3
node903.members.0.js
species
2026770

35
2323
no rank

clade
35
1783234

clade
1783273
11

1794811
4
clade

phylum
3
1752727

2
2026716
node909.members.0.js
species

species
node910.members.0.js
1618633
1

1817917
1
phylum

1
2750080
species
node912.members.0.js

clade
7
1794810

phylum
1752726
1

1618372
species
node915.members.0.js
1

phylum
1752722
3

2026804
species
node917.members.0.js
3

phylum
1752723
3

3
species
node919.members.0.js
2282149

95901
3
clade

2
1104668
no rank

2
2891165
species
node922.members.0.js

class
2497643
1

1
2497644
order

family
2497645
1

genus
1551504
1

1
673862
node927.members.0.js
species

67817
2
phylum

no rank
910038
1

1
2052149
species
node930.members.0.js

1
99276
no rank

174294
node932.members.0.js
species
1

3
1619053
phylum

class
1801911
3

1803398
3
order

3
1803397
clade

genus
1803399
3

3
node938.members.0.js
species
1735162

node939.members.0.js
phylum
95818
3
14

2171982
2
class

2
2171984
order

2171990
2
family

2
2725941
no rank

2
species
node944.members.0.js
2725942

class
2093818
4

order
4
2093819

family
4
2093822

2093823
4
genus

4
2725944
no rank

4
species
node950.members.0.js
2725945

2080739
node951.members.0.js
species
1

no rank
1895827
4

2026720
node953.members.0.js
species
3

1
node954.members.0.js
species
2572087

1
221235
phylum

1
221236
no rank

1
221237
species
node957.members.0.js

phylum
1
1052815

genus
1
1462351

2807604
1
no rank

node961.members.0.js
species
2807605
1

1930617
1
phylum

1
1962850
class

1
1962852
order

family
1962854
1

genus
187144
1

187145
1
species

880073
node968.members.0.js
strain
1

phylum
6
200918

6
188708
class

4
2419
order

family
2
188709

2335
genus
node973.members.0.js
2

family
1643950
2

2
2420
genus

1
46541
species
node976.members.0.js

1
species
node977.members.0.js
2571116

1643947
2
order

family
1643949
2

2
node980.members.0.js
genus
160798

200783
4
phylum

no rank
2202150
3

species
node983.members.0.js
2202151
3

187857
1
class

1
32069
order

node986.members.0.js
family
64898
1

1
200938
phylum

class
118001
1

189769
1
order

family
1
189770

genus
393029
1

936456
species
node992.members.0.js
1

74152
1
phylum

class
641853
1

1
641854
order

641876
1
family

genus
423604
1

423605
1
species

1
445932
node999.members.0.js
strain

1
66
node1000.members.0.js
phylum
57723

1813735
6
class

2910145
6
order

2211325
6
family

2004797
6
genus

6
species
node1005.members.0.js
1855912

305072
4
no rank

4
1978231
node1007.members.0.js
species

class
16
204432

order
10
204433

family
10
204434

genus
3
33973

species
3
33075

3
240015
node1013.members.0.js
strain

genus
388463
3

no rank
3
2637509

1
2763071
node1016.members.0.js
species

2
2703788
node1017.members.0.js
species

genus
4
940557

species
940614
1

node1020.members.0.js
strain
682795
1

no rank
3
2621151

node1022.members.0.js
species
2602070
3

332160
6
order

family
1962910
3

1649475
3
genus

species
node1026.members.0.js
1473598
3

no rank
1470392
3

3
species
node1028.members.0.js
2802972

no rank
57727
32

171953
node1030.members.0.js
species
32

533205
4
class

533206
2
order

family
533207
2

genus
2915177
2

2
species
node1035.members.0.js
2818388

2
574975
order

no rank
2
1775601

node1038.members.0.js
species
2478486
2

3
1562566
class

3
2
458032
genus
node1040.members.0.js

2851959
species
node1041.members.0.js
1

clade
352
1783257

phylum
67812
2

no rank
1047005
2

2
2035772
species
node1045.members.0.js

node1046.members.0.js
phylum
203682
4
242

class
node1047.members.0.js
203683
2
162

order
19
2691356

family
node1049.members.0.js
1763524
1
19

1763521
8
genus

8
1387353
species
node1051.members.0.js

127
2
genus

node1053.members.0.js
species
128
1
2

1
575540
node1054.members.0.js
strain

2680020
4
genus

4
node1056.members.0.js
species
2527974

genus
2
466152

466153
node1058.members.0.js
species
2

1511635
2
genus

2
species
node1060.members.0.js
406548

2
573133
no rank

2
node1062.members.0.js
species
2052181

order
47
112

no rank
40903
6

6
species
node1065.members.0.js
120965

466154
5
no rank

species
node1067.members.0.js
2603660
4

1
2723666
species
node1068.members.0.js

36
2
126
family
node1069.members.0.js

genus
3
2795776

species
node1071.members.0.js
2528027
3

genus
2795777
2

node1073.members.0.js
species
2528026
2

2795778
5
genus

5
species
node1075.members.0.js
2527983

1
2714737
genus

species
node1077.members.0.js
2527989
1

1
10
node1078.members.0.js
genus
1649453

5
node1079.members.0.js
species
2527971

1
122
node1080.members.0.js
species

2
2527978
species
node1081.members.0.js

1
2527964
species
node1082.members.0.js

no rank
8
69476

1
149589
species
node1084.members.0.js

1
node1085.members.0.js
species
149591

node1086.members.0.js
species
149590
3

species
node1087.members.0.js
69477
2

1
2026779
species
node1088.members.0.js

1
1649480
genus

node1090.members.0.js
species
2528019
1

genus
2
2795780

2
2527995
node1092.members.0.js
species

2
1936111
genus

2
1891926
species
node1094.members.0.js

order
node1095.members.0.js
2691354
1
53

2691357
node1096.members.0.js
family
34
2

1
2795605
genus

node1098.members.0.js
species
2527968
1

1579506
2
genus

980254
node1100.members.0.js
species
2

genus
2795775
1

node1102.members.0.js
species
2528010
1

genus
2807414
11

11
2528021
species
node1104.members.0.js

genus
5
123

5
125
node1106.members.0.js
species

1
2714594
genus

2527984
species
node1108.members.0.js
1

1579505
2
genus

1930273
species
node1110.members.0.js
2

265488
2
genus

species
2
265606

2
243090
strain
node1113.members.0.js

6
2795973
genus

2527979
species
node1115.members.0.js
6

node1116.members.0.js
genus
2795779
1

family
2691359
16

5
2795781
genus

5
node1119.members.0.js
species
2527972

1
9
genus
node1120.members.0.js
2691417

2650471
node1121.members.0.js
species
7

1
node1122.members.0.js
species
2528024

genus
2862451
1

1
1930276
node1124.members.0.js
species

genus
1400386
1

1
1400387
node1126.members.0.js
species

2
2691358
family

genus
2
1676125

species
node1129.members.0.js
1331910
2

39
2691355
order

1914233
node1131.members.0.js
family
39
3

no rank
2052163
2

2052164
species
node1133.members.0.js
2

genus
1382832
6

6
1123043
species
node1135.members.0.js

5
2731450
genus

species
node1137.members.0.js
2598579
5

10
2051044
genus

node1139.members.0.js
species
692036
10

5
2807415
genus

node1141.members.0.js
species
2528023
5

1
4
genus
node1142.members.0.js
113

1210884
node1143.members.0.js
species
1

2
species
node1144.members.0.js
114

genus
4
2774146

4
species
node1146.members.0.js
2774151

class
2517206
3

1127829
3
order

3
1127830
family

genus
380738
3

174633
species
node1151.members.0.js
3

class
62
666505

node1153.members.0.js
order
2483366
1
6

family
3
2483367

2483368
2
genus

2
1941349
species
node1156.members.0.js

genus
1
2690160

node1158.members.0.js
species
1851148
1

family
2
2690158

2690159
2
genus

1936003
species
node1161.members.0.js
2

order
5
666506

1500946
3
no rank

3
2052180
node1164.members.0.js
species

family
2
666507

2803301
1
genus

1
node1167.members.0.js
species
2528020

no rank
1
1385974

2026777
species
node1169.members.0.js
1

order
1771349
51

1771355
51
family

51
2807511
genus

species
node1173.members.0.js
2807512
51

473814
no rank
node1174.members.0.js
9
1

1930275
node1175.members.0.js
species
1

3
2026780
species
node1176.members.0.js

species
node1177.members.0.js
2527975
1

2
2527980
node1178.members.0.js
species

species
node1179.members.0.js
2528008
1

class
2
2897345

2897347
2
order

2897348
2
family

2897349
2
genus

2596890
node1184.members.0.js
species
2

phylum
91
74201

203494
50
class

50
48461
order

family
20
203557

2
1348508
genus

no rank
2711230
2

2
2711231
node1191.members.0.js
species

genus
16
518753

7
2824561
node1193.members.0.js
species

2728835
species
node1194.members.0.js
9

2911469
1
genus

2707525
node1196.members.0.js
species
1

genus
2735
1

species
1
2736

1
240016
strain
node1199.members.0.js

family
16
1647988

2
239934
genus

node1202.members.0.js
species
239935
2

1951308
14
no rank

14
species
node1204.members.0.js
2562705

14
134621
family

genus
14
1032526

14
1032527
species

320771
strain
node1208.members.0.js
14

no rank
1
326457

genus
1
1541670

no rank
2730358
1

1
node1212.members.0.js
species
2730359

3
134549
class

order
3
1836787

3
1836792
family

genus
3
295577

species
191863
3

strain
node1218.members.0.js
497964
3

class
1955630
1

order
717963
1

717964
1
family

1
511745
genus

species
1
431057

1
1202785
node1224.members.0.js
strain

1
417295
no rank

1
node1226.members.0.js
species
2488809

class
414999
17

2
415001
order

415002
2
family

2480624
1
genus

node1231.members.0.js
species
2200854
1

genus
2175957
1

no rank
1
2621336

2866311
node1234.members.0.js
species
1

order
415000
15

family
134623
15

genus
2100741
2

2
2628636
no rank

2
node1239.members.0.js
species
2862869

genus
2028344
1

1
1796921
species
node1241.members.0.js

genus
1961799
1

species
node1243.members.0.js
1838286
1

278955
2
no rank

2
794903
node1245.members.0.js
species

178440
node1246.members.0.js
genus
6
1

2649488
4
no rank

1882749
species
node1248.members.0.js
1

3
134634
node1249.members.0.js
species

107709
node1250.members.0.js
species
1

genus
2576890
3

3
node1252.members.0.js
species
2576891

74202
18
no rank

18
156588
species
node1254.members.0.js

phylum
256845
2

no rank
641407
1

1
2886196
node1257.members.0.js
species

class
1
1313211

order
1
278082

1
1674876
no rank

1
node1261.members.0.js
species
2094242

204428
15
phylum

class
204429
15

1963360
order
node1264.members.0.js
11
1

family
92713
9

genus
7
282132

species
node1267.members.0.js
389348
7

112987
2
genus

no rank
2643326
2

1353976
node1270.members.0.js
species
2

1
92712
family

1
420389
no rank

species
node1273.members.0.js
2698248
1

4
51291
order

809
2
family

1
1113537
no rank

810
1
genus

1
species
node1278.members.0.js
813

2822114
1
genus

1
2681469
node1280.members.0.js
species

no rank
95916
2

2
node1282.members.0.js
species
174324

clade
1802340
1

1293497
1
phylum

class
1
1293498

order
1293499
1

407032
1
family

genus
2806694
1

species
node1289.members.0.js
2705533
1

91
2981
phylum
node1290.members.0.js
1224

subphylum
68525
156

35
29547
class

order
3
235899

3
224467
family

genus
3
191301

species
291048
2

2
391592
node1297.members.0.js
strain

1
node1298.members.0.js
species
1424653

2
87828
no rank

2
120858
species
node1300.members.0.js

order
30
213849

family
12
72294

genus
194
11

node1304.members.0.js
species
28080
2

1
197
species
node1305.members.0.js

2
species
node1306.members.0.js
824

1
1965231
species

1
node1308.members.0.js
subspecies
1660067

199
node1309.members.0.js
species
1

species
201
2

2
node1311.members.0.js
subspecies
488545

206
species
node1312.members.0.js
1

node1313.members.0.js
species
260714
1

genus
57665
1

65553
1
species

1
node1316.members.0.js
strain
525898

family
5
72293

genus
209
5

123841
node1319.members.0.js
species
1

210
1
species

strain
node1321.members.0.js
1055530
1

species
node1322.members.0.js
213
3

2
8
node1323.members.0.js
family
2808963

genus
2321115
1

node1325.members.0.js
species
1462615
1

1
genus
node1326.members.0.js
28196

3
2321111
genus

1
node1328.members.0.js
species
1935204

1
1564138
node1329.members.0.js
species

node1330.members.0.js
species
28198
1

2321187
1
genus

1904463
species
node1332.members.0.js
1

2
2771472
family

genus
1
269260

1
269261
species

1
749222
strain
node1336.members.0.js

genus
1
265570

2779528
species
node1338.members.0.js
1

family
3
2771471

3
202746
genus

node1341.members.0.js
species
2590022
1

1
2509341
node1342.members.0.js
species

no rank
1
2623549

1
2894755
species
node1344.members.0.js

2
121
node1345.members.0.js
class
28221

8
1
213115
order
node1346.members.0.js

194924
7
family

genus
872
3

no rank
2593640
1

node1350.members.0.js
species
2875247
1

241368
node1351.members.0.js
species
1

node1352.members.0.js
species
44742
1

2035811
node1353.members.0.js
genus
3
1

879567
1
species

1
strain
node1355.members.0.js
1322246

1716143
node1356.members.0.js
species
1

1
2910984
genus

species
1
184917

1
573370
strain
node1359.members.0.js

order
1
453227

453228
1
family

genus
1
453229

node1363.members.0.js
species
453230
1

no rank
5
122706

species
node1365.members.0.js
2026735
5

1779134
2
order

family
1779135
1

genus
2766927
1

node1369.members.0.js
species
2292766
1

no rank
2099666
1

species
node1371.members.0.js
2600177
1

order
8
69541

213421
1
family

genus
1
890

2802975
node1375.members.0.js
species
1

5
213422
family

1
28231
genus

node1378.members.0.js
species
35554
1

1
115782
genus

313985
1
species

398767
strain
node1381.members.0.js
1

genus
3
2651583

2
2847991
species
node1383.members.0.js

1
2847989
node1384.members.0.js
species

family
2
2812024

2812025
2
genus

1
19
species

1
338963
strain
node1388.members.0.js

1
1842532
node1389.members.0.js
species

2914038
1
order

family
213468
1

1
2357
genus

1
2358
species

1
strain
node1394.members.0.js
706587

no rank
1
45456

genus
1769732
1

1
species
node1397.members.0.js
1750598

1
node1398.members.0.js
no rank
34033

10
213118
order

2886822
2
family

genus
2
2886823

2
2569540
node1402.members.0.js
species

family
2
213121

427922
1
genus

1
427923
species

1
strain
node1406.members.0.js
589865

893
1
genus

species
node1408.members.0.js
1986146
1

family
213119
6

1
2904687
genus

species
2296
1

1
node1412.members.0.js
strain
177437

231684
2
no rank

node1414.members.0.js
species
2049433
2

1
896
genus

1
897
species
node1416.members.0.js

2
497721
genus

no rank
2
2677172

2810562
species
node1419.members.0.js
2

order
29
82

3
224462
suborder

family
3
224464

1
162027
genus

species
80816
1

502025
node1425.members.0.js
strain
1

no rank
521188
2

521189
species
node1427.members.0.js
2

49
7
80811
node1428.members.0.js
suborder

family
12
1524215

node1430.members.0.js
genus
161492
2
12

2620896
10
no rank

404589
node1432.members.0.js
species
10

1524213
2
family

1524214
2
genus

node1435.members.0.js
species
1391653
2

family
15
39

genus
47
5

4
node1438.members.0.js
species
83451

1
48
node1439.members.0.js
species

2
44
genus

2
83453
species

2
node1442.members.0.js
strain
1294270

40
5
genus

1
5
node1444.members.0.js
species
41

3
378806
node1445.members.0.js
strain

node1446.members.0.js
strain
675526
1

genus
3
42

3
43
node1448.members.0.js
species

2
13
family
node1449.members.0.js
31

83461
genus
node1450.members.0.js
2

32
node1451.members.0.js
genus
8
1

5
83455
species

5
1278073
node1453.members.0.js
strain

node1454.members.0.js
species
34
2

1
224458
genus

2629105
1
no rank

2813578
node1457.members.0.js
species
1

suborder
80812
30

1055686
4
family

genus
1055688
4

species
node1461.members.0.js
927083
4

49
14
family

genus
13
39643

node1464.members.0.js
species
56
10
13

node1465.members.0.js
strain
1254432
2

1
448385
strain
node1466.members.0.js

genus
50
1

species
node1468.members.0.js
52
1

family
1524216
11

genus
1524217
11

1391654
species
node1471.members.0.js
11

no rank
1
215910

genus
1649470
1

1
species
node1474.members.0.js
888845

1267
44
1236
class
node1475.members.0.js

3
105
node1476.members.0.js
order
135622

1
267891
family

58050
1
genus

69539
species
node1479.members.0.js
1

family
49
267888

genus
1
907197

1
1266052
node1482.members.0.js
species

genus
node1483.members.0.js
53246
3
48

1
1348114
species
node1484.members.0.js

no rank
194690
1

2785910
node1486.members.0.js
species
1

1
267375
node1487.members.0.js
species

node1488.members.0.js
species
43658
1

4
node1489.members.0.js
species
43657

node1490.members.0.js
species
161398
31

node1491.members.0.js
species
882626
3

28107
1
species

1314869
node1493.members.0.js
strain
1

1
227
species
node1494.members.0.js

node1495.members.0.js
species
43662
1

family
20
267890

7
20
genus
node1497.members.0.js
22

node1498.members.0.js
species
1028752
1

node1499.members.0.js
species
2590884
1

1
node1500.members.0.js
species
256839

1
192073
species

318161
strain
node1502.members.0.js
1

species
node1503.members.0.js
1738770
1

node1504.members.0.js
species
24
1

node1505.members.0.js
species
2593655
1

271097
4
species

4
strain
node1507.members.0.js
425104

1
species
node1508.members.0.js
2487742

no rank
1
196818

species
node1510.members.0.js
2864207
1

family
267893
7

7
135575
genus

2
species
node1513.members.0.js
86102

2614829
5
no rank

4
2055892
species
node1515.members.0.js

node1516.members.0.js
species
1874361
1

267889
8
family

genus
4
28228

196834
4
no rank

node1520.members.0.js
species
2689569
1

1816218
node1521.members.0.js
species
1

2497879
node1522.members.0.js
species
1

2161872
species
node1523.members.0.js
1

2848171
1
genus

species
node1525.members.0.js
1967665
1

genus
1518149
3

2
1763536
species
node1527.members.0.js

no rank
1
2614972

2769490
species
node1529.members.0.js
1

family
267894
1

67572
1
genus

357794
1
species

1
357804
node1533.members.0.js
strain

family
72275
15

genus
1621534
1

1
197222
species
node1536.members.0.js

genus
2
1751872

node1538.members.0.js
species
1526571
2

2894574
1
genus

node1540.members.0.js
species
2172099
1

11
2903219
no rank

11
2
226
genus
node1542.members.0.js

node1543.members.0.js
no rank
2614992
2
7

2652380
node1544.members.0.js
species
2

2
node1545.members.0.js
species
1917157

species
node1546.members.0.js
2267264
1

1
species
node1547.members.0.js
28108

314275
node1548.members.0.js
species
1

267892
1
family

1
44011
genus

44012
1
species

1
550540
strain
node1552.members.0.js

2
2887327
order

family
2
1920240

2
261963
genus

species
261964
1

1
strain
node1557.members.0.js
523791

1
node1558.members.0.js
species
914150

order
22
135613

family
1046
10

genus
1227
2

species
1229
1

node1563.members.0.js
strain
323261
1

1
133539
species

1
node1565.members.0.js
strain
472759

1
85076
genus

species
1
37487

1
765910
node1568.members.0.js
strain

53392
1
genus

1
node1570.members.0.js
species
1166950

no rank
3
82569

species
node1572.members.0.js
1978339
3

2
85073
genus

1050
2
species

316276
strain
node1575.members.0.js
2

1
67575
genus

species
node1577.members.0.js
2498451
1

72276
5
family

genus
1
85108

1
1052
species
node1580.members.0.js

1
1051
genus

no rank
2684909
1

species
node1583.members.0.js
1442136
1

187271
1
no rank

2740164
species
node1585.members.0.js
1

genus
106633
1

106634
species
node1587.members.0.js
1

genus
1
1765964

1765967
species
node1589.members.0.js
1

449719
1
family

genus
1
437504

species
1
437505

1
node1593.members.0.js
strain
1192854

1
1096778
family

1
2034504
genus

1972068
node1596.members.0.js
species
1

451214
5
no rank

genus
node1598.members.0.js
1273155
5

10
244
order
node1599.members.0.js
91347

1
2812006
family

genus
1
84565

2636512
1
no rank

2697027
species
node1603.members.0.js
1

family
1903411
35

3
1565532
genus

3
1646377
species
node1606.members.0.js

genus
1
1745211

1639108
1
species

1
1441930
strain
node1609.members.0.js

613
node1610.members.0.js
genus
12
6

node1611.members.0.js
species
615
1

2
species
node1612.members.0.js
618

node1613.members.0.js
species
82996
2

47917
species
node1614.members.0.js
1

node1615.members.0.js
genus
34037
3
4

1
58169
species
node1616.members.0.js

1964366
1
genus

no rank
1
2636213

1
node1619.members.0.js
species
2126321

14
2
629
node1620.members.0.js
genus

2
28152
node1621.members.0.js
species

species group
1649845
5

5
367190
species
node1623.members.0.js

species
node1624.members.0.js
2607663
3

species
33060
1

1
349967
node1626.members.0.js
strain

1
node1627.members.0.js
species
29484

node1628.members.0.js
family
1903409
6
47

genus
5
32199

9
node1630.members.0.js
species
5
4

1241834
forma specialis
node1631.members.0.js
1

551
10
genus

3
2622719
no rank

2675378
species
node1634.members.0.js
3

species
node1635.members.0.js
65700
2

5
1922217
species
node1636.members.0.js

53335
node1637.members.0.js
genus
26
5

species group
4
1654067

4
species
node1639.members.0.js
549

3
553
node1640.members.0.js
species

2630326
4
no rank

2490851
node1642.members.0.js
species
1

1
species
node1643.members.0.js
553117

1
node1644.members.0.js
species
592316

2052056
node1645.members.0.js
species
1

59814
species
node1646.members.0.js
9

node1647.members.0.js
species
66269
1

family
10
1903410

71655
3
genus

1
2722756
species
node1650.members.0.js

1
species
node1651.members.0.js
1109412

1
55210
species

1121120
strain
node1653.members.0.js
1

2884243
1
genus

1
69223
species

1
579405
strain
node1656.members.0.js

2
6
node1657.members.0.js
genus
122277

species
node1658.members.0.js
2108399
1

2204145
node1659.members.0.js
species
2

node1660.members.0.js
species
1905730
1

node1661.members.0.js
family
543
13
127

158483
node1662.members.0.js
genus
2
1

1
species
node1663.members.0.js
158822

genus
5
1330547

1
node1665.members.0.js
species
497725

1
283686
node1666.members.0.js
species

1
species
node1667.members.0.js
1646340

2632876
1
no rank

1
2725560
node1669.members.0.js
species

species
node1670.members.0.js
1158459
1

genus
node1671.members.0.js
544
4
11

4
node1672.members.0.js
species
67824

1
1344959
species group

species
1
546

1
1333848
strain
node1675.members.0.js

1
node1676.members.0.js
no rank
2644389

67825
node1677.members.0.js
species
1

genus
590
22

28901
species
node1679.members.0.js
22
12

node1680.members.0.js
subspecies
59201
2

node1681.members.0.js
subspecies
59202
4

subspecies
node1682.members.0.js
59203
1
4

no rank
node1683.members.0.js
1151166
3

2890311
26
no rank

570
genus
node1685.members.0.js
22
7

2608929
1
no rank

1
1972757
node1687.members.0.js
species

1463165
node1688.members.0.js
species
2

548
species
node1689.members.0.js
7

1
species
node1690.members.0.js
573

3
species
node1691.members.0.js
571

244366
species
node1692.members.0.js
1

160674
node1693.members.0.js
genus
4
2

575
node1694.members.0.js
species
2

genus
2
1330545

2
node1696.members.0.js
species
61646

genus
83654
3

no rank
node1698.members.0.js
2627398
1
3

2681307
species
node1699.members.0.js
1

1
2815358
species
node1700.members.0.js

11
561
genus

562
node1702.members.0.js
species
9

1
2608889
no rank

1
node1704.members.0.js
species
2044467

208962
node1705.members.0.js
species
1

no rank
191675
2

clade
84563
2

clade
146507
2

genus
1
568988

node1710.members.0.js
species
138073
1

134287
species
node1711.members.0.js
1

node1712.members.0.js
genus
413496
2
3

413503
species
node1713.members.0.js
1

genus
2
2055880

2
species
node1715.members.0.js
566

547
node1716.members.0.js
genus
25
5

10
6
354276
species group
node1717.members.0.js

3
node1718.members.0.js
species
69218

1
node1719.members.0.js
species
550

10
2608935
no rank

node1721.members.0.js
species
2742639
4

node1722.members.0.js
species
1914861
1

1
species
node1723.members.0.js
2870346

1692238
species
node1724.members.0.js
3

399742
node1725.members.0.js
species
1

family
14
1903414

637
node1727.members.0.js
genus
1

genus
node1728.members.0.js
583
2

node1729.members.0.js
genus
586
2
10

587
species
node1730.members.0.js
2

no rank
2
2633465

2
2828763
species
node1732.members.0.js

species
node1733.members.0.js
333965
4

626
1
genus

node1735.members.0.js
species
351671
1

2
265
node1736.members.0.js
order
72274

2887365
6
family

6
2742
genus

1
node1739.members.0.js
species
1420917

species
node1740.members.0.js
1420916
2

no rank
83889
3

1415568
node1742.members.0.js
species
1

1
2488665
node1743.members.0.js
species

2304594
node1744.members.0.js
species
1

no rank
97500
2

2
332614
node1746.members.0.js
species

family
node1747.members.0.js
135621
5
255

genus
1
1654787

1
species
node1749.members.0.js
1697053

genus
1649479
1

1
node1751.members.0.js
species
1510150

subfamily
3
351

352
3
genus

3
species
node1754.members.0.js
354

286
genus
node1755.members.0.js
240
69

node1756.members.0.js
species
237610
3

2
2745495
species
node1757.members.0.js

104087
species
node1758.members.0.js
3

species group
136846
6

node1760.members.0.js
species subgroup
578833
1
6

node1761.members.0.js
species
316
2
5

1
node1762.members.0.js
strain
1196835

644801
strain
node1763.members.0.js
2

2745503
node1764.members.0.js
species
2

species
node1765.members.0.js
2745509
1

1
node1766.members.0.js
species
289370

4
18
species group
node1767.members.0.js
136849

1
53409
node1768.members.0.js
species

251698
node1769.members.0.js
species subgroup
3
1

species
47877
2

53707
no rank
node1771.members.0.js
2
1

629260
node1772.members.0.js
strain
1

251695
5
species subgroup

species
node1774.members.0.js
317
3
5

321
1
no rank

1
1324931
node1776.members.0.js
strain

node1777.members.0.js
strain
1357279
1

4
species
node1778.members.0.js
33069

node1779.members.0.js
species
36746
1

157782
node1780.members.0.js
species
1

2056231
species
node1781.members.0.js
1

2
species
node1782.members.0.js
2745518

1
1931241
species
node1783.members.0.js

1
node1784.members.0.js
species
1148509

1
198620
node1785.members.0.js
species

species
node1786.members.0.js
1785145
1

node1787.members.0.js
species
216142
1

species
node1788.members.0.js
319939
1

3
198618
node1789.members.0.js
species

node1790.members.0.js
species group
136843
1
26

node1791.members.0.js
species
200450
3

1
species
node1792.members.0.js
47878

1
76761
species

1
node1794.members.0.js
strain
1295141

1
species
node1795.members.0.js
76758

species
node1796.members.0.js
129817
6

node1797.members.0.js
species
46679
4

1
47883
node1798.members.0.js
species

4
species
node1799.members.0.js
294

380021
species
node1800.members.0.js
2
1

1
1420599
node1801.members.0.js
isolate

2
species
node1802.members.0.js
75588

95300
species
node1803.members.0.js
4

3
16
species group
node1804.members.0.js
136845

76759
species
node1805.members.0.js
2

1
species
node1806.members.0.js
2217867

10
6
303
node1807.members.0.js
species

1384061
node1808.members.0.js
strain
4

4
1461581
node1809.members.0.js
species

2774461
species
node1810.members.0.js
1

1
395598
species
node1811.members.0.js

1
101564
species

1
node1813.members.0.js
strain
741155

2894079
node1814.members.0.js
species
1

2
62104
no rank

2
species
node1816.members.0.js
114707

1
species
node1817.members.0.js
65741

species
node1818.members.0.js
1691904
2

136842
7
species group

587753
node1820.members.0.js
species
7
3

3
86192
subspecies
node1821.members.0.js

1
587851
node1822.members.0.js
subspecies

2725477
node1823.members.0.js
species
1

364197
species
node1824.members.0.js
1

136841
22
species group

species subgroup
1232139
1

node1827.members.0.js
species
301
1

species subgroup
627141
2

46680
species
node1829.members.0.js
2

species
53412
1

strain
node1831.members.0.js
1245471
1

14
13
287
species
node1832.members.0.js

381754
node1833.members.0.js
strain
1

4
43263
node1834.members.0.js
species

31
4
196821
no rank
node1835.members.0.js

2874628
species
node1836.members.0.js
1

1
2662033
node1837.members.0.js
species

2653853
species
node1838.members.0.js
3

3
species
node1839.members.0.js
2706126

1
node1840.members.0.js
species
118613

2804761
node1841.members.0.js
species
1

node1842.members.0.js
species
658641
1

3
node1843.members.0.js
species
2726989

1
node1844.members.0.js
species
1573719

1
node1845.members.0.js
species
2735906

2054919
species
node1846.members.0.js
1

1
2498848
species
node1847.members.0.js

1
species
node1848.members.0.js
2654238

2678259
species
node1849.members.0.js
2

1
2745519
species
node1850.members.0.js

1
node1851.members.0.js
species
2730847

1
1881017
species
node1852.members.0.js

2866808
node1853.members.0.js
species
1

1
node1854.members.0.js
species
2730848

1
2678260
species
node1855.members.0.js

node1856.members.0.js
species
2320867
1

genus
5
2758906

5
2213226
species
node1858.members.0.js

no rank
33811
4

947516
species
node1860.members.0.js
1

3
1913989
species
node1861.members.0.js

order
61
135623

family
641
61

4
657
genus

74109
2
species

298386
node1866.members.0.js
strain
2

38293
1
species

1
subspecies
node1868.members.0.js
85581

1
1295392
species

strain
node1870.members.0.js
658445
1

662
node1871.members.0.js
genus
55
10

1
species
node1872.members.0.js
170679

1381081
species
node1873.members.0.js
1

species group
717610
14

2
species
node1875.members.0.js
669

3
680
species
node1876.members.0.js

670
node1877.members.0.js
species
2

3
663
node1878.members.0.js
species

512649
species
node1879.members.0.js
4

1
species
node1880.members.0.js
689

1
673372
species
node1881.members.0.js

1
1435069
species
node1882.members.0.js

1
species
node1883.members.0.js
28172

1
node1884.members.0.js
species
676

1
node1885.members.0.js
species
666

2
2587862
node1886.members.0.js
species

1
55601
node1887.members.0.js
species

species
node1888.members.0.js
2662262
1

1
node1889.members.0.js
species
2711221

1
species
node1890.members.0.js
29497

1
node1891.members.0.js
species
672

1
species
node1892.members.0.js
170651

3
node1893.members.0.js
species
29495

2
1074311
node1894.members.0.js
species

no rank
2614977
10

2785746
species
node1896.members.0.js
8

1
species
node1897.members.0.js
2589990

1
2714948
node1898.members.0.js
species

51366
1
genus

1
1908198
node1900.members.0.js
species

511678
1
genus

node1902.members.0.js
species
40269
1

order
29
118969

118968
3
family

1
59195
genus

676208
species
node1906.members.0.js
1

genus
2
776

2
2676648
no rank

2749996
node1909.members.0.js
species
2

444
26
family

445
node1911.members.0.js
genus
26
2

45067
node1912.members.0.js
species
18

2
node1913.members.0.js
species
2708020

446
node1914.members.0.js
species
1

1867846
node1915.members.0.js
species
2

node1916.members.0.js
species
28082
1

15
1706369
order

family
5
1706373

genus
5
48073

1
435905
node1920.members.0.js
species

2
node1921.members.0.js
species
86173

node1922.members.0.js
species
1769779
2

1706371
10
family

genus
1
2425

node1925.members.0.js
species
2731755
1

1
2036021
genus

node1927.members.0.js
species
1737490
1

genus
447467
1

species
447471
1

1117647
strain
node1930.members.0.js
1

genus
7
10

1
7
node1932.members.0.js
no rank
2624793

species
node1933.members.0.js
1945512
2

node1934.members.0.js
species
2303332
4

order
23
72273

135616
node1936.members.0.js
family
11
1

2781121
2
genus

2
node1938.members.0.js
species
2675054

genus
933
2

2
92245
species

strain
node1941.members.0.js
717772
2

3
28884
genus

species
node1943.members.0.js
28885
3

genus
2781120
1

2675053
node1945.members.0.js
species
1

2
2039723
genus

species
node1947.members.0.js
2739063
2

family
135617
10

4
1021
genus

4
288004
species
node1950.members.0.js

1030
6
genus

2823902
species
node1952.members.0.js
2

no rank
3
2636184

3
node1954.members.0.js
species
96472

species
node1955.members.0.js
2735563
1

family
2
34064

2
262
genus

2
263
species

264
subspecies
node1959.members.0.js
2
1

strain
node1960.members.0.js
984129
1

order
742030
1

family
742031
1

1
180541
genus

no rank
2649847
1

2183911
node1965.members.0.js
species
1

118884
5
no rank

family
1400857
1

1951614
1
no rank

2053538
node1969.members.0.js
species
1

genus
1
1524249

species
node1971.members.0.js
1249552
1

112008
node1972.members.0.js
genus
1

genus
2
2738850

2
species
node1974.members.0.js
2738883

1
103
order
node1975.members.0.js
135614

node1976.members.0.js
family
32033
5
96

genus
node1977.members.0.js
83614
1
6

no rank
2629088
3

1
2901869
species
node1979.members.0.js

2508168
node1980.members.0.js
species
2

2
1176533
node1981.members.0.js
species

14
27
node1982.members.0.js
genus
40323

196198
1
no rank

1
2303750
node1984.members.0.js
species

216778
node1985.members.0.js
species
5

995085
species group
node1986.members.0.js
6
2

3
2
40324
node1987.members.0.js
species

node1988.members.0.js
strain
868597
1

species
node1989.members.0.js
2072414
1

1
species
node1990.members.0.js
128780

1
2709666
genus

node1992.members.0.js
species
2511995
1

2370
2
genus

species
2371
2

2
644357
subspecies
node1995.members.0.js

29
4
338
node1996.members.0.js
genus

node1997.members.0.js
species
56448
3

2
56454
species

453602
no rank
node1999.members.0.js
1

487905
no rank
node2000.members.0.js
1

2
node2001.members.0.js
species
48664

2
3
species
node2002.members.0.js
339

no rank
359385
1

990315
node2004.members.0.js
strain
1

node2005.members.0.js
species
56453
1

species
343
1

227946
node2007.members.0.js
no rank
1

species
node2008.members.0.js
56458
7

species group
3
643453

3
346
species

2
node2011.members.0.js
no rank
454594

no rank
node2012.members.0.js
86040
1

2
species
node2013.members.0.js
2775159

no rank
1
2643310

2081477
node2015.members.0.js
species
1

7
1
141948
node2016.members.0.js
genus

4
1463158
node2017.members.0.js
species

1
2633315
no rank

1
species
node2019.members.0.js
2714945

species
node2020.members.0.js
215691
1

14
2
68
node2021.members.0.js
genus

3
node2022.members.0.js
species
69

2795387
node2023.members.0.js
species
2

node2024.members.0.js
species
2763317
2

1
node2025.members.0.js
species
1324796

1
node2026.members.0.js
species
2591633

2635362
node2027.members.0.js
no rank
3
1

1
2781022
node2028.members.0.js
species

1
node2029.members.0.js
species
2762611

4
1
83618
node2030.members.0.js
genus

node2031.members.0.js
species
314722
1
2

strain
node2032.members.0.js
743721
1

2645906
1
no rank

species
node2034.members.0.js
1871049
1

490567
1
genus

1
node2036.members.0.js
species
370777

6
1775411
family

genus
1
231454

1
node2039.members.0.js
species
1379159

genus
75309
2

node2041.members.0.js
species
582702
2

genus
242605
2

1
242606
species

node2044.members.0.js
strain
1440763
1

2589080
species
node2045.members.0.js
1

genus
2707020
1

1
2010829
species
node2047.members.0.js

1775403
7
order

6
2689614
family

469322
6
genus

6
465721
node2051.members.0.js
species

568386
1
family

413435
1
genus

no rank
2637139
1

1
2303331
node2055.members.0.js
species

order
135618
19

1486721
1
family

no rank
2749459
1

1
species
node2059.members.0.js
417

403
18
family

1
762296
genus

1704499
species
node2062.members.0.js
1

73778
2
genus

2
1432792
node2064.members.0.js
species

genus
1
2822410

species
1
271065

1
1091494
node2067.members.0.js
strain

1
1808977
genus

species
node2069.members.0.js
1808979
1

9
413
genus

9
node2071.members.0.js
species
2681310

1
4
genus
node2072.members.0.js
416

species
421
1

857087
strain
node2074.members.0.js
1

2608980
2
no rank

2
node2076.members.0.js
species
107637

order
41
135619

family
19
28256

1
504090
genus

698828
species
node2080.members.0.js
1

1
204286
genus

1
no rank
node2082.members.0.js
2609414

15
6
2745
node2083.members.0.js
genus

2
1897729
species
node2084.members.0.js

5
1
2609666
no rank
node2085.members.0.js

1
2832500
species
node2086.members.0.js

1
1609967
node2087.members.0.js
species

node2088.members.0.js
species
2811227
1

2854257
species
node2089.members.0.js
1

2609667
species
node2090.members.0.js
2

genus
42054
1

species
158080
1

1
290398
strain
node2093.members.0.js

genus
376488
1

376489
species
node2095.members.0.js
1

family
255527
1

genus
1445504
1

species
1445505
1

1445510
node2099.members.0.js
strain
1

224379
4
family

158481
4
genus

4
158327
species

4
node2103.members.0.js
strain
349521

135620
10
family

48075
2
genus

2
2644139
no rank

2668067
node2107.members.0.js
species
2

genus
2
599651

207949
species
node2109.members.0.js
2

1
187492
genus

1
species
node2111.members.0.js
187493

genus
1537406
1

1
species
node2113.members.0.js
1249553

4
3
28253
node2114.members.0.js
genus

no rank
1
196814

1
2066133
node2116.members.0.js
species

3
191033
family

genus
3
141450

node2119.members.0.js
species
141451
3

family
4
224372

genus
4
59753

no rank
2638842
4

1
2735303
node2123.members.0.js
species

3
1872427
species
node2124.members.0.js

6
135615
order

family
868
6

genus
6
2717

node2128.members.0.js
species
2718
6

no rank
1
50423

723576
node2130.members.0.js
species
1

order
58
135625

58
10
712
family
node2132.members.0.js

1249016
2
genus

2
1032623
species
node2134.members.0.js

genus
155493
1

species
1
750

strain
node2137.members.0.js
1005058
1

75984
4
genus

1
111844
node2139.members.0.js
species

species
node2140.members.0.js
85404
1
3

node2141.members.0.js
strain
1434214
2

genus
node2142.members.0.js
697331
1

1
745
genus

754
node2144.members.0.js
species
1

5
416916
genus

1
node2146.members.0.js
species
714

no rank
2639383
3

species
node2148.members.0.js
2866570
1

2
node2149.members.0.js
species
2820817

732
species
node2150.members.0.js
1

292486
node2151.members.0.js
genus
2
1

1
node2152.members.0.js
species
762

2094023
1
genus

738
node2154.members.0.js
species
1

1
67757
no rank

1
species
node2156.members.0.js
1823760

1649317
2
genus

123824
species
node2158.members.0.js
2

genus
713
18

species
715
1

1
754347
no rank

node2162.members.0.js
strain
754259
1

4
species
node2163.members.0.js
51161

2644856
13
no rank

2774188
node2165.members.0.js
species
13

genus
node2166.members.0.js
724
2
10

1
249188
node2167.members.0.js
species

735
species
node2168.members.0.js
2

1
species
node2169.members.0.js
730

729
1
species

1
862965
strain
node2171.members.0.js

3
node2172.members.0.js
species
727

135624
27
order

84642
24
family

genus
1
43947

species
node2176.members.0.js
43948
1

19
6
642
node2177.members.0.js
genus

1
648794
node2178.members.0.js
species

node2179.members.0.js
species
650
1

node2180.members.0.js
species
644
4

1
257493
no rank

1
species
node2182.members.0.js
2033032

645
node2183.members.0.js
species
2
1

1
29491
node2184.members.0.js
subspecies

1
species
node2185.members.0.js
73010

1
species
node2186.members.0.js
196024

2
654
node2187.members.0.js
species

genus
2
2802288

2
2047966
species
node2189.members.0.js

genus
2
129577

no rank
2
2636315

node2192.members.0.js
species
511062
2

3
node2193.members.0.js
family
83763

185
2887326
order

3
185
node2195.members.0.js
family
468

genus
3
2824158

2283318
node2197.members.0.js
species
3

497
node2198.members.0.js
genus
13
2

node2199.members.0.js
species
861445
1

45610
species
node2200.members.0.js
1

4
3
196806
node2201.members.0.js
no rank

879560
species
node2202.members.0.js
1

256326
species
node2203.members.0.js
5

node2204.members.0.js
genus
469
29
92

70346
node2205.members.0.js
species
3

species
node2206.members.0.js
202956
1

1
node2207.members.0.js
species
1879050

node2208.members.0.js
species
1324350
1

2715164
node2209.members.0.js
species
1

12
202950
species

62977
node2211.members.0.js
strain
12

node2212.members.0.js
species
2053287
1

3
29430
species
node2213.members.0.js

28090
node2214.members.0.js
species
2

106649
node2215.members.0.js
species
1

1
40214
node2216.members.0.js
species

40216
species
node2217.members.0.js
4

1
node2218.members.0.js
species
108981

species
node2219.members.0.js
1789224
2

13
1
909768
node2220.members.0.js
species group

6
7
species
node2221.members.0.js
470

1
1279013
node2222.members.0.js
strain

2
species
node2223.members.0.js
48296

1
node2224.members.0.js
species
1530123

471
species
node2225.members.0.js
1

106654
node2226.members.0.js
species
1

species
node2227.members.0.js
756892
3

2662362
species
node2228.members.0.js
1

1
2136182
species
node2229.members.0.js

11
196816
no rank

5
species
node2231.members.0.js
2563897

1
2853158
species
node2232.members.0.js

1879049
species
node2233.members.0.js
2

1
species
node2234.members.0.js
2004647

472
species
node2235.members.0.js
2

74
14
475
genus
node2236.members.0.js

2
263768
no rank

2
species
node2238.members.0.js
263769

node2239.members.0.js
species
34062
48

species
node2240.members.0.js
34059
6

node2241.members.0.js
species
386891
1

478
node2242.members.0.js
species
3

1553900
16
class

order
213481
11

family
213483
11

1
9
node2246.members.0.js
genus
958

4
7
species
node2247.members.0.js
959

3
node2248.members.0.js
strain
765869

no rank
1
2633795

1
2220073
species
node2250.members.0.js

2862745
2
genus

species
node2252.members.0.js
453816
2

2024979
5
order

3
1652132
family

genus
3
1652133

3
2639665
no rank

species
node2257.members.0.js
2109558
3

family
2
263369

2
146784
genus

node2260.members.0.js
species
960
2

578
10
28216
node2261.members.0.js
class

no rank
2
33809

2
species
node2263.members.0.js
1904640

71
206351
order

4
50
node2265.members.0.js
family
481

1055692
3
genus

682798
node2267.members.0.js
species
3

1
59
genus

63
node2269.members.0.js
species
1

538
1
genus

1
species
node2271.members.0.js
539

1
1193515
genus

node2273.members.0.js
species
1196083
1

32257
4
genus

species
node2275.members.0.js
502
1

species
node2276.members.0.js
504
3

genus
3
334107

153493
species
node2278.members.0.js
3

32
13
482
genus
node2279.members.0.js

1
node2280.members.0.js
species
485

4
2
495
species
node2281.members.0.js

2
88719
subspecies
node2282.members.0.js

1
species
node2283.members.0.js
486

1
node2284.members.0.js
species
2666100

1
484
node2285.members.0.js
species

species
node2286.members.0.js
1107316
1

1
28091
species
node2287.members.0.js

2
2623750
no rank

655307
species
node2289.members.0.js
2

node2290.members.0.js
species
488
2

490
node2291.members.0.js
species
2

3
607711
node2292.members.0.js
species

no rank
1
421605

1
2014784
node2294.members.0.js
species

family
21
1499392

no rank
3
90153

535
node2297.members.0.js
genus
3
2

no rank
2641838
1

2735795
node2299.members.0.js
species
1

885864
1
genus

no rank
1
2619593

1009682
node2302.members.0.js
species
1

168470
1
genus

1
node2304.members.0.js
species
168471

genus
400060
10

10
species
node2306.members.0.js
1225769

genus
4
230666

1737446
species
node2308.members.0.js
4

57739
2
genus

no rank
2
2684990

1
2877939
species
node2311.members.0.js

1
1192162
node2312.members.0.js
species

119066
8
no rank

genus
2
2819026

2
2608985
node2315.members.0.js
species

genus
6
327159

2619054
6
no rank

2053492
node2318.members.0.js
species
6

2
19
order
node2319.members.0.js
206389

75787
4
family

1
90628
no rank

1
species
node2322.members.0.js
151985

genus
551759
1

76116
species
node2324.members.0.js
1

genus
1
1914449

species
node2326.members.0.js
1565605
1

146937
1
genus

1
2609269
no rank

species
node2329.members.0.js
1765049
1

7
2008794
family

2
1
33057
genus
node2331.members.0.js

no rank
2609274
1

1
species
node2333.members.0.js
85643

2
12960
genus

2
41977
species
node2335.members.0.js

3
2891294
genus

2
node2337.members.0.js
species
2815343

1
2016596
species
node2338.members.0.js

5
2008795
family

genus
73029
2

2
281362
species
node2341.members.0.js

genus
88875
2

2
node2343.members.0.js
species
76259

genus
1
138117

1
138118
species
node2345.members.0.js

1
2808923
family

2808942
1
genus

1
1751046
node2348.members.0.js
species

80840
order
node2349.members.0.js
444
14

family
1
995019

40544
1
genus

1
2494234
species
node2352.members.0.js

no rank
80841
2

1469502
node2354.members.0.js
species
1

1
species
node2355.members.0.js
1891238

family
node2356.members.0.js
80864
6
130

2
215579
genus

413882
species
node2358.members.0.js
2

665874
genus
node2359.members.0.js
2
1

2626134
1
no rank

node2361.members.0.js
species
1678129
1

genus
node2362.members.0.js
80865
4
7

1
80866
node2363.members.0.js
species

1
species
node2364.members.0.js
558537

node2365.members.0.js
species
180282
1

29
4
34072
node2366.members.0.js
genus

436515
species
node2367.members.0.js
1

10
9
34073
node2368.members.0.js
species

1
node2369.members.0.js
strain
1246301

663243
node2370.members.0.js
no rank
14
2

node2371.members.0.js
species
434009
1

2
node2372.members.0.js
species
1795631

207745
species
node2373.members.0.js
1

2681552
species
node2374.members.0.js
4

2
species
node2375.members.0.js
2762323

species
node2376.members.0.js
1034889
2

genus
47420
11

2610897
11
no rank

node2379.members.0.js
species
2806347
2

2806262
species
node2380.members.0.js
2

2651974
species
node2381.members.0.js
1

1
species
node2382.members.0.js
2565558

1
species
node2383.members.0.js
795665

1842537
node2384.members.0.js
species
2

species
node2385.members.0.js
2716225
2

genus
6
28065

node2387.members.0.js
species
2509614
1

no rank
1
54524

1
species
node2389.members.0.js
223188

no rank
2
2627954

species
node2391.members.0.js
2741720
1

1
2822760
species
node2392.members.0.js

1842727
node2393.members.0.js
species
2

genus
1
1649468

1
2641902
no rank

node2396.members.0.js
species
2109913
1

genus
683756
3

3
2632542
no rank

2877940
species
node2399.members.0.js
2

node2400.members.0.js
species
2714952
1

283
genus
node2401.members.0.js
14
2

2
2638500
no rank

2853257
species
node2403.members.0.js
1

2597701
species
node2404.members.0.js
1

1562974
species
node2405.members.0.js
1

363952
node2406.members.0.js
species
1

node2407.members.0.js
species
225991
3

species
32013
1

1219032
node2409.members.0.js
strain
1

species
node2410.members.0.js
379895
3

285
species
node2411.members.0.js
1

genus
219181
3

no rank
3
2645081

3
1658672
node2414.members.0.js
species

genus
238749
3

no rank
2649760
1

2714924
species
node2417.members.0.js
1

species
node2418.members.0.js
1288495
2

24
174951
genus

2617605
17
no rank

17
species
node2421.members.0.js
2732511

94132
species
node2422.members.0.js
7
4

3
node2423.members.0.js
strain
365046

12916
genus
node2424.members.0.js
19
5

1
758826
species
node2425.members.0.js

no rank
5
2684926

1
2518343
node2427.members.0.js
species

1
species
node2428.members.0.js
2738413

1
358220
node2429.members.0.js
species

2
2478662
species
node2430.members.0.js

553814
node2431.members.0.js
species
1

species
80867
7

subspecies
node2433.members.0.js
80870
7

family
node2434.members.0.js
75682
5
89

963
genus
node2435.members.0.js
11
1

no rank
75669
1

160236
node2437.members.0.js
species
1

node2438.members.0.js
species
2014887
1

863372
species
node2439.members.0.js
2

no rank
2
2624150

2
node2441.members.0.js
species
2025949

species
341045
2

2
1262470
strain
node2443.members.0.js

node2444.members.0.js
species
964
2

genus
1
303379

node2446.members.0.js
species
1809410
1

genus
1344552
1

2617509
1
no rank

2601898
species
node2449.members.0.js
1

genus
202907
4

2
node2451.members.0.js
species
279113

node2452.members.0.js
species
279058
1

1
158899
species
node2453.members.0.js

genus
1229970
1

no rank
2644401
1

species
node2456.members.0.js
2807626
1

6
53
no rank
node2457.members.0.js
2895353

genus
75654
9

no rank
9
2636909

2728020
node2460.members.0.js
species
6

species
node2461.members.0.js
2728021
3

149698
node2462.members.0.js
genus
38
4

node2463.members.0.js
species
1141883
5

2
871742
node2464.members.0.js
species

node2465.members.0.js
species
945844
4

321985
species
node2466.members.0.js
2

2
species
node2467.members.0.js
321984

2
species
node2468.members.0.js
864828

17
4
2609279
no rank
node2469.members.0.js

3
species
node2470.members.0.js
1707785

2899220
node2471.members.0.js
species
1

1
node2472.members.0.js
species
2861283

1
2861282
node2473.members.0.js
species

1
node2474.members.0.js
species
2765360

1
node2475.members.0.js
species
1678028

2852099
species
node2476.members.0.js
2

2
1337838
species
node2477.members.0.js

2769491
node2478.members.0.js
species
1

29580
node2479.members.0.js
genus
11
1

2610881
4
no rank

1
species
node2481.members.0.js
1537274

1644131
species
node2482.members.0.js
1

1
species
node2483.members.0.js
2861284

1
1236179
species
node2484.members.0.js

species
5
55508

strain
node2486.members.0.js
1349767
5

1
species
node2487.members.0.js
368607

genus
401469
2

2
1
2630295
node2489.members.0.js
no rank

1
2058624
species
node2490.members.0.js

119060
family
node2491.members.0.js
124
11

5
11
node2492.members.0.js
genus
48736

190721
species
node2493.members.0.js
1

305
species
node2494.members.0.js
1

329
node2495.members.0.js
species
4
3

1366050
node2496.members.0.js
strain
1

node2497.members.0.js
genus
32008
6
42

species
node2498.members.0.js
28095
2

node2499.members.0.js
no rank
2613784
1
4

node2500.members.0.js
species
1795874
1

species
node2501.members.0.js
2571746
2

25
8
87882
species group
node2502.members.0.js

292
species
node2503.members.0.js
3

3
60552
node2504.members.0.js
species

species
node2505.members.0.js
101571
1

node2506.members.0.js
species
95486
3
4

strain
node2507.members.0.js
1055524
1

1
species
node2508.members.0.js
1503054

95485
species
node2509.members.0.js
1

node2510.members.0.js
species
152480
2

87883
node2511.members.0.js
species
2

111527
node2512.members.0.js
species group
5
1

node2513.members.0.js
species
342113
1

28450
node2514.members.0.js
species
3

2
1827195
genus

1
no rank
node2516.members.0.js
2646786

species
node2517.members.0.js
758793
1

2
240411
genus

2594795
node2519.members.0.js
species
2

1822464
genus
node2520.members.0.js
21
3

1
2547399
species
node2521.members.0.js

node2522.members.0.js
species
60548
1

1
36873
species

1
strain
node2524.members.0.js
266265

1
75105
species
node2525.members.0.js

node2526.members.0.js
species
2211211
1

species
node2527.members.0.js
311231
1

2
node2528.members.0.js
species
1761016

1
92647
species
node2529.members.0.js

1
148447
species

node2531.members.0.js
strain
391038
1

1
311230
node2532.members.0.js
species

node2533.members.0.js
species
134537
5

node2534.members.0.js
species
134536
1

1
2571747
node2535.members.0.js
species

genus
1810868
5

2649241
3
no rank

3
species
node2538.members.0.js
2651972

species
node2539.members.0.js
1553431
2

genus
8
44013

576610
species
node2541.members.0.js
3

576611
node2542.members.0.js
species
2

2640945
3
no rank

1
2081041
node2544.members.0.js
species

1855616
node2545.members.0.js
species
1

species
node2546.members.0.js
2576928
1

93217
2
genus

species
node2548.members.0.js
656179
1

656178
node2549.members.0.js
species
1

132490
1
no rank

1
node2551.members.0.js
species
2030806

genus
47670
3

3
47671
node2553.members.0.js
species

106589
genus
node2554.members.0.js
16
4

4
164546
species
node2555.members.0.js

species
node2556.members.0.js
119219
2

1
species
node2557.members.0.js
151783

4
node2558.members.0.js
species
96344

1
1796606
species
node2559.members.0.js

57
506
family

genus
node2561.members.0.js
222
1
19

1353891
node2562.members.0.js
species
12

1353889
node2563.members.0.js
species
1

node2564.members.0.js
species
217203
1

2
species
node2565.members.0.js
85698

1
1389922
node2566.members.0.js
species

node2567.members.0.js
species
32002
1

node2568.members.0.js
genus
90243
1
3

1
90244
species
node2569.members.0.js

node2570.members.0.js
species
90245
1

517
17
genus

1331258
node2572.members.0.js
species
1

521
node2573.members.0.js
species
2

1
node2574.members.0.js
species
463040

2630031
9
no rank

1697043
node2576.members.0.js
species
7

1746199
node2577.members.0.js
species
2

94624
species
node2578.members.0.js
1

2
463014
node2579.members.0.js
species

1
node2580.members.0.js
species
103855

5
1921582
genus

4
1851544
node2582.members.0.js
species

2163011
species
node2583.members.0.js
1

152267
2
genus

2
2626614
no rank

node2586.members.0.js
species
2488560
2

1
6
genus
node2587.members.0.js
507

1
node2588.members.0.js
species
2582914

4
species
node2589.members.0.js
511

no rank
2
83496

2
2593958
species
node2591.members.0.js

305976
3
genus

no rank
1
2640016

1007105
node2594.members.0.js
species
1

species
node2595.members.0.js
2028345
2

2
208545
no rank

2
species
node2597.members.0.js
208544

224471
no rank
node2598.members.0.js
25
1

316612
genus
node2599.members.0.js
1

88
2
genus

species
2
34029

395495
strain
node2602.members.0.js
2

1
2
node2603.members.0.js
genus
212743

no rank
2640088
1

2753607
node2605.members.0.js
species
1

genus
3
28067

28068
node2607.members.0.js
species
3

2
318147
genus

2642959
2
no rank

species
node2610.members.0.js
1768242
2

93681
2
genus

2
76731
species
node2612.members.0.js

genus
114248
10

10
307486
species
node2614.members.0.js

34102
2
genus

34103
2
species

2
subspecies
node2617.members.0.js
639200

no rank
47926
2

node2619.members.0.js
species
86027
2

order
32003
22

9
2803844
family

9
1
2803845
genus
node2622.members.0.js

species
node2623.members.0.js
2732067
4

4
2732487
node2624.members.0.js
species

90627
1
family

1443590
1
genus

1188319
node2627.members.0.js
species
1

family
2008790
2

919
2
genus

species
2
36861

2
292415
strain
node2631.members.0.js

206379
5
family

35798
1
genus

species
node2634.members.0.js
1288494
1

4
914
genus

node2636.members.0.js
species
44574
1

3
node2637.members.0.js
species
1444684

2008793
3
family

311181
1
genus

species
node2640.members.0.js
311182
1

no rank
2
2211107

species
node2642.members.0.js
2211108
1

1
species
node2643.members.0.js
2496847

1
2772226
family

935200
1
genus

649841
1
species

strain
node2647.members.0.js
1163617
1

32011
1
family

genus
359407
1

species
1055487
1

1
strain
node2651.members.0.js
666681

28211
node2652.members.0.js
class
869
14

order
56
204458

3
56
node2654.members.0.js
family
76892

36
8
41275
genus
node2655.members.0.js

4
588932
species
node2656.members.0.js

1
species
node2657.members.0.js
74329

2800818
node2658.members.0.js
species
1

no rank
20
2622653

6
node2660.members.0.js
species
2861285

species
node2661.members.0.js
1938605
2

1
2579977
node2662.members.0.js
species

species
node2663.members.0.js
2591463
7

1827469
node2664.members.0.js
species
1

3
2752515
species
node2665.members.0.js

node2666.members.0.js
species
2774190
2

20
genus
node2667.members.0.js
7
1

2640670
1
no rank

1
node2669.members.0.js
species
2803784

1
species
node2670.members.0.js
2201350

284016
species
node2671.members.0.js
4
2

450851
node2672.members.0.js
strain
2

75
6
genus

no rank
2
2648921

2823693
node2675.members.0.js
species
2

1
69395
species
node2676.members.0.js

88688
node2677.members.0.js
species
2

1
species
node2678.members.0.js
2010972

76890
genus
node2679.members.0.js
4
1

78587
species
node2680.members.0.js
3
1

strain
node2681.members.0.js
573065
2

order
2800059
7

family
7
2800061

3
2004661
genus

no rank
2638931
3

3
2749086
node2686.members.0.js
species

74317
2
genus

species
74318
2

2
394221
node2689.members.0.js
strain

1649466
1
genus

1
2029849
node2691.members.0.js
species

1
1433402
genus

1
node2693.members.0.js
species
1434191

no rank
2
47925

1
node2695.members.0.js
species
91750

116865
node2696.members.0.js
species
1

204441
94
order

3
48
family
node2698.members.0.js
433

genus
2919364
1

1
node2700.members.0.js
species
436

genus
2
441

318683
node2702.members.0.js
species
2

genus
1
2791118

species
node2704.members.0.js
2558360
1

1
5
genus
node2705.members.0.js
434

subgenus
2
151157

2
node2707.members.0.js
species
435

node2708.members.0.js
species
1076596
1

node2709.members.0.js
species
2500548
1

22
10
125216
genus
node2710.members.0.js

1
node2711.members.0.js
species
257708

node2712.members.0.js
species
2768162
3

2617492
3
no rank

node2714.members.0.js
species
2897332
3

species
node2715.members.0.js
2768161
4

node2716.members.0.js
species
207340
1

genus
35812
9

2771361
node2718.members.0.js
species
9

genus
1
1434011

1
265960
species
node2720.members.0.js

genus
1
50709

1
2648610
no rank

1
406384
species
node2723.members.0.js

2804525
1
genus

1484109
node2725.members.0.js
species
1

genus
2603324
1

1969806
node2727.members.0.js
species
1

genus
153497
1

species
node2729.members.0.js
153496
1

41295
15
family

genus
2705399
2

2602016
species
node2732.members.0.js
2

13134
5
genus

species
84159
1

1
342108
node2735.members.0.js
strain

2
node2736.members.0.js
species
55518

no rank
node2737.members.0.js
2617991
1
2

1007128
node2738.members.0.js
species
1

genus
1
1081

1
1085
node2740.members.0.js
species

6
111830
genus

5
111831
node2742.members.0.js
species

1
2642886
no rank

1
2682137
species
node2744.members.0.js

1
1231242
genus

1
2594003
node2746.members.0.js
species

family
2813951
2

genus
196080
2

no rank
2614952
2

2
2560057
species
node2750.members.0.js

no rank
941843
1

1
species
node2752.members.0.js
2026786

3
2844866
family

genus
3
168934

2648997
3
no rank

3
node2756.members.0.js
species
2785911

2
2690195
family

457933
2
genus

2
node2759.members.0.js
species
2843330

5
2844601
family

93
5
genus

no rank
5
2622320

node2763.members.0.js
species
2558175
5

family
17
2829815

2478349
2
genus

2220096
node2766.members.0.js
species
2

1543705
2
genus

species
28077
2

2
1441467
node2769.members.0.js
strain

genus
node2770.members.0.js
191
5
13

192
node2771.members.0.js
species
2

node2772.members.0.js
species
1226968
1

1
1064539
node2773.members.0.js
species

no rank
3
2630922

2
species
node2775.members.0.js
709810

node2776.members.0.js
species
137722
1

node2777.members.0.js
species
2202148
1

family
1
597359

genus
1
1647175

2775080
species
node2780.members.0.js
1

2066490
1
order

family
1
2066491

2813803
1
genus

species
node2784.members.0.js
2582913
1

no rank
1
82117

213485
1
genus

species
node2787.members.0.js
349221
1

2
1921002
order

2100208
1
family

genus
2100210
1

1
node2791.members.0.js
species
1629334

family
44746
1

genus
1
2601574

2601575
node2794.members.0.js
species
1

14
362
node2795.members.0.js
order
356

2813035
5
family

3
1742974
genus

1712261
node2798.members.0.js
species
3

2
256616
genus

species
2
256618

2
node2801.members.0.js
strain
402881

family
node2802.members.0.js
2831106
1
20

node2803.members.0.js
genus
46913
3
19

3
2860336
node2804.members.0.js
species

2
2774137
species
node2805.members.0.js

no rank
196773
10

2806348
species
node2807.members.0.js
2

1736675
node2808.members.0.js
species
4

4
2083786
node2809.members.0.js
species

211433
1
no rank

1
species
node2811.members.0.js
211434

family
1
2036754

genus
28209
1

no rank
2638111
1

1
1702325
node2815.members.0.js
species

node2816.members.0.js
family
82115
7
90

genus
323620
2

species
node2818.members.0.js
352475
1

2643062
1
no rank

879274
node2820.members.0.js
species
1

genus
34019
1

1
node2822.members.0.js
species
34020

genus
1
1648508

1969821
node2824.members.0.js
species
1

genus
2
2853332

node2826.members.0.js
species
1813451
2

node2827.members.0.js
no rank
227290
17
70

genus
7
1525371

no rank
2629175
1

1
species
node2830.members.0.js
2060726

399
node2831.members.0.js
species
6

16
1
357
genus
node2832.members.0.js

359
species
node2833.members.0.js
1

node2834.members.0.js
no rank
2632611
1
2

2735528
node2835.members.0.js
species
1

373
species
node2836.members.0.js
2

species group
10
1183400

species
node2838.members.0.js
358
10

3
29
node2839.members.0.js
genus
379

species
398
1

1
node2841.members.0.js
strain
698761

1
node2842.members.0.js
species
2267833

9
1
2613769
no rank
node2843.members.0.js

2
species
node2844.members.0.js
1480001

1
node2845.members.0.js
species
2028343

1
2806346
species
node2846.members.0.js

1
species
node2847.members.0.js
2048897

2
node2848.members.0.js
species
1981173

1
species
node2849.members.0.js
1571470

node2850.members.0.js
species
240521
1

384
node2851.members.0.js
species
9
2

7
4
386
node2852.members.0.js
no rank

754523
node2853.members.0.js
strain
1

1033991
strain
node2854.members.0.js
2

451876
species
node2855.members.0.js
1

species
29449
2

538025
strain
node2857.members.0.js
2

1
species
node2858.members.0.js
1538158

species
node2859.members.0.js
648995
1

genus
1903858
1

1
1335061
node2861.members.0.js
species

no rank
227292
7

node2863.members.0.js
genus
28105
1
6

1
2613772
no rank

node2865.members.0.js
species
2613773
1

110321
2
species

366394
strain
node2867.members.0.js
2

species group
2
663276

species
380
2

1
node2870.members.0.js
strain
1185652

node2871.members.0.js
strain
1128330
1

genus
106591
1

1
1752398
node2873.members.0.js
species

49
1
119045
family
node2874.members.0.js

2
186650
genus

2
species
node2876.members.0.js
2651334

39
7
407
genus
node2877.members.0.js

1
1775910
node2878.members.0.js
species

2
species
node2879.members.0.js
270351

3
2202827
node2880.members.0.js
species

node2881.members.0.js
species
114616
1
2

1
460265
strain
node2882.members.0.js

17
species
node2883.members.0.js
570505

2
node2884.members.0.js
species
269660

species
39956
1

1
node2886.members.0.js
strain
908290

2202828
species
node2887.members.0.js
1

2
3
node2888.members.0.js
no rank
2615210

739141
species
node2889.members.0.js
1

1
7
node2890.members.0.js
genus
2282523

1
408
node2891.members.0.js
species

5
223967
node2892.members.0.js
species

9
45401
family

3
29407
genus

3
2619116
no rank

species
node2896.members.0.js
674703
3

genus
45402
2

2663263
2
no rank

2
node2899.members.0.js
species
2820278

2827482
1
genus

169176
1
species

1
1219035
strain
node2902.members.0.js

81
3
genus

1
2619925
no rank

1
species
node2905.members.0.js
2682968

1
53399
species

1
node2907.members.0.js
strain
582899

1427356
node2908.members.0.js
species
1

family
2831100
5

5
1
85413
node2910.members.0.js
genus

no rank
2
2653178

2
species
node2912.members.0.js
2015316

node2913.members.0.js
species
1526658
2

119042
4
no rank

1734920
2
genus

1235591
species
node2916.members.0.js
2

genus
2
169055

2
2623691
no rank

2712222
species
node2919.members.0.js
2

2843308
5
family

genus
261933
5

2627136
5
no rank

5
species
node2923.members.0.js
1885025

2821832
node2924.members.0.js
family
7
2

genus
478070
2

no rank
2
2648686

species
node2927.members.0.js
2021862
1

1
2590016
species
node2928.members.0.js

genus
152161
1

538381
species
node2930.members.0.js
1

1
227873
genus

121719
node2932.members.0.js
species
1

genus
1
150830

187304
node2934.members.0.js
species
1

3
2843305
family

1632780
3
genus

1868589
node2937.members.0.js
species
3

1
41292
no rank

1
1862950
node2939.members.0.js
species

31993
5
family

1
2041902
genus

1
1437009
species
node2942.members.0.js

4
133
genus

173366
node2944.members.0.js
species
1

1
655015
node2945.members.0.js
species

species
node2946.members.0.js
391905
2

family
772
8

genus
773
8

2267275
species
node2949.members.0.js
1

33045
1
species

node2951.members.0.js
strain
634504
1

4
2645622
no rank

species
node2953.members.0.js
2759660
3

node2954.members.0.js
species
159745
1

33047
1
species

subspecies
40933
1

1
1094497
node2957.members.0.js
strain

species
node2958.members.0.js
803
1

3
2831104
family

3
1406135
genus

no rank
2624130
3

3
2304600
species
node2962.members.0.js

14
335928
family

279
4
genus

280
node2965.members.0.js
species
3
1

strain
node2966.members.0.js
78245
2

1
species
node2967.members.0.js
2528964

2
204476
genus

no rank
2688601
2

2
node2970.members.0.js
species
2789216

2
152053
genus

2
2615126
no rank

2
species
node2973.members.0.js
2709380

1
6
genus

7
1
species

1
node2976.members.0.js
strain
438753

genus
556257
1

331696
species
node2978.members.0.js
1

genus
4
99

3
2626613
no rank

3
1850374
node2981.members.0.js
species

species
node2982.members.0.js
1745854
1

3
118882
family

3
1
2826938
no rank
node2984.members.0.js

528
1
genus

no rank
1
239106

1986154
node2987.members.0.js
species
1

234
1
genus

1
node2989.members.0.js
species
529

32
1
69277
family
node2990.members.0.js

genus
1
1594166

1
species
node2992.members.0.js
2742145

1
28100
genus

species
node2994.members.0.js
1867719
1

1649463
1
genus

2647570
1
no rank

2305987
node2997.members.0.js
species
1

6
27
genus
node2998.members.0.js
68287

species
2066070
1

1
266835
node3000.members.0.js
strain

17
2
325217
no rank
node3001.members.0.js

2880934
species
node3002.members.0.js
1

3
node3003.members.0.js
species
1854057

2
2493675
species
node3004.members.0.js

2108445
node3005.members.0.js
species
1

1
2483404
species
node3006.members.0.js

1
2493669
species
node3007.members.0.js

1
2493677
species
node3008.members.0.js

node3009.members.0.js
species
2493676
1

1
2654248
node3010.members.0.js
species

species
node3011.members.0.js
2589967
1

2777475
species
node3012.members.0.js
1

node3013.members.0.js
species
2589974
1

2
species
node3014.members.0.js
28104

species
39645
1

no rank
278148
1

1
strain
node3017.members.0.js
765698

genus
1
2911176

node3019.members.0.js
species
675281
1

41294
family
node3020.members.0.js
69
11

14
49
genus
node3021.members.0.js
374

5
species
node3022.members.0.js
858423

1355477
species
node3023.members.0.js
3

1
2748629
species
node3024.members.0.js

species
node3025.members.0.js
722472
1

375
species
node3026.members.0.js
2

2631580
no rank
node3027.members.0.js
12
1

376
species
node3028.members.0.js
2

1
species
node3029.members.0.js
858422

1
1325100
species
node3030.members.0.js

114615
node3031.members.0.js
species
1

2840469
species
node3032.members.0.js
1

1223566
species
node3033.members.0.js
1

1
node3034.members.0.js
species
115808

2
2715960
species
node3035.members.0.js

node3036.members.0.js
species
1325114
1

1
83637
species
node3037.members.0.js

1325090
node3038.members.0.js
species
2

2
species
node3039.members.0.js
1274631

node3040.members.0.js
species
1437360
1

species
node3041.members.0.js
1325095
5

9
1073
genus

node3043.members.0.js
species
1076
1
4

3
316057
strain
node3044.members.0.js

4
2638247
no rank

node3046.members.0.js
species
340268
4

475937
node3047.members.0.js
species
1

family
4
45404

532
1
genus

species
1
533

31994
node3051.members.0.js
subspecies
1

genus
1156568
2

2
2640612
no rank

species
node3054.members.0.js
2822761
2

genus
1
120652

species
199596
1

395965
node3057.members.0.js
strain
1

11
255475
family

genus
182269
3

no rank
2638230
3

3
2906072
node3061.members.0.js
species

genus
414371
8

no rank
2615206
6

species
node3064.members.0.js
1638162
2

3
2816454
node3065.members.0.js
species

1
node3066.members.0.js
species
2826993

species
node3067.members.0.js
2758041
2

766
10
order

1
2603433
family

genus
1
2603321

2163644
species
node3071.members.0.js
1

family
942
7

943
1
genus

species group
106178
1

1
944
node3075.members.0.js
species

tribe
4
952

genus
953
4

955
node3078.members.0.js
species
1

node3079.members.0.js
no rank
2640676
1
3

1812111
node3080.members.0.js
species
1

116598
species
node3081.members.0.js
1

genus
1
33993

species
1
33994

node3084.members.0.js
strain
1286528
1

1
768
genus

142058
1
species

node3087.members.0.js
strain
1248439
1

775
2
family

tribe
2
33988

2
780
genus

114295
1
no rank

species
337479
1

1
1182263
node3093.members.0.js
strain

1
node3094.members.0.js
species group
114277

10
178
node3095.members.0.js
order
204457

41297
node3096.members.0.js
family
135
7

1649486
7
genus

2
node3098.members.0.js
species
160791

3
2617679
no rank

2711156
species
node3100.members.0.js
3

2
node3101.members.0.js
species
1850238

genus
node3102.members.0.js
165695
2
14

1
46429
species

690566
node3104.members.0.js
strain
1

1
species
node3105.members.0.js
1332080

2
13690
species
node3106.members.0.js

332055
species
node3107.members.0.js
2
1

861109
strain
node3108.members.0.js
1

node3109.members.0.js
species
121428
1

4
2
2611147
no rank
node3110.members.0.js

1
node3111.members.0.js
species
1855519

1
627192
species
node3112.members.0.js

node3113.members.0.js
species
135719
1

genus
1763827
2

1763828
node3115.members.0.js
species
2

genus
2676233
1

2711215
node3117.members.0.js
species
1

19
82
node3118.members.0.js
genus
13687

5
species
node3119.members.0.js
2599297

1
node3120.members.0.js
species
1549858

1
653931
species
node3121.members.0.js

68569
node3122.members.0.js
species
1

species
node3123.members.0.js
1560345
2

node3124.members.0.js
species
152682
2

2
2319844
node3125.members.0.js
species

node3126.members.0.js
no rank
196159
9
37

species
node3127.members.0.js
1523415
1

node3128.members.0.js
species
2903960
1

species
node3129.members.0.js
2219696
2

2
1327635
node3130.members.0.js
species

2
node3131.members.0.js
species
1517554

2565555
node3132.members.0.js
species
5

1390395
species
node3133.members.0.js
1

2
2735134
species
node3134.members.0.js

4
1517551
species
node3135.members.0.js

1
node3136.members.0.js
species
1811332

2899123
species
node3137.members.0.js
1

1
species
node3138.members.0.js
28214

node3139.members.0.js
species
1938607
1

2
2492837
species
node3140.members.0.js

1
745310
species
node3141.members.0.js

1961362
species
node3142.members.0.js
1

2759526
species
node3143.members.0.js
1

2698828
node3144.members.0.js
species
1

1
1813879
node3145.members.0.js
species

1
2681549
species
node3146.members.0.js

1
species
node3147.members.0.js
424800

3
species
node3148.members.0.js
2698679

species
node3149.members.0.js
1045317
1

363835
species
node3150.members.0.js
3

genus
1
2823232

1
node3152.members.0.js
species
941907

1
8
node3153.members.0.js
genus
165696

1
1176536
species
node3154.members.0.js

4
2644732
no rank

1016987
species
node3156.members.0.js
1

2
2571749
node3157.members.0.js
species

1
species
node3158.members.0.js
1609758

2
158500
node3159.members.0.js
species

genus
8
165697

no rank
1
310580

1
species
node3162.members.0.js
310581

no rank
2614943
5

2486273
species
node3164.members.0.js
1

species
node3165.members.0.js
1357916
1

1
2565556
node3166.members.0.js
species

1
node3167.members.0.js
species
292913

1874061
node3168.members.0.js
species
1

2
33050
node3169.members.0.js
species

1234545
5
genus

3
2621071
no rank

3
node3172.members.0.js
species
2724527

species
node3173.members.0.js
2026624
2

2820280
5
family

56358
1
genus

2634456
1
no rank

2759707
node3177.members.0.js
species
1

no rank
2836943
3

2836944
node3179.members.0.js
species
3

1
335405
genus

no rank
1
2644549

node3182.members.0.js
species
1892855
1

family
2844881
1

541
1
genus

1
542
species

120044
1
subspecies

1
strain
node3187.members.0.js
579138

27
5
335929
node3188.members.0.js
family

1855416
2
genus

2
266951
species
node3190.members.0.js

2800788
node3191.members.0.js
no rank
12
2

genus
1111
1

2683265
1
no rank

species
node3194.members.0.js
2003315
1

node3195.members.0.js
genus
1041
1
9

no rank
2633097
1

1042
node3197.members.0.js
species
1

node3198.members.0.js
species
39960
6
7

strain
node3199.members.0.js
314225
1

genus
3
2800686

3
species
node3201.members.0.js
692370

genus
4
361177

no rank
2614945
4

2185142
species
node3204.members.0.js
1

3
2067415
species
node3205.members.0.js

1
1295327
node3206.members.0.js
genus

order
1
1191478

1191479
1
family

genus
162171
1

1124597
1
species

1
strain
node3211.members.0.js
156889

order
1
54526

1655514
1
family

genus
1
198251

2647897
1
no rank

1
1977865
species
node3216.members.0.js

7
136
order
node3217.members.0.js
204455

1
35
node3218.members.0.js
family
2854170

genus
2919626
2

species
node3220.members.0.js
311180
2

genus
1
58842

no rank
1
2624628

1
2009329
node3223.members.0.js
species

genus
92944
1

92947
node3225.members.0.js
species
1

191028
2
genus

133924
1
species

999552
strain
node3228.members.0.js
1

species
node3229.members.0.js
1396826
1

367771
2
genus

2
42444
species

988812
strain
node3232.members.0.js
2

2
263377
genus

1
node3234.members.0.js
species
1250539

1
1229727
node3235.members.0.js
species

74030
genus
node3236.members.0.js
2
1

1
540747
node3237.members.0.js
species

genus
60136
8

no rank
196795
5

1968541
species
node3240.members.0.js
1

4
1389005
node3241.members.0.js
species

2
1402135
species
node3242.members.0.js

1
species
node3243.members.0.js
1917485

2
97050
genus

no rank
2625375
2

2
species
node3246.members.0.js
2587853

299261
1
genus

species
node3248.members.0.js
299262
1

1
3
genus
node3249.members.0.js
302485

species
node3250.members.0.js
1844006
2

genus
53945
1

species
53946
1

1
391616
node3253.members.0.js
strain

genus
1
188905

1
2645469
no rank

1
2856823
node3256.members.0.js
species

genus
875170
6

4
species
node3258.members.0.js
1758178

2
875171
node3259.members.0.js
species

family
node3260.members.0.js
31989
4
94

8
55
genus
node3261.members.0.js
265

2560053
node3262.members.0.js
species
2

3
node3263.members.0.js
species
1545044

2
59779
species
node3264.members.0.js

11
147645
node3265.members.0.js
species

species
node3266.members.0.js
1945662
2

node3267.members.0.js
species
2259340
2

2688777
20
no rank

1
species
node3269.members.0.js
2500532

node3270.members.0.js
species
235898
1

9
species
node3271.members.0.js
2760307

node3272.members.0.js
species
2903900
3

2589076
species
node3273.members.0.js
6

2
species
node3274.members.0.js
266

species
node3275.members.0.js
1077935
3

2
1844015
genus

species
node3277.members.0.js
2789856
2

1
285107
node3278.members.0.js
genus

1060
3
genus

1061
1
species

1
strain
node3281.members.0.js
272942

2
196779
no rank

1
species
node3283.members.0.js
2033869

1
species
node3284.members.0.js
1221569

2683599
3
genus

3
2579971
node3286.members.0.js
species

genus
2820509
2

node3288.members.0.js
species
2820523
2

genus
2778525
1

2171755
node3290.members.0.js
species
1

genus
1
1759417

1
node3292.members.0.js
species
1591409

genus
1855413
3

3
1267768
species
node3294.members.0.js

1
2738399
genus

1
node3296.members.0.js
species
2483033

genus
5
2816884

5
species
node3298.members.0.js
2599296

1
1679449
genus

node3300.members.0.js
species
1461694
1

genus
1434002
1

no rank
1
2633719

2692330
species
node3303.members.0.js
1

1
1775705
genus

node3305.members.0.js
species
1335048
1

1
34008
genus

35806
species
node3307.members.0.js
1

119541
1
genus

node3309.members.0.js
species
441209
1

204456
2
genus

2
2840474
node3311.members.0.js
species

genus
node3312.members.0.js
1653176
1
3

species
node3313.members.0.js
1063
2

genus
1
2083206

node3315.members.0.js
species
379347
1

no rank
58840
2

1
2683284
node3317.members.0.js
species

1904441
species
node3318.members.0.js
1

order
2800060
2

family
2
69657

genus
85
1

no rank
2630699
1

2016196
node3323.members.0.js
species
1

1
2892997
genus

species
node3325.members.0.js
1759059
1

no rank
2
33807

2
2829083
node3327.members.0.js
species

1807140
2
class

order
225057
2

family
2
225058

119977
node3331.members.0.js
genus
2
1

1
1689834
species
node3332.members.0.js

no rank
2
81684

2
1977087
node3334.members.0.js
species

node3335.members.0.js
clade
1783272
25
3532

4
2109258
phylum

4
2109259
class

4
2109260
order

family
4
2109261

genus
2109263
4

species
node3341.members.0.js
1960156
4

201174
node3342.members.0.js
phylum
2007
5

15
84998
class

1643822
7
order

node3345.members.0.js
family
1643826
1
7

genus
2
84111

84112
node3347.members.0.js
species
2

genus
2005386
1

1870985
node3349.members.0.js
species
1

84108
1
genus

1
species
node3351.members.0.js
84110

genus
2
1926677

1907662
species
node3353.members.0.js
2

order
8
84999

1643824
node3355.members.0.js
family
4
1

genus
133925
3

1805478
node3357.members.0.js
species
2

2638792
1
no rank

2109685
species
node3359.members.0.js
1

84107
4
family

102106
3
genus

74426
node3362.members.0.js
species
1

species
147206
2

2
445975
node3364.members.0.js
strain

genus
33870
1

species
1
33871

strain
node3367.members.0.js
700015
1

class
node3368.members.0.js
1760
54
1946

2
622452
order

family
83778
2

2
33981
genus

species
node3372.members.0.js
131568
1
2

1
node3373.members.0.js
strain
266940

order
2
85014

2
85034
family

2838311
1
genus

1
2811108
species
node3377.members.0.js

1
58113
genus

no rank
2637084
1

1
node3380.members.0.js
species
2867006

order
85010
59

1
59
node3382.members.0.js
family
2070

1
674734
genus

no rank
1
2636053

1653480
species
node3385.members.0.js
1

genus
1
142577

species
node3387.members.0.js
530584
1

22
1813
genus

2
node3389.members.0.js
species
33910

14
1
2618356
node3390.members.0.js
no rank

8
node3391.members.0.js
species
2914159

2
2742131
species
node3392.members.0.js

1
species
node3393.members.0.js
2745196

2653857
species
node3394.members.0.js
2

1
2842453
node3395.members.0.js
species

1
4
species group
node3396.members.0.js
2893673

129921
species
node3397.members.0.js
1

2
208439
node3398.members.0.js
species

31958
node3399.members.0.js
species
1

genus
2071
10

5
103733
species
node3401.members.0.js

5
2593673
no rank

node3403.members.0.js
species
2912058
1

node3404.members.0.js
species
2781735
4

165301
2
genus

2
species
node3406.members.0.js
1586287

2029
1
genus

species
node3408.members.0.js
860235
1

genus
17
1847

species
node3410.members.0.js
240495
1
3

node3411.members.0.js
strain
675635
2

2074
species
node3412.members.0.js
1

2619320
node3413.members.0.js
no rank
13
2

species
node3414.members.0.js
445576
2

2761535
node3415.members.0.js
species
2

687424
species
node3416.members.0.js
1

6
2865833
species
node3417.members.0.js

1
2893576
genus

species
node3419.members.0.js
2665642
1

65496
genus
node3420.members.0.js
3
1

no rank
2
2644606

1612551
species
node3422.members.0.js
1

1
species
node3423.members.0.js
2072503

2039638
4
order

2162846
4
family

genus
622681
4

node3427.members.0.js
species
1884914
4

414714
2
order

2
414877
family

genus
414878
2

species
2
304895

node3432.members.0.js
strain
479433
2

1
263
node3433.members.0.js
order
85009

family
2726069
2

genus
2
182639

1544730
node3436.members.0.js
species
2

7
182
family
node3437.members.0.js
31957

genus
11
29404

no rank
1
2619695

species
node3440.members.0.js
2672569
1

29405
node3441.members.0.js
species
2
1

1032480
strain
node3442.members.0.js
1

4
2596828
node3443.members.0.js
species

4
node3444.members.0.js
species
546874

1
2717325
genus

1
2717326
no rank

node3447.members.0.js
species
2819349
1

1912216
node3448.members.0.js
genus
139
2

node3449.members.0.js
species
33011
14

1747
species
node3450.members.0.js
113
111

node3451.members.0.js
strain
909952
2

2559073
node3452.members.0.js
species
7

3
33010
node3453.members.0.js
species

1278221
1
genus

1
species
node3455.members.0.js
675864

1743
genus
node3456.members.0.js
2
1

1
node3457.members.0.js
species
556499

genus
node3458.members.0.js
72763
1
18

species
node3459.members.0.js
1332264
1

11
species
node3460.members.0.js
2161816

3
1610493
species
node3461.members.0.js

1
node3462.members.0.js
species
1285901

1
species
node3463.members.0.js
399497

1912215
1
genus

node3465.members.0.js
species
1748
1

genus
2
2801844

2
1750
node3467.members.0.js
species

4
78
family
node3468.members.0.js
85015

59
11
1839
genus
node3469.members.0.js

1
23
node3470.members.0.js
no rank
2615069

2
2898796
species
node3471.members.0.js

2
2663857
species
node3472.members.0.js

2803858
species
node3473.members.0.js
1

2017486
species
node3474.members.0.js
4

2714939
species
node3475.members.0.js
2

species
node3476.members.0.js
2582905
3

node3477.members.0.js
species
2840457
2

species
node3478.members.0.js
2736757
1

2
node3479.members.0.js
species
196162

2685869
node3480.members.0.js
species
2

2763008
species
node3481.members.0.js
1

1
2575440
node3482.members.0.js
species

4
2712223
node3483.members.0.js
species

2558918
species
node3484.members.0.js
4

species
node3485.members.0.js
2045452
2

2714938
species
node3486.members.0.js
7

2589074
node3487.members.0.js
species
1

1
449461
node3488.members.0.js
species

1
species
node3489.members.0.js
2760089

1774216
node3490.members.0.js
species
2

node3491.members.0.js
species
1804624
1

160826
node3492.members.0.js
species
1

genus
3
53387

species
node3494.members.0.js
546871
3

1
8
node3495.members.0.js
genus
2040

2662028
species
node3496.members.0.js
1

3
2633570
no rank

1
2107713
node3498.members.0.js
species

2
species
node3499.members.0.js
2663859

3
1736691
node3500.members.0.js
species

no rank
116532
1

node3502.members.0.js
species
1871072
1

1
86795
genus

1
node3504.members.0.js
species
642780

genus
2
116071

species
node3506.members.0.js
75385
2

order
85013
10

family
74712
10

1854
node3509.members.0.js
genus
10
5

1
181581
no rank

181582
species
node3511.members.0.js
1

2
298654
node3512.members.0.js
species

2
1859
species

2
node3514.members.0.js
strain
326424

order
85012
24

2012
7
family

genus
1988
6

2
46165
species
node3518.members.0.js

2750812
node3519.members.0.js
species
2

no rank
2
2626254

2591108
node3521.members.0.js
species
1

node3522.members.0.js
species
1219491
1

genus
2019
1

species
1
2020

1
471852
node3525.members.0.js
strain

8
83676
family

genus
104204
1

no rank
2635841
1

2498135
species
node3529.members.0.js
1

genus
node3530.members.0.js
2013
1
7

53437
3
species

1205910
node3532.members.0.js
strain
3

no rank
3
2649073

1
2831970
node3534.members.0.js
species

2831968
species
node3535.members.0.js
2

family
9
2004

1
2000
genus

2632669
1
no rank

1
2202249
node3539.members.0.js
species

genus
8
83681

species
node3541.members.0.js
93944
1

species
1
404386

1
node3543.members.0.js
strain
1122611

species
node3544.members.0.js
2656914
2

2593643
4
no rank

node3546.members.0.js
species
1909395
4

41
1643682
order

11
41
family
node3548.members.0.js
85030

9
1860
genus

9
8
1861
species
node3550.members.0.js

526225
strain
node3551.members.0.js
1

14
88138
genus

4
2643866
no rank

4
species
node3554.members.0.js
2851567

10
477641
node3555.members.0.js
species

no rank
3
234661

3
2596920
species
node3557.members.0.js

genus
4
38501

4
3
138336
species
node3559.members.0.js

1
1146883
node3560.members.0.js
strain

1643684
10
order

family
85031
10

genus
53460
10

2
2618968
no rank

species
node3565.members.0.js
2762325
2

7
species
node3566.members.0.js
1902245

53461
node3567.members.0.js
species
1

order
85004
21

31953
node3569.members.0.js
family
21
2

4
419014
genus

419015
node3571.members.0.js
species
4

node3572.members.0.js
genus
1678
5
13

1
158787
species

1
node3574.members.0.js
strain
1150461

node3575.members.0.js
species
1680
1

species
78448
1

1
subspecies
node3577.members.0.js
78344

no rank
3
135024

node3579.members.0.js
species
165187
3

species
node3580.members.0.js
1684
2

2701
node3581.members.0.js
genus
1

genus
1
196082

78258
1
species

1
node3584.members.0.js
strain
864564

85008
49
order

49
2
28056
family
node3586.members.0.js

genus
390988
1

1
species
node3588.members.0.js
390989

genus
17
1865

3
196914
species

3
1246995
strain
node3591.members.0.js

species
1867
1

1
strain
node3593.members.0.js
457423

9
2626549
no rank

946334
node3595.members.0.js
species
3

species
node3596.members.0.js
2836373
2

node3597.members.0.js
species
649831
4

species
3
1866

512565
node3599.members.0.js
strain
3

node3600.members.0.js
species
122358
1

genus
673534
1

no rank
2631981
1

1
2024580
species
node3603.members.0.js

1
3
genus
node3604.members.0.js
907364

node3605.members.0.js
species
1076124
2

2
338583
genus

338584
species
node3607.members.0.js
2

genus
1
168694

species
1
168695

369723
strain
node3610.members.0.js
1

35753
1
genus

1
node3612.members.0.js
species
53362

16
3
1873
genus
node3613.members.0.js

species
node3614.members.0.js
515350
2

291594
species
node3615.members.0.js
2

2294034
node3616.members.0.js
species
2

1
species
node3617.members.0.js
47853

1
47865
node3618.members.0.js
species

no rank
2617518
2

species
node3620.members.0.js
2583243
2

node3621.members.0.js
species
356852
1

2
261654
species
node3622.members.0.js

2920401
2
genus

2811111
node3624.members.0.js
species
2

2
53365
genus

2
2645785
no rank

2
node3627.members.0.js
species
2728827

genus
84593
1

no rank
1
2630859

1
2742132
node3630.members.0.js
species

5
593
node3631.members.0.js
order
85007

no rank
2
697024

2
1
741759
node3633.members.0.js
genus

1
node3634.members.0.js
species
2760083

family
10
85026

node3636.members.0.js
genus
2053
2
10

1
4
no rank
node3637.members.0.js
2657482

2059875
species
node3638.members.0.js
1

2
node3639.members.0.js
species
337191

1
1136941
species
node3640.members.0.js

2
species
node3641.members.0.js
84595

2055
node3642.members.0.js
species
1

family
85028
8

genus
2060
8

node3645.members.0.js
species
57704
3

4
species
node3646.members.0.js
47312

1
2633480
no rank

1
node3648.members.0.js
species
2201359

11
2805586
family

1847725
11
genus

species
node3651.members.0.js
1528099
11

11
85029
family

1
11
genus
node3653.members.0.js
37914

node3654.members.0.js
species
139021
2

2
499555
node3655.members.0.js
species

2617939
4
no rank

node3657.members.0.js
species
1408143
1

3
node3658.members.0.js
species
712270

2
546160
species
node3659.members.0.js

family
1
316606

1
286801
genus

1
node3662.members.0.js
species
286802

9
115
family
node3663.members.0.js
1762

670516
6
genus

404941
species
node3665.members.0.js
1

36809
species
node3666.members.0.js
2

2
3
species
node3667.members.0.js
1774

node3668.members.0.js
subspecies
2480908
1

genus
1073531
5

1788
node3670.members.0.js
species
2

3
species
node3671.members.0.js
29314

56
2
1866885
node3672.members.0.js
genus

species
node3673.members.0.js
1286181
1

3
node3674.members.0.js
species
1795

2
species
node3675.members.0.js
1431246

1
species
node3676.members.0.js
1791

2
species
node3677.members.0.js
1286180

1
species
node3678.members.0.js
212765

1
node3679.members.0.js
species
1802

1249101
species
node3680.members.0.js
2

1
species
node3681.members.0.js
53462

node3682.members.0.js
no rank
2636767
1

2
126673
species
node3683.members.0.js

39691
node3684.members.0.js
species
1

1
1797
node3685.members.0.js
species

39687
species
node3686.members.0.js
4

1
node3687.members.0.js
species
2761578

node3688.members.0.js
species
258505
1

1792
node3689.members.0.js
species
1

6
36814
species

710685
node3691.members.0.js
strain
6

node3692.members.0.js
species
1799
1

2
39692
node3693.members.0.js
species

1
1534349
species
node3694.members.0.js

node3695.members.0.js
species
85693
2

2
species
node3696.members.0.js
28047

36813
node3697.members.0.js
species
1

1
1772
species
node3698.members.0.js

species
node3699.members.0.js
1794
1

1810
2
species

1354275
strain
node3701.members.0.js
2

species
node3702.members.0.js
1793
1

319707
node3703.members.0.js
species
4

1804
1
species

strain
node3705.members.0.js
350054
1

2
758802
node3706.members.0.js
species

species
node3707.members.0.js
39694
1

genus
39
1763

1
1778
species
node3709.members.0.js

3
2249310
species group

722731
node3711.members.0.js
species
1

species
node3712.members.0.js
220927
1

1
470076
species
node3713.members.0.js

1552759
node3714.members.0.js
species
1

species
node3715.members.0.js
44010
1

2
node3716.members.0.js
species
2738409

species
node3717.members.0.js
2094119
3

no rank
node3718.members.0.js
2642494
1
12

2587868
node3719.members.0.js
species
1

1
1545728
species
node3720.members.0.js

1
species
node3721.members.0.js
1547487

2
1920667
node3722.members.0.js
species

species
node3723.members.0.js
2675524
3

species
node3724.members.0.js
1682113
3

species
node3725.members.0.js
1775
2

2
1789
species
node3726.members.0.js

species group
120793
4

node3728.members.0.js
species
1767
1

3
species
node3729.members.0.js
64667

1
species
node3730.members.0.js
1780

species
node3731.members.0.js
482462
2

species
node3732.members.0.js
152142
4

1
1809
node3733.members.0.js
species

1653
family
node3734.members.0.js
370
13

355
36
1716
genus
node3735.members.0.js

3
node3736.members.0.js
species
2763010

2
43765
node3737.members.0.js
species

4
2624378
no rank

702967
node3739.members.0.js
species
2

2778078
species
node3740.members.0.js
2

node3741.members.0.js
species
108486
3

4
node3742.members.0.js
species
2079234

9
43769
species
node3743.members.0.js

2
species
node3744.members.0.js
161899

node3745.members.0.js
species
1862358
1

136857
node3746.members.0.js
species
2

2
2079535
species
node3747.members.0.js

2488819
node3748.members.0.js
species
1

2
1223514
species

1223515
strain
node3750.members.0.js
2

1
1072256
node3751.members.0.js
species

node3752.members.0.js
species
38305
3
6

strain
node3753.members.0.js
1224164
3

species
node3754.members.0.js
1697
1

156978
species
node3755.members.0.js
27

441500
species
node3756.members.0.js
1

2
node3757.members.0.js
species
39791

species
node3758.members.0.js
258224
2

1979527
species
node3759.members.0.js
3

1
2
species
node3760.members.0.js
575200

1
node3761.members.0.js
strain
1224163

2
node3762.members.0.js
species
187491

species
1
65058

1
node3764.members.0.js
strain
1408268

4
node3765.members.0.js
species
1718

2594913
species
node3766.members.0.js
24

156976
species
node3767.members.0.js
23

1
species
node3768.members.0.js
43770

13
node3769.members.0.js
species
161879

15
53374
species
node3770.members.0.js

species
node3771.members.0.js
38289
25
27

2
306537
node3772.members.0.js
strain

38301
node3773.members.0.js
species
1

169292
node3774.members.0.js
species
22

8
node3775.members.0.js
species
161896

14
2735136
node3776.members.0.js
species

node3777.members.0.js
species
43990
3

5
node3778.members.0.js
species
322009

2
species
node3779.members.0.js
571915

node3780.members.0.js
species
28028
2

node3781.members.0.js
species
38304
16

2754725
node3782.members.0.js
species
1

node3783.members.0.js
species
1705
1

4
species
node3784.members.0.js
2675216

species
1
152794

196164
node3786.members.0.js
strain
1

13
species
node3787.members.0.js
134034

species
1
42817

strain
node3789.members.0.js
1348662
1

13
species
node3790.members.0.js
1725

1727
1
species

858619
node3792.members.0.js
strain
1

7
species
node3793.members.0.js
43768

13
401472
species
node3794.members.0.js

species
node3795.members.0.js
38302
1

no rank
2
42818

2
1164002
node3797.members.0.js
species

family
85025
60

6
36
genus
node3799.members.0.js
1827

1
species
node3800.members.0.js
38310

species group
1
2840174

334542
node3802.members.0.js
species
1

1
species
node3803.members.0.js
1828

1
21
no rank
node3804.members.0.js
192944

2806442
species
node3805.members.0.js
1

1
2567884
species
node3806.members.0.js

1302308
species
node3807.members.0.js
2

1
species
node3808.members.0.js
2742603

node3809.members.0.js
species
1653478
9

2499145
species
node3810.members.0.js
2

2507582
node3811.members.0.js
species
4

3
99653
node3812.members.0.js
species

1
132919
species

node3814.members.0.js
strain
101510
1

1
1829
species
node3815.members.0.js

37919
1
species

1
strain
node3817.members.0.js
543736

1
24
genus
node3818.members.0.js
1817

5
2749991
species
node3819.members.0.js

species
node3820.members.0.js
209247
1

1
3
species
node3821.members.0.js
37326

2
node3822.members.0.js
strain
1133849

204891
node3823.members.0.js
species
1

3
species
node3824.members.0.js
480035

node3825.members.0.js
species
455432
4

1823
node3826.members.0.js
species
1

1
37332
species
node3827.members.0.js

node3828.members.0.js
species
2382165
1

node3829.members.0.js
species
228602
3

order
85011
222

2062
node3831.members.0.js
family
222
3

1883
genus
node3832.members.0.js
215
50

1
node3833.members.0.js
species
2686304

species
node3834.members.0.js
1940
2

1
node3835.members.0.js
species
1893

2
1915
species
node3836.members.0.js

1
36818
node3837.members.0.js
species

species
node3838.members.0.js
173860
1

1971
1
species

316284
node3840.members.0.js
strain
1

1890
node3841.members.0.js
species
1

1
species
node3842.members.0.js
67267

species
2
68246

284034
subspecies
node3844.members.0.js
2

1
2710756
species
node3845.members.0.js

1
1901
node3846.members.0.js
species

2768068
species
node3847.members.0.js
5

1
45398
node3848.members.0.js
species

7
54571
node3849.members.0.js
species

1
2867120
species group

1887
species
node3851.members.0.js
1

species
node3852.members.0.js
67345
2

2599401
node3853.members.0.js
species
1

1
1854574
node3854.members.0.js
species

node3855.members.0.js
species
28894
1

2
species
node3856.members.0.js
1616117

2593676
node3857.members.0.js
no rank
68
6

species
node3858.members.0.js
2906474
1

1
2201357
species
node3859.members.0.js

2420135
node3860.members.0.js
species
1

1
node3861.members.0.js
species
2838850

node3862.members.0.js
species
1262452
10

species
node3863.members.0.js
2692234
2

1
node3864.members.0.js
species
1984801

862751
species
node3865.members.0.js
1

2
2721246
node3866.members.0.js
species

1
node3867.members.0.js
species
2306165

1
node3868.members.0.js
species
2582831

species
node3869.members.0.js
2742134
1

node3870.members.0.js
species
1495638
1

1
2585716
species
node3871.members.0.js

2730915
species
node3872.members.0.js
1

1
node3873.members.0.js
species
659352

1
1882757
node3874.members.0.js
species

1
species
node3875.members.0.js
2797167

2
node3876.members.0.js
species
1577075

species
node3877.members.0.js
2825844
1

1
465541
species
node3878.members.0.js

1661694
species
node3879.members.0.js
3

1
2203210
node3880.members.0.js
species

1
2684468
species
node3881.members.0.js

1841249
node3882.members.0.js
species
1

1264596
species
node3883.members.0.js
1

1
species
node3884.members.0.js
2607753

node3885.members.0.js
species
2880933
2

2742137
node3886.members.0.js
species
1

node3887.members.0.js
species
2078691
1

species
node3888.members.0.js
2695266
1

node3889.members.0.js
species
1488356
2

1
species
node3890.members.0.js
2684469

species
node3891.members.0.js
2742133
1

1
1476754
species
node3892.members.0.js

1
species
node3893.members.0.js
2136173

species
node3894.members.0.js
2838015
1

2059884
species
node3895.members.0.js
2

1
species
node3896.members.0.js
2840373

species
node3897.members.0.js
2072505
1

1
2487421
node3898.members.0.js
species

1851167
node3899.members.0.js
species
2

1
node3900.members.0.js
species
1751294

1
node3901.members.0.js
species
1848900

species
node3902.members.0.js
1690221
1

2
1355015
species
node3903.members.0.js

68249
node3904.members.0.js
species
2

5
629295
species group

1482596
3
species subgroup

3
1911
species

subspecies
3
67263

node3909.members.0.js
strain
455632
3

1482558
1
species subgroup

species
1908
1

1
strain
node3912.members.0.js
1172567

1482561
1
species subgroup

1
species
node3914.members.0.js
1892

node3915.members.0.js
species
285578
1

species
node3916.members.0.js
1907
3

species
1914
1

1
node3918.members.0.js
subspecies
58340

67296
species
node3919.members.0.js
1

1
2838335
species group

2184053
species
node3921.members.0.js
1

species
node3922.members.0.js
553510
3

node3923.members.0.js
species
2768069
1

1967
species
node3924.members.0.js
1

1
species
node3925.members.0.js
42881

node3926.members.0.js
species
2763006
1

node3927.members.0.js
species
80860
7

1
1977088
node3928.members.0.js
species

1
species
node3929.members.0.js
665007

1
33899
node3930.members.0.js
species

species group
2839105
2

1
1912
node3932.members.0.js
species

1
node3933.members.0.js
species
576784

species
node3934.members.0.js
1176198
1

species
node3935.members.0.js
68214
3

1
species
node3936.members.0.js
146923

species
node3937.members.0.js
2049881
1

1
species
node3938.members.0.js
1969

68175
node3939.members.0.js
species
1

node3940.members.0.js
species
362257
1

1
species
node3941.members.0.js
1442032

4
species
node3942.members.0.js
66871

node3943.members.0.js
species
1935
1

1
48665
species
node3944.members.0.js

1
node3945.members.0.js
species
47763

1927
node3946.members.0.js
species
1

1
83656
node3947.members.0.js
species

species
114687
1

1160718
node3949.members.0.js
strain
1

species group
2
2849069

species
node3951.members.0.js
53451
2

1
2717324
node3952.members.0.js
species

68270
node3953.members.0.js
species
1

1
node3954.members.0.js
species
146922

genus
4
2063

2633591
4
no rank

2018025
species
node3957.members.0.js
2

node3958.members.0.js
species
2742135
2

order
2037
47

2049
family
node3960.members.0.js
46
1

5
29
node3961.members.0.js
genus
1654

1
species
node3962.members.0.js
2321394

2744574
species
node3963.members.0.js
2

1
2057743
species
node3964.members.0.js

1
2763540
species
node3965.members.0.js

1
species
node3966.members.0.js
2057800

species
node3967.members.0.js
1852377
1

4
544580
species
node3968.members.0.js

1
1659
species
node3969.members.0.js

1
1655
node3970.members.0.js
species

1
9
node3971.members.0.js
no rank
2609248

1
species
node3972.members.0.js
712122

species
706438
2

2
706439
node3974.members.0.js
strain

1
species
node3975.members.0.js
2755559

node3976.members.0.js
species
712116
2

2
649739
species

2
649743
strain
node3978.members.0.js

2
node3979.members.0.js
species
1656

genus
node3980.members.0.js
1522056
1

genus
2529408
8

2
52773
species
node3982.members.0.js

5
1660
node3983.members.0.js
species

species
node3984.members.0.js
131110
1

2
1
28263
node3985.members.0.js
genus

1302235
species
node3986.members.0.js
1

2
2888879
genus

2
2692125
node3988.members.0.js
species

genus
2
2050

2
node3990.members.0.js
species
2051

genus
1069494
1

1
species
node3992.members.0.js
2733571

no rank
1
56769

1
node3994.members.0.js
species
239730

order
1217098
2

1217100
2
family

genus
2
281472

species
node3998.members.0.js
419479
2

order
node3999.members.0.js
85006
12
538

33
4
85020
family
node4000.members.0.js

genus
1
1161125

1
521392
species
node4002.members.0.js

36739
genus
node4003.members.0.js
7
5

2
species
node4004.members.0.js
1667168

node4005.members.0.js
genus
43668
13
21

node4006.members.0.js
species
2017484
1

43669
node4007.members.0.js
species
1

2623841
6
no rank

2571029
node4009.members.0.js
species
1

node4010.members.0.js
species
1903186
2

2887346
node4011.members.0.js
species
3

family
2
145357

genus
1
908935

no rank
2649892
1

node4015.members.0.js
species
2770551
1

genus
745364
1

1
571913
species
node4017.members.0.js

85021
33
family

genus
367298
1

no rank
2637926
1

1
node4021.members.0.js
species
2714941

3
267408
genus

no rank
2663846
3

3
1658671
species
node4024.members.0.js

53357
5
genus

5
53358
node4026.members.0.js
species

3
23
genus
node4027.members.0.js
53457

4
262209
species
node4028.members.0.js

10
53458
species
node4029.members.0.js

5
857417
node4030.members.0.js
species

2649294
1
no rank

node4032.members.0.js
species
2761047
1

genus
1
99479

2643346
1
no rank

1813880
species
node4035.members.0.js
1

14
2805590
family

genus
265976
7

1
767452
species
node4038.members.0.js

1078471
species
node4039.members.0.js
6

genus
node4040.members.0.js
125287
2
7

1
2615080
no rank

2508882
species
node4042.members.0.js
1

1
2594265
node4043.members.0.js
species

1
1288636
species
node4044.members.0.js

2
node4045.members.0.js
species
2283195

5
189
node4046.members.0.js
family
1268

genus
1
1160973

1
2634694
no rank

1
node4049.members.0.js
species
2170745

7
25
genus
node4050.members.0.js
57493

2
species
node4051.members.0.js
71999

446860
node4052.members.0.js
species
1

1275
node4053.members.0.js
species
3

1049583
node4054.members.0.js
species
5

7
72000
node4055.members.0.js
species

25
68
genus
node4056.members.0.js
1269

no rank
2620948
11

2
1811305
node4058.members.0.js
species

node4059.members.0.js
species
2856555
9

31
species
node4060.members.0.js
1270

1
species
node4061.members.0.js
1273

35
3
32207
genus
node4062.members.0.js

12
13
species
node4063.members.0.js
2047

762948
node4064.members.0.js
strain
1

396015
species
node4065.members.0.js
1

43675
node4066.members.0.js
species
11

6
172042
node4067.members.0.js
species

37923
species
node4068.members.0.js
1

14
169133
genus

no rank
2632435
14

14
node4071.members.0.js
species
2567881

node4072.members.0.js
genus
1663
3
29

1
1670
species
node4073.members.0.js

node4074.members.0.js
no rank
235627
3
20

1811254
node4075.members.0.js
species
1

1
904039
node4076.members.0.js
species

1
species
node4077.members.0.js
2830990

species
node4078.members.0.js
1571833
3

2
2830992
node4079.members.0.js
species

1588023
species
node4080.members.0.js
1

1
species
node4081.members.0.js
1704044

1
2830983
node4082.members.0.js
species

species
node4083.members.0.js
2575374
1

species
node4084.members.0.js
2830997
1

2
node4085.members.0.js
species
2079227

1477518
species
node4086.members.0.js
1

node4087.members.0.js
species
2020130
1

1
37928
node4088.members.0.js
species

species
node4089.members.0.js
156980
3

species
node4090.members.0.js
656366
1

genus
596707
1

2646595
1
no rank

1
2020377
node4093.members.0.js
species

genus
4
1742993

1
node4095.members.0.js
species
728066

no rank
node4096.members.0.js
2647000
1
3

species
node4097.members.0.js
2590774
2

genus
1742989
7

node4099.members.0.js
species
37930
1

4
162496
species
node4100.members.0.js

256701
2
species

2
861360
node4102.members.0.js
strain

family
85019
23

genus
node4104.members.0.js
1696
3
23

273384
species
node4105.members.0.js
1

1
species
node4106.members.0.js
2697565

3
node4107.members.0.js
species
1136497

no rank
2614124
8

2725563
species
node4109.members.0.js
1

species
node4110.members.0.js
2575923
6

node4111.members.0.js
species
2777556
1

node4112.members.0.js
species
1703
1

1
species
node4113.members.0.js
358099

3
199591
species
node4114.members.0.js

node4115.members.0.js
species
33889
2

157
15
85023
node4116.members.0.js
family

427753
1
genus

no rank
1
2635918

1
species
node4119.members.0.js
2282656

genus
node4120.members.0.js
33886
4
16

3
110937
species

3
strain
node4122.members.0.js
1328866

no rank
node4123.members.0.js
2609250
4
8

node4124.members.0.js
species
2609258
1

2609254
node4125.members.0.js
species
3

1
145458
species
node4126.members.0.js

9
4
1573
node4127.members.0.js
genus

3
5
node4128.members.0.js
species
28447

2
node4129.members.0.js
subspecies
1874630

1
node4130.members.0.js
genus
46352

genus
4
235888

2632331
3
no rank

2603292
node4133.members.0.js
species
3

species
node4134.members.0.js
2079791
1

genus
2680004
3

no rank
1
2680006

2759943
node4137.members.0.js
species
1

2
species
node4138.members.0.js
2419774

518733
4
genus

412690
species
node4140.members.0.js
4

2034
genus
node4141.members.0.js
13
1

species
2035
1

node4143.members.0.js
no rank
138532
1

257496
node4144.members.0.js
no rank
8
2

2070337
node4145.members.0.js
species
1

2
2841594
species
node4146.members.0.js

1
2795488
node4147.members.0.js
species

2
node4148.members.0.js
species
1561023

3
69373
node4149.members.0.js
species

genus
33877
7

1
589382
species
node4151.members.0.js

1
2592652
node4152.members.0.js
species

1
species
node4153.members.0.js
2080742

2639701
4
no rank

2
2498704
species
node4155.members.0.js

2
species
node4156.members.0.js
2781962

genus
1
1705353

species
node4158.members.0.js
708131
1

190323
2
genus

species
node4160.members.0.js
150123
1

no rank
1
2624265

1
species
node4162.members.0.js
2480625

genus
3
110932

node4164.members.0.js
no rank
2663824
1
3

1
2781978
species
node4165.members.0.js

node4166.members.0.js
species
2724914
1

node4167.members.0.js
genus
33882
15
49

3
species
node4168.members.0.js
82380

1
species
node4169.members.0.js
104336

1
2614638
node4170.members.0.js
species

species
node4171.members.0.js
162426
3

node4172.members.0.js
species
300019
1

2782167
species
node4173.members.0.js
2

936337
node4174.members.0.js
species
1

2
node4175.members.0.js
species
904291

2609290
node4176.members.0.js
no rank
16
7

2763257
species
node4177.members.0.js
1

2483401
species
node4178.members.0.js
2

1
node4179.members.0.js
species
1916917

1
2782166
node4180.members.0.js
species

node4181.members.0.js
species
2489212
1

2
2606451
species
node4182.members.0.js

2603598
species
node4183.members.0.js
1

2
1072463
species
node4184.members.0.js

node4185.members.0.js
species
743009
1

node4186.members.0.js
species
2614639
1

255204
1
genus

2615177
1
no rank

node4189.members.0.js
species
2866588
1

genus
1
110934

2641148
1
no rank

1
species
node4192.members.0.js
2773266

1649454
1
genus

no rank
1
2645362

1
species
node4195.members.0.js
2596912

9
96492
genus

no rank
9
2627005

node4198.members.0.js
species
2596916
9

881616
1
genus

no rank
2618217
1

2735133
species
node4201.members.0.js
1

120212
2
genus

2648727
2
no rank

2895559
node4204.members.0.js
species
2

genus
337004
1

279828
species
node4206.members.0.js
1

1
1655488
no rank

clade
1655489
1

529881
1
genus

no rank
2617988
1

1
species
node4211.members.0.js
1855377

447237
3
genus

3
2626248
no rank

3
species
node4214.members.0.js
1795630

no rank
90316
2

2
node4216.members.0.js
species
2880260

55968
7
genus

1
2621730
no rank

node4219.members.0.js
species
2813744
1

1
node4220.members.0.js
species
2714935

1935379
species
node4221.members.0.js
5

85022
7
family

genus
43673
7

species
node4224.members.0.js
43674
7

family
28
2805426

1
28
node4226.members.0.js
genus
57499

27
node4227.members.0.js
species
1276

1331736
3
family

node4229.members.0.js
genus
626119
1
3

1
2675754
species
node4230.members.0.js

2648503
1
no rank

2897774
node4232.members.0.js
species
1

family
125316
1

1
84756
genus

1
84757
species

1
node4236.members.0.js
strain
471853

family
node4237.members.0.js
85017
1
4

157920
3
genus

no rank
3
2624466

3
species
node4240.members.0.js
2587808

29
85016
family

7
665568
genus

7
node4243.members.0.js
species
545619

genus
node4244.members.0.js
1707
2
15

species
1
11

1
node4246.members.0.js
strain
593907

2566013
node4247.members.0.js
species
1

2
11
node4248.members.0.js
no rank
2620175

1
species
node4249.members.0.js
2704467

3
species
node4250.members.0.js
2591145

species
node4251.members.0.js
2816956
1

node4252.members.0.js
species
2819979
3

node4253.members.0.js
species
2871170
1

7
162491
genus

2619021
7
no rank

1179673
species
node4256.members.0.js
7

family
2
145358

genus
154116
2

2
species
node4259.members.0.js
2585135

family
1
2805591

2038
1
genus

2039
node4262.members.0.js
species
1

no rank
35766
3

3
species
node4264.members.0.js
152507

class
12
84992

order
84993
12

family
7
633392

genus
2789775
1

node4269.members.0.js
species
2789776
1

genus
4
1648491

no rank
4
2633173

node4272.members.0.js
species
2849779
1

2884263
node4273.members.0.js
species
3

genus
467975
2

2624035
2
no rank

node4276.members.0.js
species
2722752
2

4
2448023
family

4
682522
genus

species
4
467094

4
node4280.members.0.js
strain
1313172

310070
1
no rank

1
species
node4282.members.0.js
310071

1
1752188
no rank

species
node4284.members.0.js
2900548
1

class
8
84995

order
8
84996

family
1
2600303

2600304
1
genus

496014
species
node4289.members.0.js
1

family
84997
7

42255
7
genus

2
1
49319
species
node4292.members.0.js

1
266117
strain
node4293.members.0.js

2653852
node4294.members.0.js
species
5

class
17
1497346

17
588673
order

family
320583
1

1
191494
genus

1
2627773
no rank

1
species
node4300.members.0.js
2812560

11
361606
family

11
361607
genus

11
1097667
node4303.members.0.js
species

no rank
1363568
5

2884022
species
node4305.members.0.js
5

class
3
908620

1755823
1
order

1
1755824
family

genus
1
1755825

1
species
node4310.members.0.js
1670830

2
908621
order

908622
2
family

908623
2
genus

2
1608957
species
node4314.members.0.js

1
46
node4315.members.0.js
phylum
200795

1495646
5
class

order
1495647
5

5
1495648
family

5
1495649
genus

node4320.members.0.js
species
1495650
5

1
475962
class

1
475963
order

475964
1
family

genus
1
233191

species
133453
1

1
strain
node4326.members.0.js
926550

189774
5
no rank

2026724
node4328.members.0.js
species
5

class
8
32061

8
32064
order

suborder
1508594
8

family
1106
8

genus
1107
8

species
152260
5

326427
strain
node4335.members.0.js
5

3
1
1108
species
node4336.members.0.js

2
node4337.members.0.js
strain
324602

class
1
2682225

order
1
2682226

1
2682227
family

genus
2682228
1

1
species
node4342.members.0.js
2601677

301297
3
class

1
1202465
order

family
1202464
1

1
node4346.members.0.js
genus
61434

genus
node4347.members.0.js
670486
1
2

2638108
1
no rank

1
species
node4349.members.0.js
943347

292625
13
class

3
292629
order

family
2
292628

2
233189
genus

167964
2
species

2
node4355.members.0.js
strain
926569

no rank
1674871
1

1
species
node4357.members.0.js
2073117

no rank
9
1081798

9
species
node4359.members.0.js
2052143

1
452434
no rank

452435
node4361.members.0.js
species
1

no rank
58229
1

node4363.members.0.js
species
166587
1

388447
8
class

3
768667
order

family
768668
3

genus
768669
3

2045280
species
node4368.members.0.js
3

388448
5
order

2692416
1
family

2692417
1
genus

1
node4372.members.0.js
species
2509675

4
768649
family

genus
768650
4

no rank
2625087
4

node4376.members.0.js
species
2490863
4

node4377.members.0.js
phylum
1239
7
1203

1
47928
no rank

344338
species
node4379.members.0.js
1

159
1737404
class

no rank
1737407
9

1582879
9
genus

1852374
node4383.members.0.js
species
9

1737405
150
order

1570339
node4385.members.0.js
family
148
1

genus
48
162289

2
2637196
no rank

species
node4388.members.0.js
1912856
2

9
54006
species
node4389.members.0.js

37
node4390.members.0.js
species
54005

2
543311
genus

33033
species
node4392.members.0.js
2

genus
23
150022

23
1260
node4394.members.0.js
species

genus
node4395.members.0.js
165779
7
73

6
1287640
node4396.members.0.js
species

33037
node4397.members.0.js
species
2

1870984
species
node4398.members.0.js
31

24
27
species
node4399.members.0.js
33034

3
525919
node4400.members.0.js
strain

31983
1
genus

1
node4402.members.0.js
species
40091

2
1737406
family

2794837
2
genus

2
45497
species

2
node4406.members.0.js
strain
1288971

class
526524
6

6
526525
order

1
2810281
family

191303
1
genus

no rank
1
2638206

species
node4412.members.0.js
1712675
1

family
5
128827

genus
1
1505663

1522
species
node4415.members.0.js
1

no rank
1
544447

node4417.members.0.js
species
2676062
1

genus
node4418.members.0.js
1647
1

genus
2
1918536

1796635
species
node4420.members.0.js
2

84086
1
no rank

2707344
species
node4422.members.0.js
1

class
227
186801

68295
7
order

family
1
186814

42857
1
no rank

1
44260
genus

species
node4428.members.0.js
1525
1

543371
6
family

44000
genus
node4430.members.0.js
5
2

1
2806205
node4431.members.0.js
species

2
301953
species

node4433.members.0.js
strain
632335
2

1
28895
genus

1
29329
species

1
858215
node4436.members.0.js
strain

3
2770089
order

family
3
2770092

genus
2770094
3

3
2316383
node4440.members.0.js
species

3
213
node4441.members.0.js
order
186802

family
186804
33

2743582
1
genus

node4444.members.0.js
species
89152
1

1
1501226
genus

no rank
2626894
1

1
2724150
node4447.members.0.js
species

no rank
1411023
15

15
2081703
species
node4449.members.0.js

1849828
1
genus

node4451.members.0.js
species
1505
1

13
1257
genus

13
node4453.members.0.js
species
1261

genus
44259
2

species
2
143361

2
546269
node4456.members.0.js
strain

family
2603322
1

genus
1
2603323

node4459.members.0.js
species
2173034
1

family
2848916
2

2
171003
genus

2
116090
node4462.members.0.js
species

family
node4463.members.0.js
186803
3
31

186928
node4464.members.0.js
no rank
6
1

species
node4465.members.0.js
2785025
1

1
node4466.members.0.js
species
712991

1898203
species
node4467.members.0.js
1

2109691
node4468.members.0.js
species
1

node4469.members.0.js
species
712982
1

1
2719231
genus

1
29370
species

strain
node4472.members.0.js
1297793
1

genus
1
572511

species
node4474.members.0.js
2779518
1

1164882
2
genus

617123
species
node4476.members.0.js
2

2569097
1
genus

1
39488
node4478.members.0.js
species

genus
1506553
4

species
66219
2

node4481.members.0.js
strain
357809
2

node4482.members.0.js
species
29347
2

830
1
genus

species
1
43305

1
strain
node4485.members.0.js
515622

1
genus
node4486.members.0.js
207244

genus
2
2316020

species
1
33039

1
strain
node4489.members.0.js
657313

1
33038
species

411470
strain
node4491.members.0.js
1

genus
1
698776

29360
1
species

1
strain
node4494.members.0.js
642492

genus
5
1843210

4
1727145
node4496.members.0.js
species

2696063
species
node4497.members.0.js
1

2
33042
genus

2
116085
species

2
717962
node4500.members.0.js
strain

genus
1663717
1

1
1679721
species
node4502.members.0.js

2
28
family
node4503.members.0.js
216572

genus
1
459786

no rank
1
2629304

species
node4506.members.0.js
2763056
1

253238
1
genus

1
253239
species
node4508.members.0.js

946234
2
genus

2
292800
species
node4510.members.0.js

2
1637257
genus

2
884684
species

699246
strain
node4513.members.0.js
2

3
216851
genus

3
species
node4515.members.0.js
853

genus
2304691
2

2
1
1510
node4517.members.0.js
species

1
160385
subspecies

1
node4519.members.0.js
strain
1346611

2
2304693
genus

2
node4521.members.0.js
species
84032

1
2
node4522.members.0.js
genus
1263

1
species
node4523.members.0.js
1160721

473772
5
no rank

2
2485925
species
node4525.members.0.js

3
2799561
node4526.members.0.js
species

genus
1
2842531

1
2780922
species
node4528.members.0.js

genus
5
236752

236753
node4530.members.0.js
species
5

family
543349
1

genus
2733
1

species
1
2734

292459
node4534.members.0.js
strain
1

31984
5
family

genus
2831443
5

1
5
node4537.members.0.js
species
35701

strain
node4538.members.0.js
498761
4

990719
4
family

990721
4
genus

no rank
4
2649046

species
node4542.members.0.js
1935934
2

2
node4543.members.0.js
species
2841518

family
66
31979

1
114627
genus

species
1
208226

1
293826
strain
node4547.members.0.js

4
1769729
genus

4
1498
species
node4549.members.0.js

genus
15
1848399

species
node4551.members.0.js
1286698
15

1849822
1
genus

1
1490
node4553.members.0.js
species

44
1485
genus

2068654
node4555.members.0.js
species
1

2320868
species
node4556.members.0.js
1

59619
1
no rank

1
59620
node4558.members.0.js
species

2
node4559.members.0.js
species
36845

94869
node4560.members.0.js
species
1

394958
species
node4561.members.0.js
1

1
1216932
species
node4562.members.0.js

1491
node4563.members.0.js
species
2

1520
node4564.members.0.js
species
1

1534
species
node4565.members.0.js
1

1
species
node4566.members.0.js
1561

species
1
217159

1
strain
node4568.members.0.js
536227

1502
node4569.members.0.js
species
2

20
node4570.members.0.js
species
1529

1
node4571.members.0.js
species
1497

1
169679
node4572.members.0.js
species

2614128
4
no rank

1
node4574.members.0.js
species
1970093

1
node4575.members.0.js
species
1042156

2507159
node4576.members.0.js
species
1

1
1506
node4577.members.0.js
species

2
1493
species

node4579.members.0.js
strain
573061
2

49082
1
genus

no rank
1
2638829

1
1041504
node4582.members.0.js
species

family
9
186806

genus
1730
8

no rank
2624479
7

7
2764325
node4586.members.0.js
species

1
1736
node4587.members.0.js
species

1
33951
genus

2638182
1
no rank

1
species
node4590.members.0.js
2184575

no rank
538999
12

543314
node4592.members.0.js
family
9
1

species
1
76124

1
888721
strain
node4594.members.0.js

node4595.members.0.js
genus
86331
3
4

node4596.members.0.js
species
114527
1

species
1
143393

1
strain
node4598.members.0.js
888727

genus
2060094
2

2
node4600.members.0.js
species
2507160

2
539000
family

2
28033
genus

1
338644
node4603.members.0.js
species

1
node4604.members.0.js
species
53633

2717089
1
genus

2086584
species
node4606.members.0.js
1

1491775
1
family

2838170
1
genus

1
2731377
node4609.members.0.js
species

family
186807
14

51196
1
genus

species
51197
1

1
645991
strain
node4613.members.0.js

1
79206
genus

1
885581
species

646529
strain
node4616.members.0.js
1

2916693
1
genus

species
1
59610

1
349161
node4619.members.0.js
strain

genus
36853
4

species
3
142877

node4622.members.0.js
strain
871968
3

1
36854
species

node4624.members.0.js
strain
756499
1

genus
1562
1

species
1565
1

node4627.members.0.js
strain
868595
1

2282742
4
genus

species
58138
4

strain
node4630.members.0.js
485916
4

genus
2
2740

2
species
node4632.members.0.js
2741

39779
node4633.members.0.js
no rank
3
1

2
1898207
node4634.members.0.js
species

order
53433
4

2
1
972
node4636.members.0.js
family

1
32636
genus

31909
1
species

1
373903
strain
node4639.members.0.js

family
2
53434

2
42417
genus

species
42422
2

node4643.members.0.js
strain
748449
2

57
909932
class

1843489
37
order

37
31977
family

29465
node4647.members.0.js
genus
27
7

29466
species
node4648.members.0.js
2

no rank
2630086
2

node4650.members.0.js
species
2682455
2

12
39777
species
node4651.members.0.js

1
node4652.members.0.js
species
39778

3
species
node4653.members.0.js
248315

5
2
906
node4654.members.0.js
genus

1
node4655.members.0.js
species
2144175

species
node4656.members.0.js
907
2

5
2
39948
node4657.members.0.js
genus

2
487173
species
node4658.members.0.js

1
39950
species
node4659.members.0.js

order
1843488
1

family
909930
1

genus
33024
1

no rank
2639160
1

1
species
node4664.members.0.js
2823317

order
19
909929

17
1843490
family

genus
1
365348

2629460
1
no rank

1
node4669.members.0.js
species
484770

16
2375
genus

16
2377
species
node4671.members.0.js

2
1843491
family

genus
2
970

species
69823
2

2
546271
node4675.members.0.js
strain

745
14
91061
class
node4676.members.0.js

2
293
order
node4677.members.0.js
186826

24
186828
family

genus
3
1470540

1
2621505
no rank

1
node4681.members.0.js
species
2908211

1868794
node4682.members.0.js
species
2

genus
117563
4

137732
species
node4684.members.0.js
2

2
species
node4685.members.0.js
46124

2747
8
genus

1
2748
node4687.members.0.js
species

species
node4688.members.0.js
2751
7

9
29393
genus

9
29394
species
node4690.members.0.js

33958
family
node4691.members.0.js
96
5

genus
1
2767881

53444
species
node4693.members.0.js
1

2767878
1
genus

node4695.members.0.js
species
1303590
1

1
46254
genus

species
node4697.members.0.js
2203724
1

genus
node4698.members.0.js
1253
1

genus
node4699.members.0.js
1578
2
36

species
node4700.members.0.js
47770
29

1
species
node4701.members.0.js
33959

1
83683
node4702.members.0.js
species

1
node4703.members.0.js
species
109790

1587
species
node4704.members.0.js
2

2767885
1
genus

1
1599
species
node4706.members.0.js

node4707.members.0.js
genus
2767842
3
5

2
1590
species
node4708.members.0.js

8
4
46255
genus
node4709.members.0.js

1
2506420
species
node4710.members.0.js

1
137591
species
node4711.members.0.js

2
1583
species
node4712.members.0.js

5
2
2742598
node4713.members.0.js
genus

1
97478
node4714.members.0.js
species

species
node4715.members.0.js
1633
2

2
2767892
genus

2
240427
species
node4717.members.0.js

11
3
1243
genus
node4718.members.0.js

5
8
species
node4719.members.0.js
1244

3
node4720.members.0.js
subspecies
115778

17
1
2767879
node4721.members.0.js
genus

1
1074467
node4722.members.0.js
species

1
1612
species

1
936140
strain
node4724.members.0.js

1847728
species
node4725.members.0.js
2

6
938155
species
node4726.members.0.js

5
node4727.members.0.js
species
83526

species
node4728.members.0.js
392416
1

genus
1
2767890

1
node4730.members.0.js
species
1296540

1
2759736
genus
node4731.members.0.js

family
16
81852

node4733.members.0.js
genus
1350
6
14

1
2608891
no rank

1
species
node4735.members.0.js
2057791

1351
node4736.members.0.js
species
3

node4737.members.0.js
species
118060
1

1
species
node4738.members.0.js
160453

1
37734
node4739.members.0.js
species

node4740.members.0.js
species
71452
1

genus
2
2737

species
node4742.members.0.js
218144
1

2571750
species
node4743.members.0.js
1

130
1300
family

122
44
1301
genus
node4745.members.0.js

7
1
2608887
no rank
node4746.members.0.js

2
species
node4747.members.0.js
712633

1902136
species
node4748.members.0.js
1

2
node4749.members.0.js
species
2710759

712623
node4750.members.0.js
species
1

1304
species
node4751.members.0.js
1

1
node4752.members.0.js
species
102684

node4753.members.0.js
species
1314
1

68892
node4754.members.0.js
species
1

species
node4755.members.0.js
1305
10
14

4
strain
node4756.members.0.js
388919

28037
species
node4757.members.0.js
6
5

node4758.members.0.js
strain
365659
1

node4759.members.0.js
species
1309
2

1346
species
node4760.members.0.js
2

5
671232
species group

1338
node4762.members.0.js
species
4

1
1328
node4763.members.0.js
species

1318
2
species

2
node4765.members.0.js
strain
1114965

1307
node4766.members.0.js
species
19

species
node4767.members.0.js
1313
1

5
1308
node4768.members.0.js
species

2
species
node4769.members.0.js
1303

node4770.members.0.js
species
1343
1

1111760
node4771.members.0.js
species
1

4
1302
node4772.members.0.js
species

species group
2
119603

species
1334
1

119602
1
subspecies

strain
node4776.members.0.js
1247189
1

1
1336
species
node4777.members.0.js

1
species
node4778.members.0.js
82348

genus
8
1357

node4780.members.0.js
species
1358
4

species
node4781.members.0.js
2419773
1

node4782.members.0.js
species
1359
2

no rank
1
2643510

2592653
species
node4784.members.0.js
1

1
25
family
node4785.members.0.js
186827

46123
6
genus

46125
node4787.members.0.js
species
6

6
4
881649
node4788.members.0.js
no rank

2796928
node4789.members.0.js
species
1

2774330
node4790.members.0.js
species
1

2
6
node4791.members.0.js
genus
1375

128944
node4792.members.0.js
species
2

species
1376
2

866775
node4794.members.0.js
strain
2

genus
6
2689587

node4796.members.0.js
species
2036206
6

438
2
1385
node4797.members.0.js
order

23
168
node4798.members.0.js
family
186817

1276290
3
genus

species
node4800.members.0.js
35841
3

2685905
genus
node4801.members.0.js
5
1

species
node4802.members.0.js
2743000
1

3
species
node4803.members.0.js
2502791

2675231
1
genus

species
node4805.members.0.js
129985
1

1
2785518
genus

1
2785519
no rank

1
2844358
species
node4808.members.0.js

2837506
3
genus

1397
species
node4810.members.0.js
3

1
1221880
genus

no rank
2636677
1

1
species
node4813.members.0.js
2829187

1
2837485
genus

1
species
node4815.members.0.js
1413

74385
2
genus

node4817.members.0.js
species
2663022
2

2
9
genus
node4818.members.0.js
150247

node4819.members.0.js
species
294699
4

2
3
no rank
node4820.members.0.js
2639704

species
node4821.members.0.js
1636720
1

genus
1
1055323

species
node4823.members.0.js
33936
1

2675233
genus
node4824.members.0.js
3
1

2874282
species
node4825.members.0.js
1

1
2675274
no rank

1
species
node4827.members.0.js
2806989

3
2
129337
node4828.members.0.js
genus

1422
1
species

strain
node4830.members.0.js
272567
1

node4831.members.0.js
genus
1329200
1
2

1221500
species
node4832.members.0.js
1

18
2
2800373
node4833.members.0.js
genus

16
15
1404
node4834.members.0.js
species

1452722
node4835.members.0.js
strain
1

1
2817139
genus

1
species
node4837.members.0.js
1398

2837508
1
genus

node4839.members.0.js
species
189381
1

4
2675230
genus

1
node4841.members.0.js
species
1670641

no rank
2
2675268

node4843.members.0.js
species
2901381
2

species
node4844.members.0.js
665099
1

genus
3
2675234

79880
species
node4846.members.0.js
3

5
2651279
genus

2651280
5
no rank

2651284
species
node4849.members.0.js
5

1609627
1
genus

no rank
2773237
1

2866627
node4852.members.0.js
species
1

182709
1
genus

species
182710
1

221109
strain
node4855.members.0.js
1

genus
2675232
2

1
node4857.members.0.js
species
220684

1
1193713
node4858.members.0.js
species

459532
1
genus

386490
node4860.members.0.js
species
1

1
2837511
genus

1
species
node4862.members.0.js
79883

1
1434042
genus

1
632773
node4864.members.0.js
species

6
400634
genus

1
species
node4866.members.0.js
436516

3
node4867.members.0.js
species
2590012

node4868.members.0.js
no rank
2636778
1
2

1
species
node4869.members.0.js
2070463

genus
351195
2

2
node4871.members.0.js
species
1230341

3
2675229
genus

1
2
node4873.members.0.js
species
1478

1
strain
node4874.members.0.js
1349754

1
421767
node4875.members.0.js
species

12
60
node4876.members.0.js
genus
1386

1
1783501
species
node4877.members.0.js

653685
12
species group

species
node4879.members.0.js
96241
1

10
6
1938374
species subgroup
node4880.members.0.js

3
492670
species
node4881.members.0.js

1
1390
species

strain
node4883.members.0.js
1292358
1

1423
node4884.members.0.js
species
1

2672567
species
node4885.members.0.js
2

1
324767
species

1367477
strain
node4887.members.0.js
1

1
1408
species
node4888.members.0.js

10
6
86661
species group
node4889.members.0.js

species
node4890.members.0.js
1428
2
3

29339
1
no rank

714359
node4892.members.0.js
strain
1

1
1405
species
node4893.members.0.js

185979
17
no rank

species
node4895.members.0.js
1565991
1

node4896.members.0.js
species
1409
2

species
node4897.members.0.js
2682541
2

node4898.members.0.js
species
2587848
2

2
node4899.members.0.js
species
666686

1
node4900.members.0.js
species
2795526

2592382
node4901.members.0.js
species
1

1
species
node4902.members.0.js
2666127

node4903.members.0.js
species
1705566
1

1
node4904.members.0.js
species
486398

1
1581038
species
node4905.members.0.js

98228
node4906.members.0.js
species
2

node4907.members.0.js
species
2728853
4

186824
3
family

genus
1274351
2

species
node4910.members.0.js
1380685
1

714067
node4911.members.0.js
species
1

2689589
1
genus

1
node4913.members.0.js
species
2490858

1
186821
family

1
2077
genus

node4916.members.0.js
species
269673
1

186823
1
family

genus
29330
1

1
1450
species
node4919.members.0.js

no rank
16
539002

no rank
4
539742

4
1
33986
genus
node4922.members.0.js

2644629
3
no rank

3
360911
species
node4924.members.0.js

539738
12
no rank

1378
genus
node4926.members.0.js
12
4

species
node4927.members.0.js
1379
8

family
186820
8

genus
6
1637

no rank
2642072
2

2
species
node4931.members.0.js
1844999

1
529731
node4932.members.0.js
species

3
1
1639
node4933.members.0.js
species

strain
node4934.members.0.js
1234142
2

2
2755
genus

2
node4936.members.0.js
species
2756

3
125
family
node4937.members.0.js
90964

45669
2
genus

2
407035
species
node4939.members.0.js

4
87
node4940.members.0.js
genus
1279

4
species
node4941.members.0.js
2282419

species
node4942.members.0.js
1293
1

1
node4943.members.0.js
species
1283

species
node4944.members.0.js
29388
5

3
node4945.members.0.js
species
29378

1
species
node4946.members.0.js
29379

2
node4947.members.0.js
species
170573

1
species
node4948.members.0.js
29385

no rank
2
91994

node4950.members.0.js
species
2813777
1

1
2583989
node4951.members.0.js
species

985762
node4952.members.0.js
species
1

30
node4953.members.0.js
species
1282

node4954.members.0.js
species
1280
4

node4955.members.0.js
species
28035
5

1281
species
node4956.members.0.js
1

18
node4957.members.0.js
species
1290

3
45972
node4958.members.0.js
species

46127
species
node4959.members.0.js
1

14
489909
genus

node4961.members.0.js
species
489910
14

genus
2
2803850

1
species
node4963.members.0.js
1296

1
643214
node4964.members.0.js
species

4
1955413
genus

node4966.members.0.js
species
1817405
4

227979
6
genus

1
453585
no rank

1
453586
species
node4969.members.0.js

2630462
5
no rank

3
946435
node4971.members.0.js
species

2
species
node4972.members.0.js
2708346

7
5
69965
genus
node4973.members.0.js

69966
node4974.members.0.js
species
2

family
23
186818

2
7
node4976.members.0.js
genus
1372

1
1302659
species
node4977.members.0.js

192421
species
node4978.members.0.js
1

node4979.members.0.js
species
1499687
1

161360
2
species

2
node4981.members.0.js
strain
1185653

496496
1
genus

no rank
1
2617942

species
node4984.members.0.js
2779527
1

3
651660
genus

3
417367
node4986.members.0.js
species

genus
8
160795

8
51173
node4988.members.0.js
species

1569
genus
node4989.members.0.js
4
2

no rank
2647733
1

1
node4991.members.0.js
species
2810514

1
node4992.members.0.js
species
298596

186822
91
family

8
74
node4994.members.0.js
genus
44249

1
species
node4995.members.0.js
116718

365617
1
species

strain
node4997.members.0.js
1268072
1

node4998.members.0.js
species
61624
1
3

2
strain
node4999.members.0.js
1036673

species
3
683737

3
strain
node5001.members.0.js
1313296

2509456
species
node5002.members.0.js
2

682956
node5003.members.0.js
species
4

12
node5004.members.0.js
species
562959

node5005.members.0.js
species
169760
1

species
node5006.members.0.js
128574
4

1126833
species
node5007.members.0.js
1

2
2810347
species
node5008.members.0.js

2
79263
species
node5009.members.0.js

3
1
44251
node5010.members.0.js
species

1333534
strain
node5011.members.0.js
2

2
3
node5012.members.0.js
species
1406

1
node5013.members.0.js
strain
886882

no rank
185978
12

species
node5015.members.0.js
715179
1

2777984
node5016.members.0.js
species
1

node5017.members.0.js
species
2023772
1

1
1536775
node5018.members.0.js
species

node5019.members.0.js
species
2748863
4

1
node5020.members.0.js
species
1906272

1536769
node5021.members.0.js
species
1

1
2660554
node5022.members.0.js
species

1
1536770
species
node5023.members.0.js

1619311
node5024.members.0.js
species
3

1
2495582
node5025.members.0.js
species

1
2
node5026.members.0.js
species
1464

1
node5027.members.0.js
subspecies
1477

node5028.members.0.js
species
1178515
1

189426
node5029.members.0.js
species
1

1
node5030.members.0.js
species
375489

species group
1
2044880

1
373687
species
node5032.members.0.js

node5033.members.0.js
species
44250
1

1
2704462
species
node5034.members.0.js

genus
8
456492

2583377
node5036.members.0.js
species
8

329857
2
genus

2507935
species
node5038.members.0.js
1

1
species
node5039.members.0.js
2598458

no rank
85151
1

genus
55079
1

1
1500254
species
node5042.members.0.js

genus
6
55080

1
54913
species
node5044.members.0.js

1
species
node5045.members.0.js
1393

2796470
species
node5046.members.0.js
3

species
node5047.members.0.js
1465
1

544448
19
phylum

no rank
2799557
3

genus
1912503
3

2624196
3
no rank

species
node5052.members.0.js
1541959
3

15
31969
class

order
2085
9

family
2092
9

2093
4
genus

no rank
2683645
1

1
2725994
node5058.members.0.js
species

node5059.members.0.js
species
2098
3

genus
2
2923352

1
2099
species
node5061.members.0.js

29562
species
node5062.members.0.js
1

3
2767358
genus

28903
node5064.members.0.js
species
1

species
node5065.members.0.js
171284
1

29553
node5066.members.0.js
species
1

order
6
186328

family
33925
1

genus
46239
1

node5070.members.0.js
species
225999
1

family
5
2131

5
2132
genus

node5073.members.0.js
species
2134
2

1
2637901
no rank

1
species
node5075.members.0.js
2792084

216935
1
species

node5077.members.0.js
strain
1276246
1

species
node5078.members.0.js
2139
1

713063
1
no rank

1
2725268
node5080.members.0.js
species

phylum
1297
46

188787
46
class

order
43
118964

183710
43
family

node5085.members.0.js
genus
1298
7
43

node5086.members.0.js
species
1768108
5

2623546
5
no rank

2
2737050
species
node5088.members.0.js

3
node5089.members.0.js
species
2080419

node5090.members.0.js
species
1309411
1

species
node5091.members.0.js
980427
6

310783
node5092.members.0.js
species
2

species
node5093.members.0.js
432329
1

68909
node5094.members.0.js
species
2

3
species
node5095.members.0.js
317577

species
node5096.members.0.js
2202254
1

1182568
node5097.members.0.js
species
3

3
1211322
species
node5098.members.0.js

species
node5099.members.0.js
1299
4

order
3
68933

family
188786
3

genus
node5102.members.0.js
270
1

2
65551
genus

2
172827
species

2
node5105.members.0.js
strain
1339250

67819
7
phylum

99283
1
no rank

157466
node5108.members.0.js
species
1

no rank
1042316
2

2004468
species
node5110.members.0.js
2

4
1663419
class

order
4
1663425

family
1663426
4

no rank
2
2682140

2
node5115.members.0.js
species
2682144

genus
2
1005038

1005039
species
node5117.members.0.js
2

clade
1798711
175

1117
node5119.members.0.js
phylum
174
6

order
52604
3

family
1890500
2

genus
2
44474

2645907
2
no rank

node5124.members.0.js
species
118163
2

family
1
1890498

1
102115
genus

102116
1
species

node5128.members.0.js
strain
111780
1

14
1
2881377
node5129.members.0.js
order

2303507
8
family

1240886
1
genus

no rank
1
2627810

species
node5133.members.0.js
2692086
1

1
2055830
genus

species
node5135.members.0.js
2315793
1

node5136.members.0.js
genus
2303528
6

family
1
2881378

genus
146785
1

146786
1
species

1
2016101
node5140.members.0.js
strain

family
1890438
2

genus
47251
2

2
2650499
no rank

1
node5144.members.0.js
species
1115757

1
111781
node5145.members.0.js
species

family
2
1890436

genus
2
1152

2593292
2
no rank

2
82654
species
node5149.members.0.js

order
1890424
35

family
2881426
9

genus
1218
9

1
98167
no rank

1
159733
node5154.members.0.js
species

1
2627481
no rank

1
1501269
species
node5156.members.0.js

7
1219
species

5
node5158.members.0.js
strain
59920

146891
node5159.members.0.js
strain
1

59922
node5160.members.0.js
strain
1

1890426
23
family

genus
node5162.members.0.js
1129
4
23

4
19
no rank
node5163.members.0.js
2626047

1
2741953
node5164.members.0.js
species

3
node5165.members.0.js
species
1280380

node5166.members.0.js
species
1916956
2

1
1400864
node5167.members.0.js
species

node5168.members.0.js
species
1400865
1

1
node5169.members.0.js
species
221352

6
321327
node5170.members.0.js
species

family
2
1890429

genus
155977
2

no rank
2
2631661

2
2740837
node5174.members.0.js
species

1890431
1
family

genus
1
217161

1
1173032
species

1173020
strain
node5178.members.0.js
1

12
1890505
order

family
1
2917811

genus
1
1827277

2739056
1
species

1
strain
node5183.members.0.js
2599943

family
1890528
11

genus
11
54298

11
259956
no rank

11
species
node5187.members.0.js
259957

3
48504
no rank

species
node5189.members.0.js
1211
3

class
1
307596

order
307595
1

1890422
1
family

33071
1
genus

species
1
1416614

1
strain
node5195.members.0.js
1183438

subclass
32
1301283

21
1150
order

family
1
1892255

genus
1
241421

1
241425
species

1173022
node5201.members.0.js
strain
1

1892254
7
family

genus
1
1198

no rank
1
2609805

1
2839084
node5205.members.0.js
species

genus
1155738
2

1155739
2
species

1454205
strain
node5208.members.0.js
2

genus
4
1158

species
118323
4

4
strain
node5211.members.0.js
56110

5
1892251
family

1
63132
genus

no rank
2648775
1

1
species
node5215.members.0.js
1173025

genus
2886347
4

species
4
2886352

strain
node5218.members.0.js
1173027
4

8
1892252
family

1
1492710
genus

species
1
2874213

strain
node5222.members.0.js
2064643
1

54304
1
genus

1160
species
node5224.members.0.js
1

6
44471
genus

species
128152
4

4
1681829
node5227.members.0.js
strain

2
2642155
no rank

2
272133
node5229.members.0.js
species

order
11
1118

5
1
1890464
node5231.members.0.js
family

genus
102231
1

no rank
1
2623012

node5234.members.0.js
species
1173026
1

genus
669357
1

no rank
2625037
1

1
node5237.members.0.js
species
1615909

2
268175
genus

2
2648896
no rank

species
node5240.members.0.js
2005460
2

5
2815910
family

5
102234
genus

2629879
2
no rank

2
node5244.members.0.js
species
2054282

3
102235
species

3
strain
node5246.members.0.js
292563

1890450
1
family

genus
1
28070

species
2546356
1

1
65393
strain
node5250.members.0.js

1161
node5251.members.0.js
order
68
13

family
1
119859

111782
1
genus

2649714
1
no rank

2099387
species
node5255.members.0.js
1

2
1892263
family

genus
2
1190

no rank
2
494603

2005456
species
node5259.members.0.js
1

1
species
node5260.members.0.js
548826

2661849
8
family

1186
8
genus

species
32054
1

1
2081933
node5264.members.0.js
strain

2
938406
species

node5266.members.0.js
strain
1973478
2

no rank
2619626
5

1
2005462
species
node5268.members.0.js

2853227
node5269.members.0.js
species
2

1
1337936
node5270.members.0.js
species

1
99598
node5271.members.0.js
species

family
3
1182

1
3
genus
node5273.members.0.js
1203

2618749
2
no rank

1
1137095
node5275.members.0.js
species

species
node5276.members.0.js
2005464
1

1892259
2
family

1
genus
node5278.members.0.js
752201

110103
1
genus

1
156213
species

strain
node5281.members.0.js
2779889
1

no rank
1219117
3

3
node5283.members.0.js
species
1940762

1185
1
family

genus
373984
1

no rank
2676603
1

373994
node5287.members.0.js
species
1

35
3
1162
node5288.members.0.js
family

1163
8
genus

species
8
1165

node5291.members.0.js
strain
272123
8

22
1177
genus

no rank
node5293.members.0.js
2593658
1
19

1869241
species
node5294.members.0.js
4

2764711
node5295.members.0.js
species
3

species
node5296.members.0.js
1618022
1

4
species
node5297.members.0.js
2576904

2814655
species
node5298.members.0.js
3

2
2914041
node5299.members.0.js
species

node5300.members.0.js
species
1973475
1

224012
1
species

224013
strain
node5302.members.0.js
1

2
1
446679
node5303.members.0.js
species

2653204
node5304.members.0.js
strain
1

56106
2
genus

1
2630404
no rank

1
species
node5307.members.0.js
2005457

species
142864
1

56107
node5309.members.0.js
strain
1

1
1798710
phylum

no rank
1897007
1

1
1899017
species
node5312.members.0.js

49928
14
no rank

node5314.members.0.js
species
1869227
10

883811
2
species

2
1235284
isolate
node5316.members.0.js

species
node5317.members.0.js
1561003
1

131430
node5318.members.0.js
species
1

276155
6466
2759
superkingdom
node5319.members.0.js

1658
51250
clade
node5320.members.0.js
33154

16027
83
4751
kingdom
node5321.members.0.js

451864
subkingdom
node5322.members.0.js
15830
58

no rank
754015
1

1
species
node5324.members.0.js
754009

739
8
5204
phylum
node5325.members.0.js

1
122
node5326.members.0.js
subphylum
29000

class
6
432005

6
1
204043
order
node5328.members.0.js

family
34418
5

1
5
node5330.members.0.js
genus
203525

node5331.members.0.js
species
106018
1

105772
species
node5332.members.0.js
3

class
162480
3

3
48846
order

1
54740
family

54741
1
genus

54742
node5337.members.0.js
species
1

family
1
2877714

genus
1
2877715

2877780
node5340.members.0.js
species
1

family
1798865
1

36054
1
genus

node5343.members.0.js
species
4997
1

162484
72
class

order
node5345.members.0.js
5258
2
72

family
node5346.members.0.js
5262
3
54

51
5296
genus

1505670
node5348.members.0.js
species
3

1
node5349.members.0.js
species
27344

species
node5350.members.0.js
5297
2
27

forma specialis
56615
25

25
strain
node5352.members.0.js
418459

1
20
species
node5353.members.0.js
27350

19
168172
forma specialis
node5354.members.0.js

family
1
145793

genus
169998
1

node5357.members.0.js
species
170000
1

5259
13
family

genus
13
5260

203908
13
species

13
747676
node5361.members.0.js
strain

no rank
190595
1

genus
1304414
1

1
1304415
species
node5364.members.0.js

family
1
198672

203918
1
genus

1
species
node5367.members.0.js
1680363

1
432025
class

order
1
432026

1
165795
family

genus
34348
1

species
1
34349

1
764103
strain
node5373.members.0.js

162481
class
node5374.members.0.js
39
1

1
55070
order

55071
1
family

genus
34416
1

1
288790
species
node5378.members.0.js

231213
28
order

family
1799696
28

5533
28
genus

4
5535
species
node5382.members.0.js

species
29898
8

strain
node5384.members.0.js
578459
8

16
5
5286
species
node5385.members.0.js

1130832
strain
node5386.members.0.js
11

no rank
4
162483

family
1799785
3

3
2
1803523
genus
node5389.members.0.js

1
106019
node5390.members.0.js
species

256806
node5391.members.0.js
genus
1

order
5
231212

1163720
5
family

5
2
5277
node5394.members.0.js
genus

2
2630045
no rank

662878
species
node5396.members.0.js
2

5278
species
node5397.members.0.js
1

subphylum
2204096
2

class
2
431957

2
431958
order

family
2
431959

genus
148959
2

1
1708541
species

1
671144
node5404.members.0.js
strain

1
245174
species

1299270
strain
node5406.members.0.js
1

1
136247
no rank

1
175244
node5408.members.0.js
species

node5409.members.0.js
subphylum
5302
3
376

155616
class
node5410.members.0.js
112
2

27
90886
order

5408
27
family

genus
27
5209

node5414.members.0.js
species
5210
25

1
node5415.members.0.js
species
104410

5212
node5416.members.0.js
species
1

order
1851469
41

40
2
1759442
family
node5418.members.0.js

genus
105983
22

105713
species
node5420.members.0.js
1

252803
species
node5421.members.0.js
16

node5422.members.0.js
species
105984
5

genus
5552
6

species
82508
6

varietas
189963
6

strain
node5426.members.0.js
1186058
6

genus
1851468
5

5
species
node5428.members.0.js
143232

genus
1838142
5

879819
node5430.members.0.js
species
5

family
1851470
1

genus
1851472
1

1
node5433.members.0.js
species
211102

order
6
90883

3
1851551
family

genus
107449
3

3
species
node5437.members.0.js
264483

3
165808
family

5410
3
genus

5412
species
node5440.members.0.js
3

order
5234
36

family
1884633
32

490731
18
genus

species
324770
2

node5445.members.0.js
strain
1296121
2

324769
12
species

12
strain
node5447.members.0.js
1296100

1734106
node5448.members.0.js
species
4

14
3
5206
node5449.members.0.js
genus

species
node5450.members.0.js
5619
2

species group
5
1897064

2
5
species
node5452.members.0.js
5207

2
3
node5453.members.0.js
varietas
40410

214684
node5454.members.0.js
strain
1

species group
node5455.members.0.js
1884637
2

2
104669
species

2
1295533
strain
node5457.members.0.js

1
1910893
family

1
4998
genus

node5460.members.0.js
species
4999
1

family
1
1884640

genus
663591
1

node5463.members.0.js
species
4979
1

family
5215
2

no rank
2
195027

2
1503101
species
node5466.members.0.js

class
11
452332

order
11
28997

11
5254
family

139276
11
genus

11
1858805
species
node5471.members.0.js

node5472.members.0.js
class
155619
3
250

452335
1
subclass

1
452336
order

family
60108
1

genus
68779
1

node5477.members.0.js
species
68780
1

95
452333
subclass

1
30
order
node5479.members.0.js
68889

227334
11
suborder

family
5
389951

5
80744
genus

species
85982
5

5
341189
varietas

5
node5485.members.0.js
strain
578457

6
80634
family

6
80635
genus

80637
6
species

6
node5489.members.0.js
strain
741705

suborder
18
227332

family
18
227336

3
18
node5492.members.0.js
genus
5379

1
species
node5493.members.0.js
1912936

6
48586
species
node5494.members.0.js

species
node5495.members.0.js
48578
2

node5496.members.0.js
species
48587
1

node5497.members.0.js
species
1912939
2

3
node5498.members.0.js
species
48563

65
1
5338
order
node5499.members.0.js

family
3
5351

40144
1
genus

1
40145
species
node5502.members.0.js

genus
29882
2

species
2
29883

486041
strain
node5505.members.0.js
2

7
1
184208
family
node5506.members.0.js

5
184431
genus

species
5
5346

5
240176
node5509.members.0.js
strain

2791032
1
genus

node5511.members.0.js
species
2316362
1

3
2024004
family

3
41247
genus

2126181
node5514.members.0.js
species
3

family
104366
9

9
5320
genus

5322
species
node5517.members.0.js
9

41954
1
family

genus
1
41955

235537
node5520.members.0.js
species
1

family
5
72117

genus
5352
5

5
species
node5523.members.0.js
5353

1
40562
family

1
5399
genus

1
5400
node5526.members.0.js
species

5339
4
family

4
5340
genus

node5529.members.0.js
species
5341
4

11
862241
family

11
47424
genus

11
47428
node5532.members.0.js
species

654128
8
family

genus
34448
8

181124
species
node5535.members.0.js
8

family
71934
7

7
71935
genus

7
648681
species
node5538.members.0.js

1
930979
family

genus
71927
1

189768
1
no rank

1
species
node5542.members.0.js
189767

1
34450
family

1
86085
genus

species
node5545.members.0.js
109634
1

family
3
5332

5333
3
genus

3
1
5334
node5548.members.0.js
species

578458
strain
node5549.members.0.js
2

151
5
355688
node5550.members.0.js
no rank

4
139380
order

family
40424
4

genus
167346
4

4
208960
species

strain
node5555.members.0.js
694068
4

order
452342
27

family
1
40458

154746
1
genus

1
205917
species
node5559.members.0.js

10
103376
family

genus
5644
7

2
7
node5562.members.0.js
species
40492

5
node5563.members.0.js
strain
721885

1
55339
genus

node5565.members.0.js
species
112237
1

166471
2
genus

2
2638731
no rank

2
1321930
species
node5568.members.0.js

68767
3
family

3
68768
genus

3
node5571.members.0.js
species
154745

family
40420
11

genus
13562
11

256003
11
no rank

node5575.members.0.js
species
984962
5
11

6
747525
strain
node5576.members.0.js

1
2
family
node5577.members.0.js
5401

genus
1
5402

node5579.members.0.js
species
71688
1

5303
node5580.members.0.js
order
72
2

81064
1
family

1
5307
genus

5308
species
node5583.members.0.js
1

family
396331
9

genus
5305
9

species
9
231932

9
strain
node5587.members.0.js
650164

family
2028214
4

4
83235
genus

4
104341
species

670580
node5591.members.0.js
strain
4

no rank
183979
14

11
320360
genus

node5594.members.0.js
species
2696576
11

3
599838
genus

3
species
node5596.members.0.js
599839

10
2028212
family

genus
5629
10

species
node5599.members.0.js
5630
9
10

1314785
node5600.members.0.js
strain
1

family
4
40465

genus
40466
4

4
node5603.members.0.js
species
139825

family
28
5317

1
551032
no rank

1
1193695
species
node5606.members.0.js

114154
3
genus

species
3
114155

3
strain
node5609.members.0.js
732165

genus
node5610.members.0.js
5324
1
21

6
species
node5611.members.0.js
160864

1
node5612.members.0.js
species
230624

4
5327
node5613.members.0.js
species

species
5325
9

717944
strain
node5615.members.0.js
9

5314
3
genus

node5617.members.0.js
species
34458
1

2
29884
species
node5618.members.0.js

order
30
36064

family
2
5237

5240
1
genus

1
5241
species
node5622.members.0.js

no rank
1
156516

1
1403776
node5624.members.0.js
species

57201
4
family

genus
36065
4

4
1750568
species
node5627.members.0.js

5250
24
family

genus
1322061
16

16
species
node5630.members.0.js
456999

genus
8
5251

2600200
8
no rank

170446
species
node5633.members.0.js
8

order
56487
4

4
56488
family

160871
genus
node5636.members.0.js
3
1

160872
species
node5637.members.0.js
2

1
56489
genus
node5638.members.0.js

order
452338
6

family
908827
6

genus
6
133746

species
202698
6

741275
node5643.members.0.js
strain
6

452339
3
order

452340
3
family

genus
3
40443

3
104355
species

strain
node5648.members.0.js
670483
3

subphylum
node5649.members.0.js
452284
1
230

5257
95
class

95
5267
order

family
node5652.members.0.js
5268
9
95

1392992
6
genus

species
node5654.members.0.js
1134040
2

4
species
node5655.members.0.js
249478

genus
36
63261

84753
node5657.members.0.js
species
36

5269
7
genus

5270
6
species

237631
strain
node5660.members.0.js
6

1
185366
species
node5661.members.0.js

6
63259
genus

species
63383
6

1398559
node5664.members.0.js
strain
6

15
1804794
genus

species
15
1392244

15
1365824
strain
node5667.members.0.js

63298
5
genus

1
5
node5669.members.0.js
species
84751

4
node5670.members.0.js
strain
1277687

genus
11
63265

species
node5672.members.0.js
72558
3
5

999809
node5673.members.0.js
strain
1

forma specialis
node5674.members.0.js
72559
1

species
node5675.members.0.js
280036
4

49012
species
node5676.members.0.js
2

class
452283
49

order
5404
24

62920
5
family

genus
215249
5

5
215250
species
node5681.members.0.js

family
19
190068

215251
19
genus

17
node5684.members.0.js
species
1280837

node5685.members.0.js
species
215252
2

5
62913
order

family
5
62919

5
5280
genus

species
5
5281

strain
node5690.members.0.js
1037660
5

742846
2
order

2
742847
family

401624
2
genus

node5694.members.0.js
species
1522189
2

order
4
62914

162479
4
no rank

genus
4
1500560

4
species
node5698.members.0.js
58919

order
162475
14

162477
node5700.members.0.js
no rank
14
2

11
561108
genus

species
node5702.members.0.js
1569628
11

1981958
1
genus

1
1684307
species
node5704.members.0.js

class
1538075
85

162474
85
order

742845
85
family

genus
84
55193

20
15
76775
species
node5709.members.0.js

strain
node5710.members.0.js
425264
5

4
5
species
node5711.members.0.js
76777

1230383
node5712.members.0.js
strain
1

77020
species
node5713.members.0.js
2

4
55194
species
node5714.members.0.js

52
53
node5715.members.0.js
species
76773

1
425265
strain
node5716.members.0.js

1
1836866
no rank

1
node5718.members.0.js
species
338567

15032
8
4890
phylum
node5719.members.0.js

14994
19
716545
node5720.members.0.js
clade

subphylum
node5721.members.0.js
147538
14
14799

723
14758
node5722.members.0.js
clade
716546

clade
9122
715962

class
1
147539

39677
1
order

family
1
39678

genus
1
39679

1
node5728.members.0.js
species
2488734

9121
275
147541
class
node5729.members.0.js

451868
node5730.members.0.js
subclass
4554
4

1
47
order
node5731.members.0.js
1111111

20
1111112
family

1494215
20
genus

253628
node5734.members.0.js
species
20

family
26
5023

26
5024
genus

26
50376
node5737.members.0.js
species

order
node5738.members.0.js
92860
68
4447

4060
87
715340
suborder
node5739.members.0.js

405
38
683158
node5740.members.0.js
family

62
5453
genus

node5742.members.0.js
species
5454
62

74
749461
genus

74
species
node5744.members.0.js
749465

55170
102
genus

species
100019
86

86
1150837
node5747.members.0.js
strain

16
749589
node5748.members.0.js
species

301206
60
genus

60
species
node5750.members.0.js
301207

69
1
749880
genus
node5751.members.0.js

no rank
68
2606853

68
species
node5753.members.0.js
2802321

150
34374
family

genus
150
5021

220671
no rank
node5756.members.0.js
150
45

species
28
5022

225342
3
no rank

985895
node5759.members.0.js
strain
3

25
node5760.members.0.js
no rank
225343

77
4
220672
species
node5761.members.0.js

node5762.members.0.js
no rank
225344
37

node5763.members.0.js
no rank
225338
36

109
2
5020
node5764.members.0.js
family

1
node5765.members.0.js
genus
33174

genus
1351751
103

13684
species
node5767.members.0.js
103
92

11
strain
node5768.members.0.js
321614

3
798159
genus

3
798162
node5770.members.0.js
species

family
32
221670

genus
45141
32

45142
32
species

32
1168544
node5774.members.0.js
strain

node5775.members.0.js
family
28556
31
3275

node5776.members.0.js
genus
5027
17
95

45151
39
species

39
strain
node5778.members.0.js
426418

39
30
53485
node5779.members.0.js
species

97480
forma
node5780.members.0.js
9

5502
genus
node5781.members.0.js
2

91493
61
genus

61
93612
species

671987
strain
node5784.members.0.js
61

33194
genus
node5785.members.0.js
189
61

45130
34
species

34
node5787.members.0.js
strain
665912

species
37
101162

node5789.members.0.js
strain
930090
37

species
node5790.members.0.js
5016
11
45

34
strain
node5791.members.0.js
665024

6
40125
species

6
930091
strain
node5793.members.0.js

species
5017
6

930089
strain
node5795.members.0.js
6

525236
4
genus

461172
species
node5797.members.0.js
4

node5798.members.0.js
genus
5598
55
2890

section
4
2499238

4
species
node5800.members.0.js
29001

section
2499258
1

1
45303
species
node5802.members.0.js

section
41
2499266

node5804.members.0.js
species
119953
41

52
2499270
section

52
1187941
species
node5806.members.0.js

2692
404
2499237
node5807.members.0.js
section

98
node5808.members.0.js
species
156630

304
species
node5809.members.0.js
1187904

187734
species group
node5810.members.0.js
1885
2

5599
node5811.members.0.js
species
1883

1
species
node5812.members.0.js
160389

section
2499262
45

45
48100
node5814.members.0.js
species

genus
3
95729

1
species
node5816.members.0.js
183478

2
235069
node5817.members.0.js
species

family
2
1460697

78388
2
genus

2
node5820.members.0.js
species
1077358

no rank
51
147498

genus
51
1450170

species
1450171
51

51
1450172
strain
node5824.members.0.js

1255046
node5825.members.0.js
suborder
138
1

91
221678
family

genus
90
125369

90
species
node5828.members.0.js
1460663

1892769
1
genus

species
node5830.members.0.js
2777658
1

715496
node5831.members.0.js
family
1

45
1208339
family

genus
100048
45

45
node5834.members.0.js
species
390896

43
55176
family

genus
45153
42

species
node5837.members.0.js
318751
42

1
no rank
node5838.members.0.js
718227

family
56
548648

genus
56
548651

56
548649
species

56
1392245
strain
node5842.members.0.js

family
27
717954

27
741162
genus

27
673940
node5845.members.0.js
species

1
281124
family

1
59585
genus

node5848.members.0.js
species
59586
1

family
2126473
3

genus
227062
3

3
1547544
species
node5851.members.0.js

56
603422
order

281242
56
family

56
574786
genus

56
574789
species
node5855.members.0.js

no rank
node5856.members.0.js
159987
8
575

2810619
75
genus

1
75
node5858.members.0.js
species
61459

74
1168221
node5859.members.0.js
strain

1
1572707
order

43216
1
family

genus
358867
1

1
species
node5863.members.0.js
796411

286660
86
genus

node5865.members.0.js
species
286661
86

46
716585
order

family
46
152637

470097
3
genus

1
470100
species
node5869.members.0.js

2614217
2
no rank

2
species
node5871.members.0.js
1499996

genus
43
470095

470096
node5873.members.0.js
species
43

order
node5874.members.0.js
451869
11
316

family
192931
2

genus
1248177
2

1248288
species
node5877.members.0.js
2

family
65
1450293

genus
65
462253

species
462254
65

65
1176127
strain
node5881.members.0.js

family
1424649
1

genus
1
121621

1151445
node5884.members.0.js
species
1

14
237
node5885.members.0.js
family
45131

35724
1
genus

1
species
node5887.members.0.js
35725

genus
66739
58

node5889.members.0.js
species
45133
58

66735
node5890.members.0.js
genus
77
1

76
species
node5891.members.0.js
236234

genus
87
407951

87
species
node5893.members.0.js
310453

family
1
218400

genus
218402
1

node5896.members.0.js
species
706121
1

family
281210
1

1263494
1
genus

species
node5899.members.0.js
1709461
1

genus
1937197
1

1
node5901.members.0.js
species
1937198

32
2714147
order

241722
32
family

genus
258075
32

species
1341166
32

32
strain
node5906.members.0.js
1392243

node5907.members.0.js
genus
1837870
3
5

species
node5908.members.0.js
1423624
1

1
species
node5909.members.0.js
1423644

703495
3
order

2
281213
family

genus
1037661
2

2530477
node5913.members.0.js
species
2

1150844
1
no rank

genus
1
1960840

1960841
node5916.members.0.js
species
1

53
3716
node5917.members.0.js
subclass
451867

308
2726946
order

452563
308
family

12
308
genus
node5920.members.0.js
5498

species
node5921.members.0.js
1758478
1

470174
node5922.members.0.js
species
7

1
2511077
species
node5923.members.0.js

2831546
species
node5924.members.0.js
1

2614628
no rank
node5925.members.0.js
26
6

node5926.members.0.js
species
1452520
1

species
node5927.members.0.js
1707700
19

7
887083
node5928.members.0.js
species

species
node5929.members.0.js
29917
69

9
1116209
species
node5930.members.0.js

92950
node5931.members.0.js
species
94

29918
node5932.members.0.js
species
75

1970641
node5933.members.0.js
species
1

node5934.members.0.js
species
29919
5

2726947
order
node5935.members.0.js
2359
47

744530
128
family

genus
128
112488

species
112489
128

1314786
node5939.members.0.js
strain
128

2
281250
family

2
286616
genus

286617
species
node5942.members.0.js
2

668547
family
node5943.members.0.js
351
1

genus
2072583
186

186
species
node5945.members.0.js
245834

483074
157
genus

157
1709381
species

157
717646
node5948.members.0.js
strain

genus
286563
1

1
2643983
no rank

1485003
node5951.members.0.js
species
1

4
91942
genus

91943
node5953.members.0.js
species
4

genus
93481
1

species
node5955.members.0.js
93486
1

genus
470130
1

1
237593
species
node5957.members.0.js

node5958.members.0.js
family
93133
103
1831

node5959.members.0.js
genus
29002
105
387

88
species
node5960.members.0.js
122368

84275
species
node5961.members.0.js
103

90
438356
species
node5962.members.0.js

1
node5963.members.0.js
species
29003

395590
322
genus

species
322
395010

node5966.members.0.js
strain
1080233
322

genus
2
1515437

2
species
node5968.members.0.js
1431932

237179
113
genus

species
85929
113

113
strain
node5971.members.0.js
692275

162
3
112497
node5972.members.0.js
genus

159
112498
species
node5973.members.0.js

161
131324
genus

161
4
1873960
species
node5975.members.0.js

strain
node5976.members.0.js
383855
157

2897311
251
genus

251
5499
node5978.members.0.js
species

genus
1047167
327

1047171
species
node5980.members.0.js
327
289

6
1276538
node5981.members.0.js
strain

3
strain
node5982.members.0.js
1276537

336722
node5983.members.0.js
strain
29

2
node5984.members.0.js
genus
687950

genus
242508
1

species
node5986.members.0.js
64363
1

node5987.members.0.js
order
5014
1
993

64899
12
family

46635
2
genus

284138
species
node5990.members.0.js
2

genus
64497
10

10
64499
node5992.members.0.js
species

980
1570301
family

980
105
5579
node5994.members.0.js
genus

1592451
node5995.members.0.js
species
1

559561
128
species

node5997.members.0.js
strain
1043004
128

node5998.members.0.js
species
46634
295
411

1043003
node5999.members.0.js
strain
116

239
3
5580
node6000.members.0.js
species

1043002
strain
node6001.members.0.js
236

96
1042127
species

96
1043005
node6003.members.0.js
strain

order
1
45676

family
40995
1

genus
1
40996

40997
species
node6007.members.0.js
1

2
134362
order

2
467990
no rank

2
1487856
genus

702019
node6011.members.0.js
species
2

no rank
209473
1

1
species
node6013.members.0.js
748633

17
1217819
class

1
2126967
order

family
1
2126968

genus
1
40219

species
node6018.members.0.js
40221
1

16
1217820
order

1217822
16
family

genus
1217823
16

species
1217826
16

16
1328760
strain
node6023.members.0.js

22
3196
node6024.members.0.js
clade
715989

147550
class
node6025.members.0.js
2940
108

node6026.members.0.js
subclass
222543
43
2234

order
1
1127803

2016478
1
family

genus
1096518
1

1
species
node6030.members.0.js
1096519

order
node6031.members.0.js
5125
57
1680

family
5129
148

37
148
node6033.members.0.js
genus
5543

25
26
species
node6034.members.0.js
51453

node6035.members.0.js
strain
431241
1

16
species
node6036.members.0.js
398673

101201
species
node6037.members.0.js
14

species
node6038.members.0.js
29875
11
17

1331945
node6039.members.0.js
strain
6

5544
8
species

983964
strain
node6041.members.0.js
8

1491479
species
node6042.members.0.js
5

11
58853
species
node6043.members.0.js

14
63577
node6044.members.0.js
species

family
node6045.members.0.js
474943
5
77

44
5
45234
node6046.members.0.js
genus

node6047.members.0.js
species
114497
1
17

node6048.members.0.js
strain
1081104
16

22
16
73501
node6049.members.0.js
species

6
983644
node6050.members.0.js
strain

genus
5581
28

species
node6052.members.0.js
176275
15
28

13
655819
strain
node6053.members.0.js

family
node6054.members.0.js
474942
1
88

5
474995
genus

species
node6056.members.0.js
268505
5

genus
37
1052105

33203
species
node6058.members.0.js
37

genus
42367
25

104307
species
node6060.members.0.js
1

species
node6061.members.0.js
111463
24

genus
98402
20

20
node6063.members.0.js
species
98403

4
132
node6064.members.0.js
family
34397

22
124426
genus

node6066.members.0.js
species
1159556
22

51
12
5529
genus
node6067.members.0.js

6
7
node6068.members.0.js
species
500148

1276141
strain
node6069.members.0.js
1

92637
12
species

12
strain
node6071.members.0.js
655827

92629
17
species

1081103
strain
node6073.members.0.js
17

568076
node6074.members.0.js
species
3
1

2
strain
node6075.members.0.js
655844

node6076.members.0.js
genus
5112
13
37

3
species
node6077.members.0.js
2834323

species
14
35717

877507
strain
node6079.members.0.js
14

7
2
5113
node6080.members.0.js
species

4
node6081.members.0.js
subspecies
1616224

1
2570311
subspecies
node6082.members.0.js

5110
genus
node6083.members.0.js
3
1

node6084.members.0.js
species
5111
2

243023
11
genus

11
280754
species

1380566
isolate
node6087.members.0.js
11

42305
4
genus

4
696354
node6089.members.0.js
species

no rank
162454
43

genus
5
159075

node6092.members.0.js
species
5044
3

2689091
1
no rank

1420317
species
node6094.members.0.js
1

species
node6095.members.0.js
261921
1

genus
45244
38

38
2614577
node6097.members.0.js
species

103887
23
family

241409
23
genus

species
node6100.members.0.js
1094350
23

45492
2
family

genus
152774
2

species
node6103.members.0.js
152775
2

110618
node6104.members.0.js
family
1110
16

genus
1079112
30

species
node6106.members.0.js
1079257
30

32
140106
genus

32
78403
species
node6108.members.0.js

57138
19
genus

no rank
19
2779503

19
182845
node6111.members.0.js
species

5506
node6112.members.0.js
genus
1013
130

679429
1
species group

61284
node6114.members.0.js
species
1

569360
species group
node6115.members.0.js
117
47

17
36050
node6116.members.0.js
species

12
5
101028
species
node6117.members.0.js

1028729
node6118.members.0.js
strain
7

1
5516
node6119.members.0.js
species

26
56646
species
node6120.members.0.js

14
13
5518
node6121.members.0.js
species

1
229533
node6122.members.0.js
strain

27
48865
species
node6123.members.0.js

species group
node6124.members.0.js
232080
11
55

1147111
species
node6125.members.0.js
1

species
node6126.members.0.js
169388
19

species
24
2747968

24
node6128.members.0.js
no rank
660122

species group
node6129.members.0.js
171627
70
504

5127
node6130.members.0.js
species
49
46

node6131.members.0.js
strain
1279085
3

78861
node6132.members.0.js
species
1

1
192009
species
node6133.members.0.js

42677
node6134.members.0.js
species
53

node6135.members.0.js
species
120644
3

120
117187
species

334819
strain
node6137.members.0.js
120

species
22
948311

strain
node6139.members.0.js
1227346
22

node6140.members.0.js
species
192010
26

species
node6141.members.0.js
520446
1

1
56676
node6142.members.0.js
species

node6143.members.0.js
species
1567544
157

1
2800332
no rank

42327
species
node6145.members.0.js
1

75
10
450425
species group
node6146.members.0.js

2675880
node6147.members.0.js
species
40

node6148.members.0.js
species
231269
25

6
1042133
species
node6149.members.0.js

species group
node6150.members.0.js
171631
50
95

node6151.members.0.js
species
5507
19
40

5
660029
strain
node6152.members.0.js

4
strain
node6153.members.0.js
660027

4
654392
forma specialis
node6154.members.0.js

forma specialis
8
59765

8
strain
node6156.members.0.js
426428

species
2502994
5

5
node6158.members.0.js
strain
1089451

no rank
2593666
2

29916
species
node6160.members.0.js
1

1619569
node6161.members.0.js
species
1

order
5592
20

family
20
5593

41687
20
genus

20
563466
node6165.members.0.js
species

1028384
node6166.members.0.js
order
490
6

family
398
681950

398
30
5455
node6168.members.0.js
genus

2707350
37
no rank

species
37
80884

759273
node6171.members.0.js
strain
37

no rank
2707336
40

node6173.members.0.js
species
1095194
40

2707348
103
no rank

5
103
species
node6175.members.0.js
31870

645133
strain
node6176.members.0.js
98

2707339
33
no rank

33
5467
species
node6178.members.0.js

node6179.members.0.js
species
1209926
20

49
10
2707335
no rank
node6180.members.0.js

1209932
species
node6181.members.0.js
39

40
86
no rank
node6182.members.0.js
2707338

17
474922
species
node6183.members.0.js

11
690259
node6184.members.0.js
species

5
1215731
species
node6185.members.0.js

node6186.members.0.js
species
690256
13

family
node6187.members.0.js
1033978
1
86

15
1401161
genus

1302862
15
species

15
1314773
node6190.members.0.js
strain

69
41
1036719
genus
node6191.members.0.js

16
15
27337
node6192.members.0.js
species

498257
strain
node6193.members.0.js
1

1051616
species
node6194.members.0.js
7

species
5
1051613

5
526221
strain
node6196.members.0.js

40657
genus
node6197.members.0.js
1

147551
22
no rank

265081
22
family

genus
265082
22

1093900
node6201.members.0.js
species
22

93
222545
subclass

37989
node6203.members.0.js
order
93
3

1812770
16
family

genus
16
1812772

16
node6206.members.0.js
species
1141098

family
26
1682405

26
67608
genus

1682393
species
node6209.members.0.js
26

40
1812776
family

37840
22
genus

393283
22
species

22
strain
node6213.members.0.js
1229662

genus
152317
18

18
species
node6215.members.0.js
152316

family
106263
1

1811811
1
genus

node6218.members.0.js
species
1699843
1

family
2033035
7

genus
2652702
1

node6221.members.0.js
species
2487000
1

42360
6
genus

326645
node6223.members.0.js
species
6

subclass
node6224.members.0.js
222544
14
483

node6225.members.0.js
order
639021
2
81

55
2528436
family

48558
genus
node6227.members.0.js
55
5

148305
node6228.members.0.js
species
22

species
node6229.members.0.js
1578925
17

node6230.members.0.js
species
318829
11

81093
family
node6231.members.0.js
24
1

29849
23
genus

23
36779
species

strain
node6234.members.0.js
644352
23

2
5135
order

2
5136
family

genus
54963
2

2
2086372
species
node6238.members.0.js

5151
46
order

46
1
5152
family
node6240.members.0.js

29907
genus
node6241.members.0.js
27
16

species
7
545650

7
strain
node6243.members.0.js
1398154

species
4
29908

node6245.members.0.js
strain
1397361
4

genus
18
360145

species
226899
18

node6248.members.0.js
strain
655863
18

node6249.members.0.js
order
5139
6
208

1
95343
family

genus
1
95344

1
273648
species
node6252.members.0.js

family
2609812
40

5144
node6254.members.0.js
genus
40
27

node6255.members.0.js
species
48703
8

2587412
5
species

5
node6257.members.0.js
strain
515849

family
node6258.members.0.js
5148
4
48

5140
genus
node6259.members.0.js
23
10

1
node6260.members.0.js
species
29879

species
40127
4

4
510951
strain
node6262.members.0.js

4
8
species
node6263.members.0.js
5141

4
strain
node6264.members.0.js
367110

genus
21
5146

5147
21
species

21
node6267.members.0.js
strain
771870

35718
family
node6268.members.0.js
113
1

genus
1920207
30

species
30
78579

30
node6271.members.0.js
strain
573729

genus
29
2609811

species
2587410
29

29
578455
node6274.members.0.js
strain

5149
53
genus

38033
37
species

37
node6277.members.0.js
strain
306901

species
209285
16

16
285224
varietas

759272
strain
node6280.members.0.js
16

5114
node6281.members.0.js
order
118
3

767018
42
family

36922
node6283.members.0.js
genus
42
7

19
748121
node6284.members.0.js
species

node6285.members.0.js
species
83186
16

399129
node6286.members.0.js
family
72
11

node6287.members.0.js
no rank
218105
2
42

19
305399
genus

2625752
19
no rank

19
2029752
species
node6290.members.0.js

genus
5115
21

5116
21
species

21
node6293.members.0.js
strain
660469

19
1276216
genus

node6295.members.0.js
species
1276217
19

family
1
5117

genus
83174
1

species
105487
1

node6299.members.0.js
varietas
694572
1

order
13
1775898

family
1756146
13

genus
13
65412

13
223192
species

13
strain
node6304.members.0.js
1286976

order
292576
1

1
79807
family

genus
79808
1

node6308.members.0.js
species
91930
1

class
147548
234

order
5120
51

node6311.members.0.js
family
34371
4
51

genus
34372
34

node6313.members.0.js
species
2867405
1

34373
33
species

node6315.members.0.js
forma specialis
62690
33

genus
62701
1

62727
species
node6317.members.0.js
1

12
1
5121
node6318.members.0.js
genus

species
node6319.members.0.js
36044
5

157594
species
node6320.members.0.js
6

149
5
5178
order
node6321.members.0.js

11
2589077
family

genus
47830
11

node6324.members.0.js
species
1316788
11

family
15
2907085

5100
15
genus

species
15
5101

node6328.members.0.js
strain
857342
15

family
2656784
12

2656786
12
genus

2656787
species
node6331.members.0.js
12

family
2755564
14

genus
86026
14

14
149040
node6334.members.0.js
species

2793945
19
family

19
2081418
genus

698440
19
species

698441
node6338.members.0.js
forma specialis
19
1

1072389
node6339.members.0.js
strain
18

28983
47
family

31
10
33196
genus
node6341.members.0.js

1
1463999
species
node6342.members.0.js

139641
node6343.members.0.js
species
6

species
node6344.members.0.js
40559
7
9

2
332648
strain
node6345.members.0.js

node6346.members.0.js
species
2478750
2

3
1964551
species
node6347.members.0.js

5
38447
genus

species
node6349.members.0.js
61186
1

4
61207
node6350.members.0.js
species

11
5179
genus

5180
11
species

665079
isolate
node6353.members.0.js
11

47743
13
family

13
47747
genus

node6356.members.0.js
no rank
186449
1
13

12
2482752
species

12
node6358.members.0.js
strain
1095630

12
5181
family

genus
101851
12

101852
12
species

12
1116229
node6362.members.0.js
strain

1
1971365
family

genus
1
54692

1
node6365.members.0.js
species
296800

33
221903
no rank

family
34379
33

2
33
genus
node6368.members.0.js
78156

15
342668
node6369.members.0.js
species

species
node6370.members.0.js
655981
16

no rank
1
159638

1
node6372.members.0.js
species
592827

147547
31
class

31
1520881
clade

28
388435
subclass

2
388450
order

157824
1
suborder

node6378.members.0.js
family
1343996
1

suborder
388451
1

169285
1
family

1
168920
genus

species
node6382.members.0.js
168921
1

26
5197
order

suborder
157822
26

family
node6385.members.0.js
78060
2
18

112415
node6386.members.0.js
genus
13
9

species
node6387.members.0.js
560253
1

species
node6388.members.0.js
112416
3

1
172634
node6389.members.0.js
genus

86620
genus
node6390.members.0.js
1

172620
1
genus

node6392.members.0.js
species
172621
1

8
56478
family

genus
8
93111

8
node6395.members.0.js
species
2732470

388283
2
subclass

38074
2
order

71598
1
family

1
1130102
subfamily

1130104
1
tribe

genus
1
71599

1
71600
species
node6402.members.0.js

56757
1
family

1
56761
genus

1986089
node6405.members.0.js
species
1

1520791
1
subclass

1
452227
order

87265
1
family

genus
1
87266

284235
species
node6410.members.0.js
1

class
315355
1

order
291611
1

family
157478
1

genus
2817108
1

1
2796175
species
node6415.members.0.js

147545
class
node6416.members.0.js
1668
35

10
1308
subclass
node6417.members.0.js
451871

33183
122
order

2
5103
family

5104
node6420.members.0.js
genus
2

family
node6421.members.0.js
299071
1
40

229219
genus
node6422.members.0.js
16
10

species
5
5039

node6424.members.0.js
strain
559297
5

1
1681229
species

1
559298
strain
node6426.members.0.js

5036
genus
node6427.members.0.js
22
9

13
8
5037
node6428.members.0.js
species

3
447093
strain
node6429.members.0.js

2
544711
node6430.members.0.js
strain

genus
1745324
1

1
121128
node6432.members.0.js
species

55
5
34384
node6433.members.0.js
family

5550
genus
node6434.members.0.js
19
5

2
63400
species

strain
node6436.members.0.js
663331
2

species
5551
10

10
strain
node6438.members.0.js
559305

species
63417
2

2
strain
node6440.members.0.js
663202

genus
1915381
13

species
11
63402

node6443.members.0.js
strain
535722
11

2
2203077
species
node6444.members.0.js

34392
5
genus

species
63405
5

strain
node6447.members.0.js
554155
5

13
63399
genus

74035
species
node6449.members.0.js
13

no rank
1593277
8

38946
genus
node6451.members.0.js
7
4

1048829
2
species

502779
strain
node6453.members.0.js
2

1
121759
species

502780
node6455.members.0.js
strain
1

genus
78589
1

no rank
2688504
1

1
node6458.members.0.js
species
859378

family
33184
17

33187
6
genus

species
6
33188

6
node6462.members.0.js
strain
336963

node6463.members.0.js
genus
5500
7
11

species
199306
4

2
node6465.members.0.js
strain
222929

strain
node6466.members.0.js
443226
2

5042
node6467.members.0.js
order
1176
26

1
102
node6468.members.0.js
family
28568

genus
1132856
11

species
11
68825

strain
node6471.members.0.js
1408163
11

90
2
5094
node6472.members.0.js
genus

4
46
node6473.members.0.js
section
2752537

node6474.members.0.js
species
37727
6

128442
node6475.members.0.js
species
7

10
28564
species

strain
node6477.members.0.js
441959
10

11
28572
species
node6478.members.0.js

1196081
node6479.members.0.js
species
8

section
9
2752543

9
1131652
node6481.members.0.js
species

section
2752542
24

node6483.members.0.js
species
121627
24

9
2752540
section

9
1441469
species
node6485.members.0.js

1036
28
1131492
family
node6486.members.0.js

165
32
5073
node6487.members.0.js
genus

13
species
node6488.members.0.js
27334

1
5083
species
node6489.members.0.js

36651
species
node6490.members.0.js
12

2593313
1
no rank

species
node6492.members.0.js
1164577
1

60172
species
node6493.members.0.js
16

11
5082
node6494.members.0.js
species

node6495.members.0.js
species
5078
15

node6496.members.0.js
species
69780
1

5077
species
node6497.members.0.js
5

1
21
no rank
node6498.members.0.js
254878

species
20
1108849

20
500485
node6500.members.0.js
strain

2
species
node6501.members.0.js
81474

21
node6502.members.0.js
species
69781

1835702
species
node6503.members.0.js
14

70110
13
genus

species
13
41063

13
1073090
strain
node6506.members.0.js

196
830
node6507.members.0.js
genus
5052

9
node6508.members.0.js
species
41047

species
6
446911

6
strain
node6510.members.0.js
1448315

species
41068
14

1392248
strain
node6512.members.0.js
14

30
species
node6513.members.0.js
1810919

node6514.members.0.js
species
1069201
3

396024
14
species

1388766
node6516.members.0.js
strain
14

2720874
10
subgenus

41413
10
species

node6519.members.0.js
strain
1160497
10

288669
4
species

1448311
strain
node6521.members.0.js
4

4
node6522.members.0.js
species
91492

node6523.members.0.js
species
209559
19

species
487661
1

1
1448322
strain
node6525.members.0.js

20
1220207
node6526.members.0.js
species

5
node6527.members.0.js
species
1517512

7
132259
node6528.members.0.js
species

6
41058
species
node6529.members.0.js

node6530.members.0.js
species
5068
1

5
75553
species
node6531.members.0.js

node6532.members.0.js
species
61420
3

319630
2
species

1448313
node6534.members.0.js
strain
2

1220188
node6535.members.0.js
species
19

subgenus
36
2720873

species
36
5066

1073089
strain
node6538.members.0.js
36

25
51019
species

strain
node6540.members.0.js
1448321
25

9
species
node6541.members.0.js
1506151

species
301854
13

node6543.members.0.js
strain
1448316
13

2720870
node6544.members.0.js
subgenus
105
7

138278
17
species

1392256
node6546.members.0.js
strain
17

23
1
162425
species
node6547.members.0.js

strain
node6548.members.0.js
227321
22

176161
node6549.members.0.js
species
1

75750
40
species

1036612
strain
node6551.members.0.js
40

46472
17
species

1036611
node6553.members.0.js
strain
17

41061
3
species

3
strain
node6555.members.0.js
1509407

species
319627
11

11
strain
node6557.members.0.js
1450535

3
1340029
species

1448314
node6559.members.0.js
strain
3

19
species
node6560.members.0.js
182096

1
1549217
species
node6561.members.0.js

species
5
340412

strain
node6563.members.0.js
1392255
5

species
1
319631

1448317
strain
node6565.members.0.js
1

18
979771
species

node6567.members.0.js
strain
1450539
18

3
1341132
node6568.members.0.js
species

34381
4
species

strain
node6570.members.0.js
1448312
4

44
5
2720872
node6571.members.0.js
subgenus

5057
30
species

30
344612
strain
node6573.members.0.js

6
1
746128
species
node6574.members.0.js

5
330879
strain
node6575.members.0.js

species
3
36630

3
node6577.members.0.js
strain
331117

2715281
node6578.members.0.js
species
1

293939
species
node6579.members.0.js
8

node6580.members.0.js
species
109264
6

81
12
2720871
subgenus
node6581.members.0.js

1
5062
species
node6582.members.0.js

species
33178
31

31
341663
strain
node6584.members.0.js

6
node6585.members.0.js
species
5059

5061
node6586.members.0.js
species
2
1

1
425011
node6587.members.0.js
strain

species
22
306088

node6589.members.0.js
strain
1392250
22

5053
7
species

690307
node6591.members.0.js
strain
7

1
1405805
node6592.members.0.js
species

20
species
node6593.members.0.js
138277

40382
species
node6594.members.0.js
1

species
1196635
3

3
1448310
node6596.members.0.js
strain

41067
node6597.members.0.js
species
12

species
19
319626

19
1450537
node6599.members.0.js
strain

family
1131624
12

genus
33202
12

12
264951
species
node6602.members.0.js

1
325
subclass
node6603.members.0.js
451870

312
3
34395
order
node6604.members.0.js

family
1341112
22

22
226991
genus

species
node6607.members.0.js
856651
1

species
293227
21

node6609.members.0.js
strain
1220924
21

43219
family
node6610.members.0.js
285
10

23
43220
genus

13
43228
species

13
strain
node6613.members.0.js
1182542

10
43229
species

10
strain
node6615.members.0.js
1182541

genus
node6616.members.0.js
40354
15
45

4
species
node6617.members.0.js
856822

1
node6618.members.0.js
species
254056

species
979981
5

node6620.members.0.js
strain
1442371
5

1367422
species
node6621.members.0.js
16

4
40355
species

4
1442368
strain
node6623.members.0.js

330783
1
genus

1
330785
species
node6625.members.0.js

node6626.members.0.js
genus
5583
1
113

9
species
node6627.members.0.js
91928

node6628.members.0.js
species
348802
22

29
215243
node6629.members.0.js
species

species
5970
8

node6631.members.0.js
strain
858893
8

10
1033840
species

10
1182545
strain
node6633.members.0.js

8
node6634.members.0.js
species
212818

26
91925
species
node6635.members.0.js

19
5600
genus

node6637.members.0.js
species
39412
2

species
node6638.members.0.js
1664694
17

13
5587
genus

13
86056
species

strain
node6641.members.0.js
1442369
13

7
61
node6642.members.0.js
genus
82105

1182553
9
species

9
node6644.members.0.js
strain
1182543

18
569365
species
node6645.members.0.js

species
11
470704

11
strain
node6647.members.0.js
1182544

1
5
species
node6648.members.0.js
89940

1442370
node6649.members.0.js
strain
4

species
11
86049

11
1279043
node6651.members.0.js
strain

family
1233474
1

genus
1758290
1

1758292
node6654.members.0.js
species
1

316340
1
no rank

node6656.members.0.js
species
1715222
1

10
146291
order

10
146292
family

10
364710
genus

364733
10
species

1263415
strain
node6661.members.0.js
10

no rank
634205
2

2
node6663.members.0.js
species
1769350

17
147549
class

order
5185
17

family
4
40289

genus
36048
4

species
39416
4

4
strain
node6669.members.0.js
656061

family
5192
12

12
5193
genus

4
12
node6672.members.0.js
section
1051054

7
species
node6673.members.0.js
1174673

1174677
node6674.members.0.js
species
1

5186
family
node6675.members.0.js
1

10
189478
class

order
189479
10

family
10
47021

2811529
1
genus

1
2576060
species
node6680.members.0.js

3
9
genus
node6681.members.0.js
47022

1
2478772
species
node6682.members.0.js

species
node6683.members.0.js
1798096
2

3
2813651
species

756982
node6685.members.0.js
strain
3

subphylum
176
147537

class
4891
176

node6688.members.0.js
order
4892
4
176

family
44277
12

genus
27308
12

12
54195
species

12
1344418
strain
node6692.members.0.js

12
410830
family

genus
410829
11

11
796027
node6695.members.0.js
species

genus
1
45787

1
45607
species
node6697.members.0.js

115784
10
family

3
604195
genus

907339
node6700.members.0.js
species
2

1
4903
node6701.members.0.js
species

genus
599737
3

3
node6703.members.0.js
species
1041607

460517
4
genus

4922
species
node6705.members.0.js
4

node6706.members.0.js
family
4893
2
43

300275
5
genus

species
node6708.members.0.js
1245769
1

4
381046
species

node6710.members.0.js
strain
559295
4

genus
node6711.members.0.js
4930
1
8

4
3
4932
species
node6712.members.0.js

node6713.members.0.js
strain
1294352
1

3
species
node6714.members.0.js
27291

113604
1
genus

1071379
1
species

1
1071380
strain
node6717.members.0.js

2
278028
genus

2
27289
species

2
node6720.members.0.js
strain
1071378

genus
374468
1

clade
600669
1

species
node6723.members.0.js
5478
1

3
71245
genus

species
2
588726

node6726.members.0.js
strain
1071383
2

species
432096
1

1071382
strain
node6728.members.0.js
1

1
3
node6729.members.0.js
genus
4953

1
node6730.members.0.js
species
1365886

1
node6731.members.0.js
species
4956

4
1196389
genus

42260
node6733.members.0.js
species
4

5
1
33170
genus
node6734.members.0.js

1
33169
species
node6735.members.0.js

45285
3
species

931890
strain
node6737.members.0.js
3

genus
4
4948

1
species
node6739.members.0.js
4950

48254
node6740.members.0.js
species
3

4910
5
genus

1
node6742.members.0.js
species
4911

4
28985
species
node6743.members.0.js

5
34365
family

genus
36034
5

5
36035
node6746.members.0.js
species

family
9
34353

4
4951
genus

4
4952
node6749.members.0.js
species

genus
5
1232588

5
species
node6751.members.0.js
2606893

clade
51
2916678

21
27319
family

genus
6
27320

species
3
27322

3
280587
varietas

3
869754
node6757.members.0.js
strain

2163413
species
node6758.members.0.js
3

36910
15
genus

45354
species
node6760.members.0.js
2

45357
node6761.members.0.js
species
1

node6762.members.0.js
species
418784
1

4
498019
node6763.members.0.js
species

node6764.members.0.js
species
36911
5
7

2
strain
node6765.members.0.js
306902

3
30
node6766.members.0.js
family
766764

genus
2
766733

2
4924
species

strain
node6769.members.0.js
322104
2

4
412764
genus

340170
4
species

4
619300
node6772.members.0.js
strain

5
507510
genus

1
species
node6774.members.0.js
717741

2
4
species
node6775.members.0.js
717740

2
node6776.members.0.js
strain
984485

genus
766728
2

4929
2
species

strain
node6779.members.0.js
294746
2

clade
6
1535325

36913
2
genus

species
2
36914

node6783.members.0.js
strain
379508
2

genus
1535326
4

5482
node6785.members.0.js
species
1

5476
node6786.members.0.js
species
1

species
42374
1

strain
node6788.members.0.js
573826
1

273371
1
species

1
1136231
strain
node6790.members.0.js

4958
4
genus

58627
node6792.members.0.js
species
2

node6793.members.0.js
species
4959
1
2

1
strain
node6794.members.0.js
284592

genus
2
766765

2
2315449
species

590646
node6797.members.0.js
strain
2

genus
2
1539666

46583
2
species

2
984487
node6800.members.0.js
strain

family
1156497
10

13366
3
genus

13502
node6803.members.0.js
species
2

5007
node6804.members.0.js
species
1

3
4919
genus

4909
node6806.members.0.js
species
1

species
2
4926

763406
strain
node6808.members.0.js
2

genus
461281
4

1005962
species
node6810.members.0.js
1

3
species
node6811.members.0.js
870730

241407
9
no rank

317045
6
genus

species
6
317047

6
strain
node6815.members.0.js
1382522

genus
1910789
3

5481
species
node6817.members.0.js
3

34366
11
family

node6819.members.0.js
genus
4943
4
11

2
species
node6820.members.0.js
44092

1725355
node6821.members.0.js
species
1

2
node6822.members.0.js
species
4944

2
2636529
no rank

2
2826930
node6824.members.0.js
species

subphylum
28
451866

class
147555
10

10
5008
order

10
27330
family

27331
node6829.members.0.js
genus
10
2

6
species
node6830.members.0.js
2754530

1
species
node6831.members.0.js
48147

27332
node6832.members.0.js
species
1

no rank
5
452562

genus
5605
5

5606
5
species

5
698492
node6836.members.0.js
strain

class
147553
3

37987
3
order

family
44281
3

genus
3
4753

2
38082
node6841.members.0.js
species

species
1
263815

node6843.members.0.js
strain
1069680
1

class
10
147554

order
10
34346

10
4894
family

4895
10
genus

4
2
4897
node6848.members.0.js
species

402676
strain
node6849.members.0.js
2

2
866546
species

2
653667
strain
node6851.members.0.js

1
4899
species

strain
node6853.members.0.js
483514
1

3
node6854.members.0.js
species
4896

no rank
45426
2

1
node6856.members.0.js
species
331496

node6857.members.0.js
species
451645
1

68
112252
no rank

phylum
6029
5

no rank
469895
4

1
144516
genus

2638002
1
no rank

1
698276
species
node6863.members.0.js

genus
1633384
1

species
node6865.members.0.js
1485682
1

586132
2
genus

586133
2
species

881290
node6868.members.0.js
strain
2

1
6032
suborder

family
36734
1

genus
1
6033

1
node6872.members.0.js
species
6035

4761
15
phylum

no rank
15
2683659

451454
1
class

79272
1
order

family
109943
1

genus
1
109944

1
node6879.members.0.js
no rank
2643455

class
14
451435

1
34478
order

family
1
34479

genus
1
4815

1
109760
species

645134
node6885.members.0.js
strain
1

order
8
451442

8
1142503
no rank

genus
100474
8

species
node6889.members.0.js
109871
1
8

7
node6890.members.0.js
strain
684364

2231171
5
order

family
286113
5

genus
286114
5

5
node6894.members.0.js
species
1806994

28
1913637
phylum

1
1137986
subphylum

2212732
1
class

order
1
214503

4854
1
family

genus
1
299330

64571
node6901.members.0.js
species
1

subphylum
18
214504

class
214506
18

18
36750
order

36751
18
family

2
231054
no rank

species
node6907.members.0.js
1147122
1

species
node6908.members.0.js
231055
1

16
1129544
genus

16
node6910.members.0.js
species
588596

9
451507
subphylum

9
2212703
class

order
9
4827

suborder
1344963
5

3
1344955
family

4842
3
genus

3
58291
species

1340429
strain
node6918.members.0.js
3

family
34489
2

1
2
node6920.members.0.js
genus
4830

1
node6921.members.0.js
species
51122

family
4
499202

genus
688353
4

688394
node6924.members.0.js
species
4

phylum
1913638
20

subphylum
13
1264859

13
1399770
class

13
4857
order

node6929.members.0.js
family
34481
5
11

genus
34484
1

node6931.members.0.js
species
34485
1

genus
76016
4

1
176681
node6933.members.0.js
species

3
76017
node6934.members.0.js
species

genus
1
39186

1
species
node6936.members.0.js
39303

34486
2
family

2878373
2
genus

2
species
node6939.members.0.js
76012

7
451828
subphylum

7
2219690
class

order
7
4861

7
4862
family

7
4867
genus

61395
node6945.members.0.js
species
7

no rank
57731
46

46
node6947.members.0.js
species
175245

6
33556
node6948.members.0.js
kingdom
33208

6
6040
phylum

6
6042
class

subclass
6
1779146

order
6
6049

178475
6
family

genus
6
178513

6
400682
node6955.members.0.js
species

64
33544
clade
node6956.members.0.js
6072

10197
1
phylum

140493
1
class

order
1
50341

1
86008
family

1
86009
genus

1
species
node6962.members.0.js
1403702

clade
node6963.members.0.js
33213
3963
33380

40
10894
node6964.members.0.js
clade
33511

phylum
10219
10

class
10
10220

10
10221
family

10222
10
genus

10
10224
species
node6969.members.0.js

36
10679
node6970.members.0.js
phylum
7711

subphylum
51
7735

51
2682552
class

order
2682553
51

51
7736
family

1
51
genus
node6975.members.0.js
7737

7741
species
node6976.members.0.js
8

node6977.members.0.js
species
7740
38

species
node6978.members.0.js
7739
4

subphylum
7712
77

class
7
30302

order
2507557
7

7
41302
family

34763
7
genus

7
34765
species
node6984.members.0.js

70
7713
class

49
32436
order

49
201955
family

genus
201956
49

49
species
node6989.members.0.js
2771288

order
17
7716

family
9
7717

genus
9
7718

9
species
node6993.members.0.js
7719

family
8
30274

8
56696
genus

species
node6996.members.0.js
59560
8

order
4
7720

7721
4
family

genus
4
7724

4
species
node7000.members.0.js
7725

subphylum
89593
10515

10515
4
7742
node7002.members.0.js
clade

clade
15
1476529

117569
14
class

order
7745
14

family
7746
14

genus
7756
14

14
node7008.members.0.js
species
7757

1
117565
class

7761
1
order

family
7762
1

subfamily
1
30309

genus
1
7763

1
7764
node7014.members.0.js
species

7776
node7015.members.0.js
clade
10496
15

88
7777
class

81
7778
subclass

117893
29
superorder

order
29
7858

family
29
30475

genus
117853
29

29
node7022.members.0.js
species
386614

119203
52
infraclass

clade
52
119197

1
52
node7025.members.0.js
superorder
119195

30496
6
order

6
7850
family

subfamily
6
7844

13396
6
genus

13397
species
node7030.members.0.js
6

30503
order
node7031.members.0.js
11
1

family
8
40580

genus
34767
8

species
node7034.members.0.js
36176
8

259919
2
family

1849765
2
genus

2
node7037.members.0.js
species
259920

30483
34
order

32
7826
family

genus
7829
32

7830
species
node7041.members.0.js
32

family
2
7839

2
7840
genus

species
node7044.members.0.js
30494
2

7863
7
subclass

order
7864
7

family
7
7865

genus
7866
7

7868
species
node7049.members.0.js
7

10393
117570
clade

602
10393
node7051.members.0.js
clade
117571

3
4553
node7052.members.0.js
superclass
7898

35
4540
class
node7053.members.0.js
186623

subclass
103
32440

103
7899
order

3
103
suborder
node7056.members.0.js
186622

family
7
7911

subfamily
7
186619

tribe
186621
7

7912
7
genus

node7061.members.0.js
species
7913
7

7900
93
family

subfamily
124129
93

tribe
124130
93

genus
node7065.members.0.js
7901
1
93

92
7906
node7066.members.0.js
species

41665
subclass
node7067.members.0.js
4402
2

32443
infraclass
node7068.members.0.js
4391
1

clade
1489340
22

clade
22
186624

2
22
node7071.members.0.js
clade
32521

7925
6
order

family
7930
6

genus
7931
6

node7075.members.0.js
species
118141
6

order
14
7933

46660
1
family

subfamily
1
182222

391221
1
genus

1
node7080.members.0.js
species
391222

family
7934
13

7935
13
genus

13
species
node7083.members.0.js
7936

16
4368
node7084.members.0.js
clade
1489341

186625
no rank
node7085.members.0.js
4230
99

738
3
186634
cohort
node7086.members.0.js

1489485
1
subcohort

order
1489486
1

family
170200
1

319781
1
genus

1
1868731
species
node7091.members.0.js

668
32519
subcohort

186626
node7093.members.0.js
clade
615
2

superorder
186628
80

order
9
8002

1489620
9
suborder

family
9
30771

genus
9
8004

9
species
node7099.members.0.js
8005

order
7991
17

suborder
1489739
17

family
5
7992

subfamily
5
42595

genus
5
7993

5
7994
species
node7105.members.0.js

family
node7106.members.0.js
42495
6
12

42525
3
genus

42526
node7108.members.0.js
species
3

3
42513
genus

42514
species
node7110.members.0.js
3

54
7995
order

1489793
suborder
node7112.members.0.js
54
8

25
7996
family

genus
25
7997

25
species
node7115.members.0.js
7998

31013
10
family

641818
10
genus

species
node7118.members.0.js
1234273
10

family
11
7999

genus
30992
11

11
node7121.members.0.js
species
310915

superorder
533
186627

order
7952
533

9
532
suborder
node7124.members.0.js
30727

8
2743726
family

2743731
8
subfamily

51137
8
genus

90988
node7128.members.0.js
species
8

3
125
node7129.members.0.js
family
7953

117
1
2743694
node7130.members.0.js
subfamily

92
7961
genus

species
node7132.members.0.js
7962
92

4
10
genus
node7133.members.0.js
75365

75366
node7134.members.0.js
species
3

307959
node7135.members.0.js
species
3

14
7956
genus

7957
species
node7137.members.0.js
14

4
2743705
subfamily

4
1606679
genus

1606681
species
node7140.members.0.js
4

subfamily
2743704
1

genus
75347
1

node7143.members.0.js
species
75348
1

2743709
390
family

subfamily
2743711
390

390
21
7954
node7146.members.0.js
genus

7955
node7147.members.0.js
species
174

135
node7148.members.0.js
species
1142201

60
242068
node7149.members.0.js
species

suborder
2509193
1

family
1
7968

genus
70542
1

1
species
node7153.members.0.js
70543

53
186633
clade

order
29140
53

53
186632
suborder

53
29142
family

53
29143
genus

53
29144
species
node7159.members.0.js

subcohort
66
282425

66
32446
order

suborder
1489460
13

family
node7163.members.0.js
55118
1
13

subfamily
7
55119

genus
34772
7

34773
species
node7166.members.0.js
7

5
7948
subfamily

genus
7949
5

species
node7169.members.0.js
7950
5

suborder
53
1489459

family
299319
53

53
299320
genus

53
299321
species
node7173.members.0.js

cohort
node7174.members.0.js
1489388
41
3393

41705
clade
node7175.members.0.js
466
3

23
8007
order

family
8008
23

8009
23
genus

8010
species
node7179.members.0.js
23

8006
440
order

440
19
8015
node7181.members.0.js
family

node7182.members.0.js
subfamily
504568
6
257

8033
6
genus

6
species
node7184.members.0.js
8040

8016
genus
node7185.members.0.js
39
13

1
species
node7186.members.0.js
8018

8022
node7187.members.0.js
species
6

4
8019
node7188.members.0.js
species

8017
species
node7189.members.0.js
1

2
node7190.members.0.js
species
74940

12
8023
species
node7191.members.0.js

3
206
genus
node7192.members.0.js
8028

5
node7193.members.0.js
species
8030

8032
node7194.members.0.js
species
198

subfamily
164
504567

3
164
genus
node7196.members.0.js
27772

59861
node7197.members.0.js
species
1

no rank
2649731
160

species
node7199.members.0.js
861768
160

clade
2886
123365

123366
2886
clade

123367
2886
clade

123368
node7203.members.0.js
clade
2886
15

clade
node7204.members.0.js
123369
6
2812

2698
136
1489872
node7205.members.0.js
clade

79
1489885
clade

order
1489894
79

8224
79
family

subfamily
79
186745

79
186749
tribe

49
79
genus
node7211.members.0.js
8234

8240
node7212.members.0.js
species
9

21
node7213.members.0.js
species
8236

12
746
clade
node7214.members.0.js
1489908

1
452
node7215.members.0.js
superorder
1489913

order
170
76071

suborder
28781
170

47757
170
family

subfamily
8088
170

node7220.members.0.js
genus
8089
2
170

163
8090
node7221.members.0.js
species

species
node7222.members.0.js
30732
5

order
node7223.members.0.js
28738
2
180

suborder
node7224.members.0.js
8087
1
129

8076
19
family

subfamily
19
136836

tribe
19
136838

genus
node7228.members.0.js
28741
1
19

node7229.members.0.js
species
77115
4

14
28743
node7230.members.0.js
species

family
101
8079

586240
node7232.members.0.js
subfamily
101
3

56612
1
genus

1
188138
species
node7234.members.0.js

8082
node7235.members.0.js
genus
13
4

4
species
node7236.members.0.js
8084

5
node7237.members.0.js
species
32473

genus
75
8080

48701
species
node7239.members.0.js
4

node7240.members.0.js
species
8081
70

48699
species
node7241.members.0.js
1

genus
33527
9

9
node7243.members.0.js
species
33528

family
28756
8

genus
8
8077

8
node7246.members.0.js
species
8078

49
45443
suborder

family
405002
27

28779
27
genus

105023
species
node7250.members.0.js
27

family
22
28771

942014
1
genus

1
node7253.members.0.js
species
451745

genus
8
326431

8
species
node7255.members.0.js
37003

52669
13
genus

species
node7257.members.0.js
52670
13

8075
101
order

30700
4
family

4
32459
genus

node7261.members.0.js
species
1250792
4

family
97
270656

genus
300305
97

species
node7264.members.0.js
300306
97

clade
1489920
177

177
1
1489921
node7266.members.0.js
order

suborder
102
123349

63826
102
family

557415
102
subfamily

genus
210581
102

441366
node7271.members.0.js
species
102

56717
74
suborder

74
56718
family

74
703913
subfamily

genus
74
94311

74
181472
species
node7276.members.0.js

1489910
33
superorder

1489911
33
order

8113
node7279.members.0.js
family
33
1

clade
319095
24

24
5
318546
node7281.members.0.js
subfamily

7
2
319058
node7282.members.0.js
tribe

genus
8121
1

1
8153
node7284.members.0.js
species

genus
2
137270

node7286.members.0.js
species
137271
2

1
143623
genus

1
57445
species group

node7289.members.0.js
species
106582
1

genus
195936
1

1
node7291.members.0.js
species
303518

1315725
9
tribe

4
9
node7293.members.0.js
genus
8139

5
8128
node7294.members.0.js
species

3
319069
tribe

genus
3
32506

species
node7297.members.0.js
32507
3

8
319056
clade

318559
8
subfamily

tribe
318529
8

8
61816
genus

63155
species
node7302.members.0.js
8

1489909
72
no rank

30863
node7304.members.0.js
family
16
1

genus
1
80969

node7306.members.0.js
species
80972
1

genus
9
80992

144197
node7308.members.0.js
species
9

genus
5
80965

5
80966
species
node7310.members.0.js

family
205120
56

56
210631
genus

56
210632
node7313.members.0.js
species

1489892
clade
node7314.members.0.js
234
11

88
43697
order

suborder
129920
82

family
94233
82

genus
82
94234

node7319.members.0.js
species
205130
82

129918
6
suborder

43698
6
family

43699
6
genus

43700
node7323.members.0.js
species
6

1489900
135
order

suborder
50370
135

46
270602
family

158449
46
subfamily

45
158455
genus

45
species
node7329.members.0.js
158456

genus
1
158453

1
1628438
node7331.members.0.js
species

family
89
64142

genus
89
64143

89
64144
node7334.members.0.js
species

1489883
41
clade

order
129912
41

suborder
41
1489884

41
72045
family

subfamily
31
129914

103719
31
genus

161584
species
node7341.members.0.js
31

subfamily
10
129917

10
72046
genus

10
109280
node7344.members.0.js
species

clade
155
1489875

22
1489878
order

22
8219
suborder

8220
18
family

497678
node7349.members.0.js
subfamily
18
2

genus
5
86204

409849
species
node7351.members.0.js
5

11
150287
genus

11
species
node7353.members.0.js
150288

4
166784
family

genus
86243
4

4
species
node7356.members.0.js
357170

1489876
133
order

133
1489877
suborder

83881
133
family

subfamily
475176
133

genus
133
375763

133
species
node7362.members.0.js
375764

1489922
clade
node7363.members.0.js
991
31

order
1489931
159

node7365.members.0.js
family
8169
3
159

8174
106
genus

106
node7367.members.0.js
species
8175

50
8176
genus

50
8177
species
node7369.members.0.js

14
1489940
order

14
1545897
superfamily

8180
10
family

10
6
27705
genus
node7373.members.0.js

27706
node7374.members.0.js
species
4

family
119486
4

119487
4
genus

species
node7377.members.0.js
119488
4

order
5
1545895

family
5
30828

109904
5
genus

5
109905
species
node7381.members.0.js

order
45
31022

44
31028
suborder

44
32517
superfamily

family
44
31031

genus
4
47144

4
node7387.members.0.js
species
99883

40
31032
genus

species
node7389.members.0.js
31033
40

superfamily
1
31023

1
31024
family

genus
392898
1

node7393.members.0.js
species
392899
1

4
538
order
node7394.members.0.js
8111

14
8107
suborder

14
274692
family

subfamily
14
274705

genus
14
34820

72105
species
node7399.members.0.js
14

2
145
node7400.members.0.js
suborder
8100

1490020
15
infraorder

69291
node7402.members.0.js
family
15
3

genus
6
69292

node7404.members.0.js
species
69293
1
6

5
subspecies
node7405.members.0.js
481459

6
134919
genus

6
134920
node7407.members.0.js
species

76
2
8192
node7408.members.0.js
infraorder

7
8202
family

genus
7
433404

7
species
node7411.members.0.js
433405

56724
67
family

subfamily
181468
67

67
56725
genus

56726
species
node7415.members.0.js
67

52
1490021
infraorder

family
49
8092

genus
61642
49

49
61643
species
node7419.members.0.js

family
3
8101

181456
3
subfamily

8102
3
genus

3
8103
species
node7423.members.0.js

13
159
suborder
node7424.members.0.js
8205

71
30806
family

71
52238
genus

71
52239
node7427.members.0.js
species

4
8214
family

4
8217
genus

4
8218
species
node7430.members.0.js

4
8206
family

4
35729
genus

40690
node7433.members.0.js
species
4

family
67
36203

genus
67
56715

species
node7436.members.0.js
56716
67

suborder
200
1489943

family
30871
200

200
274794
subfamily

2
200
node7440.members.0.js
tribe
1505891

genus
node7441.members.0.js
94231
7
107

species
node7442.members.0.js
203261
1

11
species
node7443.members.0.js
310571

species
node7444.members.0.js
293821
88

genus
134629
91

node7446.members.0.js
species
160734
91

suborder
16
8112

8165
16
family

subfamily
641308
3

283033
3
genus

3
283035
node7451.members.0.js
species

subfamily
9
698016

genus
9
54318

4
909696
subgenus

node7455.members.0.js
species
54343
4

subgenus
5
909700

417921
node7457.members.0.js
species
5

641307
4
subfamily

4
1
8166
node7459.members.0.js
genus

species
node7460.members.0.js
8168
2

1
node7461.members.0.js
species
8167

order
35
1489928

family
35
8247

genus
10
202584

241271
species
node7465.members.0.js
10

98381
14
genus

14
species
node7467.members.0.js
1203425

30799
11
genus

11
node7469.members.0.js
species
56723

1489923
95
no rank

76
1
30870
node7471.members.0.js
family

genus
29
215359

240163
species
node7473.members.0.js
29

46
215357
genus

46
node7475.members.0.js
species
215358

30869
6
family

genus
75037
6

species
node7478.members.0.js
75038
6

13
42148
family

genus
13488
11

11
13489
node7481.members.0.js
species

34815
2
genus

2
node7483.members.0.js
species
34816

order
1489939
69

69
1204718
family

genus
69
8163

node7487.members.0.js
species
315492
69

48
1489874
clade

order
48
8064

family
48
8065

subfamily
48
390319

289381
48
genus

48
390379
node7493.members.0.js
species

1489904
clade
node7494.members.0.js
268
3

no rank
1489905
9

30917
1
family

320381
1
genus

1
392546
node7498.members.0.js
species

family
30876
7

genus
270536
7

species
node7501.members.0.js
941984
7

1
8184
family

8186
1
genus

species
node7504.members.0.js
8187
1

order
1489906
6

8243
6
family

genus
8244
6

6
8245
species
node7508.members.0.js

order
8252
94

30942
suborder
node7510.members.0.js
94
1

30948
7
family

genus
7
28828

28829
node7513.members.0.js
species
7

11
8256
family

genus
node7515.members.0.js
8266
1
11

node7516.members.0.js
species
8267
3

7
195615
node7517.members.0.js
species

52902
49
family

genus
49
52903

49
species
node7520.members.0.js
52904

18
30947
family

subfamily
18
603456

genus
18
106173

18
node7524.members.0.js
species
244447

family
171414
8

8
8254
genus

8255
species
node7527.members.0.js
8

156
1489907
order

69
173245
family

173246
69
genus

69
173247
species
node7531.members.0.js

87
8157
family

genus
36211
78

36212
node7534.members.0.js
species
78

9
1
8160
genus
node7535.members.0.js

species
8
302047

8
subspecies
node7537.members.0.js
1841481

clade
108
181483

order
108
1490028

family
108
47697

108
47698
subfamily

108
47699
genus

108
586833
species
node7543.members.0.js

1489838
59
clade

59
1489841
clade

clade
1489843
59

59
8043
order

1489845
59
suborder

family
8045
59

8048
59
genus

8049
species
node7551.members.0.js
59

clade
122
1489343

clade
31089
122

122
41712
order

27723
115
family

genus
114
27726

node7557.members.0.js
species
113540
114

genus
113543
1

node7559.members.0.js
species
113544
1

family
7
31092

7
91732
genus

7
species
node7562.members.0.js
1676925

infraclass
9
1489100

7914
9
order

9
7915
family

9
7916
genus

9
species
node7567.members.0.js
7918

1338366
10
class

10
8288
order

10
1
8289
node7570.members.0.js
family

8290
3
genus

node7572.members.0.js
species
55291
3

6
27686
genus

27687
node7574.members.0.js
species
6

1
5238
superclass
node7575.members.0.js
8287

clade
node7576.members.0.js
1338369
2
5230

5
7878
subclass

superorder
118077
5

order
5
2823314

suborder
5
7879

family
7884
5

genus
5
7885

7888
species
node7583.members.0.js
5

5223
32
32523
clade
node7584.members.0.js

4987
121
32524
node7585.members.0.js
clade

8457
1124
clade

7
1124
clade
node7587.members.0.js
32561

1329799
1009
clade

subclass
2841271
40

8459
40
order

suborder
8464
40

superfamily
1579275
5

5
34907
family

5
204969
genus

species
node7595.members.0.js
13735
5

4
35
node7596.members.0.js
clade
1579337

clade
7
1579336

superfamily
7
27791

4
8465
family

4
8468
genus

4
8469
species
node7601.members.0.js

family
27792
3

genus
27793
3

3
node7604.members.0.js
species
27794

24
8486
superfamily

family
328320
5

subfamily
328321
5

5
74925
genus

5
node7609.members.0.js
species
74926

15
4
8476
node7610.members.0.js
family

genus
85610
3

species
158814
3

3
subspecies
node7613.members.0.js
2587831

genus
6
8477

8479
node7615.members.0.js
species
6
2

subspecies
node7616.members.0.js
8478
4

2
34902
genus

species
34903
2

31138
node7619.members.0.js
subspecies
2

4
8487
family

genus
904181
4

4
1137846
no rank

species
node7623.members.0.js
106734
4

969
5
8492
node7624.members.0.js
clade

10
5
1294634
order
node7625.members.0.js

family
1294636
5

34915
5
subfamily

8495
5
genus

38654
node7629.members.0.js
species
3

2
species
node7630.members.0.js
8496

clade
436486
954

954
436489
clade

954
436491
clade

954
436492
clade

11
954
class
node7635.members.0.js
8782

810
25
8825
node7636.members.0.js
infraclass

order
1
8936

family
57386
1

genus
1
57420

species
node7640.members.0.js
57421
1

30458
9
order

30462
3
family

3
56312
genus

species
node7644.members.0.js
56313
3

family
6
30459

genus
6
126785

194338
species
node7647.members.0.js
6

2
56301
order

family
2
56302

genus
2
56303

121530
node7651.members.0.js
species
2

order
4
57384

family
57385
4

genus
4
57409

4
node7655.members.0.js
species
57412

8940
4
order

family
8941
4

4
33592
genus

species
node7659.members.0.js
55661
4

order
8906
6

family
8917
5

5
8918
genus

5
species
node7663.members.0.js
198806

1
8907
family

1
50391
genus

1
node7666.members.0.js
species
50402

clade
2607030
61

60
8902
order

60
48283
family

48286
58
subfamily

48284
58
genus

species
node7672.members.0.js
111811
58

2
1854010
genus

2
279965
species
node7674.members.0.js

order
1
8892

9242
1
family

9243
1
genus

species
node7678.members.0.js
9244
1

241
26
9126
node7679.members.0.js
order

superfamily
175121
60

5
9133
family

subfamily
5
37599

5
9134
genus

species
node7684.members.0.js
9135
5

7
37611
family

subfamily
7
40155

4
59728
genus

node7688.members.0.js
species
59729
4

genus
40156
3

species
3
40157

3
subspecies
node7691.members.0.js
299123

family
36256
45

genus
45
45806

45807
45
species

45
subspecies
node7695.members.0.js
1094192

family
3
9158

2823048
2
genus

2
species
node7698.members.0.js
356909

9159
1
genus

1
9160
node7700.members.0.js
species

1
1729112
family

1
44387
genus

node7703.members.0.js
species
44394
1

9170
1
family

1
9171
genus

1
node7706.members.0.js
species
9172

family
1
36291

36292
1
genus

node7709.members.0.js
species
59894
1

superfamily
63
2116661

family
43148
13

43149
13
genus

13
node7713.members.0.js
species
43150

36270
50
family

subfamily
50
330750

genus
39620
50

48156
50
species

126889
node7718.members.0.js
subspecies
50

family
9183
51

36283
50
genus

37610
node7721.members.0.js
species
50

1
9184
genus

91951
species
node7723.members.0.js
1

17
5
114313
family
node7724.members.0.js

genus
6
321397

321398
node7726.members.0.js
species
6

3
88178
genus

649802
node7728.members.0.js
species
3

genus
196026
3

node7730.members.0.js
species
296741
3

10
192204
superfamily

family
28725
10

6
10
genus
node7733.members.0.js
30420

2
species
node7734.members.0.js
1196302

node7735.members.0.js
species
68294
1

181096
1
species

1
932674
node7737.members.0.js
subspecies

1
4
family
node7738.members.0.js
9153

genus
1
156562

156563
species
node7740.members.0.js
1

181118
1
genus

1
species
node7742.members.0.js
181119

genus
9154
1

1
species
node7744.members.0.js
9157

family
node7745.members.0.js
400783
2

28728
5
family

genus
28729
5

node7748.members.0.js
species
164674
5

order
8948
6

6
8949
family

2
6
node7751.members.0.js
genus
8952

node7752.members.0.js
species
120794
3

8954
node7753.members.0.js
species
1

2
9219
order

2
9220
family

2
1517830
genus

118200
species
node7757.members.0.js
2

3
8920
order

3
56295
family

56296
3
genus

3
240206
species
node7761.members.0.js

8929
64
order

family
node7763.members.0.js
8930
1
64

61
36242
genus

177155
species
node7765.members.0.js
61

2
8931
genus

species
node7767.members.0.js
8932
2

9108
6
order

54375
3
family

3
172679
genus

species
node7771.members.0.js
187382
3

54381
1
family

54382
1
genus

node7774.members.0.js
species
54383
1

9109
2
family

2
30413
genus

species
925459
2

2
100784
subspecies
node7778.members.0.js

order
168
2558200

family
167
56259

8955
subfamily
node7781.members.0.js
167
1

genus
166
8960

species
166
8962

166
223781
node7784.members.0.js
subspecies

family
8922
1

33615
1
genus

node7787.members.0.js
species
33616
1

22
1
9223
order
node7788.members.0.js

9224
12
family

genus
10
35547

2489341
node7791.members.0.js
species
10

2
176042
genus

species
node7793.members.0.js
176057
2

family
9
1545690

genus
9
13145

13146
species
node7796.members.0.js
9

order
9
9205

1
9206
family

1
9207
genus

9209
node7800.members.0.js
species
1

family
30445
3

37044
3
genus

3
97097
node7803.members.0.js
species

family
1
8899

genus
1
56073

species
node7806.members.0.js
188379
1

30444
1
family

genus
33617
1

36300
node7809.members.0.js
species
1

family
3
33574

128389
3
genus

species
node7812.members.0.js
128390
3

order
1
57379

family
1
57380

genus
175829
1

175835
1
species

1
175836
node7817.members.0.js
subspecies

9230
2
order

family
9231
2

1
9237
genus

1
species
node7821.members.0.js
9238

genus
1
9232

1
node7823.members.0.js
species
9233

superorder
node7824.members.0.js
1549675
1
173

order
8826
79

family
node7826.members.0.js
8830
6
79

2068721
1
subfamily

1
30384
genus

1
node7829.members.0.js
species
219594

subfamily
4
2068722

8867
3
genus

8869
species
node7832.members.0.js
3

genus
1
8842

species
1
8845

381198
subspecies
node7835.members.0.js
1

subfamily
68
2068716

genus
2
8883

node7838.members.0.js
species
8884
2

genus
8835
66

66
8839
node7840.members.0.js
species

1
93
order
node7841.members.0.js
8976

family
8990
4

4
8995
genus

4
8996
node7844.members.0.js
species

9005
family
node7845.members.0.js
88
1

subfamily
9072
15

genus
9030
13

13
species
node7848.members.0.js
9031

genus
2
9053

9054
node7850.members.0.js
species
2

466544
5
subfamily

genus
5
9090

93934
node7853.members.0.js
species
5

466552
61
subfamily

genus
61
9102

node7856.members.0.js
species
9103
61

6
466585
subfamily

5
9001
genus

9002
species
node7859.members.0.js
5

genus
30409
1

30410
node7861.members.0.js
species
1

1956187
1
order

54378
1
family

genus
54379
1

1
node7865.members.0.js
species
54380

superorder
node7866.members.0.js
8783
2
133

order
4
8784

4
8788
family

genus
4
8789

8790
species
node7870.members.0.js
4

8819
124
order

family
124
8820

node7873.members.0.js
genus
8821
2
124

species
2696672
122

subspecies
node7875.members.0.js
202946
122

8798
3
order

8799
3
family

8800
3
genus

species
8801
3

3
node7880.members.0.js
subspecies
441894

class
8504
108

8509
108
order

clade
1329961
108

1329950
104
clade

1329912
104
clade

1329911
node7886.members.0.js
clade
92
3

8548
4
infraorder

clade
1330544
4

superfamily
4
1329920

family
4
8555

8556
4
genus

61221
species
node7892.members.0.js
4

8511
28
suborder

2024743
9
family

subfamily
42425
9

genus
9
8518

9
species
node7897.members.0.js
8520

83232
5
clade

family
81953
5

5
145349
subfamily

52201
5
genus

5
node7902.members.0.js
species
103695

14
2024747
family

genus
14
28376

14
species
node7905.members.0.js
28377

8570
infraorder
node7906.members.0.js
57
1

34979
11
superfamily

family
34984
11

genus
37579
11

11
176946
node7910.members.0.js
species

superfamily
node7911.members.0.js
34989
14
45

family
10
8578

subfamily
8
169863

201800
8
genus

species
node7915.members.0.js
94885
8

subfamily
169862
2

genus
34999
2

node7918.members.0.js
species
35005
2

family
8689
14

14
2
8710
subfamily
node7920.members.0.js

4
8711
genus

8720
node7922.members.0.js
species
4

7
8728
genus

7
88082
species
node7924.members.0.js

genus
103943
1

103944
node7926.members.0.js
species
1

family
7
8602

subfamily
node7928.members.0.js
42167
2
7

8672
1
genus

1
species
node7930.members.0.js
8673

4
8661
genus

4
species
node7932.members.0.js
8663

clade
12
1329976

12
1329975
clade

family
8522
12

162266
node7936.members.0.js
subfamily
12
5

2
42163
genus

node7938.members.0.js
species
64176
2

genus
141678
5

8524
species
node7940.members.0.js
5

4
8560
infraorder

superfamily
1330539
1

311552
1
family

genus
71026
1

1
node7945.members.0.js
species
288420

8561
3
family

subfamily
385256
3

genus
3
8565

3
node7949.members.0.js
species
146911

1
3742
node7950.members.0.js
class
40674

3725
4
32525
node7951.members.0.js
clade

node7952.members.0.js
clade
9347
10
3653

3600
56
1437010
node7953.members.0.js
clade

23
1299
node7954.members.0.js
superorder
314145

order
8
9971

9972
8
family

9973
node7957.members.0.js
genus
8
2

2
9974
species
node7958.members.0.js

4
143292
species
node7959.members.0.js

178
9397
order

suborder
30559
19

9398
19
family

1
19
subfamily
node7963.members.0.js
77225

9401
genus
node7964.members.0.js
7
5

node7965.members.0.js
species
143291
2

genus
9406
11

node7967.members.0.js
species
9407
11

30560
node7968.members.0.js
suborder
159
3

family
186994
7

genus
58068
7

186990
node7971.members.0.js
species
7

5
9436
family

27621
5
genus

5
27622
species
node7974.members.0.js

9431
family
node7975.members.0.js
104
2

subfamily
5
981671

genus
9432
5

5
species
node7978.members.0.js
291302

genus
1
29077

1
node7980.members.0.js
species
29078

27671
genus
node7981.members.0.js
82
1

59472
node7982.members.0.js
species
1

species
node7983.members.0.js
59474
80

9434
node7984.members.0.js
genus
14
1

1
51298
species
node7985.members.0.js

12
109478
species
node7986.members.0.js

24
5
9415
family
node7987.members.0.js

8
2
40234
node7988.members.0.js
subfamily

genus
27659
3

192404
species
node7990.members.0.js
3

3
9416
genus

3
9417
species
node7992.members.0.js

subfamily
40237
3

genus
9429
3

node7995.members.0.js
species
9430
3

subfamily
40238
8

node7997.members.0.js
genus
9422
2
8

9423
node7998.members.0.js
species
2

4
node7999.members.0.js
species
89673

family
58055
16

16
186995
subfamily

49442
node8002.members.0.js
genus
16
5

5
89399
node8003.members.0.js
species

6
species
node8004.members.0.js
59479

node8005.members.0.js
order
91561
4
492

suborder
9845
388

node8007.members.0.js
infraorder
35500
30
388

family
node8008.members.0.js
9895
6
216

subfamily
9963
69

3
9922
genus

9925
node8011.members.0.js
species
3

66
1
9935
genus
node8012.members.0.js

node8013.members.0.js
species
9940
2

species
63
37174

63
subspecies
node8015.members.0.js
112262

27592
node8016.members.0.js
subfamily
137
2

node8017.members.0.js
genus
9903
57
130

species
node8018.members.0.js
72004
22

51
node8019.members.0.js
species
9913

3
9918
genus

89462
species
node8021.members.0.js
3

1
303929
genus

1
303930
node8023.members.0.js
species

1
9900
genus

species
9901
1

1
subspecies
node8026.members.0.js
43346

9959
4
subfamily

genus
9957
4

59534
species
node8029.members.0.js
4

node8030.members.0.js
family
9850
1
142

9881
5
subfamily

genus
5
9871

5
9874
species

5
subspecies
node8034.members.0.js
9880

subfamily
136
34878

136
9859
genus

136
node8037.members.0.js
species
9860

24
2653789
suborder

9721
node8039.members.0.js
infraorder
24
4

2
9761
parvorder

2
9765
family

2
9766
genus

9767
1
species

1
subspecies
node8044.members.0.js
310752

9771
species
node8045.members.0.js
1

node8046.members.0.js
parvorder
9722
2
18

family
node8047.members.0.js
9747
1
2

genus
9748
1

9749
node8049.members.0.js
species
1

1
9740
family

1
9741
genus

42100
node8052.members.0.js
species
1

family
7
9750

genus
7
9753

9755
species
node8055.members.0.js
7

node8056.members.0.js
family
9726
4
6

2
9729
genus

2
node8058.members.0.js
species
9731

35497
67
suborder

9821
67
family

67
9822
genus

31
67
species
node8062.members.0.js
9823

415978
node8063.members.0.js
subspecies
36

suborder
9
9834

9835
9
family

2
7
genus
node8066.members.0.js
9836

3
species
node8067.members.0.js
9837

2
9838
node8068.members.0.js
species

genus
2
30539

2
species
node8070.members.0.js
30538

order
23
9362

9363
6
family

subfamily
30577
6

9364
5
genus

5
9365
node8075.members.0.js
species

9367
1
genus

1
node8077.members.0.js
species
9368

6
9376
family

6
183663
subfamily

genus
6
9379

node8081.members.0.js
species
42254
6

9373
11
family

6
9374
genus

6
50954
species
node8084.members.0.js

5
143301
genus

143302
species
node8086.members.0.js
5

549
1
33554
order
node8087.members.0.js

379584
node8088.members.0.js
suborder
434
7

family
9608
225

genus
224
9611

224
193
9612
node8091.members.0.js
species

node8092.members.0.js
subspecies
9615
31

9625
1
genus

node8094.members.0.js
species
494514
1

12
3
9709
node8095.members.0.js
family

genus
9717
2

2
species
node8097.members.0.js
9720

3
9710
genus

3
node8099.members.0.js
species
9711

9712
2
genus

2
species
node8101.members.0.js
9713

genus
node8102.members.0.js
9714
2

family
node8103.members.0.js
9632
1
9

node8104.members.0.js
genus
9639
2
4

node8105.members.0.js
species
9643
1

1
species
node8106.members.0.js
29073

genus
9645
4

node8108.members.0.js
species
9646
4

family
9702
1

genus
34883
1

1
34884
species
node8111.members.0.js

180
22
9655
family
node8112.members.0.js

1008252
63
subfamily

genus
63
9661

species
node8115.members.0.js
9662
63

6
14
subfamily
node8116.members.0.js
169418

9665
genus
node8117.members.0.js
8
1

2
9668
species

9669
node8119.members.0.js
subspecies
2

5
36723
node8120.members.0.js
species

subfamily
169417
81

genus
81
9656

species
node8123.members.0.js
9657
81

379583
node8124.members.0.js
suborder
114
1

9681
family
node8125.members.0.js
98
3

subfamily
338152
91

13124
1
genus

61383
node8128.members.0.js
species
1

genus
9682
89

89
9685
species
node8130.members.0.js

146712
1
genus

1608482
node8132.members.0.js
species
1

subfamily
1
338151

genus
1
32535

32536
species
node8135.members.0.js
1

338153
3
subfamily

3
9688
genus

9694
node8138.members.0.js
species
1

species
node8139.members.0.js
9689
2

14
9697
family

14
37031
genus

37032
node8142.members.0.js
species
14

9676
1
family

95911
1
genus

1
species
node8145.members.0.js
95912

order
9787
26

9788
24
family

8
24
genus
node8148.members.0.js
9789

6
9796
node8149.members.0.js
species

9793
species
node8150.members.0.js
10

2
9803
family

genus
2
9806

species
9807
2

73337
subspecies
node8154.members.0.js
2

superorder
node8155.members.0.js
314146
13
2245

clade
314147
987

980
5
9989
order
node8157.members.0.js

33553
170
suborder

1
170
node8159.members.0.js
family
55153

156
9991
subfamily

30
156
tribe
node8161.members.0.js
337752

2895726
29
genus

29
30640
node8163.members.0.js
species

genus
10001
97

55149
node8165.members.0.js
species
97

subfamily
337726
13

13
6
337730
tribe
node8167.members.0.js

genus
4
9992

93162
node8169.members.0.js
species
2

species
2
9993

2
subspecies
node8171.members.0.js
9994

3
1141640
genus

3
species
node8173.members.0.js
43179

node8174.members.0.js
suborder
33550
2
36

family
5
10158

genus
5
10159

5
node8177.members.0.js
species
10160

10150
5
family

5
10151
genus

34839
node8180.members.0.js
species
5

family
10167
16

genus
10180
8

8
10181
node8183.members.0.js
species

genus
8
423606

885580
species
node8185.members.0.js
8

family
8
10139

10140
8
genus

10141
species
node8188.members.0.js
8

758
1963758
suborder

superfamily
1963761
4

family
30648
4

subfamily
35737
4

4
48867
genus

4
51337
species
node8194.members.0.js

754
8
337687
node8195.members.0.js
clade

node8196.members.0.js
family
337677
8
177

subfamily
8
10026

genus
5
10035

10036
species
node8199.members.0.js
5

3
10028
genus

node8201.members.0.js
species
10029
3

subfamily
node8202.members.0.js
39087
1
51

41
10049
genus

node8204.members.0.js
species
1047088
41

10053
node8205.members.0.js
genus
9
4

2
79684
node8206.members.0.js
species

3
111838
node8207.members.0.js
species

337963
subfamily
node8208.members.0.js
110
1

7
10040
genus

species
7
10042

230844
subspecies
node8211.members.0.js
7

102
38667
genus

species
node8213.members.0.js
38674
102

family
6
337664

subfamily
6
10061

6
30636
genus

6
node8217.members.0.js
species
1026970

563
5
10066
node8218.members.0.js
family

309
326408
subfamily

genus
node8220.members.0.js
10067
33
309

83762
node8221.members.0.js
species
12

237
60746
species
node8222.members.0.js

12
node8223.members.0.js
species
60744

15
node8224.members.0.js
species
83527

39107
subfamily
node8225.members.0.js
246
5

61153
10
genus

10
61156
species
node8227.members.0.js

node8228.members.0.js
genus
10114
2
14

species
node8229.members.0.js
10116
11

10117
species
node8230.members.0.js
1

genus
4
121588

4
491861
species
node8232.members.0.js

10088
208
genus

862507
node8234.members.0.js
subgenus
208
7

201
node8235.members.0.js
species
10090

genus
30639
4

4
species
node8237.members.0.js
35658

genus
1
71161

node8239.members.0.js
species
2107264
1

10045
3
subfamily

genus
10046
3

3
node8242.members.0.js
species
10047

suborder
11
1963757

3
10015
family

subfamily
3
38662

genus
10016
3

10020
node8247.members.0.js
species
1

2
species
node8248.members.0.js
105255

29132
8
family

10184
8
genus

51338
species
node8251.members.0.js
8

9975
7
order

family
9976
5

5
1
9977
node8254.members.0.js
genus

4
node8255.members.0.js
species
130825

9979
2
family

genus
9984
2

9986
species
node8258.members.0.js
2

order
3
9392

family
3
9393

genus
9394
3

node8262.members.0.js
species
37347
1

2
species
node8263.members.0.js
246437

order
1237
9443

suborder
376913
1217

infraorder
node8266.members.0.js
314293
1
1212

1188
8
9526
node8267.members.0.js
parvorder

superfamily
node8268.members.0.js
314295
1
1136

4
1
9577
node8269.members.0.js
family

genus
325165
1

1
node8271.members.0.js
species
61853

genus
9578
2

2
node8273.members.0.js
species
81572

node8274.members.0.js
family
9604
7
1131

subfamily
607660
6

genus
6
9599

6
9601
node8277.members.0.js
species

65
1118
node8278.members.0.js
subfamily
207598

9592
3
genus

3
2
9593
species
node8280.members.0.js

1
9595
subspecies
node8281.members.0.js

9605
1032
genus

1032
9606
node8283.members.0.js
species

18
9596
genus

18
9598
node8285.members.0.js
species

superfamily
314294
44

2
44
family
node8287.members.0.js
9527

9528
node8288.members.0.js
subfamily
37
2

genus
1
9567

9568
node8290.members.0.js
species
1

9539
genus
node8291.members.0.js
21
1

16
species
node8292.members.0.js
9544

9545
species
node8293.members.0.js
3

9541
node8294.members.0.js
species
1

genus
1
9529

9531
node8296.members.0.js
species
1

genus
5
392815

2
60711
species
node8298.members.0.js

3
node8299.members.0.js
species
9534

9554
7
genus

7
9555
species
node8301.members.0.js

subfamily
5
9569

genus
542827
1

1
61622
node8304.members.0.js
species

genus
54136
4

4
node8306.members.0.js
species
54180

3
23
parvorder
node8307.members.0.js
9479

376918
7
family

genus
9504
7

37293
node8310.members.0.js
species
7

9498
node8311.members.0.js
family
13
1

subfamily
3
378850

genus
3
9520

species
node8314.members.0.js
27679
1
3

2
subspecies
node8315.members.0.js
39432

subfamily
9480
7

genus
9481
7

subgenus
1965096
7

9483
node8319.members.0.js
species
7

2
1
38070
node8320.members.0.js
subfamily

1
1532884
genus

9515
species
node8322.members.0.js
1

376912
5
infraorder

family
9475
5

genus
1868481
5

5
node8326.members.0.js
species
1868482

suborder
20
376911

2
16
node8328.members.0.js
infraorder
376915

family
30615
7

genus
13149
7

7
30608
species
node8331.members.0.js

family
6
9445

9446
6
genus

9447
node8334.members.0.js
species
6

family
1
30599

30600
1
genus

1
379532
node8337.members.0.js
species

infraorder
376917
4

family
4
40297

4
30610
genus

4
species
node8341.members.0.js
30611

30656
5
order

5
30657
family

482536
5
genus

5
482537
node8345.members.0.js
species

superorder
14
9348

order
948951
6

family
6
9359

9360
6
genus

6
9361
node8350.members.0.js
species

order
948950
8

suborder
948953
8

8
227508
family

genus
9357
8

27675
node8355.members.0.js
species
8

311790
29
superorder

28734
4
order

28735
4
family

genus
4
28736

28737
node8360.members.0.js
species
4

order
11
9774

9775
11
family

9776
11
genus

9778
11
species

11
subspecies
node8365.members.0.js
127582

9815
2
order

2
9816
family

9817
2
genus

species
9818
2

1230840
node8370.members.0.js
subspecies
2

7
9369
family

7
176113
subfamily

7
9370
genus

9371
species
node8374.members.0.js
7

family
9389
5

subfamily
745257
5

5
185452
genus

node8378.members.0.js
species
185453
5

9263
node8379.members.0.js
clade
68
1

order
38607
10

10
30660
family

genus
33561
10

10
33562
species
node8383.members.0.js

order
13
38605

13
9265
family

subfamily
13
126287

genus
126288
7

191870
node8388.members.0.js
species
7

genus
6
13615

13616
species
node8390.members.0.js
6

order
19
38608

19
9277
family

19
9304
genus

node8394.members.0.js
species
9305
19

8
25
order
node8395.members.0.js
38609

family
9307
2

2
1960649
genus

node8398.members.0.js
species
9315
2

family
9335
3

genus
3
9336

species
node8401.members.0.js
9337
3

38624
3
family

genus
38625
3

species
node8404.members.0.js
38626
3

9
9338
family

9
29138
genus

9
species
node8407.members.0.js
29139

9254
16
clade

16
1
9255
node8409.members.0.js
order

family
9259
14

9260
14
genus

9261
node8412.members.0.js
species
14

1
9256
family

9257
1
genus

species
node8415.members.0.js
9258
1

class
8292
204

order
8445
154

30380
85
family

194407
85
genus

85
species
node8420.members.0.js
194408

12
1277737
family

genus
264009
12

node8423.members.0.js
species
1415580
12

57
264006
family

genus
260994
57

57
species
node8426.members.0.js
260995

50
41666
superorder

50
8342
order

30319
30
superfamily

30
8352
family

30
8360
subfamily

genus
8353
30

subgenus
14
262014

14
node8434.members.0.js
species
8355

16
8363
subgenus

16
8364
node8436.members.0.js
species

superfamily
8431
1

1
161696
family

1
265034
genus

1
node8440.members.0.js
species
265035

suborder
8416
19

8417
7
superfamily

family
7
8382

genus
node8444.members.0.js
8383
6
7

1
species
node8445.members.0.js
8384

30352
12
superfamily

8397
4
family

genus
4
8399

121175
4
subgenus

4
species
node8450.members.0.js
8407

family
1
27577

160964
1
subfamily

1
27578
genus

462320
species
node8454.members.0.js
1

7
685122
family

7
1191378
subfamily

7
120497
genus

125878
node8458.members.0.js
species
7

clade
7
118072

7894
7
order

7
7895
family

7896
7
genus

7
node8463.members.0.js
species
7897

phylum
165
7586

clade
133550
6

class
6
35069

6
35070
subclass

order
7581
6

family
6
106121

1529449
6
subfamily

6
1529434
genus

species
node8472.members.0.js
1529436
6

clade
159
133551

superclass
13
7624

7705
1
class

1
7681
subclass

1
7682
order

1
7687
family

307971
1
genus

307972
node8480.members.0.js
species
1

class
12
7625

subclass
12
7638

superorder
12
7674

order
31184
8

1
39211
family

39214
1
genus

161071
species
node8487.members.0.js
1

family
31185
7

genus
7
7652

7654
node8490.members.0.js
species
7

order
4
2785018

infraorder
7675
4

family
31181
4

genus
7664
4

4
node8495.members.0.js
species
7668

superclass
7587
146

class
7588
146

superorder
41243
33

41166
node8499.members.0.js
order
33
1

133432
16
family

genus
133433
16

16
node8502.members.0.js
species
133434

family
7592
16

genus
35076
16

16
46514
species
node8505.members.0.js

superorder
113
41242

order
7599
113

7600
113
family

57
7601
genus

57
7604
node8510.members.0.js
species

7608
56
genus

56
7609
species
node8512.members.0.js

clade
node8513.members.0.js
33317
376
18523

clade
node8514.members.0.js
2697495
2
1008

993
1
1206795
clade
node8515.members.0.js

34
10205
phylum

34
10206
class

order
34
10207

suborder
34
193205

193206
34
superfamily

34
192924
family

genus
34
95169

node8523.members.0.js
species
95170
34

phylum
10
6340

class
1
6341

subclass
1
6427

order
1
41798

family
1
47116

1775492
no rank
node8529.members.0.js
1

9
42113
class

8
55824
subclass

order
8
6406

8
6407
family

genus
8
6411

8
species
node8535.members.0.js
6412

subclass
1
6381

2803884
1
order

suborder
1
6391

family
1
6392

1046325
1
subfamily

genus
1
6397

no rank
1050931
1

node8543.members.0.js
species
6398
1

33313
1
phylum

order
41320
1

1
41321
suborder

41372
1
family

68038
1
genus

1
2614562
species
node8549.members.0.js

phylum
6157
254

6178
94
class

subclass
6179
94

order
43
6180

superfamily
31244
43

family
43
31245

genus
node8556.members.0.js
6181
1
43

4
node8557.members.0.js
species
6182

2
species
node8558.members.0.js
6185

species
node8559.members.0.js
6183
36

order
27871
49

suborder
49
27872

49
1776223
superfamily

family
49
73421

49
57077
genus

57078
species
node8565.members.0.js
49

2
6193
order

suborder
2
6194

6196
2
family

6197
2
genus

6198
node8570.members.0.js
species
2

class
5
147100

5
166126
clade

5
6159
order

suborder
3
1292243

superfamily
3
1292253

family
31262
3

genus
55270
3

species
node8578.members.0.js
79327
3

suborder
2
55267

superfamily
166236
2

family
2
46764

2
1292246
subfamily

1070817
2
genus

2
species
node8584.members.0.js
2597323

6199
155
class

subclass
6200
155

order
43
6201

6208
2
family

genus
6209
2

species group
2212966
2

6210
node8591.members.0.js
species
2

family
6214
41

41
6215
genus

species
node8594.members.0.js
85433
41

1224679
112
order

112
28843
family

46580
112
genus

99802
species
node8598.members.0.js
112

6217
30
phylum

class
6218
30

order
30
6219

30
6222
family

genus
30
6223

30
species
node8604.members.0.js
88925

6
7568
phylum

115360
6
subphylum

class
6
115361

7570
6
order

6
115362
superfamily

33491
6
family

genus
6
7571

7574
node8612.members.0.js
species
6

6447
node8613.members.0.js
phylum
657
6

6448
node8614.members.0.js
class
304
1

216305
17
subclass

clade
216307
17

clade
977775
10

2
120490
superorder

node8619.members.0.js
order
6527
1
2

216366
1
clade

superfamily
87862
1

family
1
133184

1980934
1
subfamily

1
1980892
genus

1980912
node8625.members.0.js
species
1

clade
8
977779

superfamily
8
216441

family
8
6524

genus
8
6525

8
species
node8630.members.0.js
6526

clade
2836391
7

6497
7
order

7
216318
superfamily

6498
7
family

7
6499
genus

7
node8636.members.0.js
species
6500

subclass
69675
61

56
146277
superfamily

56
6462
family

genus
6463
56

node8641.members.0.js
species
88005
56

superfamily
5
216260

5
69676
family

72691
5
genus

5
225164
species
node8645.members.0.js

8
69555
subclass

order
75116
8

8
6475
superfamily

8
54973
family

genus
8
72702

8
node8651.members.0.js
species
400727

10
2219556
subclass

10
216274
superfamily

10
55007
family

1735270
10
genus

10
node8656.members.0.js
species
1735272

216275
207
subclass

order
2315720
193

216285
193
superfamily

family
6466
193

subfamily
node8661.members.0.js
1955429
2
193

genus
78
1093071

78
node8663.members.0.js
species
1620919

113
2072689
genus

113
216125
node8665.members.0.js
species

order
2315723
14

216276
14
superfamily

6451
14
family

14
6452
genus

6454
species
node8670.members.0.js
14

6544
335
class

subclass
2785011
335

206
6545
clade

order
106218
86

superfamily
106219
86

86
6566
family

6578
78
genus

species
node8678.members.0.js
6579
78

genus
8
186466

6573
node8680.members.0.js
species
8

order
6562
120

superfamily
120
98302

family
120
6563

genus
6564
120

110
29159
node8685.members.0.js
species

6565
species
node8686.members.0.js
10

clade
129
6599

infraclass
735337
129

superorder
2785015
129

clade
2908833
129

order
6580
11

74489
11
superfamily

family
11
6592

genus
11
6595

species
node8695.members.0.js
6596
11

2783445
118
order

118
98297
superfamily

family
118
61354

118
457750
genus

118
species
node8700.members.0.js
2589376

class
12
6605

12
6606
subclass

1
215450
superorder

551290
1
order

suborder
1
551347

6615
1
family

1
1051063
genus

1
species
node8708.members.0.js
1051067

215451
11
superorder

order
11
6638

11
6646
suborder

6647
11
family

1
11
genus
node8713.members.0.js
6643

37653
species
node8714.members.0.js
8

2
2607531
node8715.members.0.js
species

clade
2697496
13

10190
13
phylum

class
2816136
13

44578
node8719.members.0.js
subclass
13
1

order
12
104779

family
12
104780

genus
104781
12

104782
node8723.members.0.js
species
12

96
17139
node8724.members.0.js
clade
1206794

clade
16810
88770

6656
node8726.members.0.js
phylum
16810
120

197563
node8727.members.0.js
clade
11707
2

11704
173
197562
node8728.members.0.js
clade

subphylum
6657
543

superclass
110
2172819

class
110
6670

43953
110
subclass

order
110
84318

35
116574
suborder

116575
35
superfamily

35
69351
family

genus
35
69354

35
species
node8738.members.0.js
69355

suborder
84328
72

72
84329
superfamily

family
72
43954

50
399044
genus

50
399045
node8743.members.0.js
species

22
163713
genus

22
node8745.members.0.js
species
163714

suborder
116576
3

superfamily
182511
3

3
163706
family

1164977
3
genus

1164978
species
node8750.members.0.js
3

401
2172821
superclass

239
6681
class

subclass
72041
239

superorder
233
6682

order
6683
233

suborder
211
6692

1
6738
infraorder

superfamily
1
115588

1
1124584
family

genus
1
136693

species
node8761.members.0.js
306053
1

infraorder
20
6712

superfamily
6724
13

6725
12
family

subfamily
72430
12

genus
12
6726

12
6728
species
node8767.members.0.js

family
6713
1

genus
1
6714

6715
node8770.members.0.js
species
1

superfamily
37849
7

7
6704
family

genus
7
6705

6706
species
node8774.members.0.js
7

6694
185
infraorder

115583
1
superfamily

family
6701
1

6702
1
genus

1
666362
species
node8779.members.0.js

superfamily
115580
184

184
6695
family

genus
183
6696

183
species
node8783.members.0.js
159736

1
117982
genus

1
647170
node8785.members.0.js
species

5
6752
infraorder

no rank
116704
5

116706
5
no rank

superfamily
1
6804

family
1
6805

1
6806
genus

node8792.members.0.js
species
6807
1

superfamily
6774
4

4
6757
family

80835
3
genus

3
node8796.members.0.js
species
210409

genus
557253
1

1
1888338
node8798.members.0.js
species

suborder
22
6684

22
111520
superfamily

family
22
6685

3
22
genus
node8802.members.0.js
133894

6689
node8803.members.0.js
species
12

species
node8804.members.0.js
27405
3

4
6687
species
node8805.members.0.js

superorder
6820
6

6
6821
order

suborder
1732196
5

infraorder
5
1732206

superfamily
5
199476

5
199477
family

genus
5
199487

5
node8813.members.0.js
species
294128

1
2692238
suborder

infraorder
2692343
1

parvorder
1
2692351

superfamily
1
44330

134545
1
family

genus
1
58017

1518452
species
node8820.members.0.js
1

class
125
72037

subclass
6830
125

infraclass
116569
125

superorder
116570
6

6
6833
order

5
88013
family

5
88014
genus

5
88015
species
node8828.members.0.js

163470
1
family

genus
1
749187

749188
node8831.members.0.js
species
1

116571
119
superorder

order
72033
119

family
72034
119

52
217164
genus

52
node8836.members.0.js
species
217165

67
72035
genus

72036
node8838.members.0.js
species
67

class
37
116172

subclass
37
6675

19
6676
infraclass

node8842.members.0.js
superorder
2899756
1
19

order
6
2899670

2899671
6
superfamily

family
6
37806

subfamily
2899660
6

6
637416
genus

species
node8848.members.0.js
1232801
6

order
12
2899683

12
629293
family

12
36136
genus

12
41117
species
node8852.members.0.js

infraclass
37909
18

38011
18
family

genus
18
51649

18
51650
species
node8856.members.0.js

32
6658
class

116557
32
subclass

32
84337
order

6665
32
suborder

infraorder
116561
32

family
32
77658

31
6668
genus

24
6669
node8864.members.0.js
species

6
node8865.members.0.js
species
35525

1
species
node8866.members.0.js
2282166

77650
1
genus

117539
species
node8868.members.0.js
1

10988
1
6960
subphylum
node8869.members.0.js

class
50557
10967

85512
10966
clade

10966
2
7496
subclass
node8872.members.0.js

node8873.members.0.js
infraclass
33340
235
10945

627
1
33341
cohort
node8874.members.0.js

6970
17
clade

order
17
85823

1049657
17
superfamily

no rank
17
1912919

46562
9
family

105801
9
subfamily

60568
9
genus

9
species
node8882.members.0.js
105785

7501
8
family

subfamily
8
127820

tribe
127821
8

7502
8
genus

8
136037
species
node8887.members.0.js

7020
487
order

suborder
487
523712

superfamily
487
213545

213546
487
family

node8892.members.0.js
genus
61471
222
487

61472
species
node8893.members.0.js
29

28
species
node8894.members.0.js
61478

71
629358
node8895.members.0.js
species

31
species
node8896.members.0.js
61476

node8897.members.0.js
species
170557
18

14
node8898.members.0.js
species
629360

node8899.members.0.js
species
61484
12

32
61474
node8900.members.0.js
species

170555
node8901.members.0.js
species
30

27434
1
order

suborder
1
211564

1
211568
superfamily

27437
1
family

13067
genus
node8906.members.0.js
1

order
1
58557

family
244939
1

1
node8909.members.0.js
genus
58612

50622
120
order

120
1
70405
node8911.members.0.js
superfamily

family
62802
90

1
90
subfamily
node8913.members.0.js
143773

35
62803
genus

35
2014036
species
node8915.members.0.js

genus
143721
54

species
node8917.members.0.js
143722
54

family
143727
29

subfamily
29
143771

genus
29
143731

2065413
node8921.members.0.js
species
29

139
33342
cohort

30262
12
order

38130
12
suborder

superfamily
38131
1

family
1291702
1

1
1291689
genus

1
2628252
no rank

node8929.members.0.js
species
1291690
1

45049
11
superfamily

45053
11
family

11
153976
subfamily

genus
45057
5

5
161013
species
node8934.members.0.js

genus
6
45059

133901
species
node8936.members.0.js
6

2
1930602
superorder

order
30259
2

suborder
38127
1

infraorder
239225
1

family
1
50619

239265
1
genus

no rank
1
2624915

1
469673
species
node8944.members.0.js

no rank
792396
1

1
1291676
node8946.members.0.js
species

1
125
node8947.members.0.js
order
7524

33343
13
clade

33345
13
suborder

13
33347
clade

13
33349
clade

13
33351
clade

33357
9
infraorder

superfamily
9
33358

family
160513
9

9
286710
subfamily

genus
631388
8

8
node8958.members.0.js
species
1511221

286705
1
genus

1
286706
species
node8960.members.0.js

infraorder
4
33354

33356
1
superfamily

1
236445
family

genus
2019536
1

no rank
1
2629335

species
node8966.members.0.js
2019537
1

33355
3
superfamily

family
30078
3

genus
30079
3

3
species
node8970.members.0.js
79782

suborder
1955247
10

33361
9
infraorder

36151
superfamily
node8973.members.0.js
9
1

8
33362
family

subfamily
130551
8

genus
8
108930

8
species
node8977.members.0.js
108931

infraorder
1
33365

superfamily
1
33367

family
1
7033

subfamily
140060
1

tribe
1445916
1

1
1445914
genus
node8983.members.0.js

suborder
101
33373

node8985.members.0.js
infraorder
33380
3
68

33385
65
superfamily

family
65
27482

7
55
node8988.members.0.js
subfamily
133076

tribe
40
33386

genus
13163
7

node8991.members.0.js
species
13164
7

7028
30
genus

7029
node8993.members.0.js
species
30

143947
3
genus

node8995.members.0.js
species
143948
3

tribe
node8996.members.0.js
33387
1
8

genus
80764
6

subgenus
464929
6

80765
species
node8999.members.0.js
6

345571
1
genus

742174
species
node9001.members.0.js
1

subfamily
10
805116

genus
143949
10

143950
node9004.members.0.js
species
10

superfamily
9
33377

9
7036
family

subfamily
33379
9

7037
9
genus

9
7038
node9009.members.0.js
species

superfamily
33375
24

1585420
24
family

24
121844
genus

24
node9013.members.0.js
species
121845

542
9944
node9014.members.0.js
cohort
33392

2
7509
order

140693
2
suborder

129369
2
superfamily

family
7511
2

2
476429
subfamily

2
7514
genus

species
node9021.members.0.js
7515
2

85604
node9022.members.0.js
superorder
7094
33

160
30263
order

suborder
160
93875

1683728
160
infraorder

41033
160
superfamily

160
50645
family

subfamily
160
177669

160
177673
tribe

genus
node9030.members.0.js
177674
5
160

1218281
node9031.members.0.js
species
72

node9032.members.0.js
species
1271730
83

order
7088
6901

6901
41191
suborder

41196
6901
infraorder

parvorder
6901
41197

37567
clade
node9037.members.0.js
6901
182

2
599
node9038.members.0.js
clade
104430

superfamily
104435
22

family
115354
22

subfamily
22
287187

genus
287110
22

22
node9043.members.0.js
species
287375

104434
127
superfamily

family
127
106496

106499
127
subfamily

tribe
node9047.members.0.js
301641
1
66

genus
300855
37

species
node9049.members.0.js
301037
37

genus
28
106500

28
1660703
species
node9051.members.0.js

tribe
61
301638

genus
287191
61

61
748215
species
node9054.members.0.js

superfamily
351
37568

351
7139
family

94
65022
subfamily

173709
94
tribe

3
94
node9059.members.0.js
genus
572799

758717
node9060.members.0.js
species
39

52
758706
species
node9061.members.0.js

node9062.members.0.js
subfamily
81687
3
257

126
2
581389
tribe
node9063.members.0.js

59
572704
genus

node9065.members.0.js
species
1100916
59

572852
65
genus

65
node9067.members.0.js
species
1869985

tribe
47
581385

genus
581513
47

47
1594315
node9070.members.0.js
species

tribe
581387
81

82599
46
genus

46
1100963
species
node9073.members.0.js

35
581588
genus

1101027
species
node9075.members.0.js
35

superfamily
39
104437

39
252293
family

39
252294
genus

39
252295
node9079.members.0.js
species

58
104432
superfamily

186108
58
family

subfamily
1556158
58

655084
58
genus

1101072
node9084.members.0.js
species
58

423
5780
clade
node9085.members.0.js
104431

188
37569
superfamily

7089
5
family

subfamily
475327
5

5
4
7090
node9089.members.0.js
genus

1
node9090.members.0.js
species
7091

family
1
7096

subfamily
151316
1

1
151325
genus

1
151326
node9094.members.0.js
species

7128
family
node9095.members.0.js
182
1

subfamily
76
82617

523176
27
tribe

genus
283833
27

644661
species
node9099.members.0.js
27

49
523174
tribe

genus
49
82622

49
987953
node9102.members.0.js
species

100
469321
subfamily

523180
100
tribe

522847
26
genus

522848
node9106.members.0.js
species
26

genus
522835
74

74
species
node9108.members.0.js
522836

82616
5
subfamily

5
523164
tribe

7129
5
genus

7130
species
node9112.members.0.js
5

superfamily
93
104493

family
93
186111

93
186112
subfamily

genus
1594452
59

species
node9117.members.0.js
1594453
59

467773
34
genus

34
467774
node9119.members.0.js
species

82592
677
superfamily

82593
node9121.members.0.js
family
677
3

467
7
82596
subfamily
node9122.members.0.js

genus
80
214128

40
875883
species
node9124.members.0.js

722662
node9125.members.0.js
species
40

722672
29
genus

29
node9127.members.0.js
species
722673

genus
48
393392

species
node9129.members.0.js
688445
48

22
82597
genus

22
species
node9131.members.0.js
934813

genus
104475
39

39
node9133.members.0.js
species
104476

82594
69
genus

node9135.members.0.js
species
82595
69

190355
55
genus

55
species
node9137.members.0.js
190356

genus
704699
21

node9139.members.0.js
species
934829
21

genus
104473
27

node9141.members.0.js
species
104474
27

genus
572919
70

70
934894
node9143.members.0.js
species

subfamily
node9144.members.0.js
104450
1
171

59
934916
genus

59
node9146.members.0.js
species
934917

genus
1369166
1

2677324
1
no rank

1
species
node9149.members.0.js
2484454

65
873511
genus

934888
node9151.members.0.js
species
65

genus
214132
20

20
934839
node9153.members.0.js
species

104458
25
genus

25
104460
species
node9155.members.0.js

104442
36
subfamily

genus
104446
36

104447
node9158.members.0.js
species
36

37570
node9159.members.0.js
superfamily
2275
54

1484
58
7100
family
node9160.members.0.js

95182
108
subfamily

genus
7106
9

3
species
node9163.members.0.js
7108

6
node9164.members.0.js
species
69820

56365
99
genus

58
node9166.members.0.js
species
689058

41
987877
node9167.members.0.js
species

subfamily
node9168.members.0.js
95244
6
300

genus
55056
54

54
55057
node9170.members.0.js
species

genus
320087
62

62
987933
node9172.members.0.js
species

43
320089
genus

43
species
node9174.members.0.js
875884

103830
node9175.members.0.js
genus
100
1

49
species
node9176.members.0.js
987985

50
997540
node9177.members.0.js
species

988123
35
genus

988125
node9179.members.0.js
species
35

355
6
572922
subfamily
node9180.members.0.js

genus
997550
45

997551
node9182.members.0.js
species
45

1
74
genus
node9183.members.0.js
689302

37
987872
species
node9184.members.0.js

36
987866
species
node9185.members.0.js

genus
988070
59

59
node9187.members.0.js
species
988071

2492374
65
genus

node9189.members.0.js
species
2492375
65

genus
988105
65

65
988106
species
node9191.members.0.js

genus
1101105
41

node9193.members.0.js
species
1101106
41

subfamily
84
116124

genus
116129
39

116130
species
node9196.members.0.js
39

genus
45
116125

45
species
node9198.members.0.js
116126

95186
node9199.members.0.js
subfamily
104
3

node9200.members.0.js
genus
254361
1
63

41
species
node9201.members.0.js
987893

21
254363
node9202.members.0.js
species

32
254364
genus

938171
species
node9204.members.0.js
32

7110
6
genus

7111
species
node9206.members.0.js
6

373
95179
subfamily

2555566
node9208.members.0.js
tribe
256
9

3
99
node9209.members.0.js
genus
214276

39
987995
node9210.members.0.js
species

29
node9211.members.0.js
species
214277

28
753202
node9212.members.0.js
species

45
214282
genus

45
320037
node9214.members.0.js
species

103
320016
genus

987431
species
node9216.members.0.js
56

node9217.members.0.js
species
988049
47

116
1
2555556
tribe
node9218.members.0.js

214170
48
genus

48
node9220.members.0.js
species
214171

882791
26
genus

species
node9222.members.0.js
988174
26

41
95189
genus

41
node9224.members.0.js
species
875885

genus
1
320073

1223490
node9226.members.0.js
species
1

95178
9
subfamily

7112
9
genus

29058
node9229.members.0.js
species
2

7
species
node9230.members.0.js
7113

subfamily
93
95175

50
56362
genus

species
node9233.members.0.js
987859
50

43
938225
genus

43
node9235.members.0.js
species
938226

4
278
family
node9236.members.0.js
37571

319773
68
subfamily

genus
68
13633

species
node9239.members.0.js
753216
68

subfamily
19
319770

genus
19
987448

19
987449
node9242.members.0.js
species

103
319765
subfamily

genus
node9244.members.0.js
214280
3
57

26
753204
node9245.members.0.js
species

28
species
node9246.members.0.js
988002

214308
genus
node9247.members.0.js
46
7

988018
species
node9248.members.0.js
16

988019
node9249.members.0.js
species
23

45
319766
subfamily

45
56587
genus

node9252.members.0.js
species
987943
45

319762
39
subfamily

genus
39
214089

987902
node9255.members.0.js
species
39

family
node9256.members.0.js
695564
6
459

subfamily
27548
154

genus
43
319798

43
987935
species
node9259.members.0.js

33413
52
genus

335469
node9261.members.0.js
species
52

59
13122
genus

78897
species
node9263.members.0.js
59

subfamily
95217
104

genus
411962
38

411963
species
node9266.members.0.js
38

66
95218
genus

753189
species
node9268.members.0.js
66

95191
1
subfamily

genus
1
95192

1
95193
species
node9271.members.0.js

1583079
31
subfamily

genus
31
938237

938238
species
node9274.members.0.js
31

subfamily
163
30225

tribe
node9276.members.0.js
1945703
6
80

695183
47
genus

47
node9278.members.0.js
species
875881

genus
214365
27

27
875880
species
node9280.members.0.js

83
132199
tribe

83
1
464722
genus
node9282.members.0.js

35
species
node9283.members.0.js
987419

987424
node9284.members.0.js
species
47

104494
1
superfamily

family
268500
1

genus
1
268501

node9288.members.0.js
species
268502
1

104423
61
superfamily

60
104425
family

219490
60
subfamily

genus
219491
44

721137
node9293.members.0.js
species
44

16
721162
genus

node9295.members.0.js
species
721163
16

104424
1
family

genus
node9297.members.0.js
2662233
1

37572
node9298.members.0.js
superfamily
1558
31

910
19
33415
node9299.members.0.js
family

subfamily
61
100750

tribe
215788
61

genus
124410
61

species
node9303.members.0.js
270466
61

node9304.members.0.js
subfamily
42282
1
321

124
1664845
tribe

4
124
node9306.members.0.js
genus
111897

90
447833
node9307.members.0.js
species

30
111903
species
node9308.members.0.js

tribe
node9309.members.0.js
127320
1
196

subtribe
167180
4

genus
110367
4

4
110368
species
node9312.members.0.js

81
1
366209
node9313.members.0.js
subtribe

39
111915
genus

39
species
node9315.members.0.js
111917

111932
41
genus

species
node9317.members.0.js
116150
41

167177
1
subtribe

76216
1
genus

1
species
node9320.members.0.js
76217

subtribe
150883
74

74
1
111919
genus
node9322.members.0.js

subgenus
43
111885

2795564
node9324.members.0.js
species
43

no rank
30
111950

30
species
node9326.members.0.js
191418

35
150884
subtribe

genus
111922
35

35
111923
species
node9329.members.0.js

264
2
40040
subfamily
node9330.members.0.js

3
173
node9331.members.0.js
tribe
171576

3
68
genus
node9332.members.0.js
42274

334116
species
node9333.members.0.js
3

25
171605
species
node9334.members.0.js

37
42275
species
node9335.members.0.js

genus
76218
102

111880
subgenus
node9337.members.0.js
70
2

node9338.members.0.js
species
171585
37

31
111881
species
node9339.members.0.js

32
442324
subgenus

32
species
node9341.members.0.js
171594

171578
89
tribe

node9343.members.0.js
subtribe
171580
4
89

596672
26
genus

26
113330
species
node9345.members.0.js

genus
59
104514

59
113334
species
node9347.members.0.js

1
164
node9348.members.0.js
subfamily
40037

127312
tribe
node9349.members.0.js
121
8

525812
46
genus

species
node9351.members.0.js
405009
46

127313
27
genus

27
species
node9353.members.0.js
191398

genus
405031
40

species
node9355.members.0.js
405034
40

tribe
42
127322

42
33416
genus

node9358.members.0.js
species
33443
42

subfamily
81
127218

42315
78
tribe

344711
78
subtribe

node9362.members.0.js
genus
64444
17
78

26
331299
species

26
331333
node9364.members.0.js
subspecies

species
35
304554

35
subspecies
node9366.members.0.js
331302

3
30248
tribe

subtribe
3
344705

13036
3
genus

subgenus
3
151542

13037
3
species

278856
subspecies
node9372.members.0.js
3

7143
26
family

42289
26
subfamily

tribe
189315
26

genus
26
7145

20
76193
species
node9377.members.0.js

4
node9378.members.0.js
species
76194

2
66420
node9379.members.0.js
species

node9380.members.0.js
family
27544
1
291

253
15
42297
subfamily
node9381.members.0.js

genus
55
42298

203782
node9383.members.0.js
species
55

53
11
138069
genus
node9384.members.0.js

268709
species
node9385.members.0.js
32

138070
species
node9386.members.0.js
10

genus
22
1821620

988025
species
node9388.members.0.js
22

26
242266
genus

26
species
node9390.members.0.js
242267

34
91737
genus

34
node9392.members.0.js
species
91739

48
203780
genus

48
species
node9394.members.0.js
203781

subfamily
124406
37

265359
37
genus

282391
species
node9397.members.0.js
37

7114
family
node9398.members.0.js
299
2

42450
subfamily
node9399.members.0.js
52
1

genus
42295
51

species
node9401.members.0.js
72248
51

subfamily
151208
46

189907
46
genus

node9404.members.0.js
species
189913
46

199
42449
subfamily

tribe
152601
199

11
102
node9407.members.0.js
genus
7115

29
78633
node9408.members.0.js
species

species
node9409.members.0.js
7116
25

37
node9410.members.0.js
species
64459

genus
50
129396

50
node9412.members.0.js
species
129397

72244
47
genus

47
species
node9414.members.0.js
227532

1
124343
family

subfamily
1
42265

1
124372
tribe

2250070
1
genus

1
2250071
species
node9419.members.0.js

superfamily
302
37573

7135
53
family

subfamily
40082
6

6
7136
genus

6
7137
node9424.members.0.js
species

299347
40
subfamily

1101094
40
genus

species
node9427.members.0.js
1101095
40

subfamily
40083
7

680682
7
genus

680683
node9430.members.0.js
species
7

249
268499
family

subfamily
299376
1

2231935
1
genus

1
2619405
no rank

2231937
node9435.members.0.js
species
1

200
40081
subfamily

genus
61
1368975

1371681
species
node9438.members.0.js
61

51
572825
genus

1594250
node9440.members.0.js
species
51

genus
88
572808

node9442.members.0.js
species
1660579
35

node9443.members.0.js
species
1594226
53

40080
7
subfamily

29056
7
genus

7
node9446.members.0.js
species
93504

subfamily
581380
1

genus
572908
1

1
species
node9449.members.0.js
1002954

subfamily
40
299362

genus
40
687116

1594321
node9452.members.0.js
species
40

superfamily
202
40092

family
202
40093

40100
93
subfamily

genus
40102
63

218760
species
node9457.members.0.js
63

30
76212
genus

30
node9459.members.0.js
species
520884

109
40096
subfamily

2839394
node9461.members.0.js
tribe
109
1

218734
27
genus

27
species
node9463.members.0.js
291688

genus
218743
30

30
species
node9465.members.0.js
876063

218770
51
genus

node9467.members.0.js
species
272628
51

41011
63
superfamily

63
41012
family

236781
63
subfamily

63
41013
genus

28
1101063
node9472.members.0.js
species

1594354
node9473.members.0.js
species
35

37581
210
superfamily

1
173647
family

subfamily
248746
1

genus
1
545366

1477025
species
node9478.members.0.js
1

70
753492
family

genus
116120
70

70
node9481.members.0.js
species
116121

family
104
173649

104
5
655692
node9483.members.0.js
genus

1869501
node9484.members.0.js
species
54

45
species
node9485.members.0.js
2561016

family
35
2681869

116119
35
subfamily

35
262437
genus

1857958
node9489.members.0.js
species
35

67
37582
superfamily

55
687156
family

55
687147
genus

55
1870435
node9493.members.0.js
species

family
12
51653

12
51654
genus

node9496.members.0.js
species
51655
12

956
3
7399
node9497.members.0.js
order

85772
29
superfamily

family
52632
5

410274
5
subfamily

5
270857
genus

species
node9502.members.0.js
441921
1

node9503.members.0.js
species
2872261
4

27532
24
family

2591321
1
subfamily

genus
1
362122

1
1384800
node9507.members.0.js
species

subfamily
112290
10

37343
10
genus

node9510.members.0.js
species
37344
10

subfamily
112287
13

112291
13
genus

13
362091
species
node9513.members.0.js

7400
suborder
node9514.members.0.js
914
10

superfamily
1
27487

family
1
27488

genus
1
32426

1
species
node9518.members.0.js
32427

785
14
7434
node9519.members.0.js
infraorder

2153479
163
superfamily

node9521.members.0.js
family
36668
7
163

3
94
subfamily
node9522.members.0.js
34695

tribe
node9523.members.0.js
143999
4
28

genus
64782
3

species
node9525.members.0.js
103372
3

genus
6
34699

species
node9527.members.0.js
456900
6

2
6
genus
node9528.members.0.js
12956

4
species
node9529.members.0.js
520822

34717
9
genus

node9531.members.0.js
species
64791
6

1
node9532.members.0.js
species
34720

2
node9533.members.0.js
species
471704

tribe
144001
18

18
64792
genus

species
node9536.members.0.js
64793
18

tribe
144020
20

genus
30204
20

species
node9539.members.0.js
219812
20

tribe
7
1932955

genus
7
144030

node9542.members.0.js
species
144034
7

12
144017
tribe

13685
6
genus

species
node9545.members.0.js
13686
6

55077
6
genus

6
307658
species
node9547.members.0.js

6
144004
tribe

300110
5
genus

300111
node9550.members.0.js
species
5

1
29040
genus

no rank
200581
1

node9553.members.0.js
species
1297371
1

subfamily
40138
5

genus
56621
5

219809
node9556.members.0.js
species
5

43085
21
subfamily

1
21
tribe
node9558.members.0.js
141711

genus
604375
8

8
species
node9560.members.0.js
610380

6
43086
genus

486640
node9562.members.0.js
species
6

genus
6
369122

6
609295
species
node9564.members.0.js

7479
20
subfamily

72772
9
tribe

2
1
262037
genus
node9567.members.0.js

262038
node9568.members.0.js
species
1

genus
710235
7

7
species
node9570.members.0.js
613905

72773
3
tribe

13390
3
genus

104421
species
node9573.members.0.js
3

72771
8
tribe

72766
8
genus

node9576.members.0.js
species
72781
8

9
213859
subfamily

genus
2015172
9

9
node9579.members.0.js
species
2015173

subfamily
40139
7

83484
7
genus

7
83485
node9582.members.0.js
species

142
34725
superfamily

142
1
7438
family
node9584.members.0.js

subfamily
29
50638

76989
29
genus

76990
node9587.members.0.js
species
29

9
99
node9588.members.0.js
subfamily
7439

genus
node9589.members.0.js
7451
9
27

7454
node9590.members.0.js
species
13

5
species
node9591.members.0.js
30212

46
18
7440
genus
node9592.members.0.js

species
node9593.members.0.js
85443
10

node9594.members.0.js
species
85444
4

node9595.members.0.js
species
881891
14

17
3
7443
node9596.members.0.js
genus

202808
node9597.members.0.js
species
8

5
species
node9598.members.0.js
7445

1
node9599.members.0.js
species
7446

subfamily
13
7455

tribe
76984
13

node9602.members.0.js
genus
7456
2
13

3
species
node9603.members.0.js
91411

743375
node9604.members.0.js
species
4

30207
species
node9605.members.0.js
4

439
12
34735
node9606.members.0.js
superfamily

77572
120
family

subfamily
156312
2

genus
2
178049

2448183
2
subgenus

2448451
node9611.members.0.js
species
2

7
115
subfamily
node9612.members.0.js
77573

1
62
tribe
node9613.members.0.js
88544

genus
16
88466

species
node9615.members.0.js
115100
16

2
45
genus
node9616.members.0.js
88467

25
88472
subgenus

25
88531
node9618.members.0.js
species

18
88475
subgenus

18
88514
node9620.members.0.js
species

44
479730
tribe

genus
88591
44

1190790
node9623.members.0.js
species
44

2
88545
tribe

genus
115080
2

2
node9626.members.0.js
species
115081

178025
3
subfamily

3
178032
genus

3
node9629.members.0.js
species
178035

3
156309
family

subfamily
178044
3

156310
3
genus

935657
species
node9633.members.0.js
3

family
24
124286

subfamily
24
156330

tribe
14
156332

2
14
node9637.members.0.js
genus
124287

1
species
node9638.members.0.js
473952

1437190
11
species

subspecies
node9640.members.0.js
1437191
11

156331
10
tribe

genus
132116
10

10
node9643.members.0.js
species
143995

156323
41
family

41
253710
subfamily

253714
41
genus

41
species
node9647.members.0.js
253715

125
7458
family

78169
26
subfamily

tribe
26
95294

genus
95295
26

601510
species
node9652.members.0.js
26

subfamily
78170
3

78171
3
tribe

genus
3
78173

236025
3
subgenus

156304
species
node9657.members.0.js
3

70987
node9658.members.0.js
subfamily
96
3

tribe
4
83319

genus
166413
4

node9661.members.0.js
species
561572
4

83311
77
tribe

77
16
28641
genus
node9663.members.0.js

144708
7
subgenus

7
node9665.members.0.js
species
30195

subgenus
144715
1

1
species
node9667.members.0.js
396416

20
4
28653
node9668.members.0.js
subgenus

30201
species
node9669.members.0.js
8

8
207624
node9670.members.0.js
species

subgenus
node9671.members.0.js
144703
1
8

7
species
node9672.members.0.js
30191

subgenus
144700
9

node9674.members.0.js
species
65598
9

144704
16
subgenus

85660
species
node9676.members.0.js
16

1
83323
tribe

1
117248
genus

1
node9679.members.0.js
species
597456

83321
10
tribe

node9681.members.0.js
genus
7459
4
10

1
7462
node9682.members.0.js
species

node9683.members.0.js
species
7460
1

species
node9684.members.0.js
7461
3

1
node9685.members.0.js
species
7463

83310
1
tribe

genus
28644
1

1
516756
species
node9688.members.0.js

98
2153468
no rank

family
98
253718

253722
39
subfamily

288410
39
tribe

subtribe
421285
39

421297
genus
node9694.members.0.js
39
5

2495015
species
node9695.members.0.js
9

node9696.members.0.js
species
1126389
25

subfamily
29
216423

tribe
29
288382

genus
29
288388

29
node9700.members.0.js
species
1167272

subfamily
18
288404

tribe
302523
18

18
421220
genus

18
species
node9704.members.0.js
2495127

12
253723
subfamily

421418
12
tribe

genus
421423
12

12
2495085
node9708.members.0.js
species

family
16
48719

subfamily
16
205141

16
48720
genus

16
205271
subgenus

16
444401
species
node9713.members.0.js

superfamily
1803217
27

27
27515
family

subfamily
1801551
27

200613
27
genus

1667466
node9718.members.0.js
species
27

1955251
118
infraorder

superfamily
7401
67

7408
39
family

7
65149
subfamily

no rank
7
273877

7
92443
genus

species
node9725.members.0.js
32260
7

subfamily
32
65170

tribe
172511
32

genus
27520
32

2795680
node9729.members.0.js
species
32

family
28
7402

subfamily
65210
11

11
7405
genus

11
460826
node9733.members.0.js
species

65207
5
subfamily

3
1
32390
genus
node9735.members.0.js

species
node9736.members.0.js
32391
2

genus
2
51538

2
species
node9738.members.0.js
69319

subfamily
7
68883

genus
5
58738

5
64838
node9741.members.0.js
species

2
454922
genus

2
454923
species
node9743.members.0.js

node9744.members.0.js
subfamily
68882
2
5

2
37852
genus

684658
species
node9746.members.0.js
2

genus
55913
1

node9748.members.0.js
species
77494
1

superfamily
46
7422

75187
6
family

6
75190
subfamily

genus
6
84507

species
142686
6

326594
subspecies
node9754.members.0.js
6

29051
3
family

subfamily
272199
3

genus
29052
3

29053
species
node9758.members.0.js
3

family
7489
7

7490
7
genus

7
7493
node9761.members.0.js
species

30
7423
family

272242
30
subfamily

genus
7424
30

30
node9765.members.0.js
species
7425

superfamily
5
40307

family
3
44353

3
303412
subfamily

3
63429
genus

species
node9770.members.0.js
63436
3

73401
2
family

subfamily
2
1159319

2
75144
tribe

genus
1159320
2

2
2817044
node9775.members.0.js
species

4
222831
superfamily

27528
4
family

genus
4
27529

222816
node9779.members.0.js
species
4

superfamily
222823
6

6
27524
family

6
173784
genus

6
species
node9783.members.0.js
211228

7041
507
order

suborder
437
41084

infraorder
54
41087

71192
2
superfamily

50527
2
family

2
261156
subfamily

genus
2
195164

2
224129
node9791.members.0.js
species

superfamily
52
71193

7049
5
family

5
433514
subfamily

genus
7053
5

5
node9796.members.0.js
species
7054

41097
47
family

1
46
node9798.members.0.js
subfamily
433502

24
41098
genus

195172
node9800.members.0.js
species
24

41100
21
genus

21
41101
species
node9802.members.0.js

433505
1
subfamily

1
334007
genus

1
2055709
node9805.members.0.js
species

41088
infraorder
node9806.members.0.js
310
3

71528
30
superfamily

34667
17
family

subfamily
17
79514

tribe
17
192383

genus
157304
17

17
217634
species
node9812.members.0.js

13
27439
family

1
131663
subfamily

2138365
1
genus

2677818
1
no rank

1
node9817.members.0.js
species
2055550

subfamily
2
63710

tribe
63711
2

2
226742
subtribe

no rank
2
226749

genus
50385
2

species
50389
2

2
50390
node9824.members.0.js
subspecies

63707
10
subfamily

tribe
10
63712

10
7538
genus

10
species
node9828.members.0.js
7539

32
71527
superfamily

34672
1
family

1
2507001
subfamily

34750
1
genus

1
species
node9833.members.0.js
1323541

17
55098
family

genus
295984
17

17
346838
node9836.members.0.js
species

family
14
7065

1304792
13
no rank

genus
13
7069

4
7070
species
node9840.members.0.js

9
species
node9841.members.0.js
41895

subfamily
1272142
1

genus
343927
1

no rank
2685333
1

1
species
node9845.members.0.js
2055318

superfamily
55
71525

186093
55
family

353826
55
subfamily

295699
55
genus

species
node9850.members.0.js
295700
55

superfamily
71526
150

family
196992
2

subfamily
2
196991

295905
2
genus

no rank
2
2631161

2
species
node9856.members.0.js
2055371

family
1
196982

subfamily
1
432910

genus
432911
1

no rank
2638166
1

node9861.members.0.js
species
2055512
1

family
7080
93

subfamily
93
7081

93
263631
tribe

genus
115356
29

29
115357
species
node9866.members.0.js

genus
41138
23

23
41139
node9868.members.0.js
species

genus
7083
41

41
node9870.members.0.js
species
7084

family
1
196978

297219
1
subfamily

genus
1
2055681

no rank
1
2637169

node9875.members.0.js
species
2055682
1

196996
18
family

genus
18
484807

2633831
18
no rank

18
2055333
species
node9879.members.0.js

family
35
116151

subfamily
577241
28

28
1431902
genus

28
species
node9883.members.0.js
1431903

7
577242
subfamily

genus
7
116152

species
node9886.members.0.js
116153
7

40
71529
superfamily

122737
32
family

32
701798
subfamily

32
122772
genus

node9891.members.0.js
species
201766
32

family
7042
4

subfamily
55867
3

3
77156
genus

77166
node9895.members.0.js
species
3

subfamily
39812
1

genus
1
7045

species
node9898.members.0.js
7048
1

4
2878387
family

subfamily
122824
4

4
166876
genus

308863
node9902.members.0.js
species
4

4
41086
infraorder

superfamily
75546
4

7055
4
family

4
41142
subfamily

4
569643
no rank

genus
4
166331

species
node9909.members.0.js
166361
4

41085
68
infraorder

75543
68
superfamily

family
3
57514

3
82882
subfamily

57515
3
genus

3
node9915.members.0.js
species
110193

family
1
66535

66536
1
subfamily

1
701471
genus

1
2622888
no rank

2055325
species
node9920.members.0.js
1

64
29026
family

no rank
64
351514

64
82886
subfamily

64
1
295648
node9924.members.0.js
tribe

290671
63
genus

662956
node9926.members.0.js
species
63

infraorder
1
41093

superfamily
1
75547

family
1
41094

genus
1
910044

no rank
1
2627127

2055478
node9932.members.0.js
species
1

70
41071
suborder

superfamily
2
535378

family
2
50515

1
107842
subfamily

tribe
1
183333

genus
107767
1

183348
node9939.members.0.js
species
1

107841
1
subfamily

1
390869
genus

2625789
1
no rank

1
1589306
species
node9943.members.0.js

535382
68
superfamily

family
68
41073

subfamily
68
71541

879230
68
tribe

genus
41078
68

subgenus
484232
68

species
node9950.members.0.js
767470
68

7147
node9951.members.0.js
order
809
6

1
585
node9952.members.0.js
suborder
7203

infraorder
2
43734

34687
2
family

2
343564
subfamily

genus
2
343581

2
343691
species
node9957.members.0.js

43733
582
infraorder

clade
582
480118

8
582
node9960.members.0.js
clade
480117

380
4
43738
node9961.members.0.js
no rank

43742
53
no rank

superfamily
4
43753

family
7392
4

7393
4
genus

subgenus
44051
4

4
species
node9967.members.0.js
7396

superfamily
43755
39

family
7
7381

43916
7
subfamily

7384
7
genus

subgenus
321190
7

7
596942
species
node9973.members.0.js

family
19
27474

subfamily
43917
1

1
179426
tribe

1
1918309
genus

1918310
node9978.members.0.js
species
1

subfamily
17
54288

tribe
17
569116

17
569065
node9981.members.0.js
genus

54286
1
subfamily

tribe
569105
1

1
569045
genus

1
node9985.members.0.js
species
569046

family
13
7371

subfamily
13
43914

7374
node9988.members.0.js
genus
13
1

6
13632
node9989.members.0.js
species

species
node9990.members.0.js
7375
6

superfamily
43754
10

family
7366
10

43910
10
subfamily

tribe
43911
4

35569
4
genus

species
node9996.members.0.js
35570
4

57894
6
tribe

genus
7369
6

44052
6
subgenus

6
node10000.members.0.js
species
7370

no rank
node10001.members.0.js
43741
2
323

superfamily
5
43745

139644
5
family

5
139679
genus

5
139649
species
node10005.members.0.js

1
43750
superfamily

family
1
169447

genus
1
1226614

1226616
species
node10009.members.0.js
1

superfamily
44
43752

44
7211
family

34
164860
subfamily

tribe
43871
31

4
47833
genus

subgenus
4
1987911

4
28588
node10016.members.0.js
species

3
27
genus
node10017.members.0.js
27456

subgenus
12
69624

species
node10019.members.0.js
104688
12

12
3
47832
subgenus
node10020.members.0.js

2
174628
node10021.members.0.js
species

no rank
98808
3

3
59916
species
node10023.members.0.js

4
98805
no rank

node10025.members.0.js
species
27457
4

164862
3
tribe

genus
7212
3

subgenus
474492
3

3
7213
species
node10029.members.0.js

43867
10
subfamily

10
43901
tribe

subtribe
10
164882

28609
genus
node10033.members.0.js
10
1

5
28610
node10034.members.0.js
species

4
species
node10035.members.0.js
28612

242
43746
superfamily

family
node10037.members.0.js
7214
1
242

3
43846
subfamily

tribe
46881
2

subtribe
106236
2

genus
106235
2

462269
node10042.members.0.js
species
2

1
46882
tribe

46884
1
subtribe

188292
1
genus

no rank
1
757332

2518853
species
node10047.members.0.js
1

subfamily
43845
238

5
1861795
tribe

genus
5
7354

32386
5
species group

5
species
node10052.members.0.js
7225

tribe
46877
233

233
3
7215
node10054.members.0.js
genus

4
504493
no rank

4
48384
clade

clade
48301
4

species group
4
48302

species subgroup
32378
4

node10060.members.0.js
species
7222
4

1
72
node10061.members.0.js
subgenus
32281

14
32304
species group

species subgroup
32307
14

14
species
node10064.members.0.js
7291

4
12
node10065.members.0.js
species group
32335

2
7244
node10066.members.0.js
species

species
node10067.members.0.js
47314
6

32320
9
species group

species
node10069.members.0.js
198719
9

2
35
node10070.members.0.js
species group
32321

species subgroup
32324
16

170944
node10072.members.0.js
no rank
2

14
3
198037
node10073.members.0.js
no rank

4
7232
species
node10074.members.0.js

4
species
node10075.members.0.js
7263

node10076.members.0.js
species
7230
3

species subgroup
17
40364

7224
species
node10078.members.0.js
17

species group
1
32290

species subgroup
32291
1

node10081.members.0.js
species
133979
1

19
32280
subgenus

30019
species
node10083.members.0.js
19

subgenus
135
32341

node10085.members.0.js
species group
32355
1
24

species subgroup
node10086.members.0.js
32357
1
18

7241
species
node10087.members.0.js
4

2
7266
node10088.members.0.js
species

11
node10089.members.0.js
species
7282

2
5
species subgroup
node10090.members.0.js
32358

1
node10091.members.0.js
species
7237

species
node10092.members.0.js
7229
2

species group
4
32365

species subgroup
32367
4

4
7260
node10095.members.0.js
species

11
107
node10096.members.0.js
species group
32346

3
32350
species subgroup

30025
species
node10098.members.0.js
3

32347
node10099.members.0.js
species subgroup
4
1

2
545632
no rank

2
species
node10101.members.0.js
7217

1
186282
no rank

1
species
node10103.members.0.js
42026

32351
node10104.members.0.js
species subgroup
49
6

species
node10105.members.0.js
7243
2

7245
node10106.members.0.js
species
2

node10107.members.0.js
species
7227
18

species
node10108.members.0.js
7240
4

node10109.members.0.js
species
7220
8

6
species
node10110.members.0.js
7226

129105
node10111.members.0.js
species
1

species
node10112.members.0.js
7238
2

6
32349
species subgroup

species
node10114.members.0.js
29029
6

5
65962
species subgroup

5
species
node10116.members.0.js
1041015

32354
4
species subgroup

species
node10118.members.0.js
29030
4

species subgroup
32348
6

30023
node10120.members.0.js
species
6

32352
node10121.members.0.js
species subgroup
11
1

node10122.members.0.js
species
30033
6

446045
4
no rank

species
node10124.members.0.js
7274
4

species subgroup
32353
8

1486046
species
node10126.members.0.js
6

1
28584
node10127.members.0.js
species

1
species
node10128.members.0.js
125945

29
43744
superfamily

family
29
115263

subfamily
115265
29

genus
1219203
29

1219204
species
node10133.members.0.js
29

43737
194
no rank

194
43740
superfamily

194
34680
family

subfamily
115244
162

tribe
192448
39

39
173981
genus

273409
species
node10140.members.0.js
39

tribe
3
115284

genus
226148
3

species
node10143.members.0.js
226151
3

tribe
40
224230

genus
34681
26

node10146.members.0.js
species
34682
26

genus
14
224240

14
node10148.members.0.js
species
374264

tribe
115277
80

80
5
198633
genus
node10150.members.0.js

25
198635
node10151.members.0.js
species

1124515
node10152.members.0.js
species
25

25
node10153.members.0.js
species
1572519

32
43838
subfamily

115274
26
tribe

290403
3
genus

414732
3
subgenus

3
1124558
species
node10158.members.0.js

3
219506
genus

species
node10160.members.0.js
323313
3

5
414873
genus

node10162.members.0.js
species
414876
5

219538
15
genus

15
219539
node10164.members.0.js
species

tribe
224219
6

genus
92597
2

2
node10167.members.0.js
species
653684

4
192444
genus

species
node10169.members.0.js
414846
4

218
7148
suborder

infraorder
43785
2

2
560808
superfamily

family
560809
2

genus
560766
2

560767
species
node10175.members.0.js
2

infraorder
40
43784

14
1
41830
node10177.members.0.js
superfamily

node10178.members.0.js
family
7184
3
6

35571
node10179.members.0.js
genus
3
1

2
node10180.members.0.js
species
38358

33406
7
family

7
43793
subfamily

no rank
7
52723

tribe
71814
7

genus
7
153220

7
node10186.members.0.js
species
265458

superfamily
43790
26

26
52729
family

52730
26
subfamily

genus
189978
26

189979
species
node10191.members.0.js
26

43786
175
infraorder

superfamily
36
41828

family
22
41819

22
43801
subfamily

tribe
22
58262

genus
22
41820

subgenus
22
58277

node10199.members.0.js
species
179676
22

14
1
7149
family
node10200.members.0.js

54970
5
subfamily

2
72532
tribe

genus
2
82133

288873
node10204.members.0.js
species
2

tribe
3
72530

1
3
genus
node10206.members.0.js
7150

subgenus
41809
2

species
node10208.members.0.js
41810
2

305539
node10209.members.0.js
no rank
4
1

species
node10210.members.0.js
2766959
3

43808
subfamily
node10211.members.0.js
4
1

61019
1
genus

2646824
1
no rank

1
node10214.members.0.js
species
61020

315559
genus
node10215.members.0.js
2
1

1
611654
node10216.members.0.js
species

41827
139
superfamily

139
7157
family

43817
43
subfamily

tribe
1056966
26

genus
14
124916

subgenus
317811
14

14
124917
species
node10223.members.0.js

7158
12
genus

12
53541
subgenus

7159
species
node10226.members.0.js
5

7
species
node10227.members.0.js
7160

53550
17
tribe

genus
7174
17

17
53527
subgenus

6
17
node10231.members.0.js
no rank
518105

species
5
7175

42434
subspecies
node10233.members.0.js
5

7176
node10234.members.0.js
species
6

43816
96
subfamily

7164
node10236.members.0.js
genus
96
1

44543
58
subgenus

clade
44544
58

58
44547
clade

node10240.members.0.js
species
7167
58

node10241.members.0.js
subgenus
44534
1
37

27
44535
clade

27
species
node10243.members.0.js
30069

9
44537
clade

2
9
no rank
node10245.members.0.js
44542

1
species
node10246.members.0.js
7173

node10247.members.0.js
species
1518534
3

species
node10248.members.0.js
30066
2

7165
1
species

strain
node10250.members.0.js
180454
1

43789
1
infraorder

41829
1
superfamily

1
node10253.members.0.js
family
41042

34
1
85817
superorder
node10254.members.0.js

order
33
7516

suborder
33
2029065

family
32
7520

2029106
32
subfamily

genus
32
7521

32
189513
species
node10260.members.0.js

50435
1
family

50436
1
genus

no rank
2637538
1

270870
species
node10264.members.0.js
1

19
33339
infraclass

order
19
6961

suborder
50488
19

19
70894
superfamily

70895
19
family

genus
79456
19

19
197161
species
node10271.members.0.js

1
554674
subclass

order
1
29994

85513
1
family

genus
299217
1

1
node10276.members.0.js
species
299218

30001
20
class

order
1
79705

1
187607
family

574227
1
genus

species
node10281.members.0.js
574228
1

730330
node10282.members.0.js
order
19
2

superfamily
4
730333

36141
4
family

3
187620
subfamily

genus
158440
3

158441
node10287.members.0.js
species
3

subfamily
187621
1

1
281417
genus

1
1184801
node10290.members.0.js
species

1
13
superfamily
node10291.members.0.js
79707

48704
12
family

2
2267846
node10293.members.0.js
subfamily

50238
6
subfamily

1
genus
node10295.members.0.js
301509

1
187694
genus

no rank
2644615
1

1
node10298.members.0.js
species
1304854

301513
1
genus

species
node10300.members.0.js
574220
1

3
187625
genus

1
2041941
species
node10302.members.0.js

no rank
2643097
1

species
node10304.members.0.js
2013028
1

1
1690820
node10305.members.0.js
species

subfamily
4
187687

genus
1106259
2

2
species
node10308.members.0.js
2492651

181624
2
genus

2
species
node10310.members.0.js
1302335

61985
1
subphylum

class
63448
1

2082948
1
order

1
61989
family

no rank
146860
1

1
node10316.members.0.js
species
1366095

subphylum
4983
6843

class
7
6844

order
7
6845

6846
7
family

7
6849
genus

node10322.members.0.js
species
6850
7

6854
node10323.members.0.js
class
4975
1

6855
2
order

parvorder
259437
2

70336
2
superfamily

6856
2
family

genus
2
6875

2
218467
species
node10329.members.0.js

43271
5
order

clade
5
101165

suborder
43278
3

superfamily
node10333.members.0.js
101159
1
3

101160
2
family

subfamily
121215
2

genus
2
118623

118624
species
node10337.members.0.js
2

2
43288
suborder

101153
2
superfamily

family
2
1277271

genus
1006705
2

2
2685587
no rank

2
1006706
species
node10343.members.0.js

subclass
node10344.members.0.js
6933
1
4945

3490
2
6934
superorder
node10345.members.0.js

order
34634
3462

281668
3462
suborder

1723665
infraorder
node10348.members.0.js
3462
7

3416
2
1253825
node10349.members.0.js
superfamily

1
3414
family
node10350.members.0.js
34636

2
425252
subfamily

425256
2
genus

2291836
species
node10353.members.0.js
2

425253
3357
subfamily

genus
3357
1933192

node10356.members.0.js
species
34638
3357

3
no rank
node10357.members.0.js
645130

51
2
425251
subfamily
node10358.members.0.js

99210
2
genus

2
2645455
no rank

2
1712292
species
node10361.members.0.js

genus
2
84383

2
node10363.members.0.js
species
702746

genus
425255
4

node10365.members.0.js
species
425259
4

41
84381
genus

species
node10367.members.0.js
193551
8

1
node10368.members.0.js
species
1609164

84382
node10369.members.0.js
species
20

species
node10370.members.0.js
84385
4

species
node10371.members.0.js
756255
1

7
node10372.members.0.js
species
573039

superfamily
1
41439

91336
1
family

1
704019
genus

1
2626893
no rank

species
node10377.members.0.js
2051734
1

38
41438
superfamily

38
109261
family

26
38
node10380.members.0.js
genus
62624

node10381.members.0.js
species
62625
7

5
109461
node10382.members.0.js
species

order
6935
26

297308
26
superfamily

family
26
6939

subfamily
1
426441

6942
1
genus

34610
node10388.members.0.js
species
1

subfamily
426437
21

genus
3
34619

3
543639
species
node10391.members.0.js

2
18
genus
node10392.members.0.js
34630

subgenus
9
6940

9
6941
node10394.members.0.js
species

subgenus
426455
7

species group
7
578835

node10397.members.0.js
species
34632
7

subfamily
4
426442

genus
6944
4

species
node10400.members.0.js
6945
4

superorder
1454
6946

64
83137
order

1
13
suborder
node10403.members.0.js
6951

2
66561
superfamily

2
1
83156
family
node10405.members.0.js

1
node10406.members.0.js
genus
105144

223472
9
parvorder

83163
1
superfamily

1
6952
family

subfamily
1
474036

1
6953
genus

1
node10412.members.0.js
species
6956

83158
8
superfamily

8
52281
family

subfamily
8
474019

52282
8
genus

8
node10417.members.0.js
species
52283

superfamily
1
83155

family
41442
1

subfamily
1
474069

1
41443
genus

node10422.members.0.js
species
59818
1

suborder
66551
49

49
229894
infraorder

48
229794
superfamily

229795
48
family

genus
334624
26

334625
species
node10428.members.0.js
26

genus
1979940
22

1979941
species
node10430.members.0.js
22

superfamily
1
229851

1
1427652
family

1
229867
genus

1
1323719
node10434.members.0.js
species

2
66547
suborder

superfamily
386815
2

2
386820
family

386823
2
genus

no rank
2643012
2

1
2051766
species
node10440.members.0.js

1
2051767
species
node10441.members.0.js

order
1390
83136

1390
6947
suborder

clade
13
83139

superfamily
708319
4

708320
4
family

no rank
709476
4

2051819
node10448.members.0.js
species
4

superfamily
70332
3

3
1
70320
family
node10450.members.0.js

1
426503
subfamily

1
426511
tribe

1
109364
genus

no rank
1
2623685

1
2051746
species
node10455.members.0.js

1
subfamily
node10456.members.0.js
426501

superfamily
94819
6

family
6
296205

296206
genus
node10459.members.0.js
6
3

2634748
3
no rank

3
node10461.members.0.js
species
708281

infraorder
83138
1

1
83141
clade

1
83142
superfamily

family
92103
1

subfamily
406626
1

423220
1
genus

2666603
1
no rank

1
2051735
node10469.members.0.js
species

infraorder
83145
1376

188550
1376
no rank

2
188547
superfamily

188543
2
family

genus
2
188544

481310
species
node10475.members.0.js
2

83146
node10476.members.0.js
superfamily
1374
1

1373
8
32262
family
node10477.members.0.js

genus
50025
3

3
1490239
node10479.members.0.js
species

node10480.members.0.js
genus
32263
17
1362

5
93129
species
node10481.members.0.js

node10482.members.0.js
species
32264
1335

1
60960
node10483.members.0.js
species

4
node10484.members.0.js
species
93132

6893
order
node10485.members.0.js
22
1

suborder
1
6894

family
6895
1

1
1046901
genus

1
29017
node10489.members.0.js
species

20
6905
suborder

clade
20
74971

no rank
2
320450

family
2
2736621

210001
1
genus

2653069
species
node10495.members.0.js
1

no rank
2736622
1

1
species
node10497.members.0.js
1956820

9
74974
clade

superfamily
9
74975

family
34643
9

genus
114395
1

no rank
1
2642564

1
node10503.members.0.js
species
2747258

8
449632
genus

114398
species
node10505.members.0.js
8

superfamily
175332
9

9
175333
family

genus
175340
9

202533
species
node10509.members.0.js
9

1
57294
class

order
1
373319

family
1
61893

genus
258329
1

258330
species
node10514.members.0.js
1

222
6231
phylum

class
119088
10

1457286
10
subclass

6329
10
order

10
6332
family

6333
10
genus

10
6334
node10521.members.0.js
species

class
119089
212

order
6236
120

suborder
6
6274

infraorder
6
2072716

superfamily
6295
6

6
6296
family

2
6278
genus

6279
species
node10529.members.0.js
2

genus
7208
4

species
node10531.members.0.js
7209
4

65
2301116
suborder

infraorder
1
50827

1
2301094
superfamily

family
1
50861

669145
1
genus

669146
species
node10537.members.0.js
1

2301119
64
infraorder

55879
64
superfamily

6243
64
family

5
55887
subfamily

genus
42476
5

node10543.members.0.js
species
141969
5

subfamily
59
55885

genus
6237
59

15
6239
species
node10546.members.0.js

1978547
node10547.members.0.js
species
15

21
6238
species
node10548.members.0.js

species
node10549.members.0.js
31234
8

49
6300
suborder

infraorder
9
33283

superfamily
9
33284

6301
9
family

subfamily
33286
9

genus
34509
9

node10556.members.0.js
species
51029
9

infraorder
2082223
40

40
2082224
superfamily

family
25
6246

genus
node10560.members.0.js
6247
1
20

6248
node10561.members.0.js
species
3

2
node10562.members.0.js
species
34506

species
node10563.members.0.js
174720
3

11
species
node10564.members.0.js
75913

genus
5
131309

5
131310
node10566.members.0.js
species

114888
15
family

114889
15
genus

2629767
15
no rank

species
node10570.members.0.js
114890
15

92
6308
order

85
6314
superfamily

126387
85
family

genus
6288
85

species
node10575.members.0.js
6289
85

superfamily
2572558
7

33278
7
family

subfamily
53477
7

genus
51030
7

7
species
node10580.members.0.js
51031

1215728
11
superphylum

phylum
33467
11

class
11
2082883

11
2082909
order

37891
11
family

11
37847
genus

11
37621
species
node10587.members.0.js

6073
99
phylum

19
1927913
class

19
37528
order

suborder
19
1927915

family
19
1927917

genus
37533
19

node10594.members.0.js
species
313498
19

6074
2
class

subclass
2
37516

2
406427
order

1612408
2
suborder

6080
2
family

6083
2
genus

6087
node10601.members.0.js
species
2

class
6101
78

subclass
6132
7

order
40677
7

723662
7
clade

51108
7
family

7
51109
genus

node10608.members.0.js
species
151771
7

6102
71
subclass

order
node10610.members.0.js
6125
1
20

suborder
123757
14

9
46729
family

46730
5
genus

node10614.members.0.js
species
46731
5

genus
4
50428

4
50429
node10616.members.0.js
species

family
6126
5

6127
genus
node10618.members.0.js
5
1

1
45264
node10619.members.0.js
species

3
70779
node10620.members.0.js
species

suborder
5
123760

5
46736
family

genus
5
1920453

species
node10624.members.0.js
48498
5

6103
51
order

5
42822
family

6104
5
genus

5
node10628.members.0.js
species
6105

32
86626
suborder

478428
32
family

478394
32
genus

species
node10632.members.0.js
1789172
32

family
42823
8

genus
1720308
8

2652724
node10635.members.0.js
species
8

family
6
45349

45350
6
genus

6
species
node10638.members.0.js
45351

class
3
2687318

genus
3
192874

species
192875
3

595528
strain
node10642.members.0.js
3

3
127916
class

order
3
198625

3
72018
genus

species
72019
3

3
667725
node10647.members.0.js
strain

class
28009
3

order
1924738
3

family
81529
3

genus
3
86017

3
node10652.members.0.js
species
946362

2763
17
phylum

2797
6
class

265318
6
order

265316
6
family

genus
2759657
5

5
species
node10658.members.0.js
2690220

genus
1
83373

130081
species
node10660.members.0.js
1

class
2806
11

2045258
1
subclass

111859
1
order

173442
1
family

genus
111860
1

1
species
node10666.members.0.js
111861

subclass
10
2045261

order
31468
3

31469
3
family

genus
2
2774

172968
node10671.members.0.js
species
2

1
genus
node10672.members.0.js
2781

28017
4
order

29217
4
family

4
2768
genus

4
2769
node10676.members.0.js
species

29247
1
order

family
1
31501

1
2794
genus

1
node10680.members.0.js
species
31502

2
2802
order

family
2803
1

1
2008388
tribe

99900
1
genus

1
node10685.members.0.js
species
1858662

35164
1
family

genus
1
189635

1
species
node10688.members.0.js
189649

2
2608109
clade

2
2830
phylum

clade
2608131
2

73020
2
order

418966
2
family

2
2902
genus

species
2
2903

2
280463
node10696.members.0.js
strain

2
42452
no rank

2
1928008
node10698.members.0.js
species

15
1
61964
node10699.members.0.js
no rank

12
100272
species
node10700.members.0.js

1
342097
species
node10701.members.0.js

172788
species
node10702.members.0.js
1

class
10
3027

order
10
589342

10
589343
family

genus
10
55528

55529
10
species

905079
node10708.members.0.js
strain
10

kingdom
node10709.members.0.js
33090
14
206689

phylum
35493
206571

206567
131221
subphylum

6
206567
node10712.members.0.js
clade
3193

clade
3208
5

404260
5
clade

3214
4
class

subclass
4
114656

order
4
3215

3216
4
family

37414
4
genus

3218
node10720.members.0.js
species
4

113509
1
class

order
3210
1

family
1
3211

genus
1
3212

1
129213
node10725.members.0.js
species

3195
84
clade

83
186770
class

83
186774
subclass

83
28908
order

family
41837
1

node10731.members.0.js
genus
41838
1

29585
82
family

genus
3196
82

species
82
3197

82
1480154
node10735.members.0.js
subspecies

186771
1
class

subclass
186782
1

3199
1
order

71154
1
suborder

family
41845
1

genus
1
41847

1
1926836
species
node10742.members.0.js

2
206472
clade
node10743.members.0.js
58023

1521260
1
class

order
3244
1

family
3245
1

genus
1
3246

node10748.members.0.js
species
88036
1

206469
2
78536
clade
node10749.members.0.js

2
241806
class

2
1521262
subclass

order
2
693761

3272
2
family

2
37228
genus

2
397678
species
node10755.members.0.js

3
206465
clade
node10756.members.0.js
58024

9
206249
node10757.members.0.js
class
3398

261009
6
order

6
22097
family

13332
6
genus

6
13333
node10761.members.0.js
species

1437183
clade
node10762.members.0.js
206224
177

order
22
232378

family
4401
2

genus
4402
2

2
4403
species
node10766.members.0.js

node10767.members.0.js
family
4328
1
16

4329
8
genus

8
species
node10769.members.0.js
60698

1
54936
genus

node10771.members.0.js
species
54937
1

genus
6
54954

6
54955
node10773.members.0.js
species

family
4
4429

4
4430
genus

node10776.members.0.js
species
4432
4

232347
3
clade

2
3400
order

family
1
22140

1927082
1
subfamily

1927096
1
tribe

genus
235818
1

1
235821
node10783.members.0.js
species

family
1
3401

genus
3413
1

1
3415
species
node10786.members.0.js

order
3432
1

node10788.members.0.js
family
3433
1

33
41768
order

family
1
3440

subfamily
1463138
1

tribe
1
1463139

node10793.members.0.js
genus
46984
1

32
3465
family

subfamily
1462614
29

29
3468
genus

29
species
node10797.members.0.js
3469

3
41766
subfamily

genus
3466
3

species
node10800.members.0.js
3467
3

71240
6468
clade

clade
91827
6468

node10803.members.0.js
clade
1437201
48
6468

order
1
41946

1
3792
family

1
3794
genus

species
1
1573227

1708480
node10808.members.0.js
varietas
1

3524
39
order

family
3615
1

1110380
1
subfamily

tribe
1
1110383

genus
3616
1

1
62330
node10814.members.0.js
species

family
2
3563

1316647
1
subfamily

genus
1
1844275

1
221764
species
node10818.members.0.js

240072
1
genus

1
240073
node10820.members.0.js
species

4374
1
family

genus
4375
1

1
species
node10823.members.0.js
150983

family
1804623
32

subfamily
21
1307796

15
1307774
tribe

genus
3558
14

63459
species
node10828.members.0.js
14

1307780
1
genus

1
species
node10830.members.0.js
244509

tribe
6
1307775

genus
6
3561

species
node10833.members.0.js
3562
6

11
1804621
subfamily

genus
11
3554

species
node10836.members.0.js
161934
1
11

subspecies
node10837.members.0.js
3555
10

family
3568
3

tribe
3
1141492

3
3573
genus

1
2777124
no rank

1
54811
species
node10842.members.0.js

2777055
2
subgenus

2
2777062
section

39875
node10845.members.0.js
species
2

71274
clade
node10846.members.0.js
884
2

order
41934
2

family
42219
2

1
2
node10849.members.0.js
genus
4281

species
node10850.members.0.js
179117
1

clade
91882
61

4209
51
order

family
4381
1

genus
16399
1

1
node10855.members.0.js
species
103999

4210
50
family

219103
11
subfamily

11
102818
tribe

subtribe
742010
11

genus
4264
11

11
4265
species

309979
11
subspecies

node10863.members.0.js
varietas
59895
11

subfamily
219120
19

tribe
219121
19

subtribe
745063
1

50190
1
genus

50193
node10868.members.0.js
species
1

745067
1
subtribe

49743
genus
node10870.members.0.js
1

745062
17
subtribe

genus
17
4235

species
node10873.members.0.js
1552228
1

16
4236
species
node10874.members.0.js

subfamily
20
102804

14
911341
clade

tribe
102814
14

14
4231
genus

node10879.members.0.js
species
4232
14

tribe
102809
6

6
877976
clade

4
2841721
subtribe
node10882.members.0.js

2
2841728
subtribe

genus
41574
2

species
node10885.members.0.js
72917
2

order
4036
10

suborder
364270
10

9
4037
family

9
241778
subfamily

1
241780
clade

tribe
241792
1

genus
40948
1

1
species
node10893.members.0.js
52491

241789
8
tribe

241799
8
subtribe

genus
8
4038

section
8
1873447

species
4039
8

8
79200
subspecies
node10899.members.0.js

family
4050
1

genus
1
13340

137943
node10902.members.0.js
species
1

91888
clade
node10903.members.0.js
772
3

332
1
4069
node10904.members.0.js
order

family
node10905.members.0.js
4070
1
268

424551
subfamily
node10906.members.0.js
247
1

tribe
424566
1

genus
337184
1

451527
node10909.members.0.js
species
1

tribe
8
424564

8
4071
genus

7
4072
species
node10912.members.0.js

4073
species
node10913.members.0.js
1

424574
237
tribe

237
27
4107
genus
node10915.members.0.js

54
species
node10916.members.0.js
4113

node10917.members.0.js
species
50273
10

1
node10918.members.0.js
species
1398764

17
145
subgenus
node10919.members.0.js
49274

28526
species
node10920.members.0.js
55

73
4081
species
node10921.members.0.js

subfamily
20
424554

20
424562
tribe

genus
node10924.members.0.js
4085
6
20

2
species
node10925.members.0.js
4096

1
4097
node10926.members.0.js
species

5
49451
node10927.members.0.js
species

species
node10928.members.0.js
4098
6

63
4118
family

267213
63
tribe

node10931.members.0.js
genus
4119
19
63

node10932.members.0.js
species
35885
17

26
node10933.members.0.js
species
35884

1
node10934.members.0.js
species
35883

order
4055
14

family
4056
1

1
167487
subfamily

tribe
1
167498

subtribe
1498435
1

genus
4057
1

4058
species
node10941.members.0.js
1

24966
13
family

subfamily
3
169617

169660
2
tribe

25168
2
genus

2
node10946.members.0.js
species
29788

tribe
1
169653

1
60387
node10948.members.0.js
genus

subfamily
10
169618

10
1968429
clade

10
1968428
clade

tribe
10
169640

10
3
13442
genus
node10953.members.0.js

node10954.members.0.js
species
49369
1

6
species
node10955.members.0.js
13443

order
node10956.members.0.js
4143
1
423

family
4136
21

216706
21
subfamily

tribe
21
216718

subtribe
2836339
19

genus
21880
19

subgenus
2026555
19

clade
19
2026556

19
180675
node10964.members.0.js
species

subtribe
2
2836337

2
21819
genus
node10966.members.0.js

family
156152
1

216780
1
tribe

genus
4150
1

node10970.members.0.js
species
4151
1

4144
372
family

2
315
node10972.members.0.js
tribe
426106

14
4145
genus

species
14
4146

158383
14
subspecies

14
node10976.members.0.js
varietas
158386

genus
38871
299

299
56036
node10978.members.0.js
species

57
426105
tribe

genus
4147
57

species
node10981.members.0.js
660624
57

family
16
91896

tribe
216770
2

genus
36747
2

223093
species
node10985.members.0.js
2

216772
14
tribe

genus
290219
14

species
node10988.members.0.js
290220
14

family
41399
4

genus
4
1502711

4155
species
node10991.members.0.js
4

8
4180
family

8
4181
genus

node10994.members.0.js
species
4182
8

order
41945
47

13
27065
family

node10997.members.0.js
genus
4441
1
13

12
species
node10998.members.0.js
4442

family
32
25692

genus
32
35939

node11001.members.0.js
species
253017
32

family
4345
1

subfamily
1
217037

1
217059
tribe
node11004.members.0.js

1
19955
family

genus
1
13492

1
35925
node11007.members.0.js
species

46
5496
clade
node11008.members.0.js
71275

clade
node11009.members.0.js
91835
16
1284

order
72025
387

387
3803
family

subfamily
3804
7

clade
1963083
1

genus
1
53922

362788
node11015.members.0.js
species
1

1
1917730
clade

1
54873
genus

1
66096
node11018.members.0.js
species

clade
3807
5

163487
5
tribe

35715
5
genus

node11022.members.0.js
species
207710
5

subfamily
3814
380

380
5
2231393
clade
node11024.members.0.js

52
2231387
clade

52
163725
tribe

2231390
52
clade

3817
genus
node11028.members.0.js
52
21

130453
node11029.members.0.js
species
1

3818
species
node11030.members.0.js
30

256
6
2231382
clade
node11031.members.0.js

132
1
2233855
clade
node11032.members.0.js

tribe
1
163733

genus
1
53625

1
species
node11035.members.0.js
690557

163726
tribe
node11036.members.0.js
2

tribe
5
163715

5
3815
genus

5
3816
species
node11039.members.0.js

123
2
163735
node11040.members.0.js
tribe

3820
7
genus

7
3821
node11042.members.0.js
species

3883
6
genus

node11044.members.0.js
species
3885
6

95
6
3913
genus
node11045.members.0.js

35
species
node11046.members.0.js
3917

species
157791
1

1
varietas
node11048.members.0.js
3916

2
53
species
node11049.members.0.js
3914

node11050.members.0.js
varietas
157739
51

genus
3846
13

node11052.members.0.js
subgenus
1462606
6
13

1
species
node11053.members.0.js
3848

6
node11054.members.0.js
species
3847

clade
118
2233838

clade
55
2233857

tribe
55
163747

genus
55
3867

55
species
node11059.members.0.js
34305

clade
63
2233839

tribe
163722
29

genus
29
3826

29
3827
node11063.members.0.js
species

1
163743
tribe

1
3904
genus

1
3908
species

1
1042228
subspecies
node11067.members.0.js

163742
tribe
node11068.members.0.js
33
1

47081
1
genus

species
node11070.members.0.js
47082
1

3877
20
genus

20
3880
node11072.members.0.js
species

3898
11
genus

10
node11074.members.0.js
species
57577

1
97006
species
node11075.members.0.js

2231384
67
clade

clade
2231385
67

163729
67
tribe

3869
67
genus

67
3871
node11080.members.0.js
species

order
219
3744

3487
44
family

44
3497
genus

node11084.members.0.js
species
3498
3

2
85232
node11085.members.0.js
species

node11086.members.0.js
species
981085
39

family
3481
5

1
node11088.members.0.js
genus
13408

genus
4
3482

species
node11090.members.0.js
3483
4

3608
3
family

325284
3
tribe

3
72171
genus

3
326968
species
node11094.members.0.js

3745
family
node11095.members.0.js
167
1

171638
node11096.members.0.js
subfamily
17
1

no rank
1176516
10

3764
10
genus

74649
species
node11099.members.0.js
10

tribe
6
721789

subtribe
6
1184124

genus
3746
6

6
57918
species

101020
node11104.members.0.js
subspecies
6

149
171637
subfamily

tribe
31
721805

node11107.members.0.js
genus
3754
8
31

2
species
node11108.members.0.js
102107

20
species
node11109.members.0.js
3755

species
node11110.members.0.js
42229
1

721813
node11111.members.0.js
tribe
118
2

genus
3
3766

3
species
node11113.members.0.js
225117

3749
genus
node11114.members.0.js
112
66

5
node11115.members.0.js
species
3752

604297
node11116.members.0.js
species
1

40
species
node11117.members.0.js
3750

1
193311
genus
node11118.members.0.js

86
3646
order

family
4004
36

36
4005
genus

4006
node11122.members.0.js
species
36

6
3683
family

genus
node11124.members.0.js
3684
2
5

1
species
node11125.members.0.js
196689

1
node11126.members.0.js
species
197924

1
78168
species
node11127.members.0.js

45183
1
genus

1
node11129.members.0.js
species
218843

family
112800
1

1
238080
tribe

genus
85204
1

85205
node11133.members.0.js
species
1

family
node11134.members.0.js
3977
1
25

235631
subfamily
node11135.members.0.js
21
1

tribe
14
235882

genus
14
3980

3981
node11138.members.0.js
species
14

235887
3
tribe

genus
3
3995

node11141.members.0.js
species
180498
3

235883
3
tribe

3
3982
genus

3983
species
node11144.members.0.js
3

3
235629
subfamily

3
235880
tribe

genus
3
3987

3
3988
node11148.members.0.js
species

18
3688
family

18
238069
tribe

1
40685
node11151.members.0.js
genus

3689
node11152.members.0.js
genus
17
4

9
3694
node11153.members.0.js
species

node11154.members.0.js
species
43335
1

1
80863
node11155.members.0.js
species

75702
node11156.members.0.js
species
2

495
2
3502
node11157.members.0.js
order

3514
44
family

genus
27
12989

27
species
node11160.members.0.js
176864

13450
17
genus

node11162.members.0.js
species
13451
17

16714
node11163.members.0.js
family
27
6

genus
3
13402

3
species
node11165.members.0.js
32201

5
18
node11166.members.0.js
genus
16718

species
node11167.members.0.js
2249226
7

6
species
node11168.members.0.js
51240

422
4
3503
node11169.members.0.js
family

380
17
3511
genus
node11170.members.0.js

species
node11171.members.0.js
3512
1

4
node11172.members.0.js
species
58330

node11173.members.0.js
species
97700
191

1
node11174.members.0.js
species
3513

species
node11175.members.0.js
38942
3

58331
node11176.members.0.js
species
163

genus
38
21024

38
species
node11178.members.0.js
28930

4
403666
order

43873
4
family

293141
4
subfamily

221920
node11182.members.0.js
genus
2

genus
2
66651

2
66652
species
node11184.members.0.js

order
71
71239

71
3650
family

19
1003878
tribe

3
19
genus
node11188.members.0.js
3660

node11189.members.0.js
species
3661
3

8
node11190.members.0.js
species
3662

species
3663
5

5
3664
subspecies
node11192.members.0.js

tribe
1003871
1

genus
3671
1

1
node11195.members.0.js
species
3673

1003877
51
tribe

51
1
3655
node11197.members.0.js
genus

40
3656
species
node11198.members.0.js

node11199.members.0.js
species
3659
10

order
6
233875

6
4305
family

genus
6
123484

458696
species
node11203.members.0.js
6

node11204.members.0.js
clade
91836
10
4136

3485
6
41937
order
node11205.members.0.js

30
3472
family
node11206.members.0.js
4011

1
4012
node11207.members.0.js
genus

263461
2
genus

269719
species
node11209.members.0.js
2

55512
genus
node11210.members.0.js
3337
75

node11211.members.0.js
species
55513
3252

3
289741
node11212.members.0.js
species

node11213.members.0.js
species
1540414
1

1
434238
species

434239
subspecies
node11215.members.0.js
1

434234
species
node11216.members.0.js
5

1
289765
genus

1
289766
species
node11218.members.0.js

genus
83
23461

83
29780
node11220.members.0.js
species

genus
43850
1

43851
species
node11222.members.0.js
1

node11223.members.0.js
genus
43860
4
6

1
80338
species
node11224.members.0.js

991123
species
node11225.members.0.js
1

43852
node11226.members.0.js
genus
8
2

species
node11227.members.0.js
690244
1

5
4013
species
node11228.members.0.js

1
node11229.members.0.js
genus
289806

171928
2
genus

171929
node11231.members.0.js
species
2

23513
7
family

subfamily
1728959
7

node11234.members.0.js
genus
2706
4
7

2711
node11235.members.0.js
species
3

43
41944
order

34
3931
family

1699513
subfamily
node11238.members.0.js
34
1

tribe
1699524
6

genus
3932
6

6
node11241.members.0.js
species
71139

1699523
4
tribe

4
1705102
clade

genus
4
178132

4
species
node11245.members.0.js
178133

tribe
23
1699522

178174
23
genus

219896
species
node11248.members.0.js
23

9
3928
family

9
22662
genus

9
node11251.members.0.js
species
22663

order
282
41938

282
3629
family

38
7
214909
node11254.members.0.js
subfamily

1
28
genus
node11255.members.0.js
3640

node11256.members.0.js
species
3641
27

genus
3
108869

108875
node11258.members.0.js
species
3

5
214915
subfamily

genus
5
66655

node11261.members.0.js
species
66656
5

214907
node11262.members.0.js
subfamily
239
6

genus
39
47614

39
47615
species
node11264.members.0.js

genus
15
47605

node11266.members.0.js
species
106335
15

179
47
3633
genus
node11267.members.0.js

63
34284
species
node11268.members.0.js

13
node11269.members.0.js
species
3635

node11270.members.0.js
species
29730
43

node11271.members.0.js
species
3634
3

species
node11272.members.0.js
29729
10

41943
1
order

4027
1
family

1
21556
genus

species
node11276.members.0.js
163685
1

order
3699
315

family
7
301454

clade
2768677
7

7
1168313
genus

28532
node11281.members.0.js
species
7

node11282.members.0.js
family
3700
7
300

tribe
28
981070

50451
28
genus

28
50452
node11285.members.0.js
species

981100
1
tribe

98005
1
genus

species
node11288.members.0.js
72664
1

981099
20
tribe

genus
13287
20

13288
species
node11291.members.0.js
20

tribe
node11292.members.0.js
980083
4
62

5
3718
genus

species
node11294.members.0.js
81985
5

3701
genus
node11295.members.0.js
44
3

22
3702
species
node11296.members.0.js

38785
node11297.members.0.js
species
12

species
81970
1

81971
node11299.members.0.js
subspecies
1

6
node11300.members.0.js
species
59689

9
71323
genus

9
species
node11302.members.0.js
90675

tribe
node11303.members.0.js
981071
5
182

3705
genus
node11304.members.0.js
137
85

15
19
species
node11305.members.0.js
3711

4
node11306.members.0.js
subspecies
51351

node11307.members.0.js
species
3708
10

node11308.members.0.js
species
3712
22
23

1
varietas
node11309.members.0.js
109376

40
3725
genus

40
node11311.members.0.js
species
3726

3647
8
family

3648
8
genus

8
3649
node11314.members.0.js
species

no rank
30
91834

order
403667
30

family
30
3602

tribe
30
2304100

3603
node11319.members.0.js
genus
30
3

29760
node11320.members.0.js
species
27

clade
node11321.members.0.js
4447
4
199521

199484
4
1437197
node11322.members.0.js
subclass

1
4667
order

family
1
4677

1
node11325.members.0.js
genus
4688

node11326.members.0.js
clade
4734
6
199453

6
199310
order
node11327.members.0.js
38820

node11328.members.0.js
family
4479
228
199264

359160
node11329.members.0.js
clade
1554
16

subfamily
147367
1189

tribe
147380
1189

subtribe
1189
1648021

28
1188
node11333.members.0.js
genus
4527

52545
species
node11334.members.0.js
14

4533
species
node11335.members.0.js
21

species
5
83307

5
node11337.members.0.js
varietas
110450

63629
species
node11338.members.0.js
16

3
83308
species
node11339.members.0.js

node11340.members.0.js
species
4530
263
1063

348
node11341.members.0.js
no rank
39946

39947
node11342.members.0.js
no rank
452

4536
species
node11343.members.0.js
1

node11344.members.0.js
species
4537
4

6
4535
node11345.members.0.js
species

4538
node11346.members.0.js
species
5

22
node11347.members.0.js
species
4532

genus
1
35711

1
2630782
no rank

node11350.members.0.js
species
1227981
1

332
4
147368
node11351.members.0.js
subfamily

no rank
2822797
26

tribe
26
147385

genus
15367
26

25
node11355.members.0.js
species
15368

1
species
node11356.members.0.js
29664

288
1648038
no rank

288
27
147389
node11358.members.0.js
tribe

34
213
subtribe
node11359.members.0.js
1648030

4480
18
genus

species
18
37682

200361
subspecies
node11362.members.0.js
18

161
39
4564
node11363.members.0.js
genus

species
1
4571

1
subspecies
node11365.members.0.js
4567

85692
species
node11366.members.0.js
12

109
species
node11367.members.0.js
4565

1648017
48
subtribe

15492
1
genus

1
129742
node11370.members.0.js
species

genus
1
4549

node11372.members.0.js
species
4550
1

genus
node11373.members.0.js
4512
1
46

2
45
node11374.members.0.js
species
4513

subspecies
node11375.members.0.js
112509
43

no rank
1648037
14

14
1
147387
node11377.members.0.js
tribe

7
1652081
clade

7
640630
subtribe

genus
4520
6

species
node11381.members.0.js
4522
6

genus
4605
1

species
node11383.members.0.js
4608
1

6
640623
subtribe

4496
6
genus

6
4498
species
node11386.members.0.js

2
17
node11387.members.0.js
subfamily
147366

no rank
3
1648035

147376
3
tribe

subtribe
1648004
3

genus
4581
2

species
node11392.members.0.js
58923
2

1
323892
genus

node11394.members.0.js
species
323893
1

1648034
12
no rank

tribe
1149709
12

12
1648003
subtribe

338513
1
genus

species
node11399.members.0.js
411665
1

11
1
15747
node11400.members.0.js
genus

10
38705
species
node11401.members.0.js

197482
29
147370
node11402.members.0.js
clade

197442
407
147369
node11403.members.0.js
subfamily

no rank
3
1699033

753695
3
genus

753696
node11406.members.0.js
species
3

no rank
1648036
5058

5058
284
147428
tribe
node11408.members.0.js

1765
1293360
subtribe

66017
node11410.members.0.js
genus
1765
1

1010633
node11411.members.0.js
species
1764

1293365
868
subtribe

4539
genus
node11413.members.0.js
868
43

section
2100771
602

node11415.members.0.js
species
38727
602

223
2100772
section

223
206008
node11417.members.0.js
species

subtribe
1
1293363

node11419.members.0.js
genus
45618
1

2139
1
1293361
subtribe
node11420.members.0.js

genus
4583
3

1
node11422.members.0.js
species
247021

2
4543
node11423.members.0.js
species

node11424.members.0.js
genus
4554
156
2135

node11425.members.0.js
species
4556
1935

44
node11426.members.0.js
species
4555

1
1293362
subtribe

37562
1
genus

1
species
node11429.members.0.js
240449

tribe
1293356
8

1
1293357
subtribe

1
158103
genus
node11432.members.0.js

subtribe
7
1293359

147271
node11434.members.0.js
genus
7
1

5
158149
node11435.members.0.js
species

151521
species
node11436.members.0.js
1

tribe
147443
13

4
909212
genus

4
2358305
node11439.members.0.js
species

796873
node11440.members.0.js
genus
9
1

species
8
1382009

node11442.members.0.js
varietas
2358306
8

no rank
1648033
191952

882
191952
node11444.members.0.js
tribe
147429

subtribe
node11445.members.0.js
2837560
1

3
7
subtribe
node11446.members.0.js
1647998

genus
1
15314

330559
species
node11448.members.0.js
1

2
79854
genus

1393007
species
node11450.members.0.js
1

1
2491244
node11451.members.0.js
species

genus
1
1699088

species
node11453.members.0.js
2837563
1

182679
1648028
subtribe

119
182679
node11455.members.0.js
genus
4557

1
91525
species
node11456.members.0.js

132711
species
node11457.members.0.js
96

4560
node11458.members.0.js
species
12

node11459.members.0.js
species
29704
2

node11460.members.0.js
species
4558
182439
182441

node11461.members.0.js
subspecies
171959
2

node11462.members.0.js
species
29670
3

5
node11463.members.0.js
species
91528

5581
6
1648026
node11464.members.0.js
subtribe

3
23
genus
node11465.members.0.js
62336

node11466.members.0.js
species
183675
2

species
node11467.members.0.js
154761
2

species
node11468.members.0.js
200567
1

node11469.members.0.js
species
183674
2

1
74665
species
node11470.members.0.js

10
12
node11471.members.0.js
species
62337

2
1068733
subspecies

213995
node11473.members.0.js
varietas
2
1

1
91809
node11474.members.0.js
forma

genus
4546
5552

5552
242
286192
node11476.members.0.js
no rank

4314
116
128810
node11477.members.0.js
species

node11478.members.0.js
no rank
50502
5

1
node11479.members.0.js
no rank
2767070

193079
no rank
node11480.members.0.js
700

676074
no rank
node11481.members.0.js
1

1214490
no rank
node11482.members.0.js
5

node11483.members.0.js
no rank
130727
3

1
672245
no rank
node11484.members.0.js

2
672234
node11485.members.0.js
no rank

no rank
node11486.members.0.js
672404
1

3478
node11487.members.0.js
no rank
131158

672237
no rank
node11488.members.0.js
1

node11489.members.0.js
species
62335
517

4547
species
node11490.members.0.js
479

subtribe
2837557
5

genus
79830
5

167337
node11493.members.0.js
species
5

subtribe
5
1648018

genus
node11495.members.0.js
798292
1

673199
4
genus

node11497.members.0.js
species
673200
4

1699106
15
no rank

1612336
15
genus

15
50346
node11500.members.0.js
species

5
2706
node11501.members.0.js
subtribe
1648029

4
4562
genus

4563
node11503.members.0.js
species
4

4575
genus
node11504.members.0.js
2697
3

2645
2694
node11505.members.0.js
species
4577

subspecies
node11506.members.0.js
76912
1

48
subspecies
node11507.members.0.js
381124

subtribe
7
1647999

79826
4
genus

node11510.members.0.js
species
435779
4

1
66013
genus

1
node11512.members.0.js
species
79835

1
2
node11513.members.0.js
genus
66042

1
species
node11514.members.0.js
2152742

57
1648025
subtribe

4504
56
genus

56
species
node11517.members.0.js
4505

genus
1
300124

1
300125
species
node11519.members.0.js

2837558
1
subtribe

1
798297
genus

945804
species
node11522.members.0.js
1

1648014
6
subtribe

genus
80368
4

4
node11525.members.0.js
species
80369

genus
167340
1

167341
node11527.members.0.js
species
1

1
798272
genus

1
node11529.members.0.js
species
798273

147426
1
tribe

genus
1
66007

node11532.members.0.js
species
66008
1

147371
10
subfamily

tribe
1
751754

subtribe
1
751756

38730
1
genus

29706
node11537.members.0.js
species
1

tribe
1080378
1

1
42045
genus

1
42046
species
node11540.members.0.js

8
147435
tribe

4
1
751762
node11542.members.0.js
subtribe

3
15437
genus

3
species
node11544.members.0.js
1920021

751759
1
subtribe

genus
66044
1

1
1245684
node11547.members.0.js
species

1
751763
subtribe

genus
796752
1

1148796
species
node11550.members.0.js
1

2
751757
no rank

genus
2
1465621

species
node11553.members.0.js
1465622
2

subfamily
1
318921

1
153999
tribe

66023
1
genus

species
node11557.members.0.js
883138
1

family
36
4613

subfamily
1909378
36

36
4614
genus

25
36
node11561.members.0.js
species
4615

296719
varietas
node11562.members.0.js
11

family
4
4609

986140
4
subfamily

3
986145
tribe

genus
3
4610

2034327
3
clade

1982031
3
subgenus

no rank
2034351
3

2
node11570.members.0.js
species
1053340

1
4611
species
node11571.members.0.js

tribe
986143
1

46324
genus
node11573.members.0.js
1

104
1
4618
node11574.members.0.js
order

family
4637
79

79
4
4640
node11576.members.0.js
genus

4641
node11577.members.0.js
species
74
26

48
subspecies
node11578.members.0.js
214687

1
species
node11579.members.0.js
52706

family
24
4642

genus
24
4650

94328
node11582.members.0.js
species
24

order
33
40551

family
node11584.members.0.js
4710
1
33

subfamily
15
169700

169748
15
tribe

15
4719
genus

node11588.members.0.js
species
42345
15

17
169697
subfamily

tribe
169705
17

169725
3
subtribe

genus
13893
3

3
node11593.members.0.js
species
13894

subtribe
14
169729

genus
51952
14

14
species
node11596.members.0.js
51953

order
73496
19

5
4668
family

40553
5
subfamily

703248
5
tribe

node11601.members.0.js
genus
4678
1
5

node11602.members.0.js
species
4679
4

family
9
4747

subfamily
9
158332

tribe
158393
5

subtribe
158406
5

5
37818
genus

906689
species
node11608.members.0.js
5

tribe
3
158397

158424
3
subtribe

genus
36459
3

node11612.members.0.js
species
78828
3

1
158389
tribe

1
158399
subtribe

1
142980
genus

1
646075
node11616.members.0.js
species

5
40552
family

subfamily
5
703533

4685
5
genus

species
node11620.members.0.js
4686
5

40548
5
order

family
5
4671

genus
5
4672

5
29710
species

5
subspecies
node11625.members.0.js
55577

40550
2
order

25913
1
family

genus
1
2707182

1
species
node11629.members.0.js
2707183

family
1
49662

1
genus
node11631.members.0.js
85281

33
16360
order

1
16362
family

246706
1
genus

node11635.members.0.js
species
55444
1

family
32
4454

subfamily
32
284551

genus
4473
32

29656
species
node11639.members.0.js
7

51605
node11640.members.0.js
species
25

order
261007
10

4410
10
family

genus
10
4418

10
210225
species
node11644.members.0.js

213
1437180
clade

2
3372
class

subclass
2
1445966

2
3378
order

family
2
3379

2
3380
genus

1
3382
species
node11651.members.0.js

3381
node11652.members.0.js
species
1

class
29811
1

1
1445964
subclass

3308
1
order

1
3309
family

genus
1
3310

3311
species
node11658.members.0.js
1

class
210
58019

210
3313
subclass

13
2821351
clade

1446379
13
order

family
node11663.members.0.js
3367
1
13

genus
3368
9

species
node11665.members.0.js
3369
9

25613
3
genus

species
node11667.members.0.js
50187
3

clade
2821352
197

order
197
1446380

2
197
family
node11670.members.0.js
3318

1
genus
node11671.members.0.js
3321

120
29
3328
genus
node11672.members.0.js

35
3332
node11673.members.0.js
species

56
species
node11674.members.0.js
3330

genus
3325
1

54800
species
node11676.members.0.js
1

node11677.members.0.js
genus
3337
2
73

2
139272
subgenus

node11679.members.0.js
species
3342
1

1
node11680.members.0.js
species
3348

14
69
subgenus
node11681.members.0.js
139271

2
3347
species
node11682.members.0.js

2
species
node11683.members.0.js
3353

3352
species
node11684.members.0.js
17

1
3339
species
node11685.members.0.js

1
node11686.members.0.js
species
3346

14
species
node11687.members.0.js
71647

88731
node11688.members.0.js
species
9

88730
species
node11689.members.0.js
2

2
71633
species
node11690.members.0.js

3
3349
species
node11691.members.0.js

71636
species
node11692.members.0.js
2
1

node11693.members.0.js
varietas
1504333
1

class
131220
4

3172
4
order

3173
4
family

4
2
3174
node11697.members.0.js
genus

species
node11698.members.0.js
105231
1

1
species
node11699.members.0.js
3175

node11700.members.0.js
phylum
3041
1
104

1
33103
class

2546215
1
clade

order
1
31306

2682561
1
family

1
160075
genus

node11706.members.0.js
species
2202518
1

77633
1
no rank

171245
species
node11708.members.0.js
1

2692248
clade
node11709.members.0.js
84
2

35
5
75966
class
node11710.members.0.js

14
75981
no rank

2
2511081
clade

2
2304164
genus

2704667
2
no rank

species
node11715.members.0.js
2704670
1

node11716.members.0.js
species
2704668
1

genus
1293077
2

2
species
node11718.members.0.js
1293078

clade
2511161
10

41891
10
genus

no rank
2688356
8

8
2315456
node11722.members.0.js
species

2
248742
species

isolate
node11724.members.0.js
574566
2

7
135250
order

family
5
2910630

5
29646
genus

37433
species
node11728.members.0.js
5

family
135266
2

1
135261
genus

species
node11731.members.0.js
173492
1

genus
1
1940604

1
species
node11733.members.0.js
3083

42111
1
order

family
2682484
1

genus
1
106202

106203
node11737.members.0.js
species
1

order
35460
4

4
35461
family

3187
1
genus

node11741.members.0.js
species
926285
1

3
191392
genus

3
3075
species
node11743.members.0.js

4
2507901
order

2507902
4
family

13786
1
genus

node11747.members.0.js
species
103878
1

genus
114064
3

species
node11749.members.0.js
3171
3

3166
47
class

order
1
138177

family
1
2682464

1
138174
genus

species
node11754.members.0.js
332213
1

46
2812636
clade

order
3042
43

family
3065
10

9
3066
genus

9
3067
species

node11760.members.0.js
forma
3068
9

33098
1
genus

33099
species
node11762.members.0.js
1

family
11
3051

genus
11
3052

1
2034146
no rank

1653778
node11766.members.0.js
species
1

node11767.members.0.js
species
3055
9

3054
node11768.members.0.js
species
1

family
3043
21

21
1
3044
node11770.members.0.js
genus

20
species
node11771.members.0.js
257627

77634
1
no rank

genus
1
56010

node11774.members.0.js
species
56011
1

3
35491
order

family
3
35466

genus
34111
3

3
species
node11778.members.0.js
145388

14
1035538
class

14
13792
order

family
11
1525212

4
70447
genus

species
node11783.members.0.js
70448
4

41874
7
genus

7
node11785.members.0.js
species
41875

3
41873
family

genus
3
38832

node11788.members.0.js
species
296587
2

species
38833
1

1
564608
strain
node11790.members.0.js

class
2302911
3

2302912
3
order

3
2302913
family

genus
1
2302916

1883388
node11795.members.0.js
species
1

2
2302914
genus

1764295
species
node11797.members.0.js
2

clade
1
2611341

207245
1
phylum

1
5738
order

family
1
5739

1
68459
subfamily

1
5740
genus

5741
node11804.members.0.js
species
1

clade
node11805.members.0.js
554915
1
54

30
555280
phylum

30
1485168
order

clade
555407
30

family
30
33677

5754
node11810.members.0.js
genus
30
5

node11811.members.0.js
species
5755
8
25

node11812.members.0.js
strain
1257118
17

phylum
22
2605435

clade
555406
8

order
8
2682482

33084
8
family

genus
5758
8

46681
2
species

370354
strain
node11819.members.0.js
2

species
33085
5

5
370355
strain
node11821.members.0.js

1
5759
species
node11822.members.0.js

class
142796
13

clade
33083
6

order
6
2058181

2058184
2
family

2058187
2
genus

261658
species
node11828.members.0.js
2

family
4
2058183

133407
2
genus

species
2
361139

strain
node11832.members.0.js
1410327
2

2058189
2
genus

2
2086695
species

2
670386
node11835.members.0.js
strain

node11836.members.0.js
clade
33680
1
7

subclass
137627
5

5
5789
order

node11839.members.0.js
family
1115744
1
3

5790
2
genus

species
node11841.members.0.js
5791
1

289951
species
node11842.members.0.js
1

family
2
1115745

5792
2
genus

1
372585
species
node11845.members.0.js

181200
species
node11846.members.0.js
1

2682238
1
clade

order
1
208471

family
208472
1

1092570
1
genus

2715192
node11851.members.0.js
species
1

2605334
1
class

family
1
2605335

490353
1
genus

490516
node11855.members.0.js
species
1

no rank
1
590480

species
node11857.members.0.js
590481
1

clade
61
2611352

5752
12
phylum

2601529
12
clade

12
2601530
clade

10
5765
family

2
10
genus
node11863.members.0.js
5761

species
node11864.members.0.js
51637
1

5762
species
node11865.members.0.js
2

5763
node11866.members.0.js
species
5

2
1144924
family

genus
1217108
2

2633603
2
no rank

node11870.members.0.js
species
500012
2

33682
49
phylum

5653
49
class

subclass
2704647
49

2704949
49
order

family
49
5654

5690
25
genus

39700
subgenus
node11877.members.0.js
5
4

1
node11878.members.0.js
species
5691

1
47571
subgenus

species
1
5699

1
strain
node11881.members.0.js
1055687

subgenus
47570
5

1
5
node11883.members.0.js
species
5693

353153
strain
node11884.members.0.js
3

subspecies
node11885.members.0.js
85057
1

669453
12
clade

5
species
node11887.members.0.js
71804

4
67003
node11888.members.0.js
species

3
species
node11889.members.0.js
83891

subgenus
2
39701

species
node11891.members.0.js
5698
2

1581334
2
subfamily

genus
1003337
2

species
node11894.members.0.js
59799
2

22
1286322
subfamily

5
5683
genus

5
157538
node11897.members.0.js
species

5658
17
genus

node11899.members.0.js
subgenus
37616
3
5

2
node11900.members.0.js
species group
37617

1
12
subgenus
node11901.members.0.js
38568

2
4
node11902.members.0.js
species group
38574

2
species
node11903.members.0.js
5661

7
5
38582
node11904.members.0.js
species group

1
species
node11905.members.0.js
5659

species
1
5665

929439
node11907.members.0.js
strain
1

2698737
node11908.members.0.js
clade
11584
7

11451
4
33634
node11909.members.0.js
clade

4762
phylum
node11910.members.0.js
11407
83

order
10970
4776

316
10970
family
node11912.members.0.js
4777

401
5331
genus
node11913.members.0.js
4783

4787
species
node11914.members.0.js
1363
36

1327
strain
node11915.members.0.js
403677

1
1692154
node11916.members.0.js
species

node11917.members.0.js
species
360399
5

node11918.members.0.js
species
1642459
2

1
species
node11919.members.0.js
4786

2
129354
node11920.members.0.js
species

299392
node11921.members.0.js
species
3

5
1642464
species
node11922.members.0.js

1
111171
species
node11923.members.0.js

374175
species
node11924.members.0.js
2

species
4792
1548

1548
761204
node11926.members.0.js
strain

6
node11927.members.0.js
species
360400

278
4784
node11928.members.0.js
species

species
node11929.members.0.js
4785
7

2
node11930.members.0.js
species
638948

4
node11931.members.0.js
species
164328

89335
species
node11932.members.0.js
1

node11933.members.0.js
species
53987
1

283007
node11934.members.0.js
species
2

species
node11935.members.0.js
905063
1

555429
node11936.members.0.js
species
5

67593
species
node11937.members.0.js
1678

5
4796
node11938.members.0.js
species

1
100870
species
node11939.members.0.js

no rank
211524
6

1
species
node11941.members.0.js
458840

89336
species
node11942.members.0.js
3

907715
species
node11943.members.0.js
2

184462
326
genus

7
2
272952
species
node11945.members.0.js

5
strain
node11946.members.0.js
559515

319
453155
species group

319
123356
node11948.members.0.js
species

4780
453
genus

162139
species
node11950.members.0.js
2

species
node11951.members.0.js
162141
1

162140
species
node11952.members.0.js
2

8
143451
species
node11953.members.0.js

node11954.members.0.js
species
4781
440

4369
70742
genus

2
node11956.members.0.js
species
230439

4365
node11957.members.0.js
species
542832

1
species
node11958.members.0.js
622444

1
2028485
node11959.members.0.js
species

3
143452
node11960.members.0.js
genus

2
164
genus
node11961.members.0.js
230838

158
230839
node11962.members.0.js
species

1
467161
species
node11963.members.0.js

2
886949
node11964.members.0.js
species

2624089
1
no rank

node11966.members.0.js
species
1487174
1

1
8
node11967.members.0.js
genus
4778

7
node11968.members.0.js
species
4779

order
67
370421

family
65355
67

genus
65356
67

node11972.members.0.js
species
653948
57
67

10
isolate
node11973.members.0.js
890382

4763
284
order

284
2
4764
node11975.members.0.js
family

genus
node11976.members.0.js
100860
8
126

157072
species
node11977.members.0.js
42

node11978.members.0.js
species
112090
67

species
node11979.members.0.js
100861
8

601994
node11980.members.0.js
species
1

150
30
4769
node11981.members.0.js
genus

112098
59
species

node11983.members.0.js
strain
1156394
59

species
101203
61

695850
strain
node11985.members.0.js
61

genus
2
4765

species
node11987.members.0.js
1202772
2

genus
4
74556

species
node11989.members.0.js
74557
4

121069
1
order

family
4782
1

1
1448052
genus

82942
species
node11993.members.0.js
1

2
4800
order

no rank
2
2041418

node11996.members.0.js
genus
1440114
2

46625
1
no rank

1
2795716
clade

2805718
node11999.members.0.js
species
1

class
5
2683628

5
2683629
clade

42740
5
order

family
2547934
5

12967
5
genus

clade
3
944171

3
species
node12006.members.0.js
944036

species
node12007.members.0.js
12968
2

34
2696291
clade

33859
2
class

98652
2
order

family
2
88165

genus
88166
2

2
88167
node12013.members.0.js
species

class
35675
2

order
54409
2

genus
2
44055

2
44056
species
node12017.members.0.js

1
39119
class

35680
1
order

genus
35683
1

35684
species
node12021.members.0.js
1

4
11
class
node12022.members.0.js
5747

order
425074
7

425072
7
family

genus
5748
7

species
node12026.members.0.js
145522
7

2836
18
phylum

9
33849
class

9
1
33850
node12029.members.0.js
clade

order
3
33851

family
node12031.members.0.js
33852
2
3

genus
2857
1

1
908989
no rank

1
1444690
species
node12034.members.0.js

order
38748
5

family
1
265535

genus
1242273
1

species
node12038.members.0.js
2305497
1

family
38749
3

3
2849
genus

3
2850
node12041.members.0.js
species

family
1
303454

1
431369
genus
node12043.members.0.js

class
2
33853

2
33854
subclass

order
1
33855

1
33856
family

35129
1
genus

1
species
node12049.members.0.js
210441

1
426665
order

1
38755
family

genus
33648
1

1
186022
node12053.members.0.js
species

class
33836
7

3
420266
subclass

3
2
265576
order
node12056.members.0.js

1
49236
family

49237
1
genus

node12059.members.0.js
species
2082246
1

4
33846
subclass

33847
4
order

family
29202
4

35127
4
genus

35128
4
species

4
node12065.members.0.js
strain
296543

118
33630
clade

phylum
5794
86

class
50
422676

9
5863
order

6
27994
family

genus
6
5873

2
5874
species
node12072.members.0.js

2
4
node12073.members.0.js
species
68886

node12074.members.0.js
strain
869250
2

32594
3
family

3
5864
genus

3
5868
species

3
node12078.members.0.js
strain
1133968

order
5819
41

1639119
41
family

41
1
5820
genus
node12081.members.0.js

subgenus
418104
2

2
5849
node12083.members.0.js
species

subgenus
418103
14

2
5827
species
node12085.members.0.js

1
species
node12086.members.0.js
5857

2
species
node12087.members.0.js
5855

2
node12088.members.0.js
species
5858

77519
species
node12089.members.0.js
5

1
5850
node12090.members.0.js
species

1
52288
species

1
strain
node12092.members.0.js
1237626

1
208452
species
node12093.members.0.js

8
18
node12094.members.0.js
subgenus
418101

4
5
node12095.members.0.js
species
5860

1
subspecies
node12096.members.0.js
138298

species
2
5825

5826
subspecies
node12098.members.0.js
2

species
node12099.members.0.js
5861
1

node12100.members.0.js
species
5821
2

3
5
subgenus
node12101.members.0.js
418107

node12102.members.0.js
species
647221
1

880535
node12103.members.0.js
species
1

class
36
1280412

subclass
36
5796

order
75739
36

36
423054
suborder

1
35082
family

1
5806
genus

1
857276
species
node12110.members.0.js

5809
13
family

5810
6
genus

6
3
5811
node12113.members.0.js
species

398031
3
biotype

3
strain
node12115.members.0.js
432359

2
94642
genus

94643
species
node12117.members.0.js
2

genus
29175
5

species
5
29176

572307
node12120.members.0.js
strain
5

5799
22
family

genus
5800
22

16
5802
species
node12123.members.0.js

1
5804
species
node12124.members.0.js

node12125.members.0.js
species
51315
1

species
node12126.members.0.js
5801
1

44415
node12127.members.0.js
species
3

2497438
5
phylum

clade
27997
5

27998
5
order

5
27999
family

5
28000
genus

31276
5
species

423536
node12134.members.0.js
strain
5

27
5878
phylum

27
431838
subphylum

class
658449
1

2213231
1
order

1
37952
family

70074
1
genus

1
node12141.members.0.js
species
70075

6020
23
class

order
31277
14

suborder
37093
13

13
291294
family

13
5890
genus

species
13
5911

312017
strain
node12148.members.0.js
13

suborder
1
37090

genus
5931
1

1
5932
species
node12151.members.0.js

order
9
33825

family
340080
9

genus
9
5884

9
5888
species

strain
node12156.members.0.js
412030
9

33829
3
class

1
3
subclass
node12158.members.0.js
194286

33832
1
order

family
1
99916

genus
1
331615

2678605
node12162.members.0.js
species
1

693921
1
order

1
57506
family

subfamily
1001748
1

genus
94288
1

1
94289
node12167.members.0.js
species

clade
543769
8

136419
phylum
node12169.members.0.js
7
2

order
188941
2

family
2
45105

genus
2
45106

2
45107
species
node12173.members.0.js

no rank
2
243290

175275
species
node12175.members.0.js
2

order
1238681
1

family
2586441
1

genus
1
1239209

no rank
2630903
1

1
1979791
species
node12180.members.0.js

clade
2662056
1

class
65582
1

order
node12183.members.0.js
65583
1

clade
4
554296

4
172820
family

genus
4
877559

species
4
529818

4
461836
node12188.members.0.js
strain

2157
node12189.members.0.js
superkingdom
185
1

2283796
6
phylum

class
183967
6

2301
4
order

1919231
3
no rank

species
node12194.members.0.js
2268204
1

species
node12195.members.0.js
1054217
2

90142
1
family

node12197.members.0.js
genus
74968
1

2
1234666
no rank

1906666
species
node12199.members.0.js
2

phylum
28890
156

clade
117
2290931

75
183963
class

42
2235
order

family
8
1963268

genus
4
146825

no rank
2621901
4

species
node12207.members.0.js
1380432
4

1
203135
genus

1
2610901
no rank

1
2610902
node12210.members.0.js
species

genus
2
63743

1710540
node12212.members.0.js
species
2

1
genus
node12213.members.0.js
171163

family
24
2236

genus
3
367188

no rank
3
2622732

1
node12217.members.0.js
species
2884876

2
2884875
node12218.members.0.js
species

2239
7
genus

node12220.members.0.js
species
2039234
1

5
223182
species
node12221.members.0.js

1407499
species
node12222.members.0.js
1

14
1075398
genus

1
species
node12224.members.0.js
555573

2648404
13
no rank

species
node12226.members.0.js
2599399
13

1963280
10
no rank

2732368
10
genus

10
1710541
species
node12229.members.0.js

14
1644060
order

1644061
14
family

1
2765402
genus

1
745377
species
node12233.members.0.js

genus
121871
2

2631942
2
no rank

1
node12236.members.0.js
species
1699371

2816475
node12237.members.0.js
species
1

genus
1
1269201

no rank
1
2621923

1333523
species
node12240.members.0.js
1

genus
1
387342

1
387343
species

1
797210
strain
node12243.members.0.js

genus
134813
1

61858
species
node12245.members.0.js
1

genus
88723
7

node12247.members.0.js
species
69527
1

6
2622230
no rank

node12249.members.0.js
species
1710539
1

1
node12250.members.0.js
species
2878535

node12251.members.0.js
species
2871694
4

332951
1
genus

no rank
1
2633955

2817025
node12254.members.0.js
species
1

order
19
1644055

1963271
15
family

1644057
3
genus

species
node12258.members.0.js
755307
3

1450140
1
genus

1
1073996
node12260.members.0.js
species

1075397
1
genus

2634974
1
no rank

1853690
node12263.members.0.js
species
1

43927
8
genus

2743090
species
node12265.members.0.js
2

1
2872158
node12266.members.0.js
species

5
species
node12267.members.0.js
2743089

56688
2
genus

2247
1
species

416348
strain
node12270.members.0.js
1

node12271.members.0.js
species
35743
1

family
4
1644056

1911573
2
genus

no rank
2
2629969

2
node12275.members.0.js
species
2876193

2
1073986
genus

699433
species
node12277.members.0.js
2

class
42
224756

2
2191
order

family
1
88404

2192
1
genus

species
83984
1

1
strain
node12283.members.0.js
410358

family
2194
1

45989
1
genus

83986
species
node12286.members.0.js
1

order
38
94695

family
1392996
2

2
no rank
node12289.members.0.js
2013823

34
2206
family

2225
5
genus

no rank
4
2618147

4
2822137
species
node12293.members.0.js

species
29291
1

1
node12295.members.0.js
strain
259564

2207
node12296.members.0.js
genus
7
2

170861
2
species

2
1434111
node12298.members.0.js
strain

1
3
species
node12299.members.0.js
2208

1
1434106
node12300.members.0.js
strain

1
strain
node12301.members.0.js
1434107

genus
22
2220

22
536044
species
node12303.members.0.js

2
588815
no rank

2
2759911
node12305.members.0.js
species

order
2905377
2

family
143067
2

2222
genus
node12308.members.0.js
2
1

no rank
1
2620051

1
90426
node12310.members.0.js
species

class
2545688
1

order
2545689
1

family
1
2545690

genus
2545692
1

2874847
1
no rank

1
node12316.members.0.js
species
2874846

183968
14
class

order
14
2258

family
14
2259

genus
10
2263

species
6
110164

node12322.members.0.js
strain
1432656
6

no rank
2627626
2

1
species
node12324.members.0.js
758583

1
node12325.members.0.js
species
122420

1
node12326.members.0.js
species
187880

71998
species
node12327.members.0.js
1

genus
2260
1

species
1
29292

1
272844
node12330.members.0.js
strain

83867
3
genus

971279
3
species

3
1343739
strain
node12333.members.0.js

1
183988
class

1
68985
order

family
1
183713

1
2319
genus

1
2684913
node12338.members.0.js
no rank

22
2283794
clade

class
183925
22

order
2158
22

family
2159
22

genus
4
2160

1
node12344.members.0.js
species
118062

1
node12345.members.0.js
species
2162

no rank
2627676
2

species
node12347.members.0.js
2025351
1

species
node12348.members.0.js
2025350
1

genus
18
2172

node12350.members.0.js
species
2173
17

node12351.members.0.js
species
230361
1

no rank
68359
1

1
1456563
node12353.members.0.js
species

no rank
1
93506

1906665
species
node12355.members.0.js
1

clade
1783276
1

1
1801631
phylum

2490204
1
genus

1
1920749
species
node12359.members.0.js

clade
1
1935183

1936272
1
phylum

node12362.members.0.js
species
2026747
1

48510
1
no rank

species
node12364.members.0.js
743098
1

clade
18
1783275

phylum
3
651137

no rank
node12367.members.0.js
651140
2

31932
1
order

1
371948
no rank
node12369.members.0.js

13
28889
phylum

183924
11
class

2266
2
order

2
2267
family

1
2270
genus

species
1
184117

999630
strain
node12376.members.0.js
1

2276
1
genus

species
1
2277

1
384616
node12379.members.0.js
strain

order
7
2281

family
118883
7

2
69655
genus

node12383.members.0.js
species
69656
2

2
12914
genus

species
2
563177

933801
strain
node12386.members.0.js
2

genus
3
2100760

1
node12388.members.0.js
species
2286

2287
node12389.members.0.js
species
2

2
114380
order

family
2
2272

477695
1
genus

species
1200300
1

1184251
strain
node12394.members.0.js
1

genus
54253
1

species
1
54254

strain
node12397.members.0.js
633148
1

48509
2
no rank

2
node12399.members.0.js
species
29281

phylum
928852
2

1700837
2
no rank

2
node12402.members.0.js
species
2026714

14
2787854
no rank

14
28384
no rank

14
2
81077
node12405.members.0.js
no rank

no rank
node12406.members.0.js
29278
5
7

1
45778
no rank

node12408.members.0.js
species
239484
1

species
node12409.members.0.js
2676029
1

no rank
1
111786

species
node12411.members.0.js
111789
1

32630
species
node12412.members.0.js
4
